# Disruption of autism-associated *Pcdh9* gene leads to transcriptional alterations, synapses overgrowth and aberrant excitatory transmission in the CA1

**DOI:** 10.1101/2024.01.27.577576

**Authors:** Federico Miozzo, Luca Murru, Greta Maiellano, Antonio Zippo, Edoardo Moretto, Annalaura Zambrano Avendano, Verjinia D. Metodieva, Sara Riccardi, Deborah D’Aliberti, Silvia Spinelli, Tamara Canu, Linda Chaabane, Shinji Hirano, Martien J. H. Kas, Maura Francolini, Rocco Piazza, Maria Passafaro

## Abstract

Protocadherins are cell adhesion molecules with crucial role in cell-cell contacts, whose mutations or altered expression have been implicated in multiple brain disorders. In particular, growing evidence links genetic alterations in *Protocadherin 9* (*PCDH9*) gene with Autism Spectrum Disorder (ASD) and Major Depression Disorder (MDD). Furthermore, *Pcdh9* deletion induces neuronal defects in the mouse somatosensory cortex, accompanied by sensorimotor and memory impairment. However, the synaptic and molecular mechanisms underlying *Pcdh9* physiological function and its involvement in brain pathology remain largely unknown. To this aim, we conducted a comprehensive investigation of PCDH9 role in the mouse hippocampus at the ultrastructural, biochemical, transcriptomic, electrophysiological and network level. We show that PCDH9 mainly localizes at glutamatergic synapses and its expression peaks in the first week after birth, a crucial time window for synaptogenesis. Strikingly, *Pcdh9* KO neurons exhibit oversized presynaptic terminal and postsynaptic density (PSD) in the CA1. Synapse overgrowth is sustained by the broad up-regulation of synaptic genes and the dysregulation of key drivers of synapse morphogenesis, as revealed by single-nucleus RNAseq. Synaptic and transcriptional defects are accompanied by increased EPSC frequency and disturbances in the hippocampal network activity of *Pcdh9* KO mice. In conclusion, our work indicates that *Pcdh9* regulates the morphology and function of excitatory synapses in the CA1, thereby affecting glutamatergic transmission in hippocampal circuitries.

## INTRODUCTION

Synaptic connections among the approximately 90 billion neurons of the human brain must be established during neurodevelopment to give rise to functional neuronal circuitries. Failures during this process, leading to improper neuronal networks, result in neurodevelopmental and neuropsychiatric disorders. Cell adhesion molecules (CAMs) present on neuronal cell surface take a key part in neurodevelopment, regulating cell-cell recognition, cell adhesion and migration (*1*). Among CAMs, protocadherin (PCDH) family counts more than 70 single membrane-spanning glycoproteins, representing the largest subgroup of the cadherin superfamily. PCDHs play critical roles in major neurodevelopmental processes, including neurite outgrowth, axon pathfinding, synapse formation and maturation (*2*). Indeed, growing evidence links genetic alterations in *PCDH* genes to a wide range of brain pathologies (*3*, *4*).

*Protocadherin 9* (*PCDH9*) gene has been associated with ASD after the identification of *de novo* and inherited copy number variations (CNVs) in autistic individuals (*5*, *6*). Furthermore, decreased *PCDH9* transcript levels were found in lymphoblasts of ASD patients (*7*). Interestingly, meta-analysis of three genome-wide association studies (GWAS) linked the single nucleotide polymorphism (SNP) *rs9540720* in *PCDH9* gene with MDD and cognitive function impairment (*8*). *rs9540720* is predicted to lower *PCDH9* expression, and reduced *PCDH9* mRNA levels were found in MDD patients compared with healthy controls, thus leading to the identification of *PCDH9* as a susceptibility locus for MDD (*8*). A recent study also associated *PCDH9* with Essential Tremor, a common movement disorder which is also characterized by cognitive and neuropsychiatric features (*9*).

*Pcdh9* KO mice display long-term social and object recognition deficits, hyperactivity and reduced sensorimotor competence. These behavioral abnormalities are accompanied by reduced cortical thickness of the somatosensory cortex, defective dendritic arborization and increased spine density of pyramidal neurons in these regions (*10*). A second study on a distinct *Pcdh9* KO line revealed further behavioral impairments including reduced fear extinction, possibly due to defects in *Ppp1r1b^+^* neurons of the basolateral amygdala (*11*). These investigations proved *Pcdh9* involvement in cognitive, sensory and emotional functions, thus strengthening the link with neurodevelopmental and psychiatric disorders. However, very little is known about the physiological role and molecular mechanisms of *Pcdh9* in the brain, and how its alteration relates to synaptic dysfunction and brain pathology.

Here, we examined *Pcdh9* levels and subcellular localization in the mouse brain. Then, we employed a combination of morphological, transcriptional and electrophysiological approaches to study *Pcdh9* function in the hippocampus *in vivo*. We show that *Pcdh9* depletion leads to aberrant pre- and postsynaptic morphology in the CA1, accompanied by dysregulation of synaptic genes expression. At the functional level, these alterations translate into abnormal excitatory transmission and disturbances in the hippocampal network activity. Collectively, our findings establish an important role for *Pcdh9* in the formation and function of CA1 glutamatergic synapses.

## RESULTS

### PCDH9 is enriched at glutamatergic synapses and its expression peaks in the postnatal hippocampus

To examine *Pcdh9* function in vivo, we first determined its protein levels in the adult mouse brain by western blot (WB) analysis. PCDH9 is widely present in the brain areas analysed, with a more prominent expression in cortex and hippocampus, and virtually absent in the other organs assessed, supporting a specific function for PCDH9 in the central nervous system (Fig. 1A, B). Then, we prepared hippocampal lysates from mice at different ages to study PCDH9 expression during neurodevelopment. We found that PCDH9 expression sharply increases at P7 and then progressively decreases and maintains stable in the adult mouse (Fig. 1C, D). The first three postnatal weeks are a critical time window for synapses formation and maturation (*28*), therefore PCDH9 up-regulation in this time window suggests a potential role in synaptogenesis. To study PCDH9 subcellular distribution, we performed cortex fractionation and mainly detected PCDH9 in the membrane-enriched fraction, as expected for a transmembrane protein (Fig. 1E). In order to investigate PCDH9 localization at the synapses, we coimmunostained DIV18 cultures of cortical and hippocampal neurons for PCDH9 and presynaptic (VGLUT1, excitatory; VGAT1, inhibitory) or postsynaptic markers (PSD95, excitatory; GEPHYRIN, inhibitory) (Fig. 1F). PCDH9 was moderately enriched in the perinuclear region and clearly present in the neurites. Colocalization analysis (Fig. 1G, H) revealed a broad overlap of PCDH9 and VGLUT1 signals, a moderate PCDH9 colocalization with VGAT1 and PSD95, and largely exclusive profiles with GEPHYRIN, indicating that PCDH9 is mainly present at excitatory synapses, predominantly in the presynaptic compartment. Taken together, these results indicate that PCDH9 is widely present in the brain, with a peak of expression in the hippocampus during the first postnatal weeks, and mainly localizes at excitatory synapses.

**Fig. 1.**
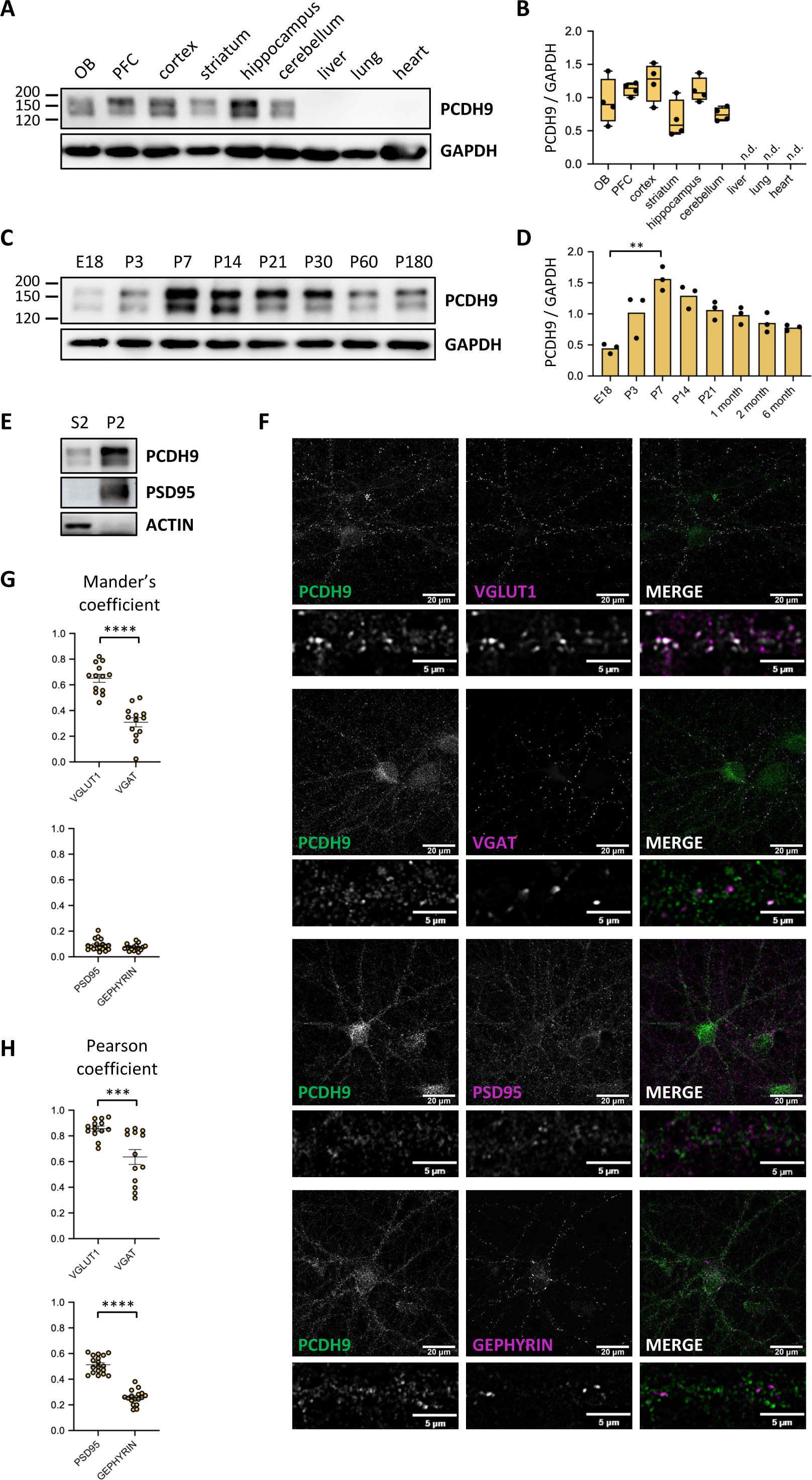
PCDH9 is enriched at glutamatergic synapses and its expression peaks in the first postnatal weeks in the hippocampus. **A** Representative PCDH9 western blot on brain regions and body organs from 2 month-old wild-type mice. OB, olfactory bulb. PFC, prefrontal cortex. **B** Quantifications of PCDH9 levels from (A). Protein levels were normalized to GAPDH. n=4. Kruskal-Wallis test, no significant differences. n.d., not detected. **C** Representative PCDH9 western blot on hippocampus from wild-type mice at different ages. E18, embryonic day 18; P3-180, postnatal day 3-180. **D** Quantifications of PCDH9 levels from (C). Protein levels were normalized to GAPDH. n=3. Kruskal-Wallis test, **p<0.01. **E** Representative PCDH9 western blot on cortex from 2 month-old wild-type mice. n=2. S2, supernatant 2 (cytosol-enriched fraction). P2, pellet 2 (membrane-enriched fraction). PSD95 and ACTIN are used as markers of P2 and S2, respectively. **F** Confocal images of DIV18 rat hippocampal neurons co-immunostained for PCDH9 and synaptic markers VGLUT1, VGAT, PSD95, GEPHYRIN. Insets show higher magnification of a dendrite. n=3 independent cultures. **G** Mander’s coefficient quantification of PCDH9 puncta colocalizing with presynaptic (top) and postsynaptic (bottom) markers analyzed in (F). VGLUT1, VGAT, n=13 neurons. PSD95, GEPHYRIN, n=17-18 neurons. **H** Pearson coefficient quantification of PCDH9 puncta colocalizing with presynaptic (upper) and postsynaptic (bottom) markers analyzed in (F). VGLUT1, VGAT, n=13 neurons. GEPHYRIN, n=17 neurons. PSD95, n=18 neurons.

### *Pcdh9* deletion leads to enlarged presynaptic and postsynaptic compartments in the CA1

To understand *Pcdh9* function in vivo, we took advantage of *Pcdh9* KO mouse line (*10*). As an initial characterization, we performed MRI analysis on *Pcdh9* KO and WT brains. We found significant changes in the volume of multiple brain areas (Suppl. Table 1), suggesting that *Pcdh9* might play a role in neural proliferation, growth or migration, similarly to other Protocadherins (*3*).

*Pcdh9* KO mice display impairments in social and object recognition (*10*). We could reproduce this evidence in our hands, thus confirming the robustness of the memory phenotype (Suppl. Fig. 1A, B). Therefore, we decided to focus our study on the hippocampus, a quintessential region for memory (*29*). Moreover, hippocampal alterations have been largely documented in MDD (*30*) and ASD (*31*, *32*), two pathologies associated with *PCDH9* in patients (*5*, *6*, *8*). Strikingly, ultrastructural analysis of excitatory synapses of the dorsal CA1 (dCA1) revealed a significant enlargement of the presynaptic terminal in *Pcdh9* KO mice compared to WT animals (Fig. 2A, B). As a consequence, the density of presynaptic vesicles was significantly reduced in *Pcdh9* KO neurons, while the pool of docked vesicles was unchanged (Suppl. Fig. 2). Importantly, the overgrowth of the presynaptic terminal was paralleled by a significant increase in the length and surface of the postsynaptic density (PSD), and a non-significant augmentation of the spine head area (Fig. 2A, B), indicating that both pre- and post-synaptic compartments are enlarged in *Pcdh9*-depleted neurons.

**Fig. 2.**
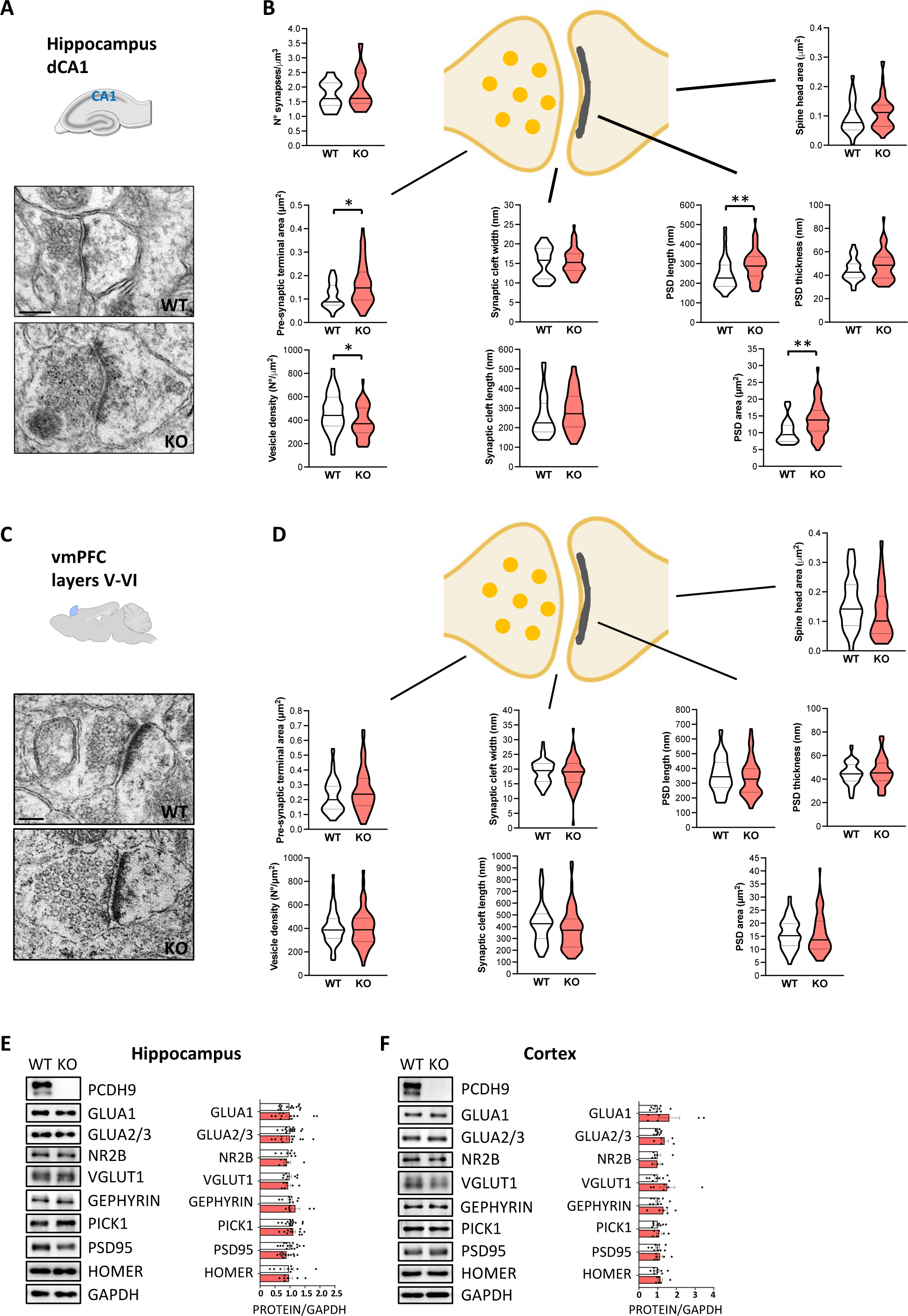
*Pcdh9* deletion leads to aberrant presynaptic and postsynaptic compartments in the CA1. **A** Representative electron micrographs of pyramidal neurons synapses from the dCA1 of 2 month-old WT and *Pcdh9* KO mice. Scale bar, 100 nm. **B** Quantifications of synaptic features from (A). n=30-50 synapses from 2 mice per genotype. Mann-Whitney test, *p<0.05, **p<0.01. **C** Representative electron micrographs of pyramidal neurons synapses from the vmPFC layer V-VI of 2 month-old WT and *Pcdh9* KO mice. Scale bar, 100 nm. **D** Quantifications of synaptic features from (C). n=50-70 synapses from 2 mice per genotype. Mann-Whitney test, no significant differences. **E** Representative western blot (left) and corresponding quantification (right) of synaptic markers levels on hippocampus from 2 month-old WT and *Pcdh9* KO mice. n=8-15. Mann-Whitney test, no significant differences. **F** Representative western blot (left) and corresponding quantification (right) of synaptic markers levels on cortices from 2 month-old WT and *Pcdh9* KO mice. n=6-9. Mann-Whitney test, no significant differences.

To understand whether *Pcdh9* role in synaptic structure was specific to CA1 or more general in the brain, we also examined the prefrontal cortex (PFC), another brain area strongly implicated in MDD and ASD (*33*, *34*). Examination of ventromedial PFC (vmPFC) synapses did not display any difference between the two genotypes, suggesting that the observed effect of *Pcdh9* KO on synaptic morphology is specific to CA1 (Fig. 2C, D). Altogether, we showed that *Pcdh9* depletion leads to an aberrant size increase both in the presynaptic terminal and in PSD, demonstrating that *Pcdh9* is important for establishing the correct morphology of CA1 excitatory synapses.

### *Pcdh9* deletions induces the dysregulation of key synaptic genes and the broad up-regulation of the synaptic transcriptome in the CA1

Then, we wanted to identify the molecular changes underlying the aberrant synaptic structure in the CA1 of *Pcdh9* KO mice. To this aim, we first performed a WB analysis of a panel of pre- and postsynaptic proteins. This approach did not reveal any obvious alteration in synapses composition between the two genotypes, neither in the hippocampus or in the cortex (Fig. 2E, F).

To gain a deeper insight into the molecular changes occurring in *Pcdh9*-depleted neurons, we conducted a single-nucleus RNAseq (snRNAseq) experiment on 16833 cells from WT and *Pcdh9* KO hippocampi. Nuclei were separated into 46 clusters, that were assigned to neuronal and non-neuronal cell types (Fig. 3A, left). Differentially expressed genes (DEG) analysis between *Pcdh9* KO and WT genotypes was implemented for each cluster. Surprisingly, *Pcdh9* was identified as an up-regulated DEG in *Pcdh9* KO neurons across multiple clusters. This was due to the increased transcription of *Pcdh9* locus downstream of the deleted exon 2, as verified by RT-qPCR (Suppl. Fig. 3A). No full-length or truncated PCDH9 proteins were detected in *Pcdh9* KO hippocampal lysates after WB with antibodies targeting PCDH9 C-terminal region, confirming the full knock-out of the protein (Suppl. Fig. 3B).

**Fig. 3.**
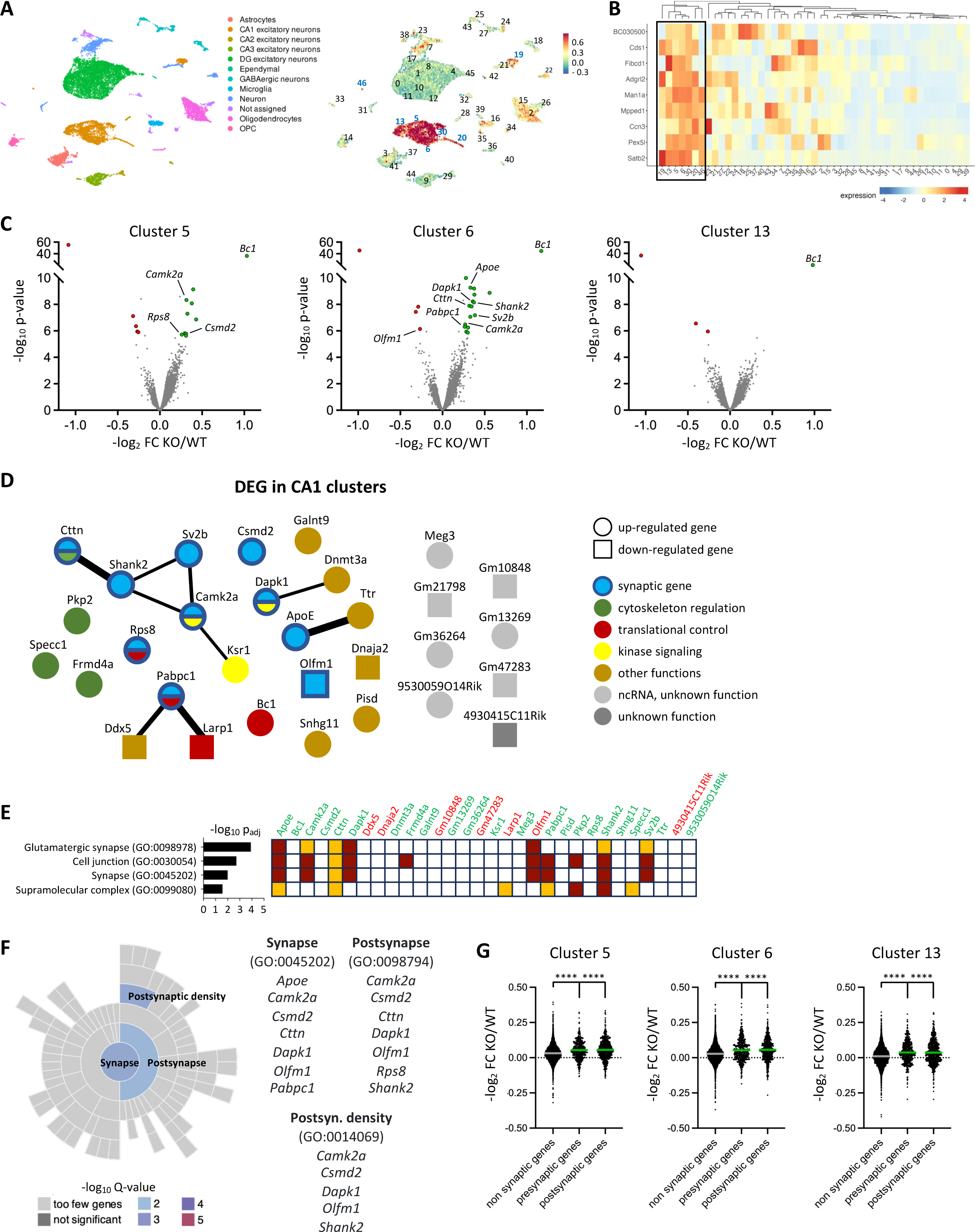
*Pcdh9* deletions induces the dysregulation of key synaptic genes and the broad up-regulation of the synaptic transcriptome in the CA1. **A** UMAP plots of snRNA-seq data. 16833 cells from WT and *Pcdh9* KO hippocampi were grouped into 46 different clusters. Cells are colored based on the attributed cell type (left) and the combined expression level of CA1 markers (right). n=2 mice per genotype. **B** Heatmap of CA1 markers fold changes among the different clusters. Clusters identified as CA1 populations are enclosed in a rectangle. **C** Volcano plots showing the KO/WT fold change and p-value distributions of genes in CA1 cluster 5 (left), 6 (center) and 13 (right). Up-regulated and down-regulated DEG are colored in green and red, respectively. **D** Network visualization of the combined 31 DEG found in CA1 clusters 5, 6 and 13. Each node represents a gene and is colored based on its function. A line connecting two nodes indicate a physical or functional interaction, as analyzed with STRING. The line thickness indicates the strength of data support. **E** The 4 GO categories (biological processes) identified with gProfiler on CA1 DEG. Green and red indicate up- and down-regulated DEG, respectively. For each GO category, the statistical significance and the associated genes are represented (brown square, inferred from experiments; orange square, inferred from sequences). **F** The 3 synaptic GO categories identified with SynGO on CA1 DEG. For each GO category, the statistical significance and the associated genes are indicated. **G** The effect of *Pcdh9* deletion on the expression levels of non-synaptic, presynaptic and postsynaptic genes in cluster 5 (left), 6 (center) and 13 (right) is shown. The horizontal line across the violin plot represents the median. Kruskal-Wallis test, ****p<0.0001.

Taking advantage of established CA1 marker genes (*19*), we identified 7 cell clusters containing CA1 excitatory neurons (Fig. 3A, B). Among the CA1 clusters, *Pcdh9* disruption led to the differential expression of 15, 21 and 4 genes in clusters 5, 6 and 13, respectively (Fig. 3C, D and Suppl. Table 2-4). Importantly, GO analysis of the combined 31 DEG from the three aforementioned CA1 clusters found enriched categories related to synapse and cell junction (Fig. 3E and Suppl. Table 5). In agreement, based on SynGO database for synapse ontology (*22*), multiple DEG were annotated as synaptic genes, mostly related to the postsynaptic compartment (Fig. 3F and Suppl. Table 6). Notably, multiple up-regulated DEG are known to promote spine formation. These include the PSD scaffold *Shank2*, a critical regulator of spine shape (*35*), the PSD components *Cctn* and *Csmd2* required for spine and dendrite morphogenesis (*36*, *37*), and the kinases *Camk2a* and *Dapk1* which act in structural synaptic plasticity (*38*, *39*). It is remarkable that the top up-regulated gene across the CA1 clusters is the non-coding RNA *Bc1*. *Bc1* has been previously implicated in the modulation of spine head and PSD size (*40*), therefore its dysregulation could also be implicated in the impaired synaptic organization of *Pcdh9*-depleted neurons.

Despite the number of DEG reaching statistical significance in CA1 clusters was relatively small, we noticed that a large majority of genes showed a moderate up-regulation in *Pcdh9* KO neurons compared to WT (Fig. 3C). We asked whether this global up-regulation was driven by synaptic genes, or rather general across all gene categories. Remarkably, pre- and postsynaptic genes showed a KO/WT fold change significantly higher than the rest of the transcriptome (Fig. 3G). In particular, genes coding for proteins localized to excitatory synapses showed a more pronounced up-regulation compared to genes related to inhibitory synapses (Suppl. Fig. 4A). Overall, our snRNAseq approach revealed that *Pcdh9* ablation leads to the dysregulation of key regulators of synapse size, and to a moderate but broad up-regulation of the synaptic transcriptome, thus sustaining the overgrowth of the pre- and postsynaptic compartments in CA1 excitatory synapses.

### *Pcdh9* deletion causes alterations in CA1 glutamatergic transmission and hippocampal network activity

Next, we asked how the synaptic and molecular alterations induced by *Pcdh9* depletion would impact on neuronal function. Electrophysiological recordings on pyramidal neurons from dCA1 acute slices revealed a significant increase in the frequency of spontaneous excitatory postsynaptic currents (EPSCs) in *Pcdh9* KO CA1, which doubled compared to WT (Fig. 4A-E). In agreement with electron microscopy observations, no differences in EPSCs properties between genotypes were observed in vmPFC slices (Fig. 4F-J). The change in mEPSCs frequency could originate from presynaptic alteration in glutamate release, therefore we performed paired-pulse ratio (PPR) experiments. No change in glutamate release probability was observed in *Pcdh9* KO mice (Fig. 4K), suggesting that a postsynaptic rather than a presynaptic defect might underlie the alteration in EPSCs frequency. Next, we examined if *Pcdh9* KO impaired long-term potentiation (LTP). Field excitatory post-synaptic potentials (fEPSPs) responses were recorded before and after high-frequency stimulation (HFS). No difference was found between the two genotypes, indicating that *Pcdh9* deletion does not affect this form of synaptic plasticity (Fig. 4L). In conclusions, we found that *Pcdh9* disruption leads to enhanced glutamatergic transmission, likely due to postsynaptic mechanisms.

**Fig. 4.**
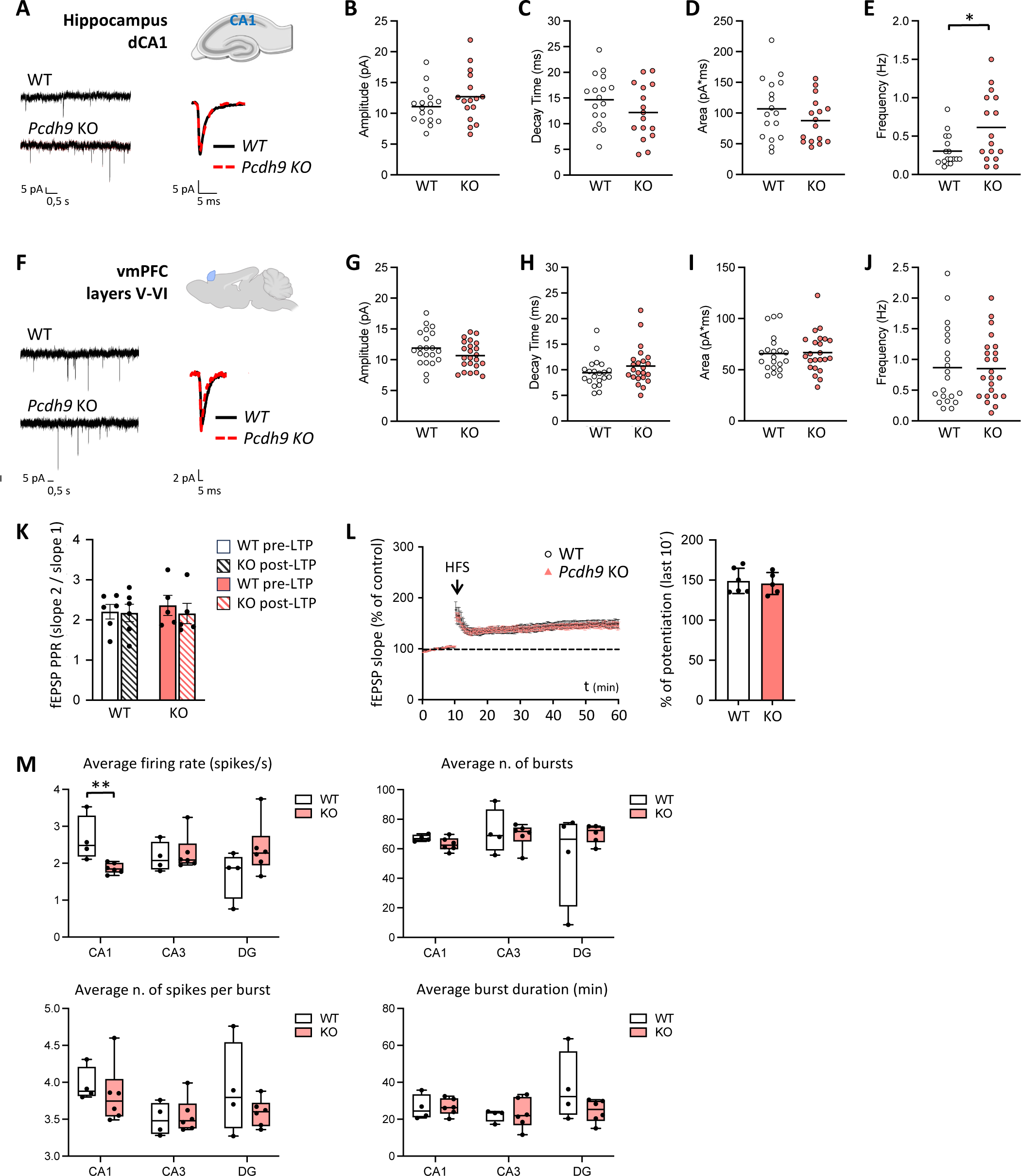
*Pcdh9* deletion leads to increased frequency of excitatory postsynaptic currents (EPSCs) and defective network activity in the CA1. **A** Representative traces of mEPSCs recorded from WT and *Pcdh9* KO CA1 pyramidal neurons. **B-E** Quantification of amplitude (B), decay time (C), area (D) and frequency (E) of mEPSCs from (A). n=16-17 recorded neurons from 5 independent animals per genotype. Mann-Whitney test, *p<0.05. **F** Representative traces of mEPSCs recorded from vmPFC pyramidal neurons from WT and *Pcdh9* KO mice. **G-J** Quantification of amplitude (G), decay time (H), area (I) and frequency (J) of mEPSCs from (F). n=21-22 recorded neurons from 5 independent mice per genotype. Mann-Whitney test, no significant differences. **K** Quantification of paired pulse ratio experiments in CA1 pyramidal neurons from WT and *Pcdh9* KO mice. n=5 independent mice per genotype. **L** Quantification of fEPSP slope before and after HFS in CA1 pyramidal neurons from WT and *Pcdh9* KO mice. n=5 independent mice per genotype. **M** Quantification of MEA recording features from WT and *Pcdh9* KO hippocampal slices. n=4-6 independent mice per genotype. Mann-Whitney test, **p<0.01.

Finally, we investigated the hippocampal network activity from *Pcdh9* KO and WT hippocampal slices using high resolution microelectrode array (MEA). Remarkably, we found a significant decrease of the average firing rate in the CA1, but not in other hippocampal regions, in *Pcdh9* KO animals (Fig. 4M). The average number of bursts, number of spikes per burst and burst duration were not affected in any hippocampal area. (Fig. 4M). These results suggest that the structural and functional alterations in CA1 synapses induced by *Pcdh9* deletion lead to disturbances in the hippocampal network activity.

## DISCUSSION

In this study, we explored the physiological role of *Pcdh9* in CA1 synapses. To this aim, we combined ultrastructural, biochemical, transcriptional, electrophysiological and hippocampal network approaches to characterize *Pcdh9* KO mice. One of the most striking findings of our work is that *Pcdh9* deletion leads to synapse overgrowth in the dCA1. Previous studies have implicated other protocadherins in spine generation or elimination (*3*), but their role in shaping synapse morphology has received less attention. Minor changes in synaptic structural features were reported in *Pcdh17* KO and *Pcdh19* mosaic mice (*41*, *42*). Here we show that *Pcdh9* ablation causes the aberrant enlargement of both presynaptic and postsynaptic compartments, demonstrating *Pcdh9* requirement for the control of synaptic size in the CA1.

We did not observe any change in dCA1 synapse number in *Pcdh9* KO mice, while previous reports found an increased spine density in the somatosensory cortex of these animals (*10*) and reduced spine counts in the ventral CA1 of a distinct *Pcdh9* KO line (*11*). Collectively, this suggests that *Pcdh9* role in spine generation is highly specific to the brain region considered, possibly depending on the particular *Pcdh9* expression profile and PCDH9 protein interactors in the different brain areas.

In rodents, synapses rapidly increase in number in the first postnatal weeks, and acquire the typical synaptic structure with normal sized synaptic vesicles and postsynaptic thickening at P7 (*28*, *43*). Our observation that PCDH9 hippocampal levels peak at P7, in line with analogous findings in the somatosensory cortex (*10*), supports the conclusion that *Pcdh9* plays a role in synaptogenesis. Therefore, we speculate that CA1 synaptic alterations observed in *Pcdh9* KO mice originate during postnatal neurodevelopment, and are then maintained in the adult brain.

Thanks to our single-nucleus transcriptional analysis, we gained insights into the molecular changes underlying synapse overgrowth in *Pcdh9* KO neurons. First, we noticed that most genes showed a mild up-regulation in *Pcdh9* KO neurons compared to WT. Remarkably, this global trend towards increased transcriptional levels was driven by synaptic genes: taken collectively, pre- and postsynaptic genes were up-regulated much more strongly than non-synaptic genes. This moderate but broad up-regulation of synaptic components could sustain the overgrowth of presynaptic terminal and PSD observed in *Pcdh9* KO neurons. Second, we identified several overexpressed DEG in *Pcdh9* KO neurons which promote synapse growth, such as *Shank2*, *Cttn*, *Camk2a*, *Dapk1* and *Csmd2*. The PSD scaffold *Shank2* recruit the actin-regulator *Cttn* to control actin dynamics and stimulate synapse morphogenesis (*36*, *44*, *45*). Notably, *Shank2* deficiency leads to a reduction in PSD thickness in the CA1 (*46*). *Camk2a* promotes activity-dependent spine enlargement by recruiting the actin cytoskeleton (*47*), and *Dapk1* also functions in structural synaptic plasticity by regulating *Camk2a* binding to the NMDA receptor (NMDAR) subunit GluN2B (*38*). The PSD component *Csmd2* promotes the development and maintenance of dendrites and synapses (*37*). Therefore, the up-regulation of *Shank2*/*Cttn* and *Camk2a*/*Dapk1* pathways and *Csmd2* gene could drive the increase in PSD size observed in *Pcdh9* KO neurons. Also, it is noteworthy that *Bc1* is the strongest up-regulated DEG in three CA1 clusters. *Bc1* is a non-coding RNA involved in the control of local protein synthesis at the synapse (*40*). In the barrel cortex, absence of *Bc1* causes the increased translation of PSD95 and other synaptic components, leading to the overgrowth of PSD, spine head and active zone. Thus, *Bc1* up-regulation might represent a compensatory mechanism at the translational level to counteract the increased transcription of synaptic genes, and synapse overgrowth, induced by *Pcdh9* deficiency.

Further studies are needed to clarify how *Pcdh9* deletion mechanistically leads to the transcriptional dysregulation of synaptic genes. Beyond their role as adhesion molecules, Protocadherins also function as signaling hubs by regulating multiple intracellular cascades, including pathways controlling gene expression such as *Wnt*/*ß-catenin* (*48*). Proteomic studies would be instrumental to determine PCDH9 binding partners and identify the signaling axis dysregulated in *Pcdh9* KO brains.

Our snRNAseq approach unveiled gene expression alterations in other neuronal and non-neuronal clusters besides CA1. DEG in these clusters only partly overlap with those identified in CA1 neurons, possibly suggesting cell-type specific effects of *Pcdh9* deletion. Further research is needed to assess *Pcdh9* role in other hippocampal regions and cell types. Through patch-clamp recording, we found that *Pcdh9* disruption positively modulates excitatory transmission. Considering that neither the pool of synaptic vesicles nor their release probability was increased in *Pcdh9* KO mice, we hypothesize that the augmented EPSCs frequency originates from alterations in the postsynaptic compartment. In agreement, the majority of the synaptic DEG that were identified localizes to the PSD. The SHANK family proteins function as scaffold at the PSD of excitatory synapses (*49*), and their dysregulation alters postsynaptic currents (*46*, *50*, *51*). Moreover, enhanced CaMK2a activity has been proved to alter glutamatergic transmission by reducing the proportion of silent synapses in the CA1 (*52*) or modifying AMPA receptors gating properties (*53*). Therefore, we believe that the upregulation of these genes in *Pcdh9* KO hippocampus might underlie the EPSCs alterations we found in *Pcdh9*-depleted CA1 neurons.

Growing evidence from human neuropathological studies and animal models indicate that autism may be conceptualized as a disease of the synapse (*54*), especially of the glutamatergic counterpart (*55*). Post-mortem research on ASD individuals revealed an increase in spine density of cortical pyramidal neurons (*56*). Furthermore, multiple genes regulating PSD organization have been associated with ASD and intellectual disabilities (*54*, *57*). To this regard, it is of great interest our finding that disruption of the autism associated gene *Pcdh9* leads to aberrantly enlarged CA1 synapses and dysregulation of genes controlling synaptic size. Moreover, 5 DEG identified in this work (*Camk2a*, *Dnmt3a*, *Galnt9*, *Pabpc1* and *Shank2*) are present in the SFARI database for ASD risk genes. Further research is required to examine whether ASD patients present dysmorphic synapses in the CA1.

In summary, our multi-level study sheds light on *Pcdh9* as a novel regulator of CA1 excitatory synapses growth and function, with relevant implications for hippocampal circuitries activity.

## MATERIALS AND METHODS

### Mice strains

*Pcdh9* KO mice were obtained from Dr. Martien Kas’ laboratory (University of Groningen, NL). Mice were maintained in rooms with 12 h light/dark cycles, temperature between 23-24 °C, and controlled humidity, with food and water provided *ad libitum*. Mice were housed at a maximum of five animals per cage in individually ventilated cages. All the experiments followed the guidelines established by the Italian Council on Animal Care and were approved by the Italian Government (protocol number 1152/2020-PR, 20/11/2020).

### Immunocytochemistry

Primary neurons were prepared from cortices and hippocampi of Sprague-Dawley E18 rat brains as previously described (*12*). Neurons were plated onto coverslips coated overnight with poly-L-lysine at 75000 neurons per well, and grown in Neurobasal plus medium (Invitrogen) supplemented with 2% B27 plus (Invitrogen), 1% L-glutamine (Invitrogen), 1% penicillin/streptomycin (Invitrogen) and 10 mM Glutamate.

Cultured neurons at DIV16-18 were washed in PBS and fixed in 4% paraformaldehyde, 10% sucrose for 15 min at room temperature (RT). After blocking/permeabilization in 10% normal goat serum (NGS), 0.1% Triton X-100, in PBS for 15 min at RT, neurons were incubated with primary antibodies (anti-PCDH9 Proteintech 25090-1-AP, 1:300; anti-PSD95 NeuroMab 75-028, 1:300; anti-GEPHYRIN Synaptic System 147021, 1:500; anti-VGLUT1 Synaptic Systems 135304, 1:200; anti-VGAT Synaptic Systems 131-003, 1:200) in GDB 1X solution (2X: 0.2% gelatin, 0.6% Triton X-100, 33 mM Na_2_HPO_4_, 0.9 M NaCl, pH 7.4) overnight at 4 °C. After three 10 min washes with high salt buffer (500 mM NaCl, 20 mM NaPO_4_(^2-^), pH 7.4), the coverslips were incubated with secondary antibodies (α-rabbit IgG Alexa-conjugated 488 Invitrogen A11029, 1:300; α-mouse IgG DyLight-conjugated 649 Jackson Immuno Research 211-492-177, 1:300; α-guinea pig IgG DyLight-conjugated 649 Jackson Immuno Research 706-175-148, 1:300) in GDB 1X solution for 1 h at RT. Neurons were washed three times with high salt buffer and incubated with DAPI (1:10000) for 5 min at RT. After washing with PBS for 5 min at RT, coverslips were mounted with Fluoromount (ThermoFisher Scientific).

### Image acquisition and analysis

Fluorescence images were acquired with an LSM800 confocal microscope (Carl Zeiss) and a 63X oil-immersion objective (numerical aperture 1.4) with sequential acquisition setting, at 1024 x 1024 pixels resolution. Images were Z-series projections of approximately 6-10 images, taken at depth intervals of 0.75 µm. Co-localization analysis was performed on randomly selected dendrites using Fiji software (*13*) with the plugin Jacop (*14*). For Manders’ colocalization coefficient, puncta were calculated by thresholding images at intensity equal to mean + 3 x St. Dev of the signals.

### Membrane-enriched fractions preparation

Cortices and hippocampi were dissected from *Pcdh9* WT and KO 2 month old animals, chopped with a razor blade in cold HEPES/sucrose buffer (4 mM HEPES pH 7.4, 320 mM sucrose) supplemented with protease inhibitor cocktails, incubated 10 min in ice and homogenized with a glass-teflon homogenizer. After centrifugation at 1000 g for 10 min at 4 °C, supernatants (S1) corresponding to total homogenate were centrifuged at 10000 g for 10 min at 4 °C. Resulting supernatants (S2) contained mainly cytosolic components while the pellet (P2) was enriched in membranes. After resuspension in HEPES/sucrose buffer, P2 fractions were centrifuged at 10000 g for 15 min at 4 °C to purify the preparation. P2 fraction were resuspended in modified RIPA buffer (50 mM TRIS-HCl, 200 mM NaCl, 1 mM EDTA, 1% NP40, 1% Triton X-100, pH 7.4) supplemented with protease inhibitor cocktail, and then quantified with bicinchoninic acid assay (BCA; Euroclone) prior to SDS-PAGE and Western blotting.

### Electron Microscopy

2.5-month-old *Pcdh9* KO and WT male mice (2 mice per genotype) were deeply anesthetized with isofluoran before transcardial perfusion with EM Buffer (2.5% glutaraldehyde and 2% paraformaldehyde in 0.1 M sodium cacodylate buffer, pH 7.4). Brains were post-fixed in EM buffer for 24 hours at 4 °C. 100 μm slices were generated with a vibratome (Leica VT1000S) and the areas of interest manually dissected. Samples were then post-fixed with 2% osmium tetroxide, rinsed, *en-bloc* stained with 1% uranyl acetate, dehydrated using increasing concentration of ethanol and finally with propylene oxide and embedded in epoxy resin (Electron Microscopy Science, Hatfield, PA, USA). Thin sections (70 nm) were obtained with a EM UC6 ultramicrotome (Leica Microsystems, Austria), stained with a saturated solution of uranyl acetate in ethanol 20% and observed under a Tecnai G2 Spirit transmission electron microscope (TEM) (FEI, Eindhoven, Netherland). Profiles of excitatory synapse used for quantitative analyses were identified on the basis of three conditions: 1) the presence in their postsynaptic terminal of the electron-dense post-synaptic density (PSD); 2) the presence of a cluster of at least 3 synaptic vesicles in the pre-synaptic compartment 3) the presence of a defined synaptic cleft. Images were acquired at a final magnification of 37000X for the morphometric analysis and of 13500X for the analysis of synapse density using a Megaview (Olympus, Munster, Germany). Morphometric analyses were performed with Fiji software (*13*) whereas, to evaluate synapse density, we used a stereological approach, as previously described (*15*).

### Western blots

Samples were loaded on 9% polyacrylamide gels and then transferred onto nitrocellulose membranes (0.22 μm, GE Healthcare). Membranes were blocked in 5% milk, 0.1% TBS Tween-20 for 1 h at RT and then incubated with the primary antibodies (anti-PCDH9 home-made rat antibody (*16*), 1:2000; anti-GLUA1 Cell Signaling 13185S, 1:1000; anti-GLUA2/3 home-made rabbit antibody, kind gift from Cecilia Gotti, 1:3000; anti-NR2B; anti-VGLUT1 Synaptic System 135303 1:15000; anti-GEPHYRIN ThermoFisher Scientific PA5-19589 1:2000; anti-PICK1 Neuromab 75040 1:2000; anti-PSD95 Cell Signaling 3450, 1:20000; anti-HOMER 1:10000; anti-GAPDH, Santa Cruz Biotechnology, 1:10000) in 0.1% TBS-Tween-20 overnight at 4 °C. After washing, the blots were incubated at room temperature for 1 h with horseradish peroxidase (HRP) - conjugated anti-rabbit (1:20000), anti-mouse (1:5000) or anti-rat (1:5000) antibodies in 0.1% TBS-Tween-20. Immunoreactive bands on blots were visualized by enhanced chemiluminescence (GE Healthcare). Quantification of bands intensity was performed with Fiji software (*13*).

### Hippocampal nuclei preparation and snRNA-seq sequencing

Hippocampal nuclei from two 2-month old mice per genotype were prepared using Chromium Nuclei Isolation Kit (10X Genomics, PN-1000494) according to manufacturer instructions, with minor modifications. Briefly, hippocampi were dissected, flash-frozen in liquid nitrogen and immediately processed for nuclear preparation. Frozen hippocampi were dissociated in Lysis Buffer with pestle, passed onto Nuclei Isolation Column and resuspended in Debris Removal Buffer. Pelleted nuclei were washed twice in Wash and Resuspension buffer and passed twice through a 40 µm cell strainer (Greiner Bio-One EASYstrainer cell sieve 542040). Nuclei integrity was examined at the confocal after DAPI staining, and the number of nuclei was estimated with a cell counting chamber. 13200 nuclei per sample were loaded on a Chromium instrument (10x Genomics). Libraries were prepared using a Single Cell 3’ Kit with v3 Chemistry following a standard protocol. Library profile and concentration were assessed using a TapeStation (Agilent). Library concentration was also analyzed using a Qubit fluorometer. All the libraries were sequenced using a Novaseq 6000 (Illumina).

### snRNAseq analysis

Raw reads were processed with Cell Ranger v7.1. The raw single cell matrices were subsequently processed with Seurat v. 4.0.44 (*17*). Only nuclei with a number of detected genes between 500 and 4000, with less than 20000 unique reads/nucleus and less than 10% reads mapped to mitochondrial DNA were retained for subsequent analyses. Integration of different experiments was achieved using the SCTransform algorithm, while regressing out the effect of Unique Molecular Identifier (UMI) and mitochondrial counts. Principal Component Analysis (PCA) was performed as an initial dimensionality reduction step. All the components required to explain a cumulative variance of > 90% were retained for downstream processing, corresponding to the first 30 PCA components. Uniform Manifold Approximation and Projection (UMAP) algorithm was run on top of PCA to end-up with a total of 2 dimensions. These dimensions were subsequently used as input for cluster identification, which was performed with the Louvain algorithm. To optimize cluster resolution identification, the Clustree algorithm was applied, and the optimal cluster resolution was selected as a compromise between maximization of the SC3 stability index and cluster number. SingleR (*18*) was subsequently applied to Log-Normalized gene counts at cluster level, using the CellDex dataset and the celldex::MouseRNAseqData as a reference to perform cell type identification, which was followed by direct manual curation. Differentially expressed genes were identified using a negative binomial generalized linear model. For each cluster, transcripts were considered to be differentially expressed in WT vs *Pcdh9* KO cells when the following three conditions were satisfied: absolute value of the log_2_FC KO/WT > 0.2, at least 10% of the cells expressing the transcript in at least one genotype, and a Bonferroni-adjusted p-value < 0.1.

CA1 clusters were identified based on CA1 marker genes from Cembrowski et al. (*19*). For interactions and GO analysis, the DEG between the two genotypes found in CA1 clusters 5, 6 and 13 were combined in a unique list of CA1 DEG, from which *Pcdh9* was excluded. The interactions of the 31 CA1 DEG were explored with STRING (*20*). GO analysis of the CA1 DEG was performed with gProfiler (*21*) and SynGO (*22*). STRING and SynGO analysis were performed on the human homologues of DEG. Genes were annotated as pre- and postsynaptic (Fig. 3g) based on SynGO ontologies. The genes coding for components of the excitatory and inhibitory PSD (Suppl. Fig. 4) were identified in Uezu et al. (*23*).

### Behavioral assays

All behavioral tests were performed at a similar time of the day (1 to 3 hours before the light off) to avoid circadian effects. The mice were allowed to habituate to the behavioral room for at least 45 min before each test. Behavioral equipment was cleaned with 70% ethanol after each test session to avoid olfactory cues.

Social recognition test was performed as in Bruining et al. (*10*) (long-term SRE). Before testing all mice were group-housed. To avoid exposure to odor prior to testing, test and intruder animals were all housed in separate cages. The day before the test, intruder animals were labeled with neutral smelling waterbased markers. Habituation and testing took place in regular cages (29 x 17 x 11 cm) with fresh bedding. On day 1, 2-month-old males (test animals) were habituated in the test cage for 5 min and initially exposed to an age-matched male conspecific (familiar intruder) for 2 min. Then, test animals were returned to their home cage. On day 2, after 24 h, the test animal was habituated again in clean cage with fresh bedding for 5 min, and then exposed at the same time with the familiar intruder of day 1 and to a novel age-matched male (novel intruder) from a different cage than the familiar intruder. Social investigation was defined as the total time that the test mice engaged in social sniffing, anogenital sniffing and allogrooming. The time spent by the test animal in investigating each intruder was manually scored from video recordings by an observer blind to the animal’s genotype.

For novel object recognition, habituation and testing took place in empty transparent plastic cages. The objects used were made of glass or plastic and were of sufficient height to discourage mice from climbing on the objects. In a pilot experiment, mice were exposed for 5 min to different couples of objects in order to select two that fulfilled the following conditions: mice had to show interest for both objects, with no overt intrinsic preference for one object compared to the other. The choice of which object to use as familiar and which one as novel was counterbalanced among the different animals tested. On day 1 and day 2, 2-month-old males were habituated in the test cage for 5 min. Then, test animals were returned to their home cage. On day 3, test animals were exposed to two identical objects (familiar objects) for 5 min, and then returned to their home cage. On day 4, after 24 h, test animals were exposed for 5 min to the familiar object and to a novel object. Object exploration was defined as approaching and having physical contact with the object: sniffing the object, turning the head towards it, touching it with the forepaws. The time spent by the animals in investigating each object was manually scored from video recordings by an observer blind to the animal’s genotype.

### *In vivo* Magnetic Resonance Imaging (MRI)

MRI experiments were conducted on a 7-Tesla scanner for rodents, fully equipped for brain MRI/MRS (Biospec, Paravision 6.0 Software Bruker-Biospin). A dedicated mouse head coil (4-channels) was used as receiver together with a volume coil as transmitter. A mixture of IsoVet (isoflurane 1-2%; Zootecnica, #104331020) with oxygen was used to anaesthetize animals and breath rate was constantly monitored to regulate the level of anesthesia. Body temperature was maintained through warm water circulating inside the bed. For cerebral anatomy analysis, T2 weighted MR images were acquired with coronal sections across the entire brain of 0.6 mm and an in-plane resolution of 71 µm (FOV = 16 × 14 mm, matrix = 224 × 200) using a fast-spin-echo sequence (TR/TE = 3500/44 ms, rare factor = 10, average = 10, 11 minutes of acquisition). T2-weighted images were analyzed using a MATLAB toolbox, the Atlas Normalization Toolbox using elastiX (ANTX, https://github.com/ChariteExpMri/antx2) (*24*), allowing to extract the cerebral volume of different brain areas following the Allen Mouse Brain Atlas (http://mouse.brain-map.org/). Areas with a cluster of at least 5 voxels were considered that correspond to a minimum volume of 0.025 mm^3^. Single brain region volume was then compared to the total brain volume of each mouse. The whole procedure was completed following the guidelines described in Koch et al. (*24*).

### RT-qPCR

mRNA was extracted from murine tissues using Nucleozol Reagent (Macherey Nagel, 15443435) following manufacturer’s instructions. A total of 1 μg of extracted mRNA was used to synthetize cDNA using SuperScript™ VILO™ cDNA Synthesis Kit (Thermo Fisher). *Pcdh9* specific sequences were amplified from cDNA with SYBR Green PCR Master Mix (Applied Biosystems) using an Applied Biosystems 7000 Real-Time thermocycler. Parallel qPCR reactions were performed on *α-actin* as reference gene. Primers were designed with Primer-BLAST online tool (https://www.ncbi.nlm.nih.gov/tools/primer-blast/): *Pcdh9* exon 1 (F: ACCATCACCCAACTCTGACG, R: GGAGTGTATCCCACCGCATC), *Pcdh9* exon 2 (F: AGACTGCCCTGGTAAGGGTT; R: AAGGAGGCATTCGGTCCAAG), *Pcdh9* exon 5 (F: CTCCTGGCTTGGGTCCATAC, R: GTGGCCGCCATTGTTGAAAT), *α-actin* (F: AGATGACCCAGATCATGTTTGAGA, R: CCTCGTAGATGGGCACAGTGT). Each reaction was performed in triplicate, and the results were analyzed using the ΔΔCT method.

### Patch-Clamp

P25-P30 *Pcdh9* KO and WT male mice were humanely sacrificed and the brain was rapidly removed and placed in an ice-cold cutting solution containing: 195 mM sucrose, 10 mM NaCl, 25 mM NaHCO_3_, 2.5 mM KCl, 1.25 mM NaH_2_PO_4_, 7 mM MgCl_2_, 0.5 mM CaCl_2_, 10 mM glucose (pH 7.3, equilibrated with 95% O_2_ and 5% CO_2_). Coronal slices from PFC and hippocampus (thickness, 250-350 μm) were prepared with a vibratome VT1000 S (Leica) and then incubated first for 40 min at 37 °C and then for 40 min at room temperature in artificial cerebrospinal fluid (aCSF), consisting of (in mM): 125 mM NaCl, 2.5 mM KCl, 1.25 mM NaH_2_PO_4_, 1 mM MgCl_2_, 2 mM CaCl_2_, 25 mM glucose, and 26 mM NaHCO_3_ (pH 7.3, equilibrated with 95% O_2_ and 5% CO_2_). Slices were transferred to a recording chamber perfused with aCSF at a rate of ∼2 ml/min at room temperature. Whole-cell patch-clamp electrophysiological recordings were performed with a Multiclamp 700B amplifier (Axon CNS molecular devices, USA) and using an infrared-differential interference contrast microscope. Patch microelectrodes (borosilicate capillaries with a filament and an outer diameter of 1.5 μm; Sutter Instruments) were prepared with a four-step horizontal puller (Sutter Instruments) and had a resistance of 3-5 MΩ. Miniature excitatory post synaptic currents (mEPSCs) were recorded from pyramidal neurons of the PFC (layers V/VI) and hippocampal CA1, as previously described (*25*).

Field excitatory postsynaptic potential (fEPSPs) were evoked (0.05 Hz of frequency) and recorded from the stratum radiatum of the hippocampal CA1 stimulating the Schaffer Collaterals (SC) using aCSF-filled monopolar glass electrodes. Stimulus strength was adjusted to give 50% maximal response. fEPSPs were acquired at 20 kHz and filtered at 5 kHz. For paired pulse experiments, pairs of stimuli were delivered at 50 ms intervals every 20 s (frequency of 0.05 Hz) and paired pulse ratio (PPR) were calculated by dividing the slope of the second response for the first one. SC-CA1 long-term potentiation (LTP) was elicited using classic stimulation protocol (100 stimuli at 100 Hz) (*26*). All the analyses were performed offline with Clampfit 10.1 software (Molecular Devices).

### Microelectrode Array (MEA)

Extracellular recordings were carried out with HD-MEA system (BiocamX, 3Brain) using Arena HD-MEA chips equipped with 4096 electrodes (21 × 21 μm^2^ in size, 42 μm pitch). Hippocampal brain slices were obtained as for Patch-Clamp experiments (400 µm thickness). Slice activity was recorded under continuous perfusion (4.5 ml/min) in aCSF solution supplemented with the potassium channel blocker 4-Aminopyridine (100 μM, Hello Bio). Recordings were performed at full-frame resolution (17855.5 Hz) and analyses of spikes rate, bursts rate, bursts duration and spikes inside the burst were conducted off-line using dedicated python routines (SpikeInterface) (*27*). When analyzing hippocampal slices, activated regions were manually identified overlapping slice images taken with a stereomicroscope (Zeiss Stemi 305) and the pseudocolor activity map visualized on the 3Brain software. For the analysis on the entire hippocampus, electrodes with a spiking rate >0.05 spike/sec were considered and slices with less than 20 active channels were discarded.

### Statistical analysis

Statistical comparisons were performed using GraphPad Prism software (San Diego, CA). The statistical significance of differences between two groups was calculated using a nonparametric two-tailed Mann-Whitney test. To compare three or more groups, Kruskal-Wallis followed by Dunn’s multiple comparisons test was used. Differences were considered significant at *p<0.05, **p<0.01, ***p<0.001, ****p<0.0001. n.s. indicates not significant. Pairwise comparisons are shown as brackets.

## FUNDING

Fondazione Cariplo, Italy (2019–3438)

## DECLARATION OF INTERESTS

The authors declare no competing interests.

## ACKOWLEDGMENTS

We thank the animal facility of the University Milano Bicocca for maintenance of mice colonies, Carlo Besta Neurological Institute for the use of the Transmission Electron Microscope, and the Experimental Imaging Centre of IRCCS-San Raffaele Hospital (Milano) for the MRI analysis. The financial support of Fondazione Cariplo, Italy (2019–3438) is gratefully acknowledged.

## AUTHOR CONTRIBUTION

F.M. and A.Z.A. performed the biochemical analysis. E.M and V.M. performed the stainings. F.M. and S.R. performed the behavioral tests. G.M. and M.F. performed and supervised the EM analysis, respectively. F.M., D.D. and S.S. performed the snRNA-seq. F.M. and R.P. analyzed snRNA-seq data. L.M. performed and analyzed the electrophysiological recordings. L.M. and A.Z. performed and analyzed MEA recordings, respectively. T.C. and L.C. performed the MRI study. S.H. and M.K. provided experimental tools. F.M. prepared all the figures and wrote the manuscript. All authors reviewed and edited the manuscript. M.P. conceptualized the study, acquired funding, and reviewed and edited the manuscript.

## FIGURES LEGENDS

**Suppl. Fig. 1. *Pcdh9* deletion causes defects in social and object recognition**

**A** Long-term social recognition assessed as time spent sniffing the familiar versus the novel intruder. n=10 mice per genotype. Mann-Whitney test, *p<0.05. n.s., non significant. **B** Long-term object recognition assessed as time spent sniffing the familiar versus the novel object. n=9-10 per genotype. Mann-Whitney test, *p<0.05. n.s., non significant.

**Suppl. Fig. 2. Evaluation of docked vesicles in *Pcdh9* KO dCA1 neurons**

Quantification of docked vesicles in dCA1 pyramidal neurons from 2 month-old WT and *Pcdh9* KO mice. The percentage of docked over total vesicles (left) and the number of docked vesicles per 100 nm of pre-synaptic length (right) are shown.

**Suppl. Fig. 3. Evaluation of *Pcdh9* mRNA and protein levels in *Pcdh9* KO hippocampus**

**A** RT-qPCR analysis of the mRNA levels of *Pcdh9* exon 1, 2 and 5 in *Pcdh9* KO and WT hippocampi. In *Pcdh9* KO mouse line, the whole second exon is replaced with a kanamycin/neomycin cassette (*10*). n=3 mice per genotype. *Pcdh9* mRNA levels were normalized over *Gapdh* mRNA levels. Unpaired t-test, **p<0.01, ***p<0.001. **B** Western blot analysis of PCDH9 levels in hippocampal lysates from 2 month-old *Pcdh9* KO and WT mice.

**Suppl. Fig. 4. *Pcdh9* deletion induces the up-regulation of excitatory PSD genes in the CA1**

The effect of *Pcdh9* deletion on the expression levels of inhibitory and excitatory PSD genes in cluster 5 (left), 6 (center) and 13 (right) is shown. Mann-Whitney test, *p<0.05, **p<0.01.

**Suppl. Table 1. MRI volumetric analysis of brain regions of *Pcdh9* KO mouse**

**Suppl. Table 2. DEG genes between *Pcdh9* KO and WT hippocampus**

**Suppl. Table 3. DEG genes between *Pcdh9* KO and WT CA1 clusters**

**Suppl. Table 4. STRING analysis of CA1 DEG genes**

**Suppl. Table 5. gProfiler analysis of CA1 DEG genes**

**Suppl. Table 6. synGO analysis of CA1 DEG genes**

## Notes

### Competing Interest Statement

The authors have declared no competing interest.

