## Supplementary figures and images for "Disruption of autism-associated *Pcdh9* gene leads to transcriptional alterations, synapses overgrowth and aberrant excitatory transmission in the CA1"

### Supplementary Figures 1-4

#### Supplementary Figure 1

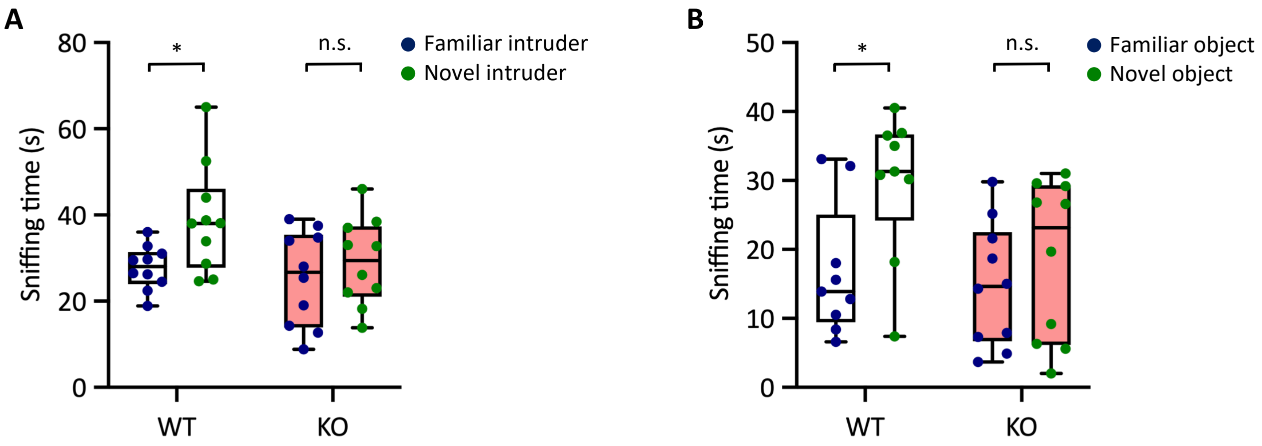


#### Supplementary Figure 2

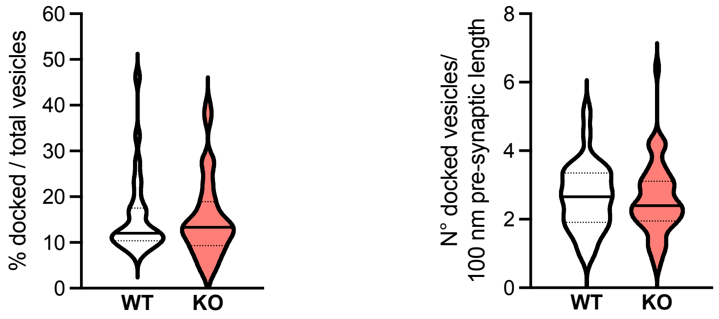


#### Supplementary Figure 3

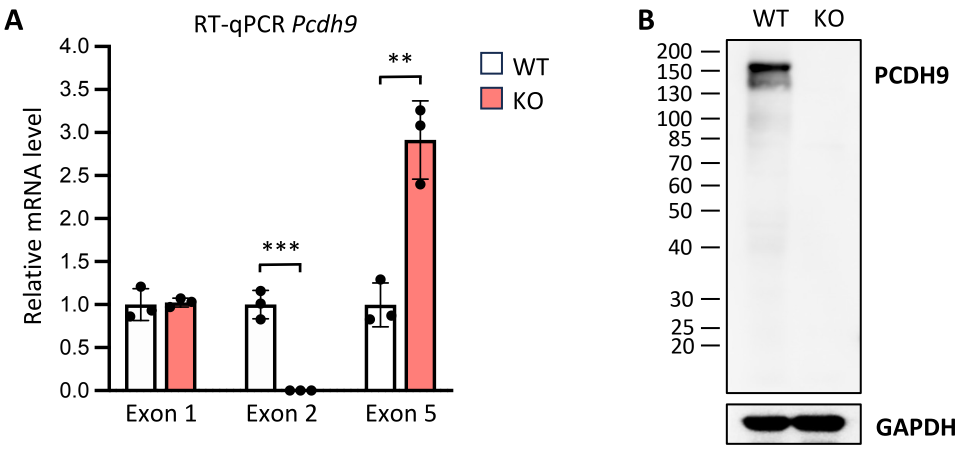


#### Supplementary Figure 4

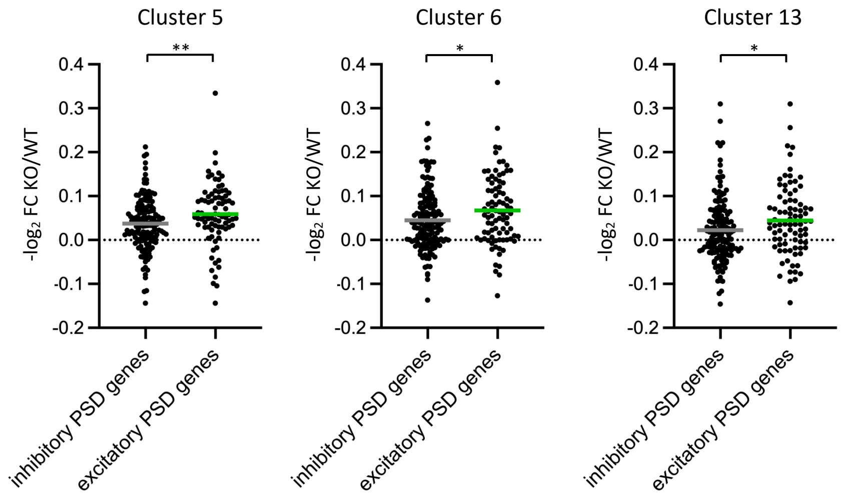
