## Supplementary Table 1 for "Disruption of autism-associated *Pcdh9* gene leads to transcriptional alterations, synapses overgrowth and aberrant excitatory transmission in the CA1"

| | Brain area | | | WT | | | KO | | | WT | | | | | | | KO | | | | | | |
|---|---|---|---|---|---|---|---|---|---|---|---|---|---|---|---|---|---|---|---|---|---|---|---|
| Significance | Region | % (KO-WT)/WT | *t-Test* | mean WT | dev.st | n° | mean KO | dev.st | n° | WT_1 | WT_2 | WT_3 | WT_4 | WT_5 | WT_6 | WT_7 | KO_1 | KO_2 | KO_3 | KO_4 | KO_5 | KO_6 | KO_7 |
| | root | -3,04 | *0,1126* | 483,65 | 17,29 | 7 | 468,94 | 14,68 | 7 | 477,0042 | 474,8807 | 464,3036 | 484,1219 | 477,7164 | 488,6861 | 518,8376 | 471,847 | 471,6581 | 455,2234 | 461,6168 | 457,9711 | 465,3195 | 498,9677 |
| | Basic cell groups and regions | -2,95 | *0,1128* | 427,75 | 14,71 | 7 | 415,15 | 12,73 | 7 | 422,2663 | 419,878 | 411,1632 | 428,4924 | 423,4197 | 431,3221 | 457,7309 | 417,5502 | 418,2195 | 403,1936 | 408,4459 | 405,7891 | 411,8353 | 441,0469 |
| | Cerebrum | -2,58 | *0,1516* | 260,71 | 8,88 | 7 | 253,98 | 7,48 | 7 | 258,6397 | 255,3111 | 250,2994 | 261,6128 | 256,5159 | 264,6557 | 277,9437 | 253,0807 | 256,3729 | 245,5828 | 249,9586 | 250,3418 | 253,3847 | 269,1194 |
| | Cerebral cortex | -2,65 | *0,1263* | 206,56 | 6,74 | 7 | 201,09 | 5,63 | 7 | 204,9577 | 201,1938 | 199,8032 | 208,0616 | 202,8003 | 209,4643 | 219,6334 | 200,7922 | 203,6219 | 195,2418 | 197,0798 | 198,1964 | 200,4481 | 212,2545 |
| | Cortical plate | -2,68 | *0,1175* | 198,22 | 6,33 | 7 | 192,90 | 5,41 | 7 | 196,4803 | 193,0968 | 191,9709 | 199,9676 | 194,8159 | 200,6887 | 210,5353 | 192,7986 | 195,3819 | 187,3608 | 188,7546 | 190,0537 | 192,3997 | 203,555 |
| | Isocortex | -2,85 | *0,0703* | 113,94 | 3,26 | 7 | 110,70 | 2,82 | 7 | 113,1973 | 111,7642 | 111,6516 | 115,1418 | 111,0278 | 114,2867 | 120,5185 | 110,7022 | 111,8859 | 108,6999 | 107,1573 | 109,0742 | 111,4933 | 115,8568 |
| | Frontal pole cerebral cortex | -17,10 | *0,1002* | 0,9763 | 0,1988 | 7 | 0,8094 | 0,1449 | 7 | 0,8733 | 1,089343 | 1,183672 | 1,250615 | 0,888514 | 0,718115 | 0,8307 | 0,800272 | 0,985886 | 0,9585 | 0,584229 | 0,666385 | 0,836785 | 0,833743 |
| * | Frontal pole layer 1 | -74,11 | *0,0282* | 0,4818 | 0,2829 | 6 | 0,1248 | 0,1444 | 5 | 0,416871 | 0,678558 | 0,535543 | 0,839829 | 0,407743 | | 0,012171 | | 0,060857 | 0,082157 | 0,103457 | | 0,003043 | 0,374271 |
| | Frontal pole layer 2/3 | 111,45 | *0,1536* | 0,2046 | 0,1642 | 4 | 0,4327 | 0,2588 | 5 | | | 0,343843 | 0,024343 | | 0,1065 | 0,343843 | 0,286029 | 0,672472 | 0,675514 | | 0,073029 | 0,456428 | |
| | Frontal pole layer 5 | -11,69 | *0,4228* | 0,3569 | 0,0648 | 7 | 0,3152 | 0,1148 | 7 | 0,401657 | 0,346886 | 0,2556 | 0,331672 | 0,392529 | 0,453386 | 0,316457 | 0,371229 | 0,179529 | 0,152143 | 0,395572 | 0,453385 | 0,2769 | 0,377314 |
| | Frontal pole layer 6a | 7,28 | *0,7782* | 0,0895 | 0,0486 | 7 | 0,0961 | 0,0347 | 7 | 0,054771 | 0,0639 | 0,048686 | 0,054771 | 0,088243 | 0,158229 | 0,158229 | 0,143014 | 0,073029 | 0,048686 | 0,0852 | 0,139971 | 0,100414 | 0,082157 |
| | Somatomotor areas | -1,71 | *0,1453* | 22,51 | 0,59 | 7 | 22,13 | 0,22 | 7 | 21,74729 | 22,03638 | 22,39848 | 22,68147 | 22,33457 | 22,81535 | 23,56997 | 22,08811 | 21,89032 | 22,22198 | 21,82338 | 22,11243 | 22,37716 | 22,36804 |
| | Primary motor area | -1,64 | *0,1933* | 10,56 | 0,30 | 7 | 10,39 | 0,13 | 7 | 10,25138 | 10,46439 | 10,27269 | 10,70173 | 10,44309 | 10,73216 | 11,08209 | 10,25748 | 10,33659 | 10,27573 | 10,33963 | 10,38527 | 10,50394 | 10,63479 |
| | Primary motor area Layer 1 | 1,12 | *0,7253* | 1,21 | 0,08 | 7 | 1,22 | 0,06 | 7 | 1,186714 | 1,131943 | 1,208015 | 1,217144 | 1,144114 | 1,144114 | 1,372329 | 1,211058 | 1,223229 | 1,274957 | 1,1076 | 1,189757 | 1,268871 | 1,2567 |
| | Primary motor area Layer 2/3 | -0,36 | *0,7837* | 3,53 | 0,11 | 7 | 3,52 | 0,04 | 7 | 3,46277 | 3,526673 | 3,392787 | 3,514502 | 3,456686 | 3,669687 | 3,687943 | 3,544931 | 3,508415 | 3,511456 | 3,465815 | 3,56927 | 3,547969 | 3,474943 |
| | Primary motor area Layer 5 | -1,43 | *0,3041* | 3,14 | 0,08 | 7 | 3,09 | 0,08 | 7 | 2,991128 | 3,164573 | 3,07633 | 3,149359 | 3,182829 | 3,18283 | 3,210214 | 2,994173 | 3,128058 | 3,03677 | 3,128058 | 3,125013 | 3,01547 | 3,2163 |
| * | Primary motor area Layer 6a | -5,05 | *0,0220* | 2,54 | 0,08 | 7 | 2,41 | 0,10 | 7 | 2,455585 | 2,507316 | 2,455587 | 2,680759 | 2,519486 | 2,552958 | 2,6199 | 2,355173 | 2,352129 | 2,294314 | 2,498187 | 2,349085 | 2,50427 | 2,540786 |
| | Primary motor area Layer 6b | -0,87 | *0,8882* | 0,1500 | 0,0197 | 7 | 0,1487 | 0,0137 | 7 | 0,155186 | 0,133886 | 0,139971 | 0,139972 | 0,139971 | 0,1491 | 0,1917 | 0,152143 | 0,124757 | 0,158229 | 0,139971 | 0,152143 | 0,167357 | 0,146057 |
| | Secondary motor area | -1,78 | *0,1725* | 11,95 | 0,34 | 7 | 11,74 | 0,17 | 7 | 11,49591 | 11,57199 | 12,12579 | 11,97974 | 11,89149 | 12,08319 | 12,48789 | 11,83064 | 11,55373 | 11,94625 | 11,48375 | 11,72717 | 11,87322 | 11,73326 |
| | Secondary motor area layer 1 | -2,58 | *0,4521* | 2,05 | 0,12 | 7 | 2,00 | 0,13 | 7 | 1,868314 | 1,965687 | 2,206072 | 2,09653 | 1,965686 | 2,096529 | 2,163472 | 2,090444 | 1,996115 | 2,142171 | 1,792243 | 1,920042 | 2,14217 | 1,907871 |
| | Secondary motor area layer 2/3 | 0,62 | *0,7465* | 3,42 | 0,14 | 7 | 3,44 | 0,09 | 7 | 3,331928 | 3,271073 | 3,380615 | 3,444516 | 3,338014 | 3,444516 | 3,712286 | 3,474945 | 3,420172 | 3,529713 | 3,255858 | 3,456684 | 3,502327 | 3,432343 |
| | Secondary motor area layer 5 | -2,07 | *0,0851* | 4,27 | 0,11 | 7 | 4,19 | 0,04 | 7 | 4,14437 | 4,162631 | 4,451701 | 4,208274 | 4,354329 | 4,275216 | 4,3239 | 4,247831 | 4,141329 | 4,226527 | 4,199144 | 4,141327 | 4,15654 | 4,186971 |
| | Secondary motor area layer 6a | -3,78 | *0,0782* | 2,14 | 0,07 | 7 | 2,06 | 0,09 | 7 | 2,093485 | 2,111744 | 2,023501 | 2,16043 | 2,157386 | 2,206072 | 2,227372 | 1,99003 | 1,953515 | 2,005242 | 2,148258 | 2,154342 | 2,020456 | 2,142171 |
| | Secondary motor area layer 6b | -17,57 | *0,1835* | 0,0643 | 0,0064 | 7 | 0,0530 | 0,0193 | 7 | 0,057814 | 0,060857 | 0,0639 | 0,069986 | 0,076071 | 0,060857 | 0,060857 | 0,027386 | 0,0426 | 0,0426 | 0,088243 | 0,054771 | 0,051729 | 0,0639 |
| * | Somatosensory areas | -4,68 | *0,0114* | 31,14 | 0,91 | 7 | 29,68 | 0,92 | 7 | 31,77655 | 31,28059 | 30,11821 | 31,64269 | 30,7359 | 29,97216 | 32,44903 | 29,89 | 30,11212 | 29,04406 | 28,82195 | 28,86149 | 29,61915 | 31,42054 |
| * | Primary somatosensory area | -4,76 | *0,0084* | 22,93 | 0,66 | 7 | 21,84 | 0,63 | 7 | 23,48476 | 23,14398 | 22,03334 | 23,33264 | 22,67841 | 22,13984 | 23,72516 | 22,02725 | 22,00899 | 21,30304 | 21,46128 | 21,30303 | 21,68339 | 23,10746 |
| | Primary somatosensory area nose | -15,38 | *0,0661* | 2,87 | 0,44 | 7 | 2,43 | 0,37 | 7 | 3,301499 | 3,137187 | 2,887672 | 3,20413 | 3,048943 | 2,111744 | 2,409943 | 2,297358 | 2,388643 | 1,971771 | 2,695972 | 2,276056 | 3,103714 |
| | Primary somatosensory area nose layer 1 | -9,59 | *0,3246* | 0,3943 | 0,0594 | 7 | 0,3564 | 0,0769 | 7 | 0,495986 | 0,432086 | 0,310372 | 0,374272 | 0,380357 | 0,356014 | 0,410786 | 0,4686 | 0,2556 | 0,292114 | 0,441214 | 0,3195 | 0,356014 | 0,3621 |
| | Primary somatosensory area nose layer 2/3 | -16,82 | *0,1267* | 0,6177 | 0,1341 | 7 | 0,5138 | 0,0985 | 7 | 0,797228 | 0,785058 | 0,541629 | 0,614658 | 0,6177 | 0,429043 | 0,538586 | 0,526415 | 0,438171 | 0,395571 | 0,666386 | 0,535543 | 0,432085 | 0,602486 |
| | Primary somatosensory area nose layer 4 | -25,68 | *0,0614* | 0,5890 | 0,1657 | 7 | 0,4377 | 0,0885 | 7 | 0,782014 | 0,715072 | 0,514243 | 0,699858 | 0,654214 | 0,356014 | 0,401657 | 0,392529 | 0,407743 | 0,349928 | 0,5751 | 0,426 | 0,365143 | 0,547714 |
| | Primary somatosensory area nose layer 5 | -22,74 | *0,0523* | 0,5773 | 0,1296 | 7 | 0,4460 | 0,0929 | 7 | 0,663343 | 0,642043 | 0,593357 | 0,684643 | 0,672471 | 0,386443 | 0,398614 | 0,3834 | 0,480771 | 0,371228 | 0,483814 | 0,377314 | 0,398614 | 0,626829 |
| | Primary somatosensory area nose layer 6a | -2,20 | *0,8587* | 0,6516 | 0,1335 | 7 | 0,6373 | 0,1601 | 7 | 0,526414 | 0,529457 | 0,858086 | 0,794186 | 0,684643 | 0,544672 | 0,623786 | 0,499029 | 0,757672 | 0,523371 | 0,4899 | 0,587271 | 0,6816 | 0,921986 |
| | Primary somatosensory area nose layer 6b | -7,29 | *0,5948* | 0,0417 | 0,0126 | 7 | 0,0387 | 0,0074 | 7 | 0,036514 | 0,033471 | 0,069986 | 0,036514 | 0,039557 | 0,039557 | 0,036514 | 0,027386 | 0,048686 | 0,039557 | 0,039557 | 0,030429 | 0,0426 | 0,0426 |
| | Primary somatosensory area barrel field | -2,16 | *0,4071* | 5,85 | 0,29 | 7 | 5,72 | 0,26 | 7 | 5,814898 | 5,677975 | 5,766216 | 5,787518 | 5,449757 | 6,107017 | 6,329143 | 5,927489 | 5,976172 | 5,796641 | 5,209373 | 5,568426 | 5,729697 | 5,839243 |
| | Primary somatosensory area barrel field layer 1 | -4,33 | *0,0778* | 0,7525 | 0,0255 | 7 | 0,7199 | 0,0362 | 7 | 0,748543 | 0,708986 | 0,730286 | 0,782015 | 0,769843 | 0,769843 | 0,757671 | 0,7242 | 0,782014 | 0,718114 | 0,657257 | 0,715071 | 0,7242 | 0,718114 |
| | Primary somatosensory area barrel field layer 2/3 | -1,42 | *0,6379* | 1,41 | 0,05 | 7 | 1,39 | 0,10 | 7 | 1,439271 | 1,378415 | 1,375372 | 1,390587 | 1,396671 | 1,396672 | 1,5123 | 1,433186 | 1,484914 | 1,360157 | 1,223229 | 1,357114 | 1,375371 | 1,515343 |
| | Primary somatosensory area barrel field layer 4 | 0,07 | *0,9813* | 1,22 | 0,05 | 7 | 1,22 | 0,08 | 7 | 1,2141 | 1,189758 | 1,232358 | 1,180629 | 1,153243 | 1,256701 | 1,287129 | 1,262786 | 1,305386 | 1,204971 | 1,074129 | 1,168457 | 1,195842 | 1,308429 |
| | Primary somatosensory area barrel field layer 5 | -0,91 | *0,7773* | 1,14 | 0,08 | 7 | 1,13 | 0,05 | 7 | 1,092385 | 1,107601 | 1,168458 | 1,128901 | 1,016314 | 1,223229 | 1,247572 | 1,159329 | 1,177586 | 1,131942 | 1,025443 | 1,150199 | 1,162371 | 1,104557 |
| | Primary somatosensory area barrel field layer 6a | -5,15 | *0,3007* | 1,19 | 0,13 | 7 | 1,13 | 0,07 | 7 | 1,183671 | 1,141072 | 1,131943 | 1,183672 | 0,9798 | 1,329729 | 1,375372 | 1,195844 | 1,0863 | 1,238442 | 1,089343 | 1,068042 | 1,144114 | 1,074129 |
| | Primary somatosensory area barrel field layer 6b | -2,24 | *0,6739* | 0,1361 | 0,0111 | 7 | 0,1330 | 0,0150 | 7 | 0,136929 | 0,152143 | 0,1278 | 0,121714 | 0,133886 | 0,130843 | 0,1491 | 0,152143 | 0,139971 | 0,143014 | 0,139971 | 0,109543 | 0,1278 | 0,118671 |
| | Primary somatosensory area lower limb | -6,21 | *0,1352* | 2,26 | 0,14 | 7 | 2,12 | 0,18 | 7 | 2,330828 | 2,385601 | 2,072186 | 2,425159 | 2,297357 | 2,087401 | 2,190857 | 2,056973 | 2,376472 | 1,840928 | 2,148258 | 2,002199 | 2,07827 | 2,306486 |
| | Primary somatosensory area lower limb layer 1 | -2,76 | *0,5869* | 0,2682 | 0,0257 | 7 | 0,2608 | 0,0237 | 7 | 0,273857 | 0,279943 | 0,286029 | 0,261686 | 0,286029 | 0,213 | 0,2769 | 0,264729 | 0,295157 | 0,231257 | 0,249514 | 0,286028 | 0,261686 | 0,237343 |
| | Primary somatosensory area lower limb layer 2/3 | -5,34 | *0,3575* | 0,6181 | 0,0524 | 7 | 0,5851 | 0,0743 | 7 | 0,645086 | 0,672472 | 0,5538 | 0,669429 | 0,648129 | 0,556843 | 0,581186 | 0,5538 | 0,632914 | 0,502071 | 0,642043 | 0,502071 | 0,565971 | 0,696814 |
| | Primary somatosensory area lower limb layer 4 | -5,90 | *0,2963* | 0,2726 | 0,0300 | 7 | 0,2565 | 0,0248 | 7 | 0,282986 | 0,286029 | 0,225172 | 0,3195 | 0,279943 | 0,264729 | 0,249514 | 0,249514 | 0,289071 | 0,222128 | 0,258643 | 0,243428 | 0,243428 | 0,289071 |
| | Primary somatosensory area lower limb layer 5 | -7,96 | *0,0970* | 0,5681 | 0,0466 | 7 | 0,5229 | 0,0473 | 7 | 0,623786 | 0,587272 | 0,480772 | 0,605529 | 0,5751 | 0,5538 | 0,550757 | 0,514243 | 0,584229 | 0,444257 | 0,499029 | 0,514243 | 0,529457 | 0,5751 |
| | Primary somatosensory area lower limb layer 6a | -7,16 | *0,1010* | 0,4860 | 0,0262 | 7 | 0,4512 | 0,0436 | 7 | 0,465557 | 0,5112 | 0,499029 | 0,523372 | 0,465557 | 0,453386 | 0,483814 | 0,441215 | 0,529457 | 0,392528 | 0,462514 | 0,422957 | 0,435128 | 0,474686 |
| | Primary somatosensory area lower limb layer 6b | -8,16 | *0,3690* | 0,0426 | 0,0075 | 7 | 0,0391 | 0,0064 | 7 | 0,039557 | 0,048686 | 0,027386 | 0,045643 | 0,0426 | 0,045643 | 0,048686 | 0,033471 | 0,045643 | 0,048686 | 0,036514 | 0,033471 | 0,0426 | 0,033471 |
| | Primary somatosensory area mouth | -1,08 | *0,7289* | 5,75 | 0,40 | 7 | 5,69 | 0,23 | 7 | 5,732741 | 5,44976 | 5,541045 | 5,525832 | 5,419329 | 6,116145 | 6,484329 | 5,872717 | 5,723615 | 5,970084 | 5,404116 | 5,681011 | 5,805768 | 5,376729 |
| | Primary somatosensory area mouth layer 1 | -3,72 | *0,4370* | 0,7368 | 0,0503 | 7 | 0,7094 | 0,0742 | 7 | 0,7455 | 0,663343 | 0,812443 | 0,778972 | 0,739414 | 0,690729 | 0,727243 | 0,605529 | 0,815486 | 0,718114 | 0,635957 | 0,693771 | 0,715071 | 0,782014 |
| | Primary somatosensory area mouth layer 2/3 | -0,95 | *0,8526* | 1,42 | 0,13 | 7 | 1,41 | 0,14 | 7 | 1,308428 | 1,214101 | 1,478829 | 1,439272 | 1,381457 | 1,557944 | 1,551857 | 1,475787 | 1,5123 | 1,509257 | 1,147158 | 1,299299 | 1,503171 | 1,390586 |
| | Primary somatosensory area mouth layer 4 | 8,98 | *0,2497* | 0,8668 | 0,1408 | 7 | 0,9446 | 0,0932 | 7 | 0,785057 | 0,733329 | 0,870257 | 0,806358 | 0,748543 | 1,0437 | 1,080214 | 1,046743 | 0,961543 | 1,034571 | 0,797229 | 0,931114 | 0,991971 | 0,848957 |
| | Primary somatosensory area mouth layer 5 | 6,29 | *0,3944* | 1,12 | 0,18 | 7 | 1,19 | 0,10 | 7 | 1,119771 | 1,055872 | 0,970672 | 0,955458 | 0,982843 | 1,329729 | 1,417972 | 1,235401 | 1,134986 | 1,274957 | 1,183672 | 1,265828 | 1,229314 | 1,0011 |
| * | Primary somatosensory area mouth layer 6a | -10,46 | *0,0381* | 1,51 | 0,13 | 7 | 1,35 | 0,12 | 7 | 1,658357 | 1,676615 | 1,335815 | 1,439272 | 1,463614 | 1,405801 | 1,609672 | 1,427101 | 1,208014 | 1,347985 | 1,554901 | 1,408842 | 1,268871 | 1,265829 |
| | Primary somatosensory area mouth layer 6b | -11,45 | *0,0853* | 0,0987 | 0,0141 | 7 | 0,0874 | 0,0055 | 7 | 0,115629 | 0,1065 | 0,073029 | 0,1065 | 0,103457 | 0,088243 | 0,097371 | 0,082157 | 0,091286 | 0,0852 | 0,0852 | 0,082157 | 0,097371 | 0,088243 |
| * | Primary somatosensory area upper limb | -4,29 | *0,0020* | 3,57 | 0,06 | 7 | 3,42 | 0,08 | 7 | 3,511456 | 3,614916 | 3,499287 | 3,669688 | 3,581443 | 3,535802 | 3,611872 | 3,33193 | 3,487115 | 3,450599 | 3,33193 | 3,426255 | 3,377569 | 3,544929 |
| * | Primary somatosensory area upper limb layer 1 | -9,34 | *0,0026* | 0,4282 | 0,0097 | 7 | 0,3882 | 0,0228 | 7 | 0,422957 | 0,422957 | 0,429043 | 0,4473 | 0,416871 | 0,432086 | 0,426 | 0,365143 | 0,401657 | 0,407743 | 0,352972 | 0,413828 | 0,380357 | 0,395571 |
| | Primary somatosensory area upper limb layer 2/3 | -1,56 | *0,4184* | 0,9176 | 0,0253 | 7 | 0,9033 | 0,0373 | 7 | 0,9159 | 0,928072 | 0,885472 | 0,931115 | 0,897643 | 0,961543 | 0,903729 | 0,879386 | 0,879386 | 0,928071 | 0,903729 | 0,903728 | 0,858085 | 0,970671 |
| * | Primary somatosensory area upper limb layer 4 | -3,69 | *0,0429* | 0,5303 | 0,0131 | 7 | 0,5108 | 0,0184 | 7 | 0,538586 | 0,541629 | 0,517286 | 0,523372 | 0,550757 | 0,520329 | 0,520329 | 0,502072 | 0,517286 | 0,514243 | 0,502072 | 0,492943 | 0,499028 | 0,547714 |
| * | Primary somatosensory area upper limb layer 5 | -6,17 | *0,0445* | 0,7890 | 0,0265 | 7 | 0,7403 | 0,0487 | 7 | 0,754628 | 0,797229 | 0,775929 | 0,833743 | 0,8094 | 0,772886 | 0,778971 | 0,712029 | 0,803314 | 0,7668 | 0,654215 | 0,730285 | 0,739414 | 0,775929 |
| | Primary somatosensory area upper limb layer 6a | -3,38 | *0,1904* | 0,8485 | 0,0447 | 7 | 0,8198 | 0,0308 | 7 | 0,818528 | 0,861129 | 0,833743 | 0,882429 | 0,836786 | 0,785057 | 0,921986 | 0,824615 | 0,824614 | 0,772885 | 0,861129 | 0,806357 | 0,852 | 0,797229 |
| | Primary somatosensory area upper limb layer 6b | -3,55 | *0,6341* | 0,0613 | 0,0057 | 7 | 0,0591 | 0,0102 | 7 | 0,060857 | 0,0639 | 0,057814 | 0,051729 | 0,069986 | 0,0639 | 0,060857 | 0,048686 | 0,060857 | 0,060857 | 0,057814 | 0,079114 | 0,048686 | 0,057814 |
| | Primary somatosensory area trunk | -9,86 | *0,3242* | 1,43 | 0,26 | 7 | 1,29 | 0,26 | 7 | 1,634014 | 1,676615 | 1,104558 | 1,503172 | 1,652271 | 1,058915 | 1,396672 | 1,335815 | 0,918943 | 1,141071 | 1,518386 | 1,147157 | 1,290171 | 1,685743 |
| | Primary somatosensory area trunk layer 1 | -10,04 | *0,3412* | 0,2034 | 0,0363 | 7 | 0,1830 | 0,0407 | 7 | 0,231257 | 0,213 | 0,155186 | 0,225172 | 0,243429 | 0,152143 | 0,203871 | 0,194743 | 0,164314 | 0,143014 | 0,173443 | 0,143014 | 0,203871 | 0,258643 |

| | | | | | | | | | | | | | | | | | | | | | | | |
|---|---|---|---|---|---|---|---|---|---|---|---|---|---|---|---|---|---|---|---|---|---|---|---|
| | Primary somatosensory area trunk layer 2/3 | -7,91 | *0,4085* | 0,4343 | 0,0728 | 7 | 0,3999 | 0,0771 | 7 | 0,495986 | 0,4686 | 0,328629 | 0,465557 | 0,514243 | 0,343843 | 0,422957 | 0,413829 | 0,273857 | 0,380357 | 0,480772 | 0,380357 | 0,365143 | 0,505114 |
| | Primary somatosensory area trunk layer 4 | -9,18 | *0,3987* | 0,1704 | 0,0332 | 7 | 0,1548 | 0,0337 | 7 | 0,194743 | 0,203872 | 0,133886 | 0,185614 | 0,194743 | 0,118671 | 0,161271 | 0,155186 | 0,115629 | 0,136929 | 0,200829 | 0,130843 | 0,143014 | 0,200829 |
| | Primary somatosensory area trunk layer 5 | -11,12 | *0,2731* | 0,3673 | 0,0656 | 7 | 0,3265 | 0,0675 | 7 | 0,407743 | 0,4473 | 0,282986 | 0,374272 | 0,416871 | 0,2769 | 0,365143 | 0,352972 | 0,213 | 0,2982 | 0,392529 | 0,279943 | 0,343843 | 0,4047 |
| | Primary somatosensory area trunk layer 6a | -11,59 | *0,3560* | 0,2213 | 0,0534 | 7 | 0,1956 | 0,0462 | 7 | 0,264728 | 0,304286 | 0,182571 | 0,213 | 0,231257 | 0,139971 | 0,213 | 0,179529 | 0,136929 | 0,161271 | 0,243429 | 0,176486 | 0,203871 | 0,267771 |
| | Primary somatosensory area trunk layer 6b | -12,20 | *0,4628* | 0,0356 | 0,0100 | 7 | 0,0313 | 0,0113 | 7 | 0,039557 | 0,039557 | 0,0213 | 0,039557 | 0,051729 | 0,027386 | 0,030429 | 0,039557 | 0,015214 | 0,0213 | 0,027386 | 0,036514 | 0,030429 | 0,048686 |
| | Primary somatosensory area unassigned | -2,25 | *0,3634* | 1,20 | 0,06 | 7 | 1,17 | 0,05 | 7 | 1,159328 | 1,201929 | 1,162372 | 1,217144 | 1,229314 | 1,122815 | 1,302343 | 1,204972 | 1,138029 | 1,131942 | 1,153243 | 1,201928 | 1,125856 | 1,250614 |
| | Primary somatosensory area unassigned layer 1 | 3,13 | *0,6165* | 0,1526 | 0,0180 | 7 | 0,1574 | 0,0167 | 7 | 0,130843 | 0,161272 | 0,139971 | 0,133886 | 0,158229 | 0,164314 | 0,179529 | 0,188657 | 0,143014 | 0,164314 | 0,152143 | 0,164314 | 0,139971 | 0,1491 |
| | Primary somatosensory area unassigned layer 2/3 | -0,40 | *0,9146* | 0,3247 | 0,0234 | 7 | 0,3234 | 0,0211 | 7 | 0,325586 | 0,343843 | 0,295157 | 0,301243 | 0,328629 | 0,316457 | 0,3621 | 0,349929 | 0,286029 | 0,310371 | 0,334714 | 0,337757 | 0,316457 | 0,328629 |
| | Primary somatosensory area unassigned layer 4 | -6,77 | *0,0757* | 0,1926 | 0,0133 | 7 | 0,1795 | 0,0117 | 7 | 0,200829 | 0,213 | 0,179529 | 0,182572 | 0,188657 | 0,179529 | 0,203871 | 0,164314 | 0,173443 | 0,1704 | 0,176486 | 0,188657 | 0,185614 | 0,197786 |
| | Primary somatosensory area unassigned layer 5 | -1,58 | *0,7126* | 0,2469 | 0,0202 | 7 | 0,2430 | 0,0185 | 7 | 0,243428 | 0,219086 | 0,246472 | 0,279943 | 0,261686 | 0,228214 | 0,249514 | 0,243429 | 0,246471 | 0,237343 | 0,2343 | 0,240386 | 0,219086 | 0,279943 |
| | Primary somatosensory area unassigned layer 6a | -4,15 | *0,4538* | 0,2621 | 0,0316 | 7 | 0,2513 | 0,0190 | 7 | 0,240386 | 0,243429 | 0,273857 | 0,307329 | 0,273857 | 0,213 | 0,282986 | 0,246472 | 0,279943 | 0,2343 | 0,240386 | 0,237343 | 0,243428 | 0,2769 |
| | Primary somatosensory area unassigned layer 6b | -12,77 | *0,4761* | 0,0204 | 0,0049 | 7 | 0,0178 | 0,0079 | 7 | 0,018257 | 0,0213 | 0,027386 | 0,012171 | 0,018257 | 0,0213 | 0,024343 | 0,012171 | 0,009129 | 0,015214 | 0,015214 | 0,033471 | 0,0213 | 0,018257 |
| * | Supplemental somatosensory area | -4,46 | *0,0426* | 8,21 | 0,28 | 7 | 7,84 | 0,32 | 7 | 8,291783 | 8,136604 | 8,084874 | 8,310048 | 8,057486 | 7,832318 | 8,723872 | 7,862747 | 8,10313 | 7,741026 | 7,360674 | 7,558453 | 7,935767 | 8,313086 |
| * | Supplemental somatosensory area layer 1 | -8,51 | *0,0124* | 1,16 | 0,07 | 7 | 1,06 | 0,05 | 7 | 1,238442 | 1,189758 | 1,138029 | 1,150201 | 1,131943 | 1,0437 | 1,2567 | 1,128901 | 1,046743 | 1,061957 | 0,995015 | 1,034571 | 1,058914 | 1,1289 |
| | Supplemental somatosensory area layer 2/3 | -4,28 | *0,1273* | 1,94 | 0,09 | 7 | 1,86 | 0,10 | 7 | 2,005242 | 1,86223 | 1,862229 | 2,038716 | 1,8744 | 1,871358 | 2,069143 | 1,764858 | 1,920043 | 1,786157 | 1,743558 | 1,850056 | 1,95047 | 1,986986 |
| | Supplemental somatosensory area layer 4 | -4,88 | *0,0718* | 1,09 | 0,05 | 7 | 1,03 | 0,05 | 7 | 1,104557 | 1,043701 | 1,071086 | 1,165415 | 1,074129 | 1,019358 | 1,122814 | 1,043701 | 1,083257 | 1,007185 | 0,9585 | 1,016314 | 1,010228 | 1,110643 |
| | Supplemental somatosensory area layer 5 | -2,95 | *0,3015* | 1,99 | 0,07 | 7 | 1,93 | 0,12 | 7 | 2,014371 | 1,965687 | 2,020458 | 1,983944 | 1,996114 | 1,843972 | 2,0874 | 1,944387 | 2,072186 | 1,901785 | 1,773986 | 1,773985 | 1,974813 | 2,060014 |
| | Supplemental somatosensory area layer 6a | -3,95 | *0,0813* | 1,85 | 0,08 | 7 | 1,77 | 0,06 | 7 | 1,749642 | 1,907872 | 1,813543 | 1,792244 | 1,801371 | 1,871358 | 1,990029 | 1,795287 | 1,798329 | 1,792242 | 1,691829 | 1,716171 | 1,755728 | 1,865271 |
| | Supplemental somatosensory area layer 6b | 0,48 | *0,8868* | 0,1808 | 0,0089 | 7 | 0,1817 | 0,0130 | 7 | 0,179529 | 0,167357 | 0,179529 | 0,179529 | 0,179529 | 0,182572 | 0,197786 | 0,185614 | 0,182571 | 0,1917 | 0,197786 | 0,167357 | 0,185614 | 0,161271 |
| | Gustatory areas | -0,85 | *0,6201* | 1,64 | 0,06 | 7 | 1,63 | 0,03 | 7 | 1,658357 | 1,597501 | 1,524472 | 1,664444 | 1,6401 | 1,673572 | 1,7253 | 1,618801 | 1,655315 | 1,652271 | 1,612715 | 1,564028 | 1,63097 | 1,652271 |
| | Gustatory areas layer 1 | -0,72 | *0,9285* | 0,1817 | 0,0309 | 7 | 0,1804 | 0,0213 | 7 | 0,219086 | 0,152143 | 0,130843 | 0,203872 | 0,179529 | 0,185614 | 0,200829 | 0,1917 | 0,194743 | 0,197786 | 0,164314 | 0,155186 | 0,203871 | 0,155186 |
| | Gustatory areas layer 2/3 | -6,70 | *0,1103* | 0,3825 | 0,0121 | 7 | 0,3569 | 0,0354 | 7 | 0,386443 | 0,380357 | 0,380357 | 0,3621 | 0,389486 | 0,401657 | 0,377314 | 0,337757 | 0,337757 | 0,322543 | 0,401657 | 0,395571 | 0,3195 | 0,3834 |
| | Gustatory areas layer 4 | -7,19 | *0,3712* | 0,1452 | 0,0154 | 7 | 0,1348 | 0,0251 | 7 | 0,146057 | 0,164314 | 0,133886 | 0,1491 | 0,133886 | 0,124757 | 0,164314 | 0,118671 | 0,1278 | 0,139971 | 0,176486 | 0,130843 | 0,097371 | 0,152143 |
| | Gustatory areas layer 5 | 6,42 | *0,2925* | 0,4743 | 0,0490 | 7 | 0,5047 | 0,0542 | 7 | 0,410786 | 0,450343 | 0,477729 | 0,514243 | 0,559886 | 0,456429 | 0,450343 | 0,480772 | 0,547714 | 0,471643 | 0,4473 | 0,453385 | 0,584228 | 0,547714 |
| | Gustatory areas layer 6a | 0,62 | *0,9259* | 0,4203 | 0,0589 | 7 | 0,4230 | 0,0423 | 7 | 0,453386 | 0,413829 | 0,365143 | 0,4047 | 0,337757 | 0,456429 | 0,5112 | 0,456429 | 0,422957 | 0,499028 | 0,392529 | 0,401657 | 0,413828 | 0,374271 |
| | Gustatory areas layer 6b | -26,19 | *0,0653* | 0,0365 | 0,0088 | 7 | 0,0270 | 0,0089 | 7 | 0,0426 | 0,036514 | 0,036514 | 0,030429 | 0,039557 | 0,048686 | 0,0213 | 0,033471 | 0,024343 | 0,0213 | 0,030429 | 0,027386 | 0,012171 | 0,039557 |
| | Visceral area | -5,82 | *0,1300* | 2,15 | 0,17 | 7 | 2,03 | 0,12 | 7 | 2,327785 | 2,212158 | 1,983944 | 2,005244 | 2,0235 | 2,105658 | 2,397772 | 2,060015 | 1,807457 | 2,078271 | 2,041758 | 2,072185 | 1,947427 | 2,1726 |
| | Visceral area layer 1 | -0,71 | *0,8918* | 0,3069 | 0,0300 | 7 | 0,3047 | 0,0285 | 7 | 0,343843 | 0,325586 | 0,279943 | 0,2769 | 0,273857 | 0,310372 | 0,337757 | 0,352972 | 0,264729 | 0,301243 | 0,322543 | 0,279943 | 0,307328 | 0,304286 |
| | Visceral area layer 2/3 | -6,68 | *0,2177* | 0,5012 | 0,0536 | 7 | 0,4677 | 0,0416 | 7 | 0,587271 | 0,517286 | 0,435129 | 0,465557 | 0,450343 | 0,520329 | 0,5325 | 0,450343 | 0,380357 | 0,486857 | 0,492943 | 0,499028 | 0,486857 | 0,477729 |
| | Visceral area layer 4 | -7,47 | *0,2234* | 0,1630 | 0,0139 | 7 | 0,1508 | 0,0206 | 7 | 0,167357 | 0,158229 | 0,161271 | 0,139972 | 0,158229 | 0,1704 | 0,185614 | 0,167357 | 0,124757 | 0,161271 | 0,1278 | 0,161271 | 0,136928 | 0,176486 |
| * | Visceral area layer 5 | -8,22 | *0,0421* | 0,6607 | 0,0524 | 7 | 0,6064 | 0,0327 | 7 | 0,684643 | 0,690729 | 0,6177 | 0,6177 | 0,657257 | 0,605529 | 0,751586 | 0,632915 | 0,581186 | 0,626828 | 0,611614 | 0,5751 | 0,565971 | 0,651171 |
| | Visceral area layer 6a | -6,27 | *0,1820* | 0,4712 | 0,0410 | 7 | 0,4416 | 0,0369 | 7 | 0,502071 | 0,477729 | 0,426 | 0,480772 | 0,432086 | 0,441214 | 0,538586 | 0,398615 | 0,416871 | 0,459471 | 0,441214 | 0,465557 | 0,407743 | 0,502071 |
| | Visceral area layer 6b | 13,64 | *0,4558* | 0,0478 | 0,0129 | 7 | 0,0543 | 0,0182 | 7 | 0,0426 | 0,0426 | 0,0639 | 0,024343 | 0,051729 | 0,057814 | 0,051729 | 0,057814 | 0,039557 | 0,0426 | 0,045643 | 0,091286 | 0,0426 | 0,060857 |
| * | Auditory areas | -5,37 | *0,0372* | 5,35 | 0,21 | 7 | 5,06 | 0,24 | 7 | 5,440627 | 5,349346 | 5,18503 | 5,358475 | 5,108957 | 5,248931 | 5,757086 | 5,230674 | 4,999415 | 5,026798 | 4,69513 | 4,908126 | 5,108954 | 5,468014 |
| * | Dorsal auditory area | -9,02 | *0,0484* | 1,15 | 0,06 | 7 | 1,04 | 0,11 | 7 | 1,204971 | 1,168458 | 1,147158 | 1,180629 | 1,122814 | 1,028486 | 1,174543 | 1,101515 | 1,089343 | 1,031528 | 0,928072 | 0,991971 | 0,934157 | 1,226271 |
| | Dorsal auditory area layer 1 | -9,35 | *0,1172* | 0,1952 | 0,0191 | 7 | 0,1769 | 0,0213 | 7 | 0,185614 | 0,179529 | 0,1917 | 0,182572 | 0,182571 | 0,216043 | 0,228214 | 0,1917 | 0,176486 | 0,197786 | 0,152143 | 0,143014 | 0,185614 | 0,1917 |
| | Dorsal auditory area layer 2/3 | -2,28 | *0,6469* | 0,3047 | 0,0150 | 7 | 0,2978 | 0,0357 | 7 | 0,3195 | 0,316457 | 0,2982 | 0,289072 | 0,282986 | 0,307329 | 0,3195 | 0,295157 | 0,301243 | 0,328628 | 0,252557 | 0,264728 | 0,286028 | 0,356014 |
| | Dorsal auditory area layer 4 | 0,00 | *1,0000* | 0,1230 | 0,0098 | 7 | 0,1230 | 0,0112 | 7 | 0,124757 | 0,118671 | 0,118671 | 0,136929 | 0,124757 | 0,1065 | 0,130843 | 0,139972 | 0,124757 | 0,118671 | 0,121714 | 0,121714 | 0,103457 | 0,130843 |
| | Dorsal auditory area layer 5 | -10,92 | *0,0642* | 0,3304 | 0,0294 | 7 | 0,2943 | 0,0362 | 7 | 0,3621 | 0,343843 | 0,331672 | 0,346886 | 0,337757 | 0,270814 | 0,3195 | 0,313414 | 0,316457 | 0,270814 | 0,258643 | 0,295157 | 0,252557 | 0,352971 |
| * | Dorsal auditory area layer 6a | -21,09 | *0,0292* | 0,1669 | 0,0296 | 7 | 0,1317 | 0,0226 | 7 | 0,182571 | 0,188657 | 0,179529 | 0,194743 | 0,164314 | 0,109543 | 0,1491 | 0,143014 | 0,143014 | 0,109543 | 0,1278 | 0,146057 | 0,094329 | 0,158229 |
| | Dorsal auditory area layer 6b | -26,23 | *0,1358* | 0,0265 | 0,0049 | 7 | 0,0196 | 0,0101 | 7 | 0,030429 | 0,0213 | 0,027386 | 0,030429 | 0,030429 | 0,018257 | 0,027386 | 0,018257 | 0,027386 | 0,006086 | 0,015214 | 0,0213 | 0,012171 | 0,036514 |
| | Primary auditory area | -0,93 | *0,7925* | 1,95 | 0,13 | 7 | 1,94 | 0,12 | 7 | 1,923085 | 1,868315 | 1,956558 | 2,069144 | 1,731386 | 1,999158 | 2,133043 | 1,965687 | 1,947429 | 1,913956 | 1,743558 | 1,831799 | 2,07827 | 2,072186 |
| | Primary auditory area layer 1 | -5,98 | *0,3496* | 0,3343 | 0,0343 | 7 | 0,3143 | 0,0421 | 7 | 0,3195 | 0,313414 | 0,343843 | 0,3834 | 0,2769 | 0,349929 | 0,352971 | 0,331672 | 0,325586 | 0,2769 | 0,246472 | 0,3621 | 0,301243 | 0,356014 |
| * | Primary auditory area layer 2/3 | -7,88 | *0,0317* | 0,4247 | 0,0263 | 7 | 0,3912 | 0,0252 | 7 | 0,407743 | 0,395572 | 0,426 | 0,474686 | 0,4047 | 0,432086 | 0,432086 | 0,401657 | 0,365143 | 0,380357 | 0,395572 | 0,365143 | 0,392528 | 0,438171 |
| | Primary auditory area layer 4 | 0,64 | *0,8806* | 0,2030 | 0,0096 | 7 | 0,2043 | 0,0201 | 7 | 0,197786 | 0,200829 | 0,1917 | 0,206914 | 0,200829 | 0,200829 | 0,222129 | 0,182572 | 0,216043 | 0,213 | 0,179529 | 0,1917 | 0,2343 | 0,213 |
| | Primary auditory area layer 5 | 0,85 | *0,8461* | 0,6633 | 0,0488 | 7 | 0,6690 | 0,0573 | 7 | 0,651171 | 0,632915 | 0,6603 | 0,693772 | 0,5964 | 0,657257 | 0,751586 | 0,678558 | 0,672472 | 0,699857 | 0,590314 | 0,605528 | 0,760714 | 0,675514 |
| | Primary auditory area layer 6a | 9,39 | *0,1172* | 0,2778 | 0,0319 | 7 | 0,3039 | 0,0254 | 7 | 0,295157 | 0,2769 | 0,279943 | 0,264729 | 0,216043 | 0,295157 | 0,316457 | 0,316457 | 0,310371 | 0,301243 | 0,279943 | 0,261686 | 0,334714 | 0,322543 |
| | Primary auditory area layer 6b | 4,24 | *0,6392* | 0,0513 | 0,0089 | 7 | 0,0535 | 0,0080 | 7 | 0,051729 | 0,048686 | 0,054771 | 0,045643 | 0,036514 | 0,0639 | 0,057814 | 0,054771 | 0,057814 | 0,0426 | 0,051729 | 0,045643 | 0,054771 | 0,066943 |
| | Posterior auditory area | -11,46 | *0,1245* | 0,5499 | 0,0683 | 7 | 0,4869 | 0,0743 | 7 | 0,584228 | 0,535543 | 0,517286 | 0,632915 | 0,629871 | 0,453386 | 0,495986 | 0,450343 | 0,629872 | 0,419914 | 0,416872 | 0,4686 | 0,495985 | 0,526414 |
| | Posterior auditory area layer 1 | -6,16 | *0,7495* | 0,0917 | 0,0248 | 7 | 0,0861 | 0,0382 | 7 | 0,079114 | 0,069986 | 0,094329 | 0,1065 | 0,121714 | 0,115629 | 0,054771 | 0,057814 | 0,136929 | 0,057814 | 0,030429 | 0,094329 | 0,109543 | 0,115629 |
| | Posterior auditory area layer 2/3 | -0,33 | *0,9817* | 0,1300 | 0,0173 | 7 | 0,1295 | 0,0452 | 7 | 0,146057 | 0,124757 | 0,121714 | 0,1491 | 0,143014 | 0,124757 | 0,100414 | 0,082157 | 0,1917 | 0,073029 | 0,100414 | 0,1704 | 0,143014 | 0,146057 |
| | Posterior auditory area layer 4 | 2,84 | *0,8716* | 0,0613 | 0,0160 | 7 | 0,0630 | 0,0227 | 7 | 0,057814 | 0,060857 | 0,066943 | 0,088243 | 0,069986 | 0,0426 | 0,0426 | 0,045643 | 0,100414 | 0,0426 | 0,045643 | 0,057814 | 0,060857 | 0,088243 |
| | Posterior auditory area layer 5 | -18,62 | *0,1105* | 0,1704 | 0,0374 | 7 | 0,1387 | 0,0310 | 7 | 0,197786 | 0,185614 | 0,155186 | 0,1917 | 0,188657 | 0,091286 | 0,182571 | 0,161272 | 0,164314 | 0,146057 | 0,167357 | 0,0852 | 0,109543 | 0,136929 |
| * | Posterior auditory area layer 6a | -31,35 | *0,0113* | 0,0804 | 0,0111 | 7 | 0,0552 | 0,0184 | 7 | 0,0852 | 0,076071 | 0,069986 | 0,082157 | 0,091286 | 0,0639 | 0,094329 | 0,079114 | 0,033471 | 0,079114 | 0,051729 | 0,048686 | 0,057814 | 0,036514 |
| | Posterior auditory area layer 6b | -10,81 | *0,6419* | 0,0161 | 0,0038 | 7 | 0,0143 | 0,0087 | 7 | 0,018257 | 0,018257 | 0,009129 | 0,015214 | 0,015214 | 0,015214 | 0,0213 | 0,024343 | 0,003043 | 0,0213 | 0,0213 | 0,012171 | 0,015214 | 0,003043 |
| | Ventral auditory area | -6,04 | *0,2013* | 1,70 | 0,16 | 7 | 1,60 | 0,12 | 7 | 1,728342 | 1,77703 | 1,564029 | 1,475787 | 1,624886 | 1,767901 | 1,953514 | 1,71313 | 1,332772 | 1,661399 | 1,606629 | 1,615756 | 1,600542 | 1,643143 |
| | Ventral auditory area layer 1 | 0,17 | *0,9780* | 0,2517 | 0,0339 | 7 | 0,2521 | 0,0225 | 7 | 0,2556 | 0,2343 | 0,279943 | 0,182572 | 0,270814 | 0,273857 | 0,267771 | 0,252557 | 0,261686 | 0,240386 | 0,225172 | 0,237343 | 0,295157 | 0,252557 |
| | Ventral auditory area layer 2/3 | -8,55 | *0,1613* | 0,3812 | 0,0454 | 7 | 0,3486 | 0,0354 | 7 | 0,398614 | 0,4047 | 0,398614 | 0,282986 | 0,386443 | 0,377314 | 0,419914 | 0,356014 | 0,301243 | 0,346886 | 0,304286 | 0,359057 | 0,398614 | 0,374271 |
| | Ventral auditory area layer 4 | -6,17 | *0,4770* | 0,1974 | 0,0330 | 7 | 0,1852 | 0,0288 | 7 | 0,182571 | 0,197786 | 0,1491 | 0,1704 | 0,222129 | 0,213 | 0,246471 | 0,1704 | 0,161271 | 0,203871 | 0,194743 | 0,222128 | 0,139971 | 0,203871 |
| | Ventral auditory area layer 5 | -3,98 | *0,4534* | 0,5577 | 0,0597 | 7 | 0,5355 | 0,0463 | 7 | 0,569014 | 0,5751 | 0,499029 | 0,5325 | 0,477729 | 0,599443 | 0,651171 | 0,578143 | 0,441214 | 0,538586 | 0,578143 | 0,544671 | 0,523371 | 0,544671 |
| | Ventral auditory area layer 6a | -10,49 | *0,3092* | 0,2652 | 0,0384 | 7 | 0,2373 | 0,0572 | 7 | 0,267771 | 0,304286 | 0,209957 | 0,273857 | 0,222129 | 0,264729 | 0,313414 | 0,310372 | 0,139971 | 0,289071 | 0,264729 | 0,206914 | 0,216043 | 0,2343 |
| | Ventral auditory area layer 6b | -18,10 | *0,1633* | 0,0456 | 0,0122 | 7 | 0,0374 | 0,0080 | 7 | 0,054771 | 0,060857 | 0,027386 | 0,033471 | 0,045643 | 0,0426 | 0,054771 | 0,045643 | 0,027386 | 0,0426 | 0,039557 | 0,045643 | 0,027386 | 0,033471 |
| | Visual areas | -2,89 | *0,2557* | 12,60 | 0,59 | 7 | 12,24 | 0,55 | 7 | 12,65524 | 12,28402 | 11,86715 | 12,99301 | 12,02537 | 12,83478 | 13,53767 | 12,46051 | 12,1836 | 11,94625 | 11,66936 | 11,69065 | 12,44224 | 13,25773 |
| | Anterolateral visual area | -11,23 | *0,1165* | 0,7277 | 0,0615 | 7 | 0,6460 | 0,1090 | 7 | 0,727243 | 0,651172 | 0,721157 | 0,833743 | 0,7455 | 0,754629 | 0,6603 | 0,569015 | 0,8307 | 0,587271 | 0,584229 | 0,562928 | 0,614657 | 0,772886 |
| | Anterolateral visual area layer 1 | 2,87 | *0,8625* | 0,1061 | 0,0307 | 7 | 0,1091 | 0,0336 | 7 | 0,082157 | 0,073029 | 0,133886 | 0,094329 | 0,088243 | 0,158229 | 0,112586 | 0,112586 | 0,136929 | 0,115629 | 0,054771 | 0,076071 | 0,152143 | 0,115629 |
| | Anterolateral visual area layer 2/3 | -8,55 | *0,5785* | 0,1830 | 0,0578 | 7 | 0,1674 | 0,0436 | 7 | 0,164314 | 0,152143 | 0,161271 | 0,194743 | 0,167357 | 0,307329 | 0,133886 | 0,130843 | 0,2343 | 0,136929 | 0,136929 | 0,152143 | 0,155186 | 0,225171 |
| | Anterolateral visual area layer 4 | 1,40 | *0,9113* | 0,0935 | 0,0086 | 7 | 0,0948 | 0,0286 | 7 | 0,091286 | 0,100414 | 0,0852 | 0,088243 | 0,109543 | 0,091286 | 0,088243 | 0,073029 | 0,139971 | 0,076071 | 0,0852 | 0,088243 | 0,069986 | 0,130843 |
| | Anterolateral visual area layer 5 | -13,10 | *0,3277* | 0,1991 | 0,0522 | 7 | 0,1730 | 0,0429 | 7 | 0,219086 | 0,173443 | 0,194743 | 0,267772 | 0,246471 | 0,109543 | 0,182571 | 0,139972 | 0,231257 | 0,152143 | 0,173443 | 0,1491 | 0,130843 | 0,2343 |
| * | Anterolateral visual area layer 6a | -28,57 | *0,0077* | 0,1217 | 0,0240 | 7 | 0,0869 | 0,0124 | 7 | 0,136929 | 0,121714 | 0,121714 | 0,155186 | 0,121714 | 0,076071 | 0,118671 | 0,094329 | 0,079114 | 0,091286 | 0,103457 | 0,079114 | 0,094329 | 0,066943 |

| | | | | | | | | | | | | | | | | | | | | | | | |
|---|---|---|---|---|---|---|---|---|---|---|---|---|---|---|---|---|---|---|---|---|---|---|---|
| | Anterolateral visual area layer 6b | -29,17 | *0,1493* | 0,0243 | 0,0091 | 7 | 0,0172 | 0,0074 | 6 | 0,033471 | 0,030429 | 0,024343 | 0,033471 | 0,012171 | 0,012171 | 0,024343 | 0,018257 | 0,009129 | 0,015214 | 0,030429 | 0,018257 | 0,012171 | |
| | Anteromedial visual area | 4,15 | *0,6178* | 0,7546 | 0,1281 | 7 | 0,7859 | 0,0981 | 7 | 0,715071 | 0,708986 | 0,8094 | 0,590315 | 0,632914 | 0,912858 | 0,912857 | 0,842872 | 0,751586 | 0,848957 | 0,629872 | 0,812442 | 0,915899 | 0,699857 |
| | Anteromedial visual area layer 1 | 16,04 | *0,1366* | 0,1274 | 0,0195 | 7 | 0,1478 | 0,0274 | 7 | 0,118671 | 0,109543 | 0,1491 | 0,109543 | 0,115629 | 0,130843 | 0,158229 | 0,194743 | 0,124757 | 0,124757 | 0,130843 | 0,146057 | 0,176486 | 0,136929 |
| | Anteromedial visual area layer 2/3 | 5,61 | *0,5847* | 0,2091 | 0,0389 | 7 | 0,2208 | 0,0393 | 7 | 0,200829 | 0,197786 | 0,213 | 0,155186 | 0,179529 | 0,249514 | 0,267771 | 0,240386 | 0,188657 | 0,258643 | 0,164314 | 0,228214 | 0,270814 | 0,194743 |
| | Anteromedial visual area layer 4 | 0,00 | *1,0000* | 0,0565 | 0,0155 | 7 | 0,0565 | 0,0136 | 7 | 0,0426 | 0,048686 | 0,0639 | 0,048686 | 0,039557 | 0,079114 | 0,073029 | 0,054771 | 0,048686 | 0,076071 | 0,054771 | 0,060857 | 0,066943 | 0,033471 |
| | Anteromedial visual area layer 5 | -3,29 | *0,7142* | 0,2247 | 0,0425 | 7 | 0,2173 | 0,0300 | 7 | 0,206914 | 0,206914 | 0,237343 | 0,173443 | 0,188657 | 0,282986 | 0,2769 | 0,222129 | 0,219086 | 0,237343 | 0,161271 | 0,216043 | 0,258643 | 0,206914 |
| | Anteromedial visual area layer 6a | 2,96 | *0,6711* | 0,1174 | 0,0161 | 7 | 0,1208 | 0,0137 | 7 | 0,130843 | 0,118671 | 0,121714 | 0,088243 | 0,1065 | 0,136929 | 0,118671 | 0,118671 | 0,124757 | 0,136929 | 0,094329 | 0,124757 | 0,130843 | 0,115629 |
| | Anteromedial visual area layer 6b | 15,56 | *0,6417* | 0,0196 | 0,0099 | 7 | 0,0226 | 0,0136 | 7 | 0,015214 | 0,027386 | 0,024343 | 0,015214 | 0,003043 | 0,033471 | 0,018257 | 0,012171 | 0,045643 | 0,015214 | 0,024343 | 0,036514 | 0,012171 | 0,012171 |
| | Lateral visual area | -1,94 | *0,7111* | 1,19 | 0,13 | 7 | 1,16 | 0,10 | 7 | 1,244528 | 1,198886 | 1,119772 | 1,125858 | 0,976757 | 1,244529 | 1,387543 | 1,250615 | 1,040657 | 1,250614 | 1,195843 | 1,028485 | 1,232356 | 1,138029 |
| | Lateral visual area layer 1 | -2,79 | *0,7439* | 0,1713 | 0,0215 | 7 | 0,1665 | 0,0310 | 7 | 0,209957 | 0,173443 | 0,146057 | 0,185614 | 0,1704 | 0,155186 | 0,158229 | 0,143014 | 0,176486 | 0,139971 | 0,203871 | 0,176486 | 0,124757 | 0,200829 |
| | Lateral visual area layer 2/3 | -8,44 | *0,2836* | 0,3039 | 0,0502 | 7 | 0,2782 | 0,0329 | 7 | 0,334714 | 0,325586 | 0,313414 | 0,310372 | 0,264729 | 0,213 | 0,365143 | 0,2769 | 0,264729 | 0,304286 | 0,286029 | 0,216043 | 0,3195 | 0,279943 |
| | Lateral visual area layer 4 | -6,94 | *0,5226* | 0,1817 | 0,0226 | 7 | 0,1691 | 0,0447 | 7 | 0,1704 | 0,182572 | 0,179529 | 0,173443 | 0,155186 | 0,182572 | 0,228214 | 0,216043 | 0,118671 | 0,225171 | 0,167357 | 0,136929 | 0,197786 | 0,121714 |
| | Lateral visual area layer 5 | -2,28 | *0,8288* | 0,3047 | 0,0634 | 7 | 0,2978 | 0,0540 | 7 | 0,307328 | 0,301243 | 0,273857 | 0,2982 | 0,194743 | 0,377314 | 0,380357 | 0,365143 | 0,219086 | 0,349928 | 0,325586 | 0,273857 | 0,304286 | 0,246471 |
| | Lateral visual area layer 6a | 8,97 | *0,4013* | 0,1891 | 0,0438 | 7 | 0,2060 | 0,0264 | 7 | 0,185614 | 0,182572 | 0,176486 | 0,136929 | 0,158229 | 0,273857 | 0,209957 | 0,203872 | 0,219086 | 0,1917 | 0,173443 | 0,179528 | 0,243428 | 0,231257 |
| * | Lateral visual area layer 6b | 28,75 | *0,0242* | 0,0348 | 0,0080 | 7 | 0,0448 | 0,0063 | 7 | 0,036514 | 0,033471 | 0,030429 | 0,0213 | 0,033471 | 0,0426 | 0,045643 | 0,045643 | 0,0426 | 0,039557 | 0,039557 | 0,045643 | 0,0426 | 0,057814 |
| | Primary visual area | -3,26 | *0,2518* | 6,57 | 0,30 | 7 | 6,35 | 0,36 | 7 | 6,581698 | 6,441732 | 6,091802 | 6,925547 | 6,423472 | 6,523888 | 6,968143 | 6,399132 | 6,313929 | 6,110055 | 5,991388 | 6,28654 | 6,256111 | 7,102029 |
| | Primary visual area layer 1 | -0,52 | *0,9539* | 1,16 | 0,07 | 7 | 1,15 | 0,26 | 7 | 1,1502 | 1,101515 | 1,113686 | 1,162372 | 1,2993 | 1,204972 | 1,0863 | 0,982843 | 1,3419 | 0,964585 | 0,9798 | 1,159328 | 0,998057 | 1,649229 |
| | Primary visual area layer 2/3 | -7,54 | *0,0526* | 1,90 | 0,15 | 7 | 1,76 | 0,09 | 7 | 1,977857 | 1,920044 | 1,770943 | 2,136087 | 1,871357 | 1,682701 | 1,9596 | 1,81963 | 1,707043 | 1,694871 | 1,700958 | 1,673571 | 1,80137 | 1,917 |
| | Primary visual area layer 4 | -3,81 | *0,2941* | 0,9572 | 0,0705 | 7 | 0,9207 | 0,0522 | 7 | 0,985885 | 0,991972 | 0,836786 | 0,991972 | 0,909814 | 0,931115 | 1,052829 | 0,995015 | 0,818529 | 0,921985 | 0,918943 | 0,9372 | 0,931114 | 0,921986 |
| | Primary visual area layer 5 | 2,84 | *0,5787* | 1,42 | 0,17 | 7 | 1,46 | 0,08 | 7 | 1,357114 | 1,439272 | 1,372329 | 1,317558 | 1,208014 | 1,527515 | 1,731386 | 1,545772 | 1,332772 | 1,524471 | 1,402758 | 1,451442 | 1,524471 | 1,454486 |
| | Primary visual area layer 6a | -7,26 | *0,2178* | 0,9759 | 0,1170 | 7 | 0,9050 | 0,0826 | 7 | 0,973714 | 0,812443 | 0,8733 | 1,180629 | 1,007186 | 1,019358 | 0,964586 | 0,882429 | 0,982843 | 0,833743 | 0,8307 | 0,921985 | 0,839828 | 1,0437 |
| | Primary visual area layer 6b | 1,76 | *0,8238* | 0,1478 | 0,0214 | 7 | 0,1504 | 0,0214 | 7 | 0,136929 | 0,176486 | 0,124757 | 0,136929 | 0,1278 | 0,158229 | 0,173443 | 0,173443 | 0,130843 | 0,1704 | 0,158229 | 0,143014 | 0,161271 | 0,115629 |
| | Posterolateral visual area | -16,18 | *0,0564* | 0,7629 | 0,1369 | 7 | 0,6394 | 0,0460 | 7 | 0,778971 | 0,7668 | 0,629872 | 0,937201 | 0,7242 | 0,5751 | 0,928071 | 0,678558 | 0,602486 | 0,684643 | 0,690729 | 0,608571 | 0,635957 | 0,5751 |
| * | Posterolateral visual area layer 1 | -51,27 | *0,0461* | 0,1713 | 0,0875 | 7 | 0,0835 | 0,0519 | 7 | 0,155186 | 0,124757 | 0,1278 | 0,2982 | 0,2769 | 0,051729 | 0,164314 | 0,069986 | 0,194743 | 0,079114 | 0,082157 | 0,0639 | 0,030429 | 0,0639 |
| * | Posterolateral visual area layer 2/3 | -54,41 | *0,0226* | 0,2708 | 0,1252 | 7 | 0,1235 | 0,0692 | 7 | 0,377314 | 0,240386 | 0,258643 | 0,413829 | 0,161271 | 0,073029 | 0,371229 | 0,133886 | 0,036514 | 0,222128 | 0,164314 | 0,161271 | 0,112586 | 0,033471 |
| * | Posterolateral visual area layer 4 | 73,11 | *0,0284* | 0,0259 | 0,0137 | 6 | 0,0448 | 0,0130 | 7 | 0,012171 | 0,036514 | 0,015214 | | 0,015214 | 0,045643 | 0,030429 | 0,051729 | 0,039557 | 0,036514 | 0,066943 | 0,027386 | 0,039557 | 0,051729 |
| | Posterolateral visual area layer 5 | 58,04 | *0,0559* | 0,2000 | 0,1103 | 7 | 0,3160 | 0,0937 | 7 | 0,185614 | 0,337757 | 0,100414 | 0,076071 | 0,1065 | 0,273857 | 0,3195 | 0,413829 | 0,185614 | 0,334714 | 0,3621 | 0,2556 | 0,429043 | 0,231257 |
| | Posterolateral visual area layer 6a | -17,19 | *0,6951* | 0,0835 | 0,0545 | 7 | 0,0691 | 0,0769 | 7 | 0,033471 | 0,0213 | 0,109543 | 0,109543 | 0,152143 | 0,130843 | 0,027386 | 0,006086 | 0,146057 | 0,009129 | 0,009129 | 0,094329 | 0,024343 | 0,194743 |
| * | Posterolateral visual area layer 6b | -74,29 | *0,0365* | 0,0178 | 0,0115 | 6 | 0,0046 | 0,0018 | 4 | 0,015214 | 0,006086 | 0,018257 | 0,039557 | 0,012171 | | 0,015214 | 0,003043 | | 0,003043 | 0,006086 | 0,006086 | | |
| | posteromedial visual area | -6,34 | *0,1480* | 0,9941 | 0,0804 | 7 | 0,9311 | 0,0717 | 7 | 1,052828 | 0,952415 | 0,921986 | 1,131944 | 1,0224 | 0,903729 | 0,973714 | 0,903729 | 0,9585 | 0,864171 | 0,9372 | 0,867214 | 0,912857 | 1,074129 |
| | posteromedial visual area layer 1 | -5,24 | *0,2548* | 0,1826 | 0,0181 | 7 | 0,1730 | 0,0105 | 7 | 0,1704 | 0,152143 | 0,188657 | 0,1917 | 0,185614 | 0,179529 | 0,209957 | 0,158229 | 0,179529 | 0,188657 | 0,161271 | 0,173443 | 0,176486 | 0,173443 |
| | posteromedial visual area layer 2/3 | -7,41 | *0,2907* | 0,2817 | 0,0253 | 7 | 0,2608 | 0,0425 | 7 | 0,292114 | 0,273857 | 0,258643 | 0,3195 | 0,307329 | 0,267772 | 0,252557 | 0,209957 | 0,270814 | 0,240386 | 0,270814 | 0,2343 | 0,2556 | 0,343843 |
| | posteromedial visual area layer 4 | -6,94 | *0,3517* | 0,0939 | 0,0062 | 7 | 0,0874 | 0,0163 | 7 | 0,097371 | 0,097371 | 0,091286 | 0,103457 | 0,094329 | 0,0852 | 0,088243 | 0,097371 | 0,094329 | 0,060857 | 0,094329 | 0,076071 | 0,079114 | 0,109543 |
| | posteromedial visual area layer 5 | -3,71 | *0,5311* | 0,2695 | 0,0320 | 7 | 0,2595 | 0,0256 | 7 | 0,286028 | 0,258643 | 0,237343 | 0,322543 | 0,289071 | 0,231257 | 0,261686 | 0,2982 | 0,2769 | 0,222128 | 0,270814 | 0,237343 | 0,249514 | 0,261686 |
| | posteromedial visual area layer 6a | -8,56 | *0,3103* | 0,1421 | 0,0234 | 7 | 0,1300 | 0,0193 | 7 | 0,173443 | 0,139972 | 0,115629 | 0,1704 | 0,1278 | 0,118671 | 0,1491 | 0,124757 | 0,115629 | 0,1278 | 0,1065 | 0,136929 | 0,130843 | 0,167357 |
| | posteromedial visual area layer 6b | -16,07 | *0,3563* | 0,0243 | 0,0077 | 7 | 0,0204 | 0,0076 | 7 | 0,033471 | 0,030429 | 0,030429 | 0,024343 | 0,018257 | 0,0213 | 0,012171 | 0,015214 | 0,0213 | 0,024343 | 0,033471 | 0,009129 | 0,0213 | 0,018257 |
| | Laterointermediate area | 1,59 | *0,8106* | 0,4634 | 0,0593 | 7 | 0,4708 | 0,0534 | 7 | 0,453386 | 0,459472 | 0,486857 | 0,429043 | 0,371229 | 0,477729 | 0,565971 | 0,556843 | 0,422957 | 0,486857 | 0,450343 | 0,401657 | 0,514243 | 0,462514 |
| | Laterointermediate area layer 1 | -10,80 | *0,4244* | 0,0765 | 0,0231 | 7 | 0,0682 | 0,0123 | 7 | 0,097371 | 0,0852 | 0,066943 | 0,097371 | 0,054771 | 0,039557 | 0,094329 | 0,082157 | 0,054771 | 0,082157 | 0,079114 | 0,060857 | 0,060857 | 0,057814 |
| | Laterointermediate area layer 2/3 | -3,28 | *0,7714* | 0,1191 | 0,0255 | 7 | 0,1152 | 0,0237 | 7 | 0,1065 | 0,133886 | 0,1278 | 0,115629 | 0,091286 | 0,094329 | 0,164314 | 0,158229 | 0,088243 | 0,121714 | 0,118671 | 0,100414 | 0,124757 | 0,094329 |
| | Laterointermediate area layer 4 | -4,35 | *0,7209* | 0,0600 | 0,0161 | 7 | 0,0574 | 0,0097 | 7 | 0,060857 | 0,060857 | 0,0639 | 0,033471 | 0,045643 | 0,076071 | 0,079114 | 0,066943 | 0,048686 | 0,060857 | 0,066943 | 0,0426 | 0,0639 | 0,051729 |
| | Laterointermediate area layer 5 | 13,38 | *0,3378* | 0,1300 | 0,0349 | 7 | 0,1474 | 0,0300 | 7 | 0,112586 | 0,1065 | 0,139971 | 0,1065 | 0,103457 | 0,200829 | 0,139971 | 0,1704 | 0,118671 | 0,143014 | 0,118671 | 0,121714 | 0,194743 | 0,164314 |
| | Laterointermediate area layer 6a | 5,33 | *0,5175* | 0,0652 | 0,0068 | 7 | 0,0687 | 0,0119 | 7 | 0,0639 | 0,0639 | 0,076071 | 0,057814 | 0,060857 | 0,060857 | 0,073029 | 0,0639 | 0,088243 | 0,0639 | 0,057814 | 0,066943 | 0,057814 | 0,082157 |
| | Laterointermediate area layer 6b | 10,34 | *0,6133* | 0,0126 | 0,0041 | 7 | 0,0139 | 0,0052 | 7 | 0,012171 | 0,009129 | 0,012171 | 0,018257 | 0,015214 | 0,006086 | 0,015214 | 0,015214 | 0,024343 | 0,015214 | 0,009129 | 0,009129 | 0,012171 | 0,012171 |
| | Postrhinal area | 8,95 | *0,1593* | 1,15 | 0,14 | 7 | 1,25 | 0,12 | 7 | 1,101514 | 1,104558 | 1,0863 | 1,019358 | 1,1289 | 1,442315 | 1,141072 | 1,259744 | 1,262786 | 1,113685 | 1,189758 | 1,122814 | 1,360156 | 1,433186 |
| | Postrhinal area layer 1 | 7,40 | *0,5629* | 0,2408 | 0,0605 | 7 | 0,2586 | 0,0511 | 7 | 0,282986 | 0,261686 | 0,237343 | 0,182572 | 0,136929 | 0,279943 | 0,304286 | 0,337757 | 0,179529 | 0,301243 | 0,246472 | 0,225171 | 0,2556 | 0,264729 |
| | Postrhinal area layer 2/3 | 8,79 | *0,3562* | 0,3412 | 0,0637 | 7 | 0,3712 | 0,0525 | 7 | 0,343843 | 0,343843 | 0,286029 | 0,286029 | 0,322543 | 0,474686 | 0,331671 | 0,3408 | 0,337757 | 0,334714 | 0,377314 | 0,349928 | 0,374271 | 0,483814 |
| | Postrhinal area layer 4 | 21,60 | *0,2388* | 0,0543 | 0,0131 | 7 | 0,0661 | 0,0211 | 7 | 0,051729 | 0,045643 | 0,060857 | 0,0426 | 0,057814 | 0,079114 | 0,0426 | 0,045643 | 0,073029 | 0,045643 | 0,048686 | 0,0639 | 0,088243 | 0,097371 |
| | Postrhinal area layer 5 | 0,90 | *0,8824* | 0,3391 | 0,0169 | 7 | 0,3421 | 0,0498 | 7 | 0,3195 | 0,325586 | 0,352972 | 0,3195 | 0,359057 | 0,349929 | 0,346886 | 0,322543 | 0,435129 | 0,304286 | 0,334714 | 0,282986 | 0,371228 | 0,343843 |
| | Postrhinal area layer 6a | 33,33 | *0,1128* | 0,1395 | 0,0604 | 7 | 0,1860 | 0,0369 | 7 | 0,088243 | 0,1065 | 0,115629 | 0,136929 | 0,216043 | 0,231257 | 0,082157 | 0,1917 | 0,197786 | 0,121714 | 0,167357 | 0,173443 | 0,237343 | 0,213 |
| | Postrhinal area layer 6b | -20,83 | *0,3136* | 0,0313 | 0,0117 | 7 | 0,0248 | 0,0114 | 7 | 0,015214 | 0,0213 | 0,033471 | 0,051729 | 0,036514 | 0,027386 | 0,033471 | 0,0213 | 0,039557 | 0,006086 | 0,015214 | 0,027386 | 0,033471 | 0,030429 |
| | Anterior cingulate area | 1,28 | *0,6631* | 5,21 | 0,29 | 7 | 5,28 | 0,26 | 7 | 5,008541 | 4,801631 | 5,382816 | 5,248932 | 4,981157 | 5,629288 | 5,443672 | 5,063317 | 5,394986 | 4,972027 | 5,075487 | 5,656669 | 5,255011 | 5,544086 |
| | Anterior cingulate area dorsal part | 0,86 | *0,7764* | 2,95 | 0,20 | 7 | 2,97 | 0,12 | 7 | 2,811599 | 2,638159 | 3,121972 | 3,01243 | 2,8116 | 3,191959 | 3,027643 | 2,933316 | 2,972872 | 2,875499 | 2,845072 | 3,198041 | 2,957655 | 3,009386 |
| | Anterior cingulate area dorsal part layer 1 | -0,80 | *0,8377* | 0,6003 | 0,0438 | 7 | 0,5955 | 0,0416 | 7 | 0,569014 | 0,535543 | 0,648129 | 0,6177 | 0,565971 | 0,648129 | 0,6177 | 0,593357 | 0,593357 | 0,562928 | 0,538586 | 0,672471 | 0,605528 | 0,602486 |
| | Anterior cingulate area dorsal part layer 2/3 | -1,42 | *0,7120* | 0,5829 | 0,0486 | 7 | 0,5747 | 0,0308 | 7 | 0,587271 | 0,505115 | 0,620743 | 0,608572 | 0,5325 | 0,642043 | 0,584229 | 0,5538 | 0,559886 | 0,565971 | 0,538586 | 0,632914 | 0,584228 | 0,587271 |
| | Anterior cingulate area dorsal part layer 5 | 2,28 | *0,4027* | 1,12 | 0,07 | 7 | 1,15 | 0,03 | 7 | 1,055871 | 1,019358 | 1,183672 | 1,162372 | 1,110643 | 1,214101 | 1,113686 | 1,144115 | 1,180629 | 1,1076 | 1,122815 | 1,201928 | 1,128899 | 1,153243 |
| | Anterior cingulate area dorsal part layer 6a | 1,46 | *0,6487* | 0,6234 | 0,0479 | 7 | 0,6325 | 0,0172 | 7 | 0,584229 | 0,569015 | 0,651172 | 0,608572 | 0,590314 | 0,6603 | 0,699857 | 0,626829 | 0,620743 | 0,620743 | 0,623786 | 0,666385 | 0,623785 | 0,645086 |
| | Anterior cingulate area dorsal part layer 6b | 22,22 | *0,2095* | 0,0156 | 0,0059 | 7 | 0,0191 | 0,0034 | 7 | 0,015214 | 0,009129 | 0,018257 | 0,015214 | 0,012171 | 0,027386 | 0,012171 | 0,015214 | 0,018257 | 0,018257 | 0,0213 | 0,024343 | 0,015214 | 0,0213 |
| | Anterior cingulate area ventral part | 1,82 | *0,6014* | 2,27 | 0,11 | 7 | 2,31 | 0,17 | 7 | 2,196942 | 2,163473 | 2,260844 | 2,236501 | 2,169557 | 2,43733 | 2,416029 | 2,130001 | 2,422115 | 2,096528 | 2,230415 | 2,458627 | 2,297356 | 2,5347 |
| | Anterior cingulate area ventral part layer 1 | -0,83 | *0,8002* | 0,4738 | 0,0237 | 7 | 0,4699 | 0,0321 | 7 | 0,4473 | 0,4686 | 0,508157 | 0,465557 | 0,4473 | 0,499029 | 0,480771 | 0,432086 | 0,505114 | 0,426 | 0,465557 | 0,480771 | 0,471643 | 0,508157 |
| | Anterior cingulate area ventral part layer 2/3 | 3,23 | *0,4088* | 0,4986 | 0,0348 | 7 | 0,5147 | 0,0355 | 7 | 0,480771 | 0,471643 | 0,483814 | 0,492943 | 0,465557 | 0,5538 | 0,541629 | 0,4686 | 0,547714 | 0,477728 | 0,4899 | 0,535543 | 0,526414 | 0,556843 |
| | Anterior cingulate area ventral part layer 5 | 2,57 | *0,5562* | 0,9146 | 0,0541 | 7 | 0,9381 | 0,0865 | 7 | 0,897643 | 0,861129 | 0,897643 | 0,894601 | 0,870257 | 0,9798 | 1,0011 | 0,858086 | 0,991972 | 0,836785 | 0,885472 | 1,010228 | 0,915899 | 1,068043 |
| | Anterior cingulate area ventral part 6a | 3,11 | *0,3868* | 0,3212 | 0,0135 | 7 | 0,3312 | 0,0258 | 7 | 0,316457 | 0,301243 | 0,322543 | 0,3195 | 0,3195 | 0,346886 | 0,322543 | 0,3195 | 0,325586 | 0,295157 | 0,334714 | 0,377314 | 0,3195 | 0,346886 |
| | Anterior cingulate area ventral part 6b | -7,19 | *0,2133* | 0,0604 | 0,0073 | 7 | 0,0561 | 0,0046 | 7 | 0,054771 | 0,060857 | 0,048686 | 0,0639 | 0,066943 | 0,057814 | 0,069986 | 0,051729 | 0,051729 | 0,060857 | 0,054771 | 0,054771 | 0,0639 | 0,054771 |
| | Prelimbic area | -5,78 | *0,1065* | 2,21 | 0,11 | 7 | 2,08 | 0,16 | 7 | 2,172599 | 2,072187 | 2,346044 | 2,361259 | 2,2365 | 2,178687 | 2,120872 | 2,047844 | 1,986986 | 1,889614 | 2,230415 | 2,139128 | 1,965685 | 2,333871 |
| | Prelimbic area layer 1 | -5,70 | *0,1299* | 0,4803 | 0,0287 | 7 | 0,4530 | 0,0339 | 7 | 0,471643 | 0,438172 | 0,456429 | 0,502072 | 0,471643 | 0,505115 | 0,517286 | 0,486857 | 0,450343 | 0,419914 | 0,429043 | 0,483814 | 0,410785 | 0,4899 |
| | Prelimbic area layer 2/3 | -9,37 | *0,1064* | 0,4082 | 0,0209 | 7 | 0,3699 | 0,0515 | 7 | 0,4047 | 0,395572 | 0,395572 | 0,432086 | 0,441214 | 0,4047 | 0,3834 | 0,374272 | 0,286029 | 0,322543 | 0,401657 | 0,392528 | 0,371228 | 0,441214 |
| | Prelimbic area layer 5 | -3,47 | *0,4498* | 0,8889 | 0,0827 | 7 | 0,8581 | 0,0636 | 7 | 0,906771 | 0,8094 | 1,028486 | 0,946329 | 0,903729 | 0,806357 | 0,821571 | 0,818529 | 0,827657 | 0,815485 | 0,9585 | 0,858085 | 0,794185 | 0,934157 |
| | Prelimbic area layer 6a | -7,03 | *0,1810* | 0,4143 | 0,0334 | 7 | 0,3851 | 0,0425 | 7 | 0,371228 | 0,416872 | 0,435129 | 0,456429 | 0,407743 | 0,441214 | 0,371229 | 0,343843 | 0,407743 | 0,3195 | 0,416872 | 0,392528 | 0,374271 | 0,441214 |
| | Prelimbic area layer 6b | -10,42 | *0,5604* | 0,0209 | 0,0071 | 7 | 0,0187 | 0,0064 | 7 | 0,018257 | 0,012171 | 0,030429 | 0,024343 | 0,012171 | 0,0213 | 0,027386 | 0,024343 | 0,015214 | 0,012171 | 0,024343 | 0,012171 | 0,015214 | 0,027386 |

| | | | | | | | | | | | | | | | | | | | | | | | |
|---|---|---|---|---|---|---|---|---|---|---|---|---|---|---|---|---|---|---|---|---|---|---|---|
| | Infralimbic area | -12,88 | *0,1701* | 0,8337 | 0,1200 | 7 | 0,7264 | 0,1524 | 7 | 0,943285 | 0,934158 | 0,815486 | 0,885472 | 0,909814 | 0,721157 | 0,626829 | 0,590315 | 0,693772 | 0,569014 | 0,973715 | 0,708985 | 0,654214 | 0,8946 |
| | Infralimbic area layer 1 | -12,71 | *0,4600* | 0,1574 | 0,0426 | 7 | 0,1374 | 0,0545 | 7 | 0,228214 | 0,185614 | 0,124757 | 0,118671 | 0,1704 | 0,164314 | 0,109543 | 0,130843 | 0,121714 | 0,082157 | 0,240386 | 0,173443 | 0,088243 | 0,124757 |
| | Infralimbic area layer 2/3 | -23,53 | *0,3039* | 0,1256 | 0,0431 | 7 | 0,0961 | 0,0583 | 7 | 0,1917 | 0,146057 | 0,118671 | 0,112586 | 0,139971 | 0,121714 | 0,048686 | 0,0852 | 0,060857 | 0,033471 | 0,203871 | 0,1278 | 0,048686 | 0,112586 |
| | Infralimbic area layer 5 | -22,21 | *0,1262* | 0,3582 | 0,0958 | 7 | 0,2786 | 0,0848 | 7 | 0,407743 | 0,4473 | 0,349929 | 0,426 | 0,426 | 0,225172 | 0,225171 | 0,1917 | 0,243429 | 0,225171 | 0,410786 | 0,2343 | 0,258643 | 0,386443 |
| | Infralimbic area layer 6a | 12,02 | *0,4695* | 0,1808 | 0,0454 | 7 | 0,2026 | 0,0619 | 7 | 0,100414 | 0,1491 | 0,216043 | 0,222129 | 0,167357 | 0,188657 | 0,222129 | 0,164314 | 0,261686 | 0,209957 | 0,103457 | 0,161271 | 0,249514 | 0,267771 |
| | Infralimbic area layer 6b | 0,00 | *1,0000* | 0,0117 | 0,0073 | 7 | 0,0117 | 0,0059 | 7 | 0,015214 | 0,006086 | 0,006086 | 0,006086 | 0,006086 | 0,0213 | 0,0213 | 0,018257 | 0,006086 | 0,018257 | 0,015214 | 0,012171 | 0,009129 | 0,003043 |
| | Orbital area | -0,46 | *0,8967* | 5,24 | 0,43 | 7 | 5,21 | 0,20 | 7 | 4,704256 | 4,783374 | 5,303702 | 5,309789 | 5,042014 | 5,638417 | 5,872715 | 5,188074 | 5,413244 | 5,075484 | 4,841187 | 5,32804 | 5,270226 | 5,370643 |
| | Orbital area lateral part | -0,41 | *0,9194* | 2,33 | 0,15 | 7 | 2,32 | 0,19 | 7 | 2,081314 | 2,154344 | 2,428201 | 2,443416 | 2,394729 | 2,349087 | 2,446457 | 2,154344 | 2,437329 | 2,193899 | 2,120872 | 2,221285 | 2,473841 | 2,629029 |
| | Orbital area lateral part layer 1 | -4,43 | *0,7751* | 0,2847 | 0,0914 | 7 | 0,2721 | 0,0680 | 7 | 0,213 | 0,273857 | 0,441214 | 0,380357 | 0,209957 | 0,213 | 0,261686 | 0,203872 | 0,310371 | 0,325586 | 0,188657 | 0,213 | 0,307328 | 0,356014 |
| | Orbital area lateral part layer 2/3 | 24,90 | *0,2845* | 0,5464 | 0,2579 | 7 | 0,6825 | 0,1893 | 7 | 0,541628 | 0,410786 | 0,416872 | 0,337757 | 0,307329 | 0,852 | 0,9585 | 0,721158 | 0,769843 | 0,882428 | 0,562929 | 0,730285 | 0,797228 | 0,313414 |
| | Orbital area lateral part layer 5 | -22,21 | *0,1575* | 1,14 | 0,30 | 7 | 0,8863 | 0,3295 | 7 | 1,083257 | 1,168458 | 1,229315 | 1,405801 | 1,548814 | 0,824615 | 0,715071 | 0,815486 | 0,687686 | 0,517286 | 1,040657 | 0,9159 | 0,699857 | 1,527514 |
| | Orbital area lateral part layer 6a | 23,89 | *0,1568* | 0,3238 | 0,0443 | 7 | 0,4012 | 0,1224 | 7 | 0,243428 | 0,301243 | 0,3408 | 0,3195 | 0,328629 | 0,346886 | 0,386443 | 0,2982 | 0,550757 | 0,3621 | 0,289072 | 0,2982 | 0,578143 | 0,432086 |
| | Orbital area medial part | 6,32 | *0,4290* | 1,24 | 0,22 | 7 | 1,32 | 0,12 | 7 | 1,028485 | 1,083258 | 1,229315 | 1,204972 | 1,080214 | 1,457529 | 1,627929 | 1,402758 | 1,354072 | 1,417971 | 1,1076 | 1,433185 | 1,329728 | 1,217143 |
| | Orbital area medial part layer 1 | 2,67 | *0,8256* | 0,3738 | 0,0839 | 7 | 0,3838 | 0,0821 | 7 | 0,252557 | 0,359057 | 0,435129 | 0,450343 | 0,295157 | 0,346886 | 0,477729 | 0,365143 | 0,5112 | 0,453386 | 0,258643 | 0,325586 | 0,386443 | 0,386443 |
| | Orbital area medial part layer 2/3 | 23,99 | *0,2860* | 0,2482 | 0,1166 | 7 | 0,3078 | 0,0776 | 7 | 0,1917 | 0,1704 | 0,167357 | 0,1704 | 0,2343 | 0,322543 | 0,480771 | 0,337757 | 0,331671 | 0,343843 | 0,1917 | 0,307328 | 0,419914 | 0,222129 |
| | Orbital area medial part layer 5 | 33,30 | *0,1358* | 0,3956 | 0,1817 | 7 | 0,5273 | 0,1150 | 7 | 0,398614 | 0,2343 | 0,3408 | 0,313414 | 0,200829 | 0,687686 | 0,593357 | 0,614657 | 0,480771 | 0,608571 | 0,465557 | 0,687685 | 0,483814 | 0,349929 |
| * | Orbital area medial part layer 6a | -55,56 | *0,0386* | 0,2269 | 0,1077 | 7 | 0,1008 | 0,0946 | 7 | 0,185614 | 0,3195 | 0,286029 | 0,270814 | 0,349929 | 0,100414 | 0,076071 | 0,0852 | 0,015214 | 0,012171 | 0,1917 | 0,112586 | 0,030429 | 0,258643 |
| | Orbital area ventrolateral part | -5,59 | *0,1062* | 1,66 | 0,11 | 7 | 1,57 | 0,08 | 7 | 1,594457 | 1,545772 | 1,646186 | 1,661401 | 1,567071 | 1,831801 | 1,798329 | 1,630972 | 1,621843 | 1,463614 | 1,612715 | 1,673571 | 1,466656 | 1,524471 |
| | Orbital area ventrolateral part layer 1 | 0,23 | *0,9880* | 0,3699 | 0,0993 | 7 | 0,3708 | 0,1117 | 7 | 0,286028 | 0,282986 | 0,407743 | 0,426 | 0,237343 | 0,4899 | 0,459471 | 0,395572 | 0,477729 | 0,474686 | 0,313414 | 0,416871 | 0,3621 | 0,155186 |
| | Orbital area ventrolateral part layer 2/3 | 10,28 | *0,2751* | 0,4356 | 0,0925 | 7 | 0,4803 | 0,0415 | 7 | 0,401657 | 0,392529 | 0,343843 | 0,352972 | 0,444257 | 0,514243 | 0,599443 | 0,508157 | 0,514243 | 0,407743 | 0,441214 | 0,477728 | 0,517285 | 0,495986 |
| | Orbital area ventrolateral part layer 5 | -10,60 | *0,0972* | 0,5864 | 0,0519 | 7 | 0,5242 | 0,0742 | 7 | 0,666386 | 0,581186 | 0,632914 | 0,5751 | 0,517286 | 0,5964 | 0,535543 | 0,562929 | 0,492943 | 0,432086 | 0,629872 | 0,587271 | 0,441214 | 0,523371 |
| | Orbital area ventrolateral part layer 6a | -28,64 | *0,0514* | 0,2717 | 0,0551 | 7 | 0,1939 | 0,0763 | 7 | 0,240386 | 0,289072 | 0,261686 | 0,307329 | 0,368186 | 0,231257 | 0,203871 | 0,164314 | 0,130843 | 0,1491 | 0,228214 | 0,1917 | 0,143014 | 0,349929 |
| | Agranular insular area | -0,59 | *0,7543* | 7,26 | 0,30 | 7 | 7,21 | 0,17 | 7 | 7,202441 | 7,059432 | 6,974231 | 7,14159 | 7,047257 | 7,640618 | 7,734943 | 7,047261 | 7,318072 | 7,105069 | 7,105074 | 7,071596 | 7,372839 | 7,482386 |
| | Agranular insular area dorsal part | -0,81 | *0,6894* | 3,44 | 0,16 | 7 | 3,41 | 0,09 | 7 | 3,42017 | 3,322802 | 3,380615 | 3,414088 | 3,240643 | 3,599702 | 3,687943 | 3,338016 | 3,362358 | 3,426256 | 3,334973 | 3,438427 | 3,587527 | 3,383657 |
| | Agranular insular area dorsal part layer 1 | 3,63 | *0,6102* | 0,3947 | 0,0605 | 7 | 0,4090 | 0,0394 | 7 | 0,343843 | 0,325586 | 0,492943 | 0,444257 | 0,346886 | 0,407743 | 0,401657 | 0,374272 | 0,422957 | 0,438171 | 0,356014 | 0,377314 | 0,435128 | 0,459471 |
| | Agranular insular area dorsal part layer 2/3 | 8,61 | *0,2828* | 0,8377 | 0,1454 | 7 | 0,9098 | 0,0838 | 7 | 0,8094 | 0,782015 | 0,754629 | 0,742458 | 0,690729 | 1,013272 | 1,071086 | 0,961543 | 0,925029 | 0,967628 | 0,870257 | 0,903728 | 0,995014 | 0,7455 |
| * | Agranular insular area dorsal part layer 5 | -7,10 | *0,0095* | 1,45 | 0,04 | 7 | 1,35 | 0,07 | 7 | 1,494042 | 1,509258 | 1,417972 | 1,478829 | 1,390586 | 1,436229 | 1,430143 | 1,281044 | 1,268872 | 1,311471 | 1,366243 | 1,351028 | 1,372328 | 1,484914 |
| | Agranular insular area dorsal part layer 6a | 0,62 | *0,8367* | 0,7012 | 0,0384 | 7 | 0,7055 | 0,0387 | 7 | 0,6816 | 0,654215 | 0,663343 | 0,699858 | 0,751586 | 0,708986 | 0,748543 | 0,693772 | 0,705943 | 0,6816 | 0,705943 | 0,7455 | 0,760714 | 0,645086 |
| | Agranular insular area dorsal part layer 6b | -29,27 | *0,1039* | 0,0535 | 0,0192 | 7 | 0,0378 | 0,0132 | 7 | 0,091286 | 0,051729 | 0,051729 | 0,048686 | 0,060857 | 0,033471 | 0,036514 | 0,027386 | 0,039557 | 0,027386 | 0,036514 | 0,060857 | 0,024343 | 0,048686 |
| | Agranular insular area posterior part | -0,33 | *0,8951* | 2,12 | 0,10 | 7 | 2,11 | 0,10 | 7 | 2,126956 | 2,087401 | 1,999158 | 1,996115 | 2,163471 | 2,218244 | 2,239543 | 2,038715 | 2,175643 | 2,011328 | 2,111744 | 2,09957 | 2,053927 | 2,291271 |
| | Agranular insular area posterior part layer 1 | 0,23 | *0,9358* | 0,3738 | 0,0212 | 7 | 0,3747 | 0,0182 | 7 | 0,389486 | 0,377314 | 0,368186 | 0,337757 | 0,3621 | 0,377314 | 0,4047 | 0,374272 | 0,365143 | 0,386443 | 0,371229 | 0,368186 | 0,349928 | 0,407743 |
| | Agranular insular area posterior part layer 2/3 | -7,00 | *0,0586* | 0,7020 | 0,0457 | 7 | 0,6529 | 0,0421 | 7 | 0,663343 | 0,648129 | 0,712029 | 0,678558 | 0,7668 | 0,687686 | 0,757671 | 0,611615 | 0,678557 | 0,578143 | 0,684643 | 0,684643 | 0,654214 | 0,678557 |
| | Agranular insular area posterior part layer 5 | 4,08 | *0,2296* | 0,6499 | 0,0406 | 7 | 0,6764 | 0,0377 | 7 | 0,690728 | 0,639 | 0,578143 | 0,629872 | 0,657257 | 0,699857 | 0,654214 | 0,663343 | 0,7029 | 0,648128 | 0,678557 | 0,663343 | 0,632914 | 0,7455 |
| | Agranular insular area posterior part layer 6a | 3,69 | *0,4861* | 0,3773 | 0,0412 | 7 | 0,3912 | 0,0301 | 7 | 0,365143 | 0,413829 | 0,325586 | 0,3408 | 0,352971 | 0,432086 | 0,410786 | 0,3834 | 0,413829 | 0,380357 | 0,356014 | 0,365143 | 0,395571 | 0,444257 |
| | Agranular insular area posterior part layer 6b | 5,56 | *0,7763* | 0,0156 | 0,0059 | 7 | 0,0165 | 0,0052 | 7 | 0,018257 | 0,009129 | 0,015214 | 0,009129 | 0,024343 | 0,0213 | 0,012171 | 0,006086 | 0,015214 | 0,018257 | 0,0213 | 0,018257 | 0,0213 | 0,015214 |
| | Agranular insular area ventral part | -0,46 | *0,8730* | 1,70 | 0,09 | 7 | 1,69 | 0,09 | 7 | 1,655314 | 1,649229 | 1,594458 | 1,731387 | 1,643143 | 1,822672 | 1,807457 | 1,670529 | 1,780072 | 1,667485 | 1,658358 | 1,533599 | 1,731385 | 1,807457 |
| | Agranular insular area ventral part layer 1 | 6,65 | *0,3091* | 0,2417 | 0,0351 | 7 | 0,2578 | 0,0180 | 7 | 0,219086 | 0,225172 | 0,216043 | 0,2343 | 0,213 | 0,292114 | 0,292114 | 0,237343 | 0,282986 | 0,258643 | 0,252557 | 0,2343 | 0,273857 | 0,264729 |
| * | Agranular insular area ventral part layer 2/3 | -10,70 | *0,0073* | 0,6173 | 0,0396 | 7 | 0,5512 | 0,0369 | 7 | 0,608571 | 0,590315 | 0,611614 | 0,672472 | 0,6603 | 0,620743 | 0,556843 | 0,587272 | 0,5538 | 0,520328 | 0,605529 | 0,502071 | 0,529457 | 0,559886 |
| | Agranular insular area ventral part layer 5 | 9,49 | *0,3002* | 0,6412 | 0,0613 | 7 | 0,7020 | 0,1324 | 7 | 0,550757 | 0,5964 | 0,611614 | 0,663343 | 0,635957 | 0,721157 | 0,708986 | 0,544672 | 0,827657 | 0,654214 | 0,599443 | 0,602485 | 0,8094 | 0,876343 |
| | Agranular insular area ventral part layer 6a | -8,57 | *0,6300* | 0,1978 | 0,0545 | 7 | 0,1808 | 0,0723 | 7 | 0,273857 | 0,231257 | 0,152143 | 0,158229 | 0,130843 | 0,188657 | 0,249514 | 0,301243 | 0,115629 | 0,2343 | 0,197786 | 0,1917 | 0,118671 | 0,1065 |
| | Retrosplenial area | -2,60 | *0,1221* | 9,72 | 0,34 | 7 | 9,47 | 0,19 | 7 | 9,956225 | 9,785834 | 9,39026 | 9,670206 | 9,417643 | 9,499804 | 10,34876 | 9,715848 | 9,575873 | 9,24724 | 9,28376 | 9,308095 | 9,502837 | 9,667157 |
| * | Retrosplenial area lateral agranular part | -6,43 | *0,0463* | 2,23 | 0,13 | 7 | 2,08 | 0,11 | 7 | 2,434285 | 2,133044 | 2,163472 | 2,343001 | 2,181729 | 2,053929 | 2,269972 | 2,063058 | 2,306486 | 1,959599 | 2,032629 | 2,072185 | 2,050885 | 2,093486 |
| | Retrosplenial area lateral agranular part layer 1 | -9,28 | *0,2768* | 0,4873 | 0,0772 | 7 | 0,4421 | 0,0711 | 7 | 0,520328 | 0,456429 | 0,480772 | 0,541629 | 0,611614 | 0,395572 | 0,4047 | 0,416872 | 0,541629 | 0,371228 | 0,450343 | 0,4047 | 0,374271 | 0,535543 |
| * | Retrosplenial area lateral agranular part layer 2/3 | -19,06 | *0,0071* | 0,6729 | 0,0869 | 7 | 0,5447 | 0,0276 | 7 | 0,772885 | 0,584229 | 0,651172 | 0,803315 | 0,678557 | 0,578143 | 0,642043 | 0,547715 | 0,587272 | 0,499028 | 0,556843 | 0,556843 | 0,538585 | 0,526414 |
| | Retrosplenial area lateral agranular part layer 5 | 1,86 | *0,7974* | 0,7012 | 0,1100 | 7 | 0,7142 | 0,0711 | 7 | 0,794185 | 0,782015 | 0,6177 | 0,666386 | 0,5538 | 0,639 | 0,855043 | 0,775929 | 0,651172 | 0,775928 | 0,748543 | 0,730285 | 0,733328 | 0,584229 |
| | Retrosplenial area lateral agranular part layer 6a | 4,42 | *0,6935* | 0,3347 | 0,0466 | 7 | 0,3495 | 0,0842 | 7 | 0,325586 | 0,273857 | 0,386443 | 0,310372 | 0,310371 | 0,407743 | 0,328629 | 0,289072 | 0,483814 | 0,282986 | 0,246472 | 0,352971 | 0,368186 | 0,422957 |
| | Retrosplenial area lateral agranular part layer 6b | 8,82 | *0,4770* | 0,0296 | 0,0072 | 7 | 0,0322 | 0,0060 | 7 | 0,0213 | 0,036514 | 0,027386 | 0,0213 | 0,027386 | 0,033471 | 0,039557 | 0,033471 | 0,0426 | 0,030429 | 0,030429 | 0,027386 | 0,036514 | 0,024343 |
| | Retrosplenial area dorsal part | -2,25 | *0,1800* | 3,51 | 0,11 | 7 | 3,44 | 0,09 | 7 | 3,557099 | 3,639259 | 3,313673 | 3,526674 | 3,511457 | 3,426259 | 3,627086 | 3,468859 | 3,411043 | 3,374527 | 3,386701 | 3,350184 | 3,432341 | 3,624043 |
| | Retrosplenial area dorsal part layer 1 | -6,17 | *0,1602* | 0,7885 | 0,0743 | 7 | 0,7398 | 0,0399 | 7 | 0,803314 | 0,861129 | 0,645086 | 0,7881 | 0,848957 | 0,7455 | 0,827657 | 0,754629 | 0,708986 | 0,748543 | 0,803315 | 0,705943 | 0,690728 | 0,7668 |
| | Retrosplenial area dorsal part layer 2/3 | -0,71 | *0,7997* | 0,9163 | 0,0568 | 7 | 0,9098 | 0,0339 | 7 | 0,9372 | 0,967629 | 0,864172 | 0,852001 | 0,900686 | 0,885472 | 1,007186 | 0,970672 | 0,885472 | 0,903728 | 0,888515 | 0,882428 | 0,894599 | 0,943286 |
| | Retrosplenial area dorsal part layer 5 | 0,28 | *0,9093* | 1,10 | 0,02 | 7 | 1,11 | 0,06 | 7 | 1,098471 | 1,138029 | 1,095429 | 1,116729 | 1,080214 | 1,074129 | 1,1289 | 1,141072 | 1,016314 | 1,101514 | 1,058915 | 1,071085 | 1,192799 | 1,1715 |
| | Retrosplenial area dorsal part layer 6a | -3,17 | *0,4746* | 0,6712 | 0,0358 | 7 | 0,6499 | 0,0666 | 7 | 0,6816 | 0,626829 | 0,687686 | 0,733329 | 0,651171 | 0,678557 | 0,639 | 0,578143 | 0,760714 | 0,5964 | 0,605529 | 0,663343 | 0,632914 | 0,712029 |
| | Retrosplenial area dorsal part layer 6b | -16,67 | *0,1970* | 0,0339 | 0,0090 | 7 | 0,0283 | 0,0060 | 7 | 0,036514 | 0,045643 | 0,0213 | 0,036514 | 0,030429 | 0,0426 | 0,024343 | 0,024343 | 0,039557 | 0,024343 | 0,030429 | 0,027386 | 0,0213 | 0,030429 |
| | Retrosplenial area ventral part | -0,76 | *0,7647* | 3,98 | 0,23 | 7 | 3,95 | 0,12 | 7 | 3,964842 | 4,013531 | 3,913116 | 3,800531 | 3,724457 | 4,019616 | 4,4517 | 4,183931 | 3,858343 | 3,913113 | 3,86443 | 3,885727 | 4,019612 | 3,949629 |
| | Retrosplenial area ventral part layer 1 | -0,92 | *0,7700* | 0,9007 | 0,0650 | 7 | 0,8924 | 0,0321 | 7 | 0,8946 | 0,943286 | 0,9159 | 0,821572 | 0,806357 | 0,943286 | 0,9798 | 0,934158 | 0,882429 | 0,861128 | 0,909815 | 0,900685 | 0,915899 | 0,842871 |
| | Retrosplenial area ventral part layer 2/3 | -0,51 | *0,8260* | 0,9337 | 0,0528 | 7 | 0,9289 | 0,0172 | 7 | 0,964585 | 0,937201 | 0,891557 | 0,882429 | 0,900686 | 0,925029 | 1,034571 | 0,955458 | 0,912857 | 0,909814 | 0,9372 | 0,9159 | 0,943285 | 0,928071 |
| | Retrosplenial area ventral part layer 5 | -0,11 | *0,9700* | 1,53 | 0,11 | 7 | 1,52 | 0,05 | 7 | 1,497085 | 1,512301 | 1,475786 | 1,466658 | 1,436229 | 1,530558 | 1,758772 | 1,606629 | 1,484914 | 1,545771 | 1,475786 | 1,466656 | 1,557942 | 1,527514 |
| | Retrosplenial area ventral part layer 6a | -2,18 | *0,5514* | 0,5790 | 0,0311 | 7 | 0,5664 | 0,0444 | 7 | 0,5538 | 0,593357 | 0,590314 | 0,587272 | 0,526414 | 0,578143 | 0,623786 | 0,639 | 0,5325 | 0,550757 | 0,505114 | 0,5751 | 0,559885 | 0,602486 |
| | Retrosplenial area ventral part layer 6b | -6,73 | *0,5449* | 0,0452 | 0,0103 | 7 | 0,0422 | 0,0077 | 7 | 0,054771 | 0,027386 | 0,039557 | 0,0426 | 0,054771 | 0,0426 | 0,054771 | 0,048686 | 0,045643 | 0,045643 | 0,036514 | 0,027386 | 0,0426 | 0,048686 |
| | Posterior parietal association areas | 2,75 | *0,7270* | 2,26 | 0,32 | 7 | 2,32 | 0,33 | 7 | 2,044799 | 1,923087 | 2,455587 | 2,172601 | 1,932214 | 2,71423 | 2,580343 | 2,349087 | 2,881586 | 2,321699 | 1,813543 | 2,361256 | 2,449499 | 2,081314 |
| | Anterior area | 3,84 | *0,7228* | 1,32 | 0,27 | 7 | 1,37 | 0,25 | 7 | 1,0863 | 1,068043 | 1,591415 | 1,250615 | 1,049786 | 1,640101 | 1,573157 | 1,372329 | 1,764857 | 1,433185 | 1,016315 | 1,512299 | 1,427099 | 1,089343 |
| | Anterior area layer 1 | 0,99 | *0,9190* | 0,2204 | 0,0440 | 7 | 0,2226 | 0,0335 | 7 | 0,179529 | 0,182572 | 0,2556 | 0,188657 | 0,1917 | 0,267772 | 0,2769 | 0,206914 | 0,267771 | 0,2343 | 0,182571 | 0,261686 | 0,216043 | 0,188657 |
| | Anterior area layer 2/3 | -0,22 | *0,9860* | 0,3951 | 0,0913 | 7 | 0,3943 | 0,0903 | 7 | 0,337757 | 0,279943 | 0,462514 | 0,368186 | 0,3195 | 0,520329 | 0,477729 | 0,386443 | 0,514243 | 0,422957 | 0,237343 | 0,4686 | 0,395571 | 0,334714 |
| | Anterior area layer 4 | 4,85 | *0,6766* | 0,1434 | 0,0297 | 7 | 0,1504 | 0,0312 | 7 | 0,118671 | 0,124757 | 0,182571 | 0,124757 | 0,115629 | 0,182572 | 0,155186 | 0,155186 | 0,200829 | 0,146057 | 0,1065 | 0,164314 | 0,161271 | 0,118671 |
| | Anterior area layer 5 | 7,23 | *0,4926* | 0,3308 | 0,0589 | 7 | 0,3547 | 0,0671 | 7 | 0,282986 | 0,273857 | 0,392529 | 0,325586 | 0,264729 | 0,4047 | 0,371229 | 0,371229 | 0,471643 | 0,356014 | 0,2769 | 0,377314 | 0,356014 | 0,273857 |
| | Anterior area layer 6a | 12,22 | *0,3610* | 0,1956 | 0,0497 | 7 | 0,2195 | 0,0443 | 7 | 0,136929 | 0,164314 | 0,2556 | 0,209957 | 0,136929 | 0,219086 | 0,246471 | 0,219086 | 0,273857 | 0,243428 | 0,182571 | 0,206914 | 0,261686 | 0,1491 |
| | Anterior area layer 6b | -13,95 | *0,2104* | 0,0374 | 0,0092 | 7 | 0,0322 | 0,0043 | 7 | 0,030429 | 0,0426 | 0,0426 | 0,033471 | 0,0213 | 0,045643 | 0,045643 | 0,033471 | 0,036514 | 0,030429 | 0,030429 | 0,033471 | 0,036514 | 0,024343 |
| | Rostrolateral visual area | 1,21 | *0,8309* | 0,9376 | 0,0810 | 7 | 0,9489 | 0,1102 | 7 | 0,9585 | 0,855043 | 0,864172 | 0,921986 | 0,882429 | 1,074129 | 1,007186 | 0,976758 | 1,116729 | 0,888514 | 0,797229 | 0,848957 | 1,022399 | 0,991971 |
| | Rostrolateral area layer 1 | -1,67 | *0,7821* | 0,1561 | 0,0158 | 7 | 0,1534 | 0,0186 | 7 | 0,182571 | 0,1491 | 0,152143 | 0,133886 | 0,152143 | 0,152143 | 0,1704 | 0,1491 | 0,161271 | 0,158229 | 0,130843 | 0,143014 | 0,188657 | 0,143014 |

| | | | | | | | | | | | | | | | | | | | | | | | |
|---|---|---|---|---|---|---|---|---|---|---|---|---|---|---|---|---|---|---|---|---|---|---|---|
| | **Rostrolateral area layer 2/3** | -1,47 | *0,7415* | 0,2656 | 0,0173 | 7 | 0,2617 | 0,0252 | 7 | 0,2769 | 0,249514 | 0,2343 | 0,270814 | 0,273857 | 0,282986 | 0,270814 | 0,2769 | 0,270814 | 0,264728 | 0,2343 | 0,219086 | 0,279943 | 0,286029 |
| | **Rostrolateral area layer 4** | -0,61 | *0,9241* | 0,1426 | 0,0190 | 7 | 0,1417 | 0,0139 | 7 | 0,152143 | 0,136929 | 0,103457 | 0,161272 | 0,143014 | 0,155186 | 0,146057 | 0,146057 | 0,164314 | 0,1278 | 0,130843 | 0,130843 | 0,136928 | 0,155186 |
| | **Rostrolateral area layer 5** | 2,99 | *0,7581* | 0,2182 | 0,0392 | 7 | 0,2247 | 0,0383 | 7 | 0,206914 | 0,179529 | 0,213 | 0,213 | 0,176486 | 0,286029 | 0,252557 | 0,243429 | 0,295157 | 0,194743 | 0,179529 | 0,203871 | 0,2343 | 0,222129 |
| | **Rostrolateral area layer 6a** | 3,30 | *0,7478* | 0,1317 | 0,0195 | 7 | 0,1361 | 0,0289 | 7 | 0,118671 | 0,118671 | 0,139971 | 0,115629 | 0,121714 | 0,1704 | 0,136929 | 0,1278 | 0,182571 | 0,109543 | 0,097371 | 0,130843 | 0,1491 | 0,155186 |
| * | **Rostrolateral area layer 6b** | 33,33 | *0,0361* | 0,0235 | 0,0052 | 7 | 0,0313 | 0,0070 | 7 | 0,0213 | 0,0213 | 0,0213 | 0,027386 | 0,015214 | 0,027386 | 0,030429 | 0,033471 | 0,0426 | 0,033471 | 0,024343 | 0,0213 | 0,033471 | 0,030429 |
| | **Temporal association areas** | 0,99 | *0,7654* | 2,76 | 0,19 | 7 | 2,79 | 0,15 | 7 | 2,823771 | 2,656416 | 2,699015 | 2,625987 | 2,543829 | 2,878544 | 3,094586 | 2,802473 | 2,723358 | 2,78117 | 2,711187 | 2,559042 | 2,963741 | 2,972871 |
| | **Temporal association areas layer 1** | -3,70 | *0,4081* | 0,4582 | 0,0480 | 7 | 0,4412 | 0,0179 | 7 | 0,459471 | 0,410786 | 0,474686 | 0,505115 | 0,435129 | 0,395572 | 0,526414 | 0,465557 | 0,432086 | 0,422957 | 0,416872 | 0,456428 | 0,444257 | 0,450343 |
| | **Temporal association areas layer 2/3** | -3,57 | *0,5308* | 0,6216 | 0,0671 | 7 | 0,5994 | 0,0613 | 7 | 0,642043 | 0,584229 | 0,550757 | 0,620743 | 0,626829 | 0,572057 | 0,754629 | 0,584229 | 0,544672 | 0,648128 | 0,620743 | 0,517285 | 0,584228 | 0,696814 |
| | **Temporal association areas layer 4** | -11,76 | *0,0552* | 0,2882 | 0,0312 | 7 | 0,2543 | 0,0285 | 7 | 0,243428 | 0,292114 | 0,249514 | 0,310372 | 0,286029 | 0,313414 | 0,322543 | 0,289072 | 0,243429 | 0,273857 | 0,279943 | 0,206914 | 0,246471 | 0,240386 |
| | **Temporal association areas layer 5** | 2,08 | *0,6755* | 0,8572 | 0,0891 | 7 | 0,8750 | 0,0638 | 7 | 0,867214 | 0,839829 | 0,876343 | 0,8094 | 0,7029 | 0,988929 | 0,9159 | 0,900686 | 0,818529 | 0,839828 | 0,824615 | 0,842871 | 0,998057 | 0,900686 |
| | **Temporal association areas layer 6a** | 11,68 | *0,1121* | 0,4690 | 0,0642 | 7 | 0,5238 | 0,0548 | 7 | 0,529457 | 0,456429 | 0,474686 | 0,3621 | 0,413829 | 0,508157 | 0,538586 | 0,474686 | 0,587272 | 0,508157 | 0,483814 | 0,459471 | 0,5751 | 0,578143 |
| * | **Temporal association areas layer 6b** | 42,11 | *0,0457* | 0,0661 | 0,0285 | 7 | 0,0939 | 0,0135 | 7 | 0,082157 | 0,073029 | 0,073029 | 0,018257 | 0,079114 | 0,100414 | 0,036514 | 0,088243 | 0,097371 | 0,088243 | 0,0852 | 0,076071 | 0,115629 | 0,1065 |
| | **Perirhinal area** | -0,83 | *0,9063* | 0,6290 | 0,0717 | 7 | 0,6238 | 0,0895 | 7 | 0,520328 | 0,584229 | 0,590314 | 0,699858 | 0,693771 | 0,608572 | 0,705943 | 0,5112 | 0,727243 | 0,5325 | 0,593357 | 0,587271 | 0,699857 | 0,715071 |
| | **Perirhinal area layer 1** | -6,21 | *0,4454* | 0,1821 | 0,0249 | 7 | 0,1708 | 0,0286 | 7 | 0,139971 | 0,182572 | 0,161271 | 0,185614 | 0,216043 | 0,1917 | 0,197786 | 0,124757 | 0,176486 | 0,155186 | 0,167357 | 0,161271 | 0,206914 | 0,203871 |
| | **Perirhinal area layer 2/3** | -0,17 | *0,9840* | 0,2560 | 0,0453 | 7 | 0,2556 | 0,0330 | 7 | 0,179529 | 0,225172 | 0,2556 | 0,286029 | 0,304286 | 0,240386 | 0,301243 | 0,228214 | 0,292114 | 0,213 | 0,243429 | 0,261686 | 0,246471 | 0,304286 |
| | **Perirhinal area layer 5** | 3,64 | *0,7466* | 0,1313 | 0,0225 | 7 | 0,1361 | 0,0308 | 7 | 0,133886 | 0,118671 | 0,112586 | 0,1704 | 0,109543 | 0,121714 | 0,152143 | 0,115629 | 0,179529 | 0,109543 | 0,1278 | 0,109543 | 0,179528 | 0,130843 |
| | **Perirhinal area layer 6a** | -5,61 | *0,5712* | 0,0465 | 0,0052 | 7 | 0,0439 | 0,0105 | 7 | 0,051729 | 0,045643 | 0,048686 | 0,045643 | 0,051729 | 0,045643 | 0,036514 | 0,033471 | 0,057814 | 0,036514 | 0,033471 | 0,0426 | 0,045643 | 0,057814 |
| | **Perirhinal area layer 6b** | 33,33 | *0,0708* | 0,0130 | 0,0029 | 7 | 0,0174 | 0,0049 | 7 | 0,015214 | 0,012171 | 0,012171 | 0,012171 | 0,012171 | 0,009129 | 0,018257 | 0,009129 | 0,0213 | 0,018257 | 0,0213 | 0,012171 | 0,0213 | 0,018257 |
| | **Ectorhinal area** | -2,83 | *0,5956* | 1,45 | 0,13 | 7 | 1,40 | 0,15 | 7 | 1,3419 | 1,314515 | 1,433186 | 1,430144 | 1,466657 | 1,408843 | 1,722257 | 1,238444 | 1,536643 | 1,278 | 1,281043 | 1,478828 | 1,396671 | 1,621843 |
| | **Ectorhinal area/Layer 1** | -0,78 | *0,9029* | 0,2217 | 0,0182 | 7 | 0,2200 | 0,0319 | 7 | 0,225171 | 0,200829 | 0,222129 | 0,209957 | 0,243429 | 0,203872 | 0,246471 | 0,173443 | 0,228214 | 0,209957 | 0,194743 | 0,219086 | 0,243428 | 0,270814 |
| | **Ectorhinal area/Layer 2/3** | -8,06 | *0,4235* | 0,3938 | 0,0810 | 7 | 0,3621 | 0,0603 | 7 | 0,322543 | 0,301243 | 0,380357 | 0,389486 | 0,456429 | 0,368186 | 0,538586 | 0,301243 | 0,407743 | 0,3408 | 0,310372 | 0,416871 | 0,310371 | 0,4473 |
| | **Ectorhinal area/Layer 5** | -4,80 | *0,3017* | 0,4525 | 0,0364 | 7 | 0,4308 | 0,0389 | 7 | 0,438171 | 0,4473 | 0,422957 | 0,444257 | 0,419914 | 0,4686 | 0,526414 | 0,429043 | 0,453386 | 0,365143 | 0,410786 | 0,486857 | 0,416871 | 0,453386 |
| | **Ectorhinal area/Layer 6a** | 0,26 | *0,9587* | 0,3356 | 0,0223 | 7 | 0,3365 | 0,0371 | 7 | 0,307328 | 0,328629 | 0,371229 | 0,349929 | 0,316457 | 0,325586 | 0,349929 | 0,295157 | 0,380357 | 0,313414 | 0,316457 | 0,304286 | 0,368186 | 0,377314 |
| * | **Ectorhinal area/Layer 6b** | 32,29 | *0,0397* | 0,0417 | 0,0102 | 7 | 0,0552 | 0,0116 | 7 | 0,048686 | 0,036514 | 0,036514 | 0,036514 | 0,030429 | 0,0426 | 0,060857 | 0,039557 | 0,066943 | 0,048686 | 0,048686 | 0,051729 | 0,057814 | 0,073029 |
| | **Olfactory areas** | -2,22 | *0,3454* | 43,78 | 1,89 | 7 | 42,81 | 1,81 | 7 | 43,82017 | 41,44069 | 41,42547 | 43,58591 | 44,76652 | 44,77262 | 46,67743 | 42,51787 | 43,53416 | 40,20221 | 42,93777 | 42,71256 | 41,6719 | 46,1145 |
| | **Main olfactory bulb** | -4,02 | *0,1309* | 15,47 | 0,73 | 7 | 14,85 | 0,70 | 7 | 15,7833 | 14,02758 | 15,00738 | 15,55814 | 15,81981 | 16,03586 | 16,0815 | 15,21125 | 14,89479 | 13,97584 | 14,82481 | 15,55508 | 13,87238 | 15,62811 |
| | **Accessory olfactory bulb** | -14,69 | *0,1530* | 0,6538 | 0,0666 | 7 | 0,5577 | 0,1473 | 7 | 0,639 | 0,678558 | 0,635957 | 0,739415 | 0,730286 | 0,562929 | 0,590314 | 0,556843 | 0,4686 | 0,365143 | 0,678557 | 0,590314 | 0,4473 | 0,797229 |
| | **Accessory olfactory bulb glomerular layer** | -42,82 | *0,1626* | 0,2373 | 0,0317 | 5 | 0,1357 | 0,1322 | 5 | 0,209957 | 0,240386 | 0,200829 | 0,261686 | 0,273857 | | | | 0,039557 | | 0,249514 | 0,054771 | 0,027386 | 0,307329 |
| | **Accessory olfactory bulb granular layer** | -12,32 | *0,2563* | 0,2117 | 0,0451 | 7 | 0,1856 | 0,0361 | 7 | 0,182571 | 0,246472 | 0,237343 | 0,273857 | 0,167357 | 0,152143 | 0,222129 | 0,143014 | 0,185614 | 0,176486 | 0,194743 | 0,155186 | 0,188657 | 0,2556 |
| | **Accessory olfactory bulb mitral layer** | 0,96 | *0,9560* | 0,2726 | 0,0876 | 7 | 0,2752 | 0,0857 | 7 | 0,246471 | 0,1917 | 0,197786 | 0,203872 | 0,289071 | 0,410786 | 0,368186 | 0,413829 | 0,243429 | 0,188657 | 0,2343 | 0,380357 | 0,231257 | 0,2343 |
| * | **Anterior olfactory nucleus** | -7,13 | *0,0173* | 4,91 | 0,15 | 7 | 4,56 | 0,28 | 7 | 4,886827 | 4,783374 | 4,69513 | 5,017674 | 4,902043 | 4,944645 | 5,157643 | 4,603845 | 4,570372 | 4,305641 | 4,832059 | 4,94464 | 4,126112 | 4,552114 |
| | **Taenia tecta** | -8,71 | *0,2189* | 1,45 | 0,16 | 7 | 1,32 | 0,20 | 7 | 1,305385 | 1,518387 | 1,399715 | 1,594458 | 1,700957 | 1,311472 | 1,302343 | 1,204972 | 1,3845 | 1,2354 | 1,159329 | 1,183671 | 1,347985 | 1,734429 |
| * | **Taenia tecta dorsal part** | -20,01 | *0,0215* | 0,7646 | 0,1123 | 7 | 0,6116 | 0,1042 | 7 | 0,657257 | 0,8307 | 0,800272 | 0,879386 | 0,891557 | 0,672472 | 0,620743 | 0,562929 | 0,587272 | 0,5538 | 0,587272 | 0,581185 | 0,562928 | 0,845914 |
| | **Taenia tecta ventral part** | 3,95 | *0,6076* | 0,6829 | 0,0673 | 7 | 0,7099 | 0,1163 | 7 | 0,648128 | 0,687686 | 0,599443 | 0,715072 | 0,8094 | 0,639 | 0,6816 | 0,642043 | 0,797229 | 0,6816 | 0,572057 | 0,602485 | 0,785057 | 0,888514 |
| | **Dorsal peduncular area** | -8,41 | *0,1488* | 0,5425 | 0,0372 | 7 | 0,4969 | 0,0671 | 7 | 0,605528 | 0,526415 | 0,514243 | 0,523372 | 0,559886 | 0,569015 | 0,499029 | 0,517286 | 0,486857 | 0,392528 | 0,584229 | 0,535543 | 0,535543 | 0,426 |
| | **Piriform area** | 0,61 | *0,8252* | 10,61 | 0,61 | 7 | 10,68 | 0,44 | 7 | 10,6287 | 10,37615 | 9,867989 | 10,19358 | 10,44917 | 11,05166 | 11,72109 | 10,37311 | 10,98472 | 10,26051 | 10,74738 | 10,19052 | 10,76258 | 11,41984 |
| | **Nucleus of the lateral olfactory tract** | 4,19 | *0,7672* | 0,3634 | 0,1036 | 7 | 0,3786 | 0,0831 | 7 | 0,492943 | 0,462515 | 0,429043 | 0,243429 | 0,307329 | 0,371229 | 0,237343 | 0,334714 | 0,459471 | 0,4047 | 0,459472 | 0,243428 | 0,435128 | 0,313414 |
| | **Nucleus of the lateral olfactory tract molecular layer** | -2,37 | *0,8960* | 0,1282 | 0,0394 | 7 | 0,1252 | 0,0457 | 7 | 0,164314 | 0,158229 | 0,158229 | 0,066943 | 0,1491 | 0,0852 | 0,115629 | 0,066943 | 0,167357 | 0,118671 | 0,158229 | 0,0852 | 0,185614 | 0,094329 |
| | **Nucleus of the lateral olfactory tract pyramidal layer** | 5,29 | *0,7810* | 0,1808 | 0,0768 | 7 | 0,1904 | 0,0440 | 7 | 0,270814 | 0,246472 | 0,219086 | 0,136929 | 0,100414 | 0,219086 | 0,073029 | 0,203872 | 0,225171 | 0,222128 | 0,231257 | 0,112586 | 0,185614 | 0,152143 |
| | **Nucleus of the lateral olfactory tract layer 3** | 16,00 | *0,0753* | 0,0543 | 0,0087 | 7 | 0,0630 | 0,0080 | 7 | 0,057814 | 0,057814 | 0,051729 | 0,039557 | 0,057814 | 0,066943 | 0,048686 | 0,0639 | 0,066943 | 0,0639 | 0,069986 | 0,045643 | 0,0639 | 0,066943 |
| | **Cortical amygdalar area** | -3,66 | *0,3020* | 3,19 | 0,22 | 7 | 3,07 | 0,18 | 7 | 3,146313 | 2,878544 | 2,99113 | 3,155445 | 3,347143 | 3,274116 | 3,547972 | 3,155445 | 3,0033 | 2,863328 | 3,027644 | 3,094584 | 2,957655 | 3,420171 |
| | **Cortical amygdalar area anterior part** | -3,97 | *0,5547* | 0,8537 | 0,0974 | 7 | 0,8198 | 0,1108 | 7 | 0,821571 | 0,7668 | 0,7029 | 0,903729 | 0,888514 | 0,897643 | 0,995014 | 0,879386 | 0,678557 | 0,815485 | 0,785057 | 0,921985 | 0,690728 | 0,967629 |
| | **Cortical amygdalar area posterior part** | -3,55 | *0,2718* | 2,34 | 0,14 | 7 | 2,25 | 0,13 | 7 | 2,324742 | 2,111744 | 2,288229 | 2,251716 | 2,458629 | 2,376472 | 2,552957 | 2,276058 | 2,324743 | 2,047842 | 2,242586 | 2,172599 | 2,266927 | 2,452543 |
| | **Cortical amygdalar area posterior part lateral zone** | -2,19 | *0,5584* | 1,05 | 0,07 | 7 | 1,03 | 0,07 | 7 | 1,016314 | 0,925029 | 1,0437 | 1,061958 | 1,095429 | 1,089343 | 1,147157 | 1,065001 | 1,040657 | 0,946328 | 0,995015 | 1,010228 | 0,988928 | 1,1715 |
| | **Cortical amygdalar area posterior part medial zone** | -4,67 | *0,1708* | 1,28 | 0,08 | 7 | 1,22 | 0,07 | 7 | 1,308428 | 1,186715 | 1,244529 | 1,189758 | 1,3632 | 1,287129 | 1,4058 | 1,211058 | 1,284086 | 1,101514 | 1,247572 | 1,162371 | 1,277999 | 1,281043 |
| | **Piriform-amygdalar area** | -1,81 | *0,6217* | 1,10 | 0,07 | 7 | 1,08 | 0,08 | 7 | 1,159328 | 1,168458 | 1,046743 | 1,019358 | 1,0437 | 1,168458 | 1,119771 | 1,074129 | 1,098472 | 1,034571 | 1,189758 | 1,010228 | 0,988928 | 1,189757 |
| | **Postpiriform transition area** | -6,70 | *0,1786* | 1,15 | 0,11 | 7 | 1,07 | 0,09 | 7 | 1,186714 | 1,058915 | 1,141072 | 1,019358 | 1,180629 | 1,095429 | 1,360157 | 0,940243 | 1,1928 | 0,988928 | 1,116729 | 1,089342 | 1,134985 | 1,040657 |
| | **Hippocampal formation** | -2,73 | *0,1721* | 40,50 | 1,64 | 7 | 39,39 | 1,14 | 7 | 39,4628 | 39,89188 | 38,89381 | 41,23987 | 39,0216 | 41,62935 | 43,33942 | 39,57846 | 39,96185 | 38,45866 | 38,65951 | 38,26695 | 39,23458 | 41,58369 |
| | **Hippocampal region** | -0,52 | *0,8378* | 22,62 | 0,92 | 7 | 22,50 | 1,15 | 7 | 21,87509 | 22,42587 | 21,48258 | 22,65104 | 22,23416 | 24,08118 | 23,58214 | 22,02725 | 23,856 | 20,84052 | 21,98161 | 21,73816 | 23,3174 | 23,75559 |
| | **Ammon's horn** | -2,37 | *0,2892* | 15,95 | 0,55 | 7 | 15,57 | 0,71 | 7 | 15,3786 | 15,72245 | 15,35731 | 15,96588 | 15,87459 | 16,74485 | 16,60792 | 15,19908 | 16,44665 | 14,57224 | 15,24472 | 15,26905 | 15,72852 | 16,54097 |
| | **Field CA1** | -2,02 | *0,4353* | 9,42 | 0,36 | 7 | 9,23 | 0,50 | 7 | 9,107269 | 9,238119 | 9,372003 | 9,396348 | 9,019029 | 10,02926 | 9,776701 | 8,882105 | 9,791916 | 8,666054 | 8,958174 | 9,073796 | 9,192466 | 10,04447 |
| | **Field CA2** | -17,48 | *0,0917* | 0,4551 | 0,0901 | 7 | 0,3756 | 0,0702 | 7 | 0,593357 | 0,520329 | 0,334714 | 0,495986 | 0,441214 | 0,438172 | 0,3621 | 0,356014 | 0,313414 | 0,371228 | 0,523372 | 0,343843 | 0,328628 | 0,392529 |
| | **Field CA3** | -1,80 | *0,5171* | 6,08 | 0,33 | 7 | 5,97 | 0,28 | 7 | 5,67797 | 5,964003 | 5,650588 | 6,073546 | 6,414343 | 6,277417 | 6,469115 | 5,96096 | 6,341315 | 5,534955 | 5,763173 | 5,851411 | 6,207425 | 6,103972 |
| | **Dentate gyrus** | 3,77 | *0,3084* | 6,50 | 0,38 | 7 | 6,75 | 0,48 | 7 | 6,313927 | 6,554318 | 5,985302 | 6,539104 | 6,222643 | 7,126374 | 6,779486 | 6,660818 | 7,226787 | 6,103969 | 6,563445 | 6,289583 | 7,38501 | 7,0077 |
| | **Dentate gyrus molecular layer** | 10,34 | *0,0582* | 4,20 | 0,38 | 7 | 4,64 | 0,39 | 7 | 4,25087 | 4,390845 | 3,879644 | 4,050045 | 3,767057 | 4,923345 | 4,159586 | 4,634274 | 4,987244 | 4,05917 | 4,643402 | 4,177841 | 5,102869 | 4,859443 |
| * | **Dentate gyrus polymorph layer** | -27,78 | *0,0441* | 0,7307 | 0,2104 | 7 | 0,5277 | 0,0647 | 7 | 0,559886 | 0,620743 | 0,6603 | 0,946329 | 0,912857 | 0,444257 | 0,970671 | 0,569015 | 0,5325 | 0,602486 | 0,492943 | 0,593357 | 0,432085 | 0,471643 |
| | **Dentate gyrus granule cell layer** | 0,86 | *0,8571* | 1,57 | 0,10 | 7 | 1,58 | 0,16 | 7 | 1,503171 | 1,542729 | 1,445358 | 1,542729 | 1,542729 | 1,758772 | 1,649229 | 1,457529 | 1,707043 | 1,442314 | 1,4271 | 1,518385 | 1,850056 | 1,676614 |
| | **Fasciola cinerea** | 17,52 | *0,2937* | 0,0596 | 0,0216 | 7 | 0,0700 | 0,0123 | 7 | 0,066943 | 0,057814 | 0,0426 | 0,030429 | 0,048686 | 0,094329 | 0,076071 | 0,0639 | 0,0852 | 0,079114 | 0,0639 | 0,0639 | 0,082157 | 0,051729 |
| | **Induseum griseum** | 6,15 | *0,5209* | 0,1061 | 0,0132 | 7 | 0,1126 | 0,0223 | 7 | 0,115629 | 0,091286 | 0,097371 | 0,115629 | 0,088243 | 0,115629 | 0,118671 | 0,103457 | 0,097371 | 0,0852 | 0,109543 | 0,115629 | 0,121714 | 0,155186 |
| * | **Retrohippocampal region** | -6,23 | *0,0392* | 17,46 | 0,95 | 7 | 16,37 | 0,79 | 7 | 17,13128 | 16,97002 | 17,15259 | 18,33323 | 16,49837 | 16,93959 | 19,20652 | 17,0674 | 15,69202 | 17,17084 | 16,12411 | 15,99933 | 15,29643 | 17,27126 |
| | **Entorhinal area** | -1,75 | *0,4928* | 10,34 | 0,52 | 7 | 10,16 | 0,43 | 7 | 10,01708 | 10,04752 | 9,892332 | 10,49482 | 9,895372 | 10,85996 | 11,2038 | 10,45222 | 9,746273 | 10,36093 | 10,04447 | 10,03838 | 9,624552 | 10,87517 |
| | **Entorhinal area lateral part** | -4,76 | *0,1076* | 5,63 | 0,29 | 7 | 5,36 | 0,29 | 7 | 5,504527 | 5,455846 | 5,489316 | 5,681018 | 5,331086 | 5,757088 | 6,204386 | 5,334131 | 5,312829 | 5,413241 | 5,315873 | 5,352383 | 4,917254 | 5,9001 |
| * | **Entorhinal area lateral part layer 1** | -8,67 | *0,0264* | 0,8424 | 0,0608 | 7 | 0,7694 | 0,0447 | 7 | 0,794185 | 0,821572 | 0,8094 | 0,888515 | 0,791143 | 0,833743 | 0,9585 | 0,708986 | 0,782014 | 0,742457 | 0,772886 | 0,760714 | 0,763757 | 0,855043 |
| | **Entorhinal area lateral part layer 2** | -4,34 | *0,3490* | 1,37 | 0,09 | 7 | 1,31 | 0,14 | 7 | 1,296257 | 1,278001 | 1,369286 | 1,387544 | 1,354071 | 1,384501 | 1,542729 | 1,424058 | 1,293214 | 1,421014 | 1,259743 | 1,308428 | 1,052828 | 1,436229 |
| * | **Entorhinal area lateral part layer 3** | -5,46 | *0,0405* | 1,15 | 0,05 | 7 | 1,09 | 0,05 | 7 | 1,165414 | 1,144115 | 1,098472 | 1,119772 | 1,134986 | 1,174543 | 1,244529 | 1,083258 | 1,092386 | 1,071085 | 1,068043 | 1,107599 | 1,019357 | 1,198886 |
| | **Entorhinal area lateral part layer 5** | -5,55 | *0,1242* | 1,28 | 0,09 | 7 | 1,21 | 0,06 | 7 | 1,238442 | 1,238444 | 1,229315 | 1,329729 | 1,147157 | 1,320601 | 1,433186 | 1,189758 | 1,208014 | 1,168457 | 1,201929 | 1,165414 | 1,162371 | 1,344943 |
| | **Entorhinal area lateral part layer 6a** | -0,18 | *0,9506* | 0,9850 | 0,0471 | 7 | 0,9833 | 0,0553 | 7 | 1,010228 | 0,973715 | 0,982843 | 0,955458 | 0,903729 | 1,0437 | 1,025443 | 0,928072 | 0,9372 | 1,010228 | 1,013272 | 1,010228 | 0,918942 | 1,065 |
| | **Entorhinal area medial part dorsal zone** | 1,84 | *0,5229* | 4,71 | 0,26 | 7 | 4,80 | 0,23 | 7 | 4,512556 | 4,591674 | 4,403016 | 4,813803 | 4,564286 | 5,102874 | 4,999415 | 5,118089 | 4,433443 | 4,947684 | 4,728602 | 4,685998 | 4,707297 | 4,975072 |
| | **Entorhinal area medial part dorsal zone layer 1** | -4,57 | *0,4400* | 1,19 | 0,14 | 7 | 1,13 | 0,11 | 7 | 1,168457 | 1,195844 | 1,065 | 1,357115 | 0,961543 | 1,229315 | 1,347986 | 1,183672 | 0,967629 | 1,265828 | 1,2141 | 1,113685 | 1,016314 | 1,183671 |

| | | | | | | | | | | | | | | | | | | | | | | | |
|---|---|---|---|---|---|---|---|---|---|---|---|---|---|---|---|---|---|---|---|---|---|---|---|
| | Entorhinal area medial part dorsal zone layer 2 | 9,47 | *0,1755* | 1,18 | 0,18 | 7 | 1,29 | 0,10 | 7 | 1,296257 | 1,296258 | 1,141072 | 0,906772 | 1,092386 | 1,439272 | 1,0863 | 1,454487 | 1,165414 | 1,375371 | 1,268872 | 1,214099 | 1,302342 | 1,259743 |
| | Entorhinal area medial part dorsal zone layer 3 | 10,45 | *0,3427* | 0,8446 | 0,1814 | 7 | 0,9329 | 0,1510 | 7 | 0,635957 | 0,687686 | 0,775929 | 0,824615 | 1,065 | 1,116729 | 0,806357 | 0,897643 | 1,055872 | 0,751585 | 0,754629 | 0,885471 | 1,128899 | 1,055871 |
| | Entorhinal area medial part dorsal zone layer 5 | -12,43 | *0,1192* | 0,9129 | 0,1497 | 7 | 0,7994 | 0,0929 | 7 | 0,797228 | 0,785058 | 1,0437 | 1,049786 | 0,964586 | 0,696815 | 1,052829 | 0,867215 | 0,797229 | 0,879385 | 0,8094 | 0,842871 | 0,602485 | 0,797229 |
| | Entorhinal area medial part dorsal zone layer 6 | 9,35 | *0,3426* | 0,5860 | 0,1160 | 7 | 0,6407 | 0,0891 | 7 | 0,614657 | 0,626829 | 0,377314 | 0,675515 | 0,480771 | 0,620743 | 0,705943 | 0,715072 | 0,4473 | 0,675514 | 0,6816 | 0,629871 | 0,657257 | 0,678557 |
| | Parasubiculum | -1,26 | *0,8908* | 0,8637 | 0,1954 | 7 | 0,8529 | 0,0503 | 7 | 0,748543 | 0,754629 | 0,572057 | 1,052829 | 1,034571 | 0,797229 | 1,0863 | 0,855043 | 0,864172 | 0,8307 | 0,815486 | 0,812442 | 0,958499 | 0,833743 |
| | Postsubiculum | -5,74 | *0,4888* | 1,03 | 0,17 | 7 | 0,9702 | 0,1379 | 7 | 1,1715 | 1,098472 | 0,897643 | 1,061958 | 0,842871 | 0,852 | 1,281043 | 1,153243 | 0,7455 | 1,065 | 1,040657 | 0,912857 | 1,010228 | 0,864171 |
| * | Presubiculum | -28,60 | *0,0394* | 1,12 | 0,28 | 7 | 0,7977 | 0,2316 | 7 | 1,177585 | 1,046743 | 1,618801 | 1,235401 | 0,751586 | 0,861129 | 1,1289 | 0,928072 | 0,599443 | 1,125857 | 0,663343 | 0,836785 | 0,465557 | 0,964586 |
| * | Subiculum | -23,48 | *0,0090* | 2,07 | 0,35 | 7 | 1,58 | 0,13 | 7 | 1,828757 | 1,807458 | 2,330829 | 2,537744 | 2,181729 | 1,554901 | 2,233457 | 1,554901 | 1,834843 | 1,664442 | 1,484915 | 1,515342 | 1,469699 | 1,551857 |
| | Prosubiculum | -3,17 | *0,7922* | 1,37 | 0,34 | 7 | 1,33 | 0,26 | 7 | 1,685742 | 1,685744 | 1,0863 | 1,183672 | 0,921986 | 1,271915 | 1,764857 | 1,567072 | 0,909814 | 1,615757 | 1,533601 | 1,299299 | 1,110642 | 1,259743 |
| | Hippocampo-amygdalar transition area | -0,80 | *0,8112* | 0,4364 | 0,0297 | 7 | 0,4330 | 0,0232 | 7 | 0,413828 | 0,459472 | 0,419914 | 0,444257 | 0,416871 | 0,410786 | 0,4899 | 0,441215 | 0,477729 | 0,432086 | 0,432086 | 0,407743 | 0,410785 | 0,429043 |
| | Area prostriata | 6,95 | *0,8673* | 0,2313 | 0,1685 | 7 | 0,2473 | 0,1836 | 7 | 0,088243 | 0,069986 | 0,334714 | 0,322543 | 0,453386 | 0,331672 | 0,018257 | 0,115629 | 0,514243 | 0,076071 | 0,109543 | 0,176486 | 0,246471 | 0,492943 |
| | Cortical subplate | -1,76 | *0,4855* | 8,34 | 0,46 | 7 | 8,19 | 0,27 | 7 | 8,477397 | 8,097047 | 7,832317 | 8,094005 | 7,984457 | 8,775604 | 9,098143 | 7,99359 | 8,240058 | 7,880997 | 8,32526 | 8,142682 | 8,048353 | 8,699529 |
| | Claustrum | -2,14 | *0,7946* | 0,5473 | 0,0909 | 7 | 0,5355 | 0,0729 | 7 | 0,593357 | 0,550757 | 0,4686 | 0,480772 | 0,435129 | 0,614657 | 0,687686 | 0,599443 | 0,453386 | 0,642043 | 0,535543 | 0,569014 | 0,499028 | 0,450343 |
| | Endopiriform nucleus | -0,03 | *0,9901* | 2,50 | 0,11 | 7 | 2,50 | 0,15 | 7 | 2,534699 | 2,446458 | 2,379515 | 2,449501 | 2,455586 | 2,540787 | 2,7051 | 2,358216 | 2,522529 | 2,400813 | 2,498187 | 2,467756 | 2,446456 | 2,8116 |
| | Endopiriform nucleus dorsal part | 2,69 | *0,4157* | 1,60 | 0,06 | 7 | 1,65 | 0,12 | 7 | 1,567071 | 1,557944 | 1,545772 | 1,597501 | 1,588371 | 1,634015 | 1,728343 | 1,524472 | 1,676615 | 1,536642 | 1,594458 | 1,618799 | 1,700956 | 1,868314 |
| | Endopiriform nucleus ventral part | -4,88 | *0,1867* | 0,8989 | 0,0554 | 7 | 0,8550 | 0,0617 | 7 | 0,967628 | 0,888515 | 0,833743 | 0,852001 | 0,867214 | 0,906772 | 0,976757 | 0,833743 | 0,845914 | 0,864171 | 0,903729 | 0,848957 | 0,7455 | 0,943286 |
| | Lateral amygdalar nucleus | 1,07 | *0,8532* | 0,7694 | 0,0870 | 7 | 0,7777 | 0,0760 | 7 | 0,8307 | 0,827658 | 0,769843 | 0,715072 | 0,605529 | 0,861129 | 0,775929 | 0,842872 | 0,721157 | 0,760714 | 0,815486 | 0,803314 | 0,642042 | 0,858086 |
| | Basolateral amygdalar nucleus | -3,05 | *0,4184* | 1,69 | 0,15 | 7 | 1,64 | 0,06 | 7 | 1,634014 | 1,524472 | 1,545772 | 1,710087 | 1,710086 | 1,783115 | 1,953514 | 1,606629 | 1,6614 | 1,560985 | 1,691829 | 1,606628 | 1,618799 | 1,752686 |
| | Basolateral amygdalar nucleus anterior part | -3,49 | *0,4639* | 0,6977 | 0,0722 | 7 | 0,6733 | 0,0441 | 7 | 0,657257 | 0,648129 | 0,608572 | 0,687686 | 0,712029 | 0,742457 | 0,827657 | 0,6177 | 0,678557 | 0,632914 | 0,696815 | 0,687685 | 0,651171 | 0,748543 |
| | Basolateral amygdalar nucleus posterior part | 1,26 | *0,6845* | 0,6225 | 0,0371 | 7 | 0,6303 | 0,0330 | 7 | 0,645086 | 0,578143 | 0,648129 | 0,605529 | 0,5751 | 0,672472 | 0,632914 | 0,626829 | 0,672472 | 0,602486 | 0,6603 | 0,578143 | 0,623785 | 0,648129 |
| | Basolateral amygdalar nucleus ventral part | -9,41 | *0,2641* | 0,3743 | 0,0743 | 7 | 0,3391 | 0,0176 | 7 | 0,331671 | 0,2982 | 0,289072 | 0,416872 | 0,422957 | 0,368186 | 0,492943 | 0,3621 | 0,310371 | 0,325586 | 0,334714 | 0,3408 | 0,343843 | 0,356014 |
| | Basomedial amygdalar nucleus | -3,47 | *0,3468* | 1,45 | 0,10 | 7 | 1,40 | 0,10 | 7 | 1,442314 | 1,381458 | 1,302343 | 1,469701 | 1,481871 | 1,475786 | 1,615757 | 1,293215 | 1,424057 | 1,253657 | 1,430143 | 1,411885 | 1,490999 | 1,5123 |
| | Basomedial amygdalar nucleus anterior part | -6,76 | *0,3401* | 0,7529 | 0,1123 | 7 | 0,7020 | 0,0743 | 7 | 0,675514 | 0,6603 | 0,629872 | 0,815486 | 0,855043 | 0,708986 | 0,925029 | 0,6177 | 0,654214 | 0,614657 | 0,718115 | 0,751585 | 0,806357 | 0,751586 |
| | Basomedial amygdalar nucleus posterior part | 0,06 | *0,9878* | 0,6999 | 0,0543 | 7 | 0,7003 | 0,0497 | 7 | 0,7668 | 0,721158 | 0,672472 | 0,654215 | 0,626829 | 0,7668 | 0,690729 | 0,675515 | 0,769843 | 0,639 | 0,712029 | 0,6603 | 0,684642 | 0,760714 |
| | Posterior amygdalar nucleus | -1,11 | *0,8227* | 1,02 | 0,11 | 7 | 1,01 | 0,07 | 7 | 1,138028 | 1,058915 | 1,080215 | 0,873301 | 0,909814 | 1,134986 | 0,946329 | 0,982843 | 1,131943 | 0,967628 | 1,074129 | 0,921985 | 1,004142 | 0,9798 |
| | Cerebral nuclei | -2,34 | *0,2942* | 54,15 | 2,33 | 7 | 52,89 | 1,97 | 7 | 53,68207 | 54,11724 | 50,49623 | 53,55127 | 53,71556 | 55,19137 | 58,31028 | 52,28849 | 52,75098 | 50,34101 | 52,87879 | 52,14542 | 52,93656 | 56,86492 |
| | Striatum | -3,11 | *0,1465* | 44,78 | 1,79 | 7 | 43,39 | 1,55 | 7 | 44,07577 | 44,9126 | 41,94276 | 44,41966 | 44,62046 | 45,56681 | 47,91283 | 43,03211 | 42,94081 | 41,39806 | 43,50374 | 43,12639 | 43,15986 | 46,55572 |
| | Striatum dorsal region | -3,80 | *0,0697* | 24,86 | 0,95 | 7 | 23,91 | 0,82 | 7 | 24,52846 | 25,27094 | 23,42696 | 24,65324 | 24,35503 | 25,31354 | 26,4546 | 23,99294 | 23,22613 | 23,18048 | 24,40068 | 23,83469 | 23,28697 | 25,46263 |
| | Caudoputamen | -3,80 | *0,0697* | 24,86 | 0,95 | 7 | 23,91 | 0,82 | 7 | 24,52846 | 25,27094 | 23,42696 | 24,65324 | 24,35503 | 25,31354 | 26,4546 | 23,99294 | 23,22613 | 23,18048 | 24,40068 | 23,83469 | 23,28697 | 25,46263 |
| | Striatum ventral region | -2,33 | *0,4159* | 8,84 | 0,41 | 7 | 8,64 | 0,50 | 7 | 8,641712 | 8,574776 | 8,15486 | 8,964262 | 9,238115 | 8,976432 | 9,341572 | 8,46219 | 8,410458 | 7,99054 | 8,398289 | 8,629539 | 9,055538 | 9,502843 |
| | Nucleus accumbens | -3,71 | *0,1747* | 4,56 | 0,15 | 7 | 4,39 | 0,26 | 7 | 4,606884 | 4,585588 | 4,296516 | 4,527774 | 4,454743 | 4,658616 | 4,777286 | 4,527774 | 4,168715 | 4,150456 | 4,393887 | 4,348241 | 4,229569 | 4,905086 |
| | Fundus of striatum | 19,52 | *0,4210* | 0,3073 | 0,1151 | 7 | 0,3673 | 0,1512 | 7 | 0,295157 | 0,225172 | 0,209957 | 0,3408 | 0,547714 | 0,286029 | 0,246471 | 0,179529 | 0,514243 | 0,225171 | 0,3195 | 0,316457 | 0,5964 | 0,419914 |
| | Olfactory tubercle | -2,44 | *0,4892* | 3,98 | 0,26 | 7 | 3,88 | 0,25 | 7 | 3,73967 | 3,764016 | 3,648387 | 4,095688 | 4,235657 | 4,031787 | 4,317815 | 3,754888 | 3,727501 | 3,614913 | 3,684901 | 3,964841 | 4,229569 | 4,177843 |
| | Lateral septal complex | -1,53 | *0,5735* | 3,98 | 0,22 | 7 | 3,92 | 0,17 | 7 | 3,879642 | 4,101774 | 3,73663 | 3,803574 | 3,834 | 4,256959 | 4,263043 | 3,834002 | 4,077429 | 3,706199 | 3,742716 | 4,010484 | 3,931369 | 4,147414 |
| | Lateral septal nucleus | -1,81 | *0,4757* | 3,39 | 0,14 | 7 | 3,33 | 0,17 | 7 | 3,252813 | 3,441473 | 3,261944 | 3,301502 | 3,334972 | 3,578402 | 3,572315 | 3,298459 | 3,441472 | 3,140228 | 3,155444 | 3,362355 | 3,286284 | 3,630129 |
| | Lateral septal nucleus caudal (caudodorsal) part | -3,02 | *0,4764* | 0,6899 | 0,0461 | 7 | 0,6690 | 0,0591 | 7 | 0,602486 | 0,715072 | 0,7029 | 0,699858 | 0,669429 | 0,751586 | 0,687686 | 0,614657 | 0,751586 | 0,602486 | 0,611614 | 0,684643 | 0,708985 | 0,708986 |
| | Lateral septal nucleus rostral (rostroventral) part | 0,91 | *0,7516* | 2,06 | 0,10 | 7 | 2,08 | 0,11 | 7 | 2,026542 | 2,105658 | 1,947429 | 1,956558 | 2,011329 | 2,175644 | 2,196943 | 2,087401 | 2,136086 | 1,980899 | 1,980901 | 2,081313 | 1,986985 | 2,297357 |
| * | Lateral septal nucleus ventral part | -9,21 | *0,0011* | 0,6420 | 0,0259 | 7 | 0,5829 | 0,0259 | 7 | 0,623786 | 0,620743 | 0,611614 | 0,645086 | 0,654214 | 0,651172 | 0,687686 | 0,5964 | 0,5538 | 0,556843 | 0,562929 | 0,5964 | 0,590314 | 0,623786 |
| | Septofimbrial nucleus | -4,85 | *0,4425* | 0,5647 | 0,0726 | 7 | 0,5373 | 0,0549 | 7 | 0,593357 | 0,635957 | 0,474686 | 0,502072 | 0,4899 | 0,626829 | 0,629871 | 0,486857 | 0,547714 | 0,520328 | 0,5538 | 0,605528 | 0,593357 | 0,453386 |
| | Septohippocampal nucleus | 48,91 | *0,1658* | 0,0359 | 0,0208 | 5 | 0,0535 | 0,0179 | 7 | 0,033471 | 0,024343 | | | 0,009129 | 0,051729 | 0,060857 | 0,048686 | 0,088243 | 0,045643 | 0,033471 | 0,0426 | 0,051729 | 0,0639 |
| | Striatum-like amygdalar nuclei | -5,47 | *0,0733* | 4,49 | 0,27 | 7 | 4,25 | 0,18 | 7 | 4,460827 | 4,393888 | 4,180887 | 4,442574 | 4,564286 | 4,351288 | 5,0481 | 4,180888 | 4,080472 | 4,202184 | 4,436487 | 4,193055 | 4,077426 | 4,552114 |
| | Anterior amygdalar area | -1,00 | *0,9310* | 0,6064 | 0,1088 | 7 | 0,6003 | 0,1457 | 7 | 0,715071 | 0,7029 | 0,626829 | 0,520329 | 0,413829 | 0,675515 | 0,590314 | 0,672472 | 0,538586 | 0,705943 | 0,715072 | 0,505114 | 0,337757 | 0,727243 |
| | Bed nucleus of the accessory olfactory tract | -44,00 | *0,0998* | 0,0685 | 0,0242 | 4 | 0,0383 | 0,0221 | 5 | | | | 0,066943 | 0,076071 | 0,036514 | 0,094329 | 0,048686 | | 0,018257 | | 0,0639 | 0,012171 | 0,048686 |
| | Central amygdalar nucleus | -9,35 | *0,0541* | 1,28 | 0,12 | 7 | 1,16 | 0,07 | 7 | 1,189757 | 1,180629 | 1,186715 | 1,320601 | 1,314514 | 1,226272 | 1,527514 | 1,101515 | 1,110643 | 1,083257 | 1,195843 | 1,223228 | 1,122814 | 1,271914 |
| | Central amygdalar nucleus capsular part | -3,50 | *0,7248* | 0,2608 | 0,0595 | 7 | 0,2517 | 0,0295 | 7 | 0,246471 | 0,200829 | 0,209957 | 0,310372 | 0,313414 | 0,203872 | 0,3408 | 0,222129 | 0,264729 | 0,203871 | 0,252557 | 0,273857 | 0,289071 | 0,2556 |
| | Central amygdalar nucleus lateral part | -2,89 | *0,6141* | 0,2404 | 0,0274 | 7 | 0,2334 | 0,0225 | 7 | 0,222129 | 0,225172 | 0,225172 | 0,240386 | 0,216043 | 0,261686 | 0,292114 | 0,209957 | 0,243429 | 0,219086 | 0,222129 | 0,216043 | 0,252557 | 0,270814 |
| * | Central amygdalar nucleus medial part | -13,32 | *0,0075* | 0,7768 | 0,0555 | 7 | 0,6733 | 0,0643 | 7 | 0,721157 | 0,754629 | 0,751586 | 0,769843 | 0,785057 | 0,760715 | 0,8946 | 0,669429 | 0,602486 | 0,6603 | 0,721157 | 0,733328 | 0,581185 | 0,7455 |
| | Intercalated amygdalar nucleus | 18,11 | *0,1447* | 0,1608 | 0,0303 | 7 | 0,1900 | 0,0387 | 7 | 0,213 | 0,161272 | 0,1704 | 0,124757 | 0,152143 | 0,176486 | 0,1278 | 0,139972 | 0,243429 | 0,155186 | 0,219086 | 0,158228 | 0,206914 | 0,206914 |
| | Medial amygdalar nucleus | -5,71 | *0,1092* | 2,41 | 0,19 | 7 | 2,27 | 0,07 | 7 | 2,342999 | 2,349087 | 2,196944 | 2,409944 | 2,607729 | 2,236501 | 2,708143 | 2,218244 | 2,187815 | 2,239542 | 2,306486 | 2,242585 | 2,39777 | 2,297357 |
| | Pallidum | 1,34 | *0,6701* | 9,37 | 0,58 | 7 | 9,50 | 0,49 | 7 | 9,606297 | 9,204648 | 8,553474 | 9,13162 | 9,0951 | 9,624561 | 10,39744 | 9,256376 | 9,810173 | 8,942954 | 9,375046 | 9,019024 | 9,776694 | 10,3092 |
| * | Pallidum dorsal region | -6,62 | *0,0439* | 1,91 | 0,13 | 7 | 1,78 | 0,07 | 7 | 1,913957 | 1,880487 | 1,795286 | 1,94743 | 1,813543 | 1,853101 | 2,166514 | 1,837887 | 1,795286 | 1,734428 | 1,722258 | 1,749642 | 1,728342 | 1,917 |
| | Globus pallidus external segment | -5,40 | *0,2117* | 1,50 | 0,12 | 7 | 1,42 | 0,11 | 7 | 1,5762 | 1,481872 | 1,4058 | 1,457529 | 1,338857 | 1,536644 | 1,688786 | 1,497087 | 1,317557 | 1,408842 | 1,390586 | 1,411885 | 1,274956 | 1,6188 |
| | Globus pallidus internal segment | -11,08 | *0,2452* | 0,4121 | 0,0703 | 7 | 0,3664 | 0,0695 | 7 | 0,337757 | 0,398615 | 0,389486 | 0,4899 | 0,474686 | 0,316457 | 0,477729 | 0,3408 | 0,477729 | 0,325586 | 0,331672 | 0,337757 | 0,453385 | 0,2982 |
| | Pallidum ventral region | -0,06 | *0,9817* | 3,59 | 0,16 | 7 | 3,59 | 0,19 | 7 | 3,684899 | 3,426259 | 3,429301 | 3,544931 | 3,596657 | 3,547973 | 3,891815 | 3,490159 | 3,715329 | 3,465813 | 3,398873 | 3,435384 | 3,709241 | 3,891814 |
| | Substantia innominata | -0,38 | *0,8945* | 3,20 | 0,15 | 7 | 3,19 | 0,18 | 7 | 3,280199 | 3,039816 | 3,05503 | 3,173702 | 3,182829 | 3,173701 | 3,4932 | 3,131102 | 3,334972 | 3,067199 | 3,015472 | 3,01547 | 3,252812 | 3,496243 |
| | Magnocellular nucleus | 2,57 | *0,4772* | 0,3891 | 0,0169 | 7 | 0,3990 | 0,0314 | 7 | 0,4047 | 0,386443 | 0,374272 | 0,371229 | 0,413829 | 0,374272 | 0,398614 | 0,359057 | 0,380357 | 0,398614 | 0,3834 | 0,419914 | 0,456428 | 0,395571 |
| | Pallidum medial region | 12,82 | *0,1234* | 1,59 | 0,29 | 7 | 1,80 | 0,10 | 7 | 1,713128 | 1,652272 | 1,253658 | 1,308429 | 1,3419 | 1,956558 | 1,926129 | 1,847015 | 1,795286 | 1,737471 | 1,813543 | 1,643142 | 1,78007 | 1,965686 |
| | Medial septal complex | 1,22 | *0,8000* | 1,25 | 0,14 | 7 | 1,26 | 0,07 | 7 | 1,268871 | 1,281044 | 1,068043 | 1,147158 | 1,138029 | 1,427101 | 1,411886 | 1,323644 | 1,220186 | 1,281042 | 1,278 | 1,189757 | 1,189756 | 1,366243 |
| | Medial septal nucleus | 12,31 | *0,1780* | 0,4060 | 0,0734 | 7 | 0,4560 | 0,0555 | 7 | 0,459471 | 0,456429 | 0,267772 | 0,368186 | 0,374271 | 0,459472 | 0,456429 | 0,4473 | 0,459471 | 0,441214 | 0,5112 | 0,371228 | 0,422957 | 0,538586 |
| | Diagonal band nucleus | -4,13 | *0,3557* | 0,8429 | 0,0835 | 7 | 0,8081 | 0,0444 | 7 | 0,8094 | 0,824615 | 0,800272 | 0,778972 | 0,763757 | 0,967629 | 0,955457 | 0,876343 | 0,760714 | 0,839828 | 0,7668 | 0,818528 | 0,7668 | 0,827657 |
| * | Triangular nucleus of septum | 54,92 | *0,0196* | 0,3443 | 0,1593 | 7 | 0,5334 | 0,0602 | 7 | 0,444257 | 0,371229 | 0,185614 | 0,161272 | 0,203871 | 0,529457 | 0,514243 | 0,523372 | 0,5751 | 0,456428 | 0,535543 | 0,453385 | 0,590314 | 0,599443 |
| | Pallidum caudal region | 4,82 | *0,4068* | 1,21 | 0,07 | 7 | 1,27 | 0,16 | 7 | 1,226271 | 1,211058 | 1,055872 | 1,241486 | 1,226271 | 1,259743 | 1,244529 | 1,089343 | 1,329729 | 1,074128 | 1,439272 | 1,134985 | 1,384499 | 1,421014 |
| | Bed nuclei of the stria terminalis | 5,46 | *0,3510* | 1,20 | 0,07 | 7 | 1,27 | 0,16 | 7 | 1,226271 | 1,211058 | 1,055872 | 1,208015 | 1,208014 | 1,259743 | 1,244529 | 1,089343 | 1,329729 | 1,074128 | 1,439272 | 1,134985 | 1,384499 | 1,421014 |
| | Brain stem | -3,31 | *0,0951* | 117,12 | 4,03 | 7 | 113,24 | 3,98 | 7 | 115,8263 | 116,0698 | 113,1365 | 115,7899 | 116,1702 | 116,9675 | 125,8495 | 115,1357 | 114,2197 | 109,488 | 111,5025 | 110,766 | 110,5652 | 120,984 |
| * | Interbrain | -4,50 | *0,0420* | 35,36 | 1,24 | 7 | 33,77 | 1,37 | 7 | 35,16629 | 35,30629 | 33,81224 | 35,21501 | 35,05372 | 35,05373 | 37,91096 | 34,01612 | 34,12869 | 32,2847 | 33,01805 | 33,32231 | 33,07888 | 36,53863 |
| * | Thalamus | -5,50 | *0,0114* | 19,38 | 0,37 | 7 | 18,32 | 0,80 | 7 | 19,35865 | 19,2826 | 18,92658 | 19,56254 | 19,45907 | 19,05134 | 20,04634 | 18,48841 | 18,58882 | 17,44469 | 18,08979 | 17,74898 | 17,98023 | 19,88203 |
| * | Thalamus sensory-motor cortex related | -6,70 | *0,0077* | 6,31 | 0,19 | 7 | 5,89 | 0,28 | 7 | 6,362612 | 6,146575 | 6,286545 | 6,323061 | 6,256114 | 6,094845 | 6,6882 | 5,988346 | 5,830115 | 5,55017 | 5,836202 | 5,86054 | 5,69014 | 6,441729 |
| * | Ventral group of the dorsal thalamus | -5,89 | *0,0357* | 4,55 | 0,14 | 7 | 4,28 | 0,25 | 7 | 4,579499 | 4,360417 | 4,48213 | 4,567331 | 4,567329 | 4,457788 | 4,816843 | 4,299559 | 4,153501 | 3,992227 | 4,244787 | 4,278255 | 4,186969 | 4,801629 |
| | Ventral anterior-lateral complex of the thalamus | -4,30 | *0,3604* | 0,7481 | 0,0489 | 7 | 0,7159 | 0,0743 | 7 | 0,733328 | 0,699858 | 0,797229 | 0,8094 | 0,7881 | 0,693772 | 0,715071 | 0,684643 | 0,7668 | 0,608571 | 0,666386 | 0,687685 | 0,775928 | 0,821571 |

| | | | | | | | | | | | | | | | | | | | | | | | |
|---|---|---|---|---|---|---|---|---|---|---|---|---|---|---|---|---|---|---|---|---|---|---|---|
| | Ventral medial nucleus of the thalamus | -5,16 | *0,2391* | 0,8676 | 0,0831 | 7 | 0,8229 | 0,0442 | 7 | 0,845914 | 0,818529 | 0,806357 | 0,900686 | 0,903729 | 0,775929 | 1,0224 | 0,839829 | 0,803314 | 0,778971 | 0,906772 | 0,797228 | 0,791142 | 0,842871 |
| * | Ventral posterior complex of the thalamus | -9,94 | *0,0017* | 2,68 | 0,07 | 7 | 2,41 | 0,14 | 7 | 2,723356 | 2,56513 | 2,695972 | 2,695973 | 2,732486 | 2,595558 | 2,756829 | 2,464716 | 2,376472 | 2,318656 | 2,321701 | 2,510356 | 2,245627 | 2,6625 |
| | Ventral posterolateral nucleus of the thalamus | -12,46 | *0,0936* | 0,8198 | 0,1201 | 7 | 0,7177 | 0,0848 | 7 | 0,760714 | 0,699858 | 0,900686 | 0,952415 | 0,9798 | 0,721157 | 0,7242 | 0,663343 | 0,806357 | 0,6177 | 0,642043 | 0,754628 | 0,699857 | 0,839829 |
| | Ventral posterolateral nucleus of the thalamus parvicellular par | -15,32 | *0,3658* | 0,1022 | 0,0224 | 7 | 0,0865 | 0,0375 | 7 | 0,103457 | 0,079114 | 0,124757 | 0,118671 | 0,088243 | 0,073029 | 0,1278 | 0,115629 | 0,030429 | 0,1065 | 0,1065 | 0,109543 | 0,033471 | 0,103457 |
| * | Ventral posteromedial nucleus of the thalamus | -10,88 | *0,0005* | 1,53 | 0,07 | 7 | 1,37 | 0,06 | 7 | 1,545771 | 1,500129 | 1,487958 | 1,506215 | 1,570114 | 1,460572 | 1,667486 | 1,427101 | 1,329729 | 1,347985 | 1,287129 | 1,414928 | 1,317556 | 1,445357 |
| | Ventral posteromedial nucleus of the thalamus parvicellular par | 8,12 | *0,6481* | 0,2247 | 0,0960 | 7 | 0,2430 | 0,0332 | 7 | 0,313414 | 0,286029 | 0,182571 | 0,118671 | 0,094329 | 0,3408 | 0,237343 | 0,258643 | 0,209957 | 0,246471 | 0,286029 | 0,231257 | 0,194743 | 0,273857 |
| | Posterior triangular thalamic nucleus | 30,16 | *0,1356* | 0,2508 | 0,0921 | 7 | 0,3265 | 0,0845 | 7 | 0,2769 | 0,2769 | 0,182571 | 0,161272 | 0,143014 | 0,392529 | 0,322543 | 0,310372 | 0,206914 | 0,286028 | 0,349929 | 0,282986 | 0,374271 | 0,474686 |
| | Subparafascicular nucleus | 21,79 | *0,1345* | 0,1895 | 0,0606 | 7 | 0,2308 | 0,0251 | 7 | 0,209957 | 0,2343 | 0,194743 | 0,139972 | 0,076071 | 0,237343 | 0,2343 | 0,252557 | 0,200829 | 0,237343 | 0,2343 | 0,222128 | 0,200828 | 0,267771 |
| | Subparafascicular nucleus magnocellular part | -20,11 | *0,2154* | 0,0800 | 0,0136 | 7 | 0,0639 | 0,0288 | 7 | 0,069986 | 0,088243 | 0,088243 | 0,094329 | 0,066943 | 0,060857 | 0,091286 | 0,100414 | 0,030429 | 0,082157 | 0,0426 | 0,0852 | 0,030429 | 0,076071 |
| * | Subparafascicular nucleus parvicellular part | 52,38 | *0,0477* | 0,1095 | 0,0606 | 7 | 0,1669 | 0,0204 | 7 | 0,139971 | 0,146057 | 0,1065 | 0,045643 | 0,009129 | 0,176486 | 0,143014 | 0,152143 | 0,1704 | 0,155186 | 0,1917 | 0,136929 | 0,1704 | 0,1917 |
| * | Subparafascicular area | -61,76 | *0,0266* | 0,1565 | 0,0877 | 7 | 0,0598 | 0,0235 | 6 | 0,091286 | 0,076071 | 0,252557 | 0,228214 | 0,2556 | 0,051729 | 0,139971 | 0,066943 | 0,0639 | 0,079114 | 0,024343 | 0,0852 | | 0,039557 |
| | Peripeduncular nucleus | 29,73 | *0,2618* | 0,0483 | 0,0264 | 7 | 0,0626 | 0,0181 | 7 | 0,045643 | 0,051729 | 0,024343 | 0,036514 | 0,088243 | 0,076071 | 0,015214 | 0,054771 | 0,091286 | 0,036514 | 0,060857 | 0,051729 | 0,079114 | 0,0639 |
| * | Geniculate group dorsal thalamus | -7,73 | *0,0156* | 1,37 | 0,08 | 7 | 1,26 | 0,05 | 7 | 1,436228 | 1,424058 | 1,332772 | 1,351029 | 1,268871 | 1,271915 | 1,481872 | 1,314515 | 1,3206 | 1,204971 | 1,271915 | 1,223228 | 1,223228 | 1,268871 |
| | Medial geniculate complex | -4,05 | *0,3255* | 0,6555 | 0,0346 | 7 | 0,6290 | 0,0583 | 7 | 0,678557 | 0,639 | 0,654215 | 0,626829 | 0,663343 | 0,611615 | 0,715071 | 0,620743 | 0,7455 | 0,5538 | 0,614657 | 0,608571 | 0,645085 | 0,614657 |
| | Medial geniculate complex dorsal part | 8,82 | *0,3040* | 0,1478 | 0,0118 | 7 | 0,1608 | 0,0291 | 7 | 0,130843 | 0,1491 | 0,161271 | 0,139972 | 0,164314 | 0,146057 | 0,143014 | 0,136929 | 0,213 | 0,133886 | 0,136929 | 0,164314 | 0,182571 | 0,158229 |
| * | Medial geniculate complex ventral part | -19,71 | *0,0281* | 0,2669 | 0,0443 | 7 | 0,2143 | 0,0328 | 7 | 0,267771 | 0,222129 | 0,289072 | 0,310372 | 0,313414 | 0,194743 | 0,270814 | 0,197786 | 0,279943 | 0,179529 | 0,1917 | 0,216043 | 0,225171 | 0,209957 |
| | Medial geniculate complex medial part | 5,42 | *0,5518* | 0,2408 | 0,0506 | 7 | 0,2539 | 0,0232 | 7 | 0,279943 | 0,267772 | 0,203871 | 0,176486 | 0,185614 | 0,270814 | 0,301243 | 0,286029 | 0,252557 | 0,240386 | 0,286029 | 0,228214 | 0,237343 | 0,246471 |
| * | Dorsal part of the lateral geniculate complex | -11,12 | *0,0234* | 0,7112 | 0,0654 | 7 | 0,6320 | 0,0443 | 7 | 0,757671 | 0,785058 | 0,678557 | 0,7242 | 0,605529 | 0,6603 | 0,7668 | 0,693772 | 0,5751 | 0,651171 | 0,657257 | 0,614657 | 0,578143 | 0,654214 |
| * | Dorsal part of the lateral geniculate complex shell | -40,69 | *0,0038* | 0,2404 | 0,0595 | 7 | 0,1426 | 0,0252 | 7 | 0,261686 | 0,279943 | 0,2556 | 0,295157 | 0,203871 | 0,121714 | 0,264729 | 0,161272 | 0,1065 | 0,158229 | 0,1491 | 0,1704 | 0,109543 | 0,143014 |
| | Dorsal part of the lateral geniculate complex core | -0,23 | *0,9573* | 0,3856 | 0,0313 | 7 | 0,3847 | 0,0282 | 7 | 0,4047 | 0,407743 | 0,359057 | 0,359057 | 0,3408 | 0,416872 | 0,410786 | 0,435129 | 0,377314 | 0,380357 | 0,401657 | 0,343843 | 0,3834 | 0,371229 |
| | Dorsal part of the lateral geniculate complex ipsilateral zone | 22,96 | *0,0923* | 0,0852 | 0,0217 | 7 | 0,1048 | 0,0180 | 7 | 0,091286 | 0,097371 | 0,0639 | 0,069986 | 0,060857 | 0,121714 | 0,091286 | 0,097371 | 0,091286 | 0,112586 | 0,1065 | 0,100414 | 0,0852 | 0,139971 |
| * | Thalamus polymodal association cortex related | -4,66 | *0,0277* | 12,25 | 0,28 | 7 | 11,68 | 0,51 | 7 | 12,22924 | 12,34184 | 11,6937 | 12,34792 | 12,34183 | 12,17752 | 12,61264 | 11,78195 | 12,05276 | 11,17032 | 11,53243 | 11,09121 | 11,57198 | 12,54874 |
| * | Lateral group of the dorsal thalamus | -9,91 | *0,0121* | 3,09 | 0,17 | 7 | 2,79 | 0,21 | 7 | 3,158485 | 3,222387 | 2,872458 | 3,024602 | 3,015472 | 2,96983 | 3,383657 | 3,079373 | 2,604686 | 2,893756 | 3,018515 | 2,726399 | 2,565127 | 2,613814 |
| | Lateral posterior nucleus of the thalamus | -5,44 | *0,4588* | 1,26 | 0,18 | 7 | 1,19 | 0,16 | 7 | 1,378414 | 1,439272 | 1,007186 | 1,171501 | 1,104557 | 1,259743 | 1,472743 | 1,387544 | 0,970672 | 1,326685 | 1,329729 | 1,174542 | 1,052828 | 1,110643 |
| * | Posterior complex of the thalamus | -14,24 | *0,0005* | 1,21 | 0,07 | 7 | 1,04 | 0,07 | 7 | 1,162371 | 1,147158 | 1,2567 | 1,302343 | 1,1289 | 1,253658 | 1,232357 | 1,104558 | 1,113686 | 1,013271 | 1,098472 | 1,043699 | 0,967628 | 0,934157 |
| * | Posterior limiting nucleus of the thalamus | -14,92 | *0,0077* | 0,2069 | 0,0183 | 7 | 0,1761 | 0,0179 | 7 | 0,200829 | 0,194743 | 0,222129 | 0,2343 | 0,219086 | 0,185614 | 0,1917 | 0,173443 | 0,161271 | 0,164314 | 0,182571 | 0,158228 | 0,209957 | 0,182571 |
| * | Suprageniculate nucleus | -34,67 | *0,0021* | 0,1730 | 0,0329 | 7 | 0,1130 | 0,0158 | 7 | 0,167357 | 0,173443 | 0,197786 | 0,182572 | 0,200829 | 0,103457 | 0,185614 | 0,124757 | 0,1278 | 0,112586 | 0,124757 | 0,112586 | 0,082157 | 0,1065 |
| | Ethmoid nucleus of the thalamus | 10,75 | *0,2631* | 0,2386 | 0,0515 | 7 | 0,2643 | 0,0235 | 7 | 0,249514 | 0,267772 | 0,188657 | 0,182572 | 0,188657 | 0,292114 | 0,301243 | 0,289072 | 0,231257 | 0,2769 | 0,282986 | 0,237343 | 0,252557 | 0,279943 |
| | Anterior group of the dorsal thalamus | -2,18 | *0,6313* | 2,21 | 0,06 | 7 | 2,17 | 0,25 | 7 | 2,148256 | 2,193901 | 2,230415 | 2,279101 | 2,2578 | 2,248672 | 2,136086 | 2,148258 | 2,346043 | 1,962642 | 2,029586 | 1,822671 | 2,31257 | 2,5347 |
| | Anteroventral nucleus of thalamus | -0,60 | *0,9075* | 0,4377 | 0,0388 | 7 | 0,4351 | 0,0433 | 7 | 0,456428 | 0,459472 | 0,386443 | 0,377315 | 0,450343 | 0,471643 | 0,462514 | 0,514243 | 0,386443 | 0,453386 | 0,456429 | 0,407743 | 0,4047 | 0,422957 |
| | Anteromedial nucleus | 9,61 | *0,2294* | 0,3664 | 0,0339 | 7 | 0,4017 | 0,0639 | 7 | 0,377314 | 0,346886 | 0,3621 | 0,325586 | 0,337757 | 0,392529 | 0,422957 | 0,349929 | 0,438171 | 0,389486 | 0,389486 | 0,304286 | 0,444257 | 0,495986 |
| | Anteromedial nucleus dorsal part | 4,78 | *0,5268* | 0,2365 | 0,0382 | 7 | 0,2478 | 0,0250 | 7 | 0,267771 | 0,249514 | 0,206914 | 0,188657 | 0,197786 | 0,258643 | 0,286029 | 0,237343 | 0,246471 | 0,258643 | 0,243429 | 0,203871 | 0,258643 | 0,286029 |
| | Anteromedial nucleus ventral part | 18,39 | *0,2102* | 0,1300 | 0,0197 | 7 | 0,1539 | 0,0423 | 7 | 0,109543 | 0,097371 | 0,155186 | 0,136929 | 0,139971 | 0,133886 | 0,136929 | 0,112586 | 0,1917 | 0,130843 | 0,146057 | 0,100414 | 0,185614 | 0,209957 |
| | Anterodorsal nucleus | -15,40 | *0,0695* | 0,1834 | 0,0271 | 7 | 0,1552 | 0,0259 | 7 | 0,200829 | 0,222129 | 0,133886 | 0,176486 | 0,179529 | 0,1917 | 0,179529 | 0,188657 | 0,1491 | 0,173443 | 0,164314 | 0,167357 | 0,115629 | 0,1278 |
| | Interanteromedial nucleus of the thalamus | -6,67 | *0,5585* | 0,0626 | 0,0139 | 7 | 0,0584 | 0,0100 | 5 | 0,082157 | 0,076071 | 0,060857 | 0,060857 | 0,045643 | 0,045643 | 0,066943 | 0,073029 | | 0,054771 | 0,051729 | 0,0639 | | 0,048686 |
| * | Interanterodorsal nucleus of the thalamus | -31,96 | *0,0145* | 0,1374 | 0,0337 | 7 | 0,0935 | 0,0200 | 7 | 0,121714 | 0,115629 | 0,155186 | 0,185614 | 0,173443 | 0,100414 | 0,109543 | 0,112586 | 0,0639 | 0,097371 | 0,109543 | 0,1065 | 0,066943 | 0,097371 |
| | Lateral dorsal nucleus of thalamus | 1,19 | *0,9110* | 1,03 | 0,10 | 7 | 1,04 | 0,26 | 7 | 0,909814 | 0,973715 | 1,131943 | 1,153244 | 1,071086 | 1,046743 | 0,8946 | 0,909815 | 1,308429 | 0,794185 | 0,858086 | 0,772885 | 1,281042 | 1,3419 |
| | Medial group of the dorsal thalamus | 9,90 | *0,1202* | 1,79 | 0,23 | 7 | 1,97 | 0,15 | 7 | 1,962642 | 1,980901 | 1,554901 | 1,567072 | 1,524471 | 2,063058 | 1,889614 | 1,974815 | 1,923086 | 1,913956 | 1,974815 | 1,850056 | 1,859185 | 2,288229 |
| | Intermediodorsal nucleus of the thalamus | 23,08 | *0,1381* | 0,1582 | 0,0502 | 7 | 0,1947 | 0,0331 | 7 | 0,188657 | 0,197786 | 0,091286 | 0,1065 | 0,118671 | 0,203872 | 0,200829 | 0,182572 | 0,158229 | 0,228214 | 0,213 | 0,197786 | 0,1491 | 0,2343 |
| | Mediodorsal nucleus of thalamus | 6,84 | *0,2333* | 1,27 | 0,15 | 7 | 1,36 | 0,10 | 7 | 1,375371 | 1,393629 | 1,104558 | 1,104558 | 1,116729 | 1,445358 | 1,351029 | 1,436229 | 1,278 | 1,305385 | 1,390586 | 1,277999 | 1,287128 | 1,524471 |
| | Submedial nucleus of the thalamus | 8,07 | *0,3123* | 0,2369 | 0,0300 | 7 | 0,2560 | 0,0373 | 7 | 0,264728 | 0,261686 | 0,243429 | 0,246472 | 0,200829 | 0,252557 | 0,188657 | 0,225172 | 0,292114 | 0,231257 | 0,228214 | 0,231257 | 0,264728 | 0,3195 |
| * | Perireunensis nucleus | 27,49 | *0,0347* | 0,1265 | 0,0246 | 7 | 0,1613 | 0,0296 | 7 | 0,133886 | 0,1278 | 0,115629 | 0,109543 | 0,088243 | 0,161271 | 0,1491 | 0,130843 | 0,194743 | 0,1491 | 0,143014 | 0,143014 | 0,158228 | 0,209957 |
| | Midline group of the dorsal thalamus | -6,33 | *0,3057* | 1,16 | 0,13 | 7 | 1,09 | 0,12 | 7 | 1,253657 | 1,268872 | 0,9798 | 1,043701 | 1,037614 | 1,253658 | 1,290172 | 1,150201 | 0,928072 | 1,153242 | 1,119772 | 1,192799 | 0,894599 | 1,174543 |
| | Paraventricular nucleus of the thalamus | -6,47 | *0,5114* | 0,4703 | 0,0967 | 7 | 0,4399 | 0,0687 | 7 | 0,538586 | 0,556843 | 0,3408 | 0,3621 | 0,4047 | 0,538586 | 0,550757 | 0,4686 | 0,346886 | 0,486857 | 0,474686 | 0,514243 | 0,3408 | 0,4473 |
| * | Parataenial nucleus | -17,93 | *0,0407* | 0,2400 | 0,0212 | 7 | 0,1969 | 0,0425 | 7 | 0,246471 | 0,264729 | 0,237343 | 0,258643 | 0,200829 | 0,243429 | 0,228214 | 0,231257 | 0,136929 | 0,197786 | 0,231257 | 0,225171 | 0,136928 | 0,219086 |
| | Nucleus of reuniens | -3,30 | *0,4304* | 0,3817 | 0,0307 | 7 | 0,3691 | 0,0269 | 7 | 0,395571 | 0,368186 | 0,343843 | 0,356014 | 0,374271 | 0,398614 | 0,435129 | 0,3834 | 0,365143 | 0,377314 | 0,328629 | 0,380357 | 0,3408 | 0,407743 |
| * | Xiphoid thalamic nucleus | 18,24 | *0,0393* | 0,0691 | 0,0086 | 7 | 0,0817 | 0,0114 | 7 | 0,073029 | 0,079114 | 0,057814 | 0,066943 | 0,057814 | 0,073029 | 0,076071 | 0,066943 | 0,079114 | 0,091286 | 0,0852 | 0,073029 | 0,076071 | 0,100414 |
| | Intralaminar nuclei of the dorsal thalamus | -14,50 | *0,0764* | 1,46 | 0,24 | 7 | 1,25 | 0,16 | 7 | 1,305385 | 1,287129 | 1,576201 | 1,700958 | 1,798329 | 1,159329 | 1,414929 | 1,192801 | 1,570115 | 1,116728 | 1,104558 | 1,195842 | 1,354071 | 1,223229 |
| | Rhomboid nucleus | -20,10 | *0,1713* | 0,0887 | 0,0264 | 7 | 0,0709 | 0,0184 | 7 | 0,039557 | 0,0852 | 0,094329 | 0,100414 | 0,118671 | 0,073029 | 0,109543 | 0,069986 | 0,039557 | 0,094329 | 0,076071 | 0,088243 | 0,057814 | 0,069986 |
| | Central medial nucleus of the thalamus | 2,70 | *0,7352* | 0,1774 | 0,0197 | 7 | 0,1821 | 0,0306 | 7 | 0,203871 | 0,179529 | 0,146057 | 0,1704 | 0,188657 | 0,161271 | 0,1917 | 0,185614 | 0,228214 | 0,139971 | 0,152143 | 0,182571 | 0,209957 | 0,176486 |
| * | Paracentral nucleus | 34,20 | *0,0328* | 0,1678 | 0,0270 | 7 | 0,2252 | 0,0537 | 7 | 0,182571 | 0,173443 | 0,1278 | 0,143014 | 0,188657 | 0,203872 | 0,155186 | 0,182572 | 0,289071 | 0,173443 | 0,243429 | 0,155186 | 0,273857 | 0,258643 |
| | Central lateral nucleus of the thalamus | 11,28 | *0,2114* | 0,3660 | 0,0762 | 7 | 0,4073 | 0,0229 | 7 | 0,401657 | 0,407743 | 0,282986 | 0,2769 | 0,301243 | 0,441214 | 0,450343 | 0,438172 | 0,3834 | 0,413828 | 0,410786 | 0,371228 | 0,422957 | 0,410786 |
| * | Parafascicular nucleus | -57,74 | *0,0428* | 0,4886 | 0,2855 | 7 | 0,2065 | 0,1255 | 7 | 0,328628 | 0,286029 | 0,7881 | 0,785058 | 0,775929 | 0,112586 | 0,343843 | 0,1917 | 0,459471 | 0,164314 | 0,057814 | 0,231257 | 0,209957 | 0,130843 |
| | Posterior intralaminar thalamic nucleus | -8,96 | *0,3488* | 0,1747 | 0,0359 | 7 | 0,1591 | 0,0221 | 7 | 0,1491 | 0,155186 | 0,136929 | 0,225172 | 0,225171 | 0,167357 | 0,164314 | 0,124757 | 0,1704 | 0,130843 | 0,164314 | 0,167357 | 0,179528 | 0,176486 |
| | Reticular nucleus of the thalamus | 3,00 | *0,7195* | 1,22 | 0,16 | 7 | 1,25 | 0,21 | 7 | 1,077171 | 1,058915 | 1,162372 | 1,351029 | 1,481871 | 1,290172 | 1,098471 | 1,101515 | 1,463614 | 1,031528 | 1,131943 | 1,092385 | 1,460571 | 1,494043 |
| * | Geniculate group ventral thalamus | -14,63 | *0,0207* | 0,5377 | 0,0645 | 7 | 0,4590 | 0,0399 | 7 | 0,5538 | 0,556843 | 0,541629 | 0,541629 | 0,4899 | 0,435129 | 0,645086 | 0,410786 | 0,453386 | 0,419914 | 0,502072 | 0,517285 | 0,441214 | 0,4686 |
| | Intergeniculate leaflet of the lateral geniculate complex | 5,30 | *0,7157* | 0,0574 | 0,0143 | 7 | 0,0604 | 0,0161 | 7 | 0,051729 | 0,036514 | 0,048686 | 0,066943 | 0,079114 | 0,051729 | 0,051729 | 0,051729 | 0,079114 | 0,045643 | 0,045643 | 0,048686 | 0,069986 | 0,082157 |
| | Intermediate geniculate nucleus | 26,39 | *0,4606* | 0,0299 | 0,0216 | 6 | 0,0378 | 0,0135 | 7 | 0,0426 | 0,045643 | 0,003043 | 0,003043 | | 0,051729 | 0,033471 | 0,048686 | 0,0213 | 0,030429 | 0,057814 | 0,036514 | 0,024343 | 0,045643 |
| * | Ventral part of the lateral geniculate complex | -16,81 | *0,0299* | 0,4164 | 0,0629 | 7 | 0,3465 | 0,0367 | 7 | 0,435128 | 0,422957 | 0,429043 | 0,429043 | 0,377314 | 0,307329 | 0,514243 | 0,2982 | 0,352971 | 0,316457 | 0,380357 | 0,4047 | 0,3408 | 0,331671 |
| * | Subgeniculate nucleus | -56,25 | *0,0189* | 0,0383 | 0,0175 | 7 | 0,0167 | 0,0092 | 6 | 0,024343 | 0,051729 | 0,060857 | 0,0426 | 0,033471 | 0,009129 | 0,045643 | 0,012171 | | 0,027386 | 0,018257 | 0,027386 | 0,006086 | 0,009129 |
| * | Epithalamus | -8,50 | *0,0066* | 0,7725 | 0,0327 | 7 | 0,7068 | 0,0410 | 7 | 0,769843 | 0,772886 | 0,775929 | 0,839829 | 0,736371 | 0,757672 | 0,754629 | 0,7242 | 0,763757 | 0,678557 | 0,651172 | 0,693771 | 0,684642 | 0,751586 |
| | Medial habenula | -7,10 | *0,1957* | 0,3856 | 0,0327 | 7 | 0,3582 | 0,0413 | 7 | 0,349928 | 0,356014 | 0,416872 | 0,438172 | 0,395571 | 0,368186 | 0,374271 | 0,343843 | 0,410786 | 0,3195 | 0,316457 | 0,334714 | 0,365143 | 0,416871 |
| * | Lateral habenula | -9,89 | *0,0166* | 0,3869 | 0,0293 | 7 | 0,3486 | 0,0203 | 7 | 0,419914 | 0,416872 | 0,359057 | 0,401657 | 0,3408 | 0,389486 | 0,380357 | 0,380357 | 0,352971 | 0,359057 | 0,334714 | 0,359057 | 0,3195 | 0,334714 |
| | Hypothalamus | -3,28 | *0,2360* | 15,98 | 0,92 | 7 | 15,45 | 0,61 | 7 | 15,80764 | 16,02369 | 14,88566 | 15,65247 | 15,59464 | 16,00239 | 17,86462 | 15,52771 | 15,53987 | 14,84001 | 14,92826 | 15,57334 | 15,09865 | 16,6566 |
| | Periventricular zone | 3,41 | *0,3102* | 0,7777 | 0,0441 | 7 | 0,8042 | 0,0494 | 7 | 0,778971 | 0,769843 | 0,742457 | 0,7242 | 0,812443 | 0,760715 | 0,855043 | 0,763758 | 0,852 | 0,739414 | 0,836786 | 0,794185 | 0,772885 | 0,870257 |
| | Supraoptic nucleus | 2,14 | *0,8276* | 0,0609 | 0,0135 | 7 | 0,0622 | 0,0074 | 7 | 0,079114 | 0,054771 | 0,073029 | 0,051729 | 0,060857 | 0,039557 | 0,066943 | 0,057814 | 0,073029 | 0,051729 | 0,0639 | 0,057814 | 0,069986 | 0,060857 |
| | Paraventricular hypothalamic nucleus | 26,52 | *0,0560* | 0,1426 | 0,0231 | 7 | 0,1804 | 0,0399 | 7 | 0,1704 | 0,1278 | 0,112586 | 0,143014 | 0,152143 | 0,121714 | 0,1704 | 0,143014 | 0,182571 | 0,146057 | 0,206914 | 0,158228 | 0,1704 | 0,2556 |
| | Periventricular hypothalamic nucleus anterior part | 19,38 | *0,4204* | 0,0561 | 0,0289 | 7 | 0,0669 | 0,0183 | 7 | 0,076071 | 0,0852 | 0,073029 | 0,033471 | 0,012171 | 0,079114 | 0,033471 | 0,073029 | 0,0639 | 0,073029 | 0,0852 | 0,054771 | 0,033471 | 0,0852 |
| | Periventricular hypothalamic nucleus intermediate part | -13,06 | *0,2271* | 0,2230 | 0,0494 | 7 | 0,1939 | 0,0343 | 7 | 0,1704 | 0,206914 | 0,179529 | 0,249514 | 0,279943 | 0,185614 | 0,289071 | 0,1704 | 0,182571 | 0,1704 | 0,167357 | 0,240386 | 0,246471 | 0,179529 |

| | | | | | | | | | | | | | | | | | | | | | | | |
|---|---|---|---|---|---|---|---|---|---|---|---|---|---|---|---|---|---|---|---|---|---|---|---|
| | Arcuate hypothalamic nucleus | 4,85 | *0,4189* | 0,2869 | 0,0313 | 7 | 0,3008 | 0,0309 | 7 | 0,282986 | 0,295157 | 0,304286 | 0,231257 | 0,282986 | 0,334714 | 0,2769 | 0,3195 | 0,349929 | 0,2982 | 0,313414 | 0,282986 | 0,252557 | 0,289071 |
| | Periventricular region | 1,92 | *0,6498* | 2,02 | 0,13 | 7 | 2,06 | 0,18 | 7 | 1,923085 | 2,035673 | 1,816586 | 2,111744 | 2,063057 | 1,968729 | 2,209114 | 1,886572 | 2,163472 | 1,847014 | 1,874401 | 2,178685 | 2,212156 | 2,2365 |
| | Anteroventral preoptic nucleus | -27,58 | *0,1612* | 0,1561 | 0,0496 | 7 | 0,1130 | 0,0578 | 7 | 0,164314 | 0,179529 | 0,136929 | 0,155186 | 0,0639 | 0,164314 | 0,228214 | 0,146057 | 0,045643 | 0,1491 | 0,1278 | 0,155186 | 0,015214 | 0,152143 |
| | Anteroventral periventricular nucleus | -2,00 | *0,7978* | 0,1960 | 0,0304 | 7 | 0,1921 | 0,0252 | 7 | 0,200829 | 0,200829 | 0,1917 | 0,182572 | 0,139971 | 0,222129 | 0,2343 | 0,213 | 0,194743 | 0,209957 | 0,1704 | 0,182571 | 0,152143 | 0,222129 |
| | Dorsomedial nucleus of the hypothalamus | 25,00 | *0,0805* | 0,3530 | 0,1062 | 7 | 0,4412 | 0,0515 | 7 | 0,477728 | 0,456429 | 0,374272 | 0,307329 | 0,213 | 0,413829 | 0,228214 | 0,419915 | 0,492943 | 0,410786 | 0,456429 | 0,3834 | 0,401657 | 0,523371 |
| | Median preoptic nucleus | -7,61 | *0,4707* | 0,0400 | 0,0077 | 7 | 0,0369 | 0,0075 | 7 | 0,039557 | 0,039557 | 0,048686 | 0,030429 | 0,051729 | 0,033471 | 0,036514 | 0,033471 | 0,039557 | 0,039557 | 0,039557 | 0,024343 | 0,033471 | 0,048686 |
| | Medial preoptic area | -0,61 | *0,8733* | 0,5747 | 0,0443 | 7 | 0,5712 | 0,0349 | 7 | 0,559886 | 0,6177 | 0,5538 | 0,535543 | 0,523371 | 0,587272 | 0,645086 | 0,559886 | 0,608572 | 0,5751 | 0,535543 | 0,523371 | 0,578143 | 0,6177 |
| | Vascular organ of the lamina terminalis | 60,00 | *0,3339* | 0,0061 | 0,0047 | 6 | 0,0097 | 0,0066 | 5 | 0,015214 | 0,006086 | 0,003043 | 0,003043 | | 0,003043 | 0,006086 | 0,006086 | 0,018257 | 0,003043 | 0,015214 | | | 0,006086 |
| | Posterodorsal preoptic nucleus | 12,00 | *0,8855* | 0,0109 | 0,0080 | 7 | 0,0122 | 0,0133 | 3 | 0,003043 | 0,006086 | 0,003043 | 0,0213 | 0,018257 | 0,006086 | 0,018257 | 0,006086 | | 0,003043 | | 0,027386 | | |
| | Parastrial nucleus | -31,58 | *0,4491* | 0,0743 | 0,0592 | 7 | 0,0509 | 0,0529 | 7 | 0,006086 | 0,100414 | 0,073029 | 0,121714 | 0,164314 | 0,006086 | 0,048686 | 0,003043 | 0,088243 | 0,006086 | 0,006086 | 0,1278 | 0,100414 | 0,024343 |
| | Periventricular hypothalamic nucleus posterior part | 3,56 | *0,8607* | 0,1221 | 0,0533 | 7 | 0,1265 | 0,0353 | 7 | 0,091286 | 0,082157 | 0,109543 | 0,112586 | 0,139971 | 0,0852 | 0,2343 | 0,0852 | 0,1278 | 0,082157 | 0,118671 | 0,179528 | 0,155186 | 0,136929 |
| | Periventricular hypothalamic nucleus preoptic part | 1,14 | *0,8536* | 0,1521 | 0,0200 | 7 | 0,1539 | 0,0138 | 7 | 0,139971 | 0,143014 | 0,130843 | 0,158229 | 0,1917 | 0,143014 | 0,158229 | 0,146057 | 0,158229 | 0,136929 | 0,139971 | 0,161271 | 0,176486 | 0,158229 |
| | Subparaventricular zone | 8,70 | *0,2129* | 0,1000 | 0,0085 | 7 | 0,1087 | 0,0150 | 7 | 0,091286 | 0,091286 | 0,091286 | 0,103457 | 0,109543 | 0,109543 | 0,103457 | 0,100414 | 0,1065 | 0,100414 | 0,091286 | 0,136929 | 0,1065 | 0,118671 |
| | Suprachiasmatic nucleus | -2,11 | *0,9224* | 0,0617 | 0,0196 | 7 | 0,0604 | 0,0285 | 7 | 0,051729 | 0,051729 | 0,036514 | 0,073029 | 0,091286 | 0,048686 | 0,079114 | 0,060857 | 0,082157 | 0,015214 | 0,036514 | 0,073029 | 0,100414 | 0,054771 |
| | Subfornical organ | 46,55 | *0,3405* | 0,0353 | 0,0337 | 5 | 0,0517 | 0,0081 | 7 | 0,012171 | | 0,006086 | 0,015214 | | 0,079114 | 0,0639 | 0,054771 | 0,045643 | 0,048686 | 0,051729 | 0,057814 | 0,0639 | 0,039557 |
| | Ventromedial preoptic nucleus | 48,61 | *0,2705* | 0,0365 | 0,0214 | 6 | 0,0543 | 0,0302 | 6 | 0,039557 | 0,0213 | 0,030429 | 0,036514 | 0,076071 | | 0,015214 | | 0,082157 | 0,0213 | 0,054771 | 0,051729 | 0,094329 | 0,0213 |
| | Ventrolateral preoptic nucleus | -15,44 | *0,5654* | 0,0648 | 0,0342 | 7 | 0,0548 | 0,0288 | 7 | 0,030429 | 0,039557 | 0,027386 | 0,1065 | 0,073029 | 0,066943 | 0,109543 | 0,051729 | 0,012171 | 0,045643 | 0,030429 | 0,094329 | 0,082157 | 0,066943 |
| | Hypothalamic medial zone | -7,42 | *0,1471* | 4,01 | 0,44 | 7 | 3,71 | 0,22 | 7 | 3,773142 | 3,849216 | 3,748801 | 4,028745 | 4,129157 | 3,611873 | 4,932472 | 3,563188 | 3,788358 | 3,46277 | 3,499287 | 3,690984 | 3,922241 | 4,062214 |
| | Anterior hypothalamic nucleus | -1,20 | *0,8365* | 0,7638 | 0,0816 | 7 | 0,7546 | 0,0803 | 7 | 0,812443 | 0,794186 | 0,757672 | 0,7029 | 0,611614 | 0,8307 | 0,836786 | 0,800272 | 0,693772 | 0,736371 | 0,6816 | 0,812442 | 0,672471 | 0,885471 |
| | Mammillary body | -6,41 | *0,5882* | 1,07 | 0,29 | 7 | 1,00 | 0,14 | 7 | 0,876343 | 0,864172 | 0,879386 | 1,265829 | 1,329729 | 0,772886 | 1,509257 | 0,976758 | 0,903729 | 0,900685 | 0,839829 | 1,013271 | 1,138028 | 1,244529 |
| | Lateral mammillary nucleus | -7,08 | *0,7932* | 0,0922 | 0,0529 | 7 | 0,0856 | 0,0364 | 7 | 0,027386 | 0,054771 | 0,045643 | 0,136929 | 0,152143 | 0,079114 | 0,1491 | 0,097371 | 0,0426 | 0,094329 | 0,027386 | 0,112586 | 0,100414 | 0,124757 |
| | Medial mammillary nucleus | -9,54 | *0,5350* | 0,5786 | 0,2049 | 7 | 0,5234 | 0,0952 | 7 | 0,438171 | 0,4047 | 0,419914 | 0,772886 | 0,727243 | 0,410786 | 0,876343 | 0,5112 | 0,450343 | 0,465557 | 0,435129 | 0,5538 | 0,5325 | 0,715071 |
| | Medial mammillary nucleus median part | -21,29 | *0,6155* | 0,0786 | 0,0722 | 6 | 0,0619 | 0,0290 | 6 | 0,027386 | 0,003043 | 0,009129 | 0,1491 | 0,139971 | | 0,143014 | 0,036514 | 0,057814 | 0,027386 | | 0,057814 | 0,091286 | 0,100414 |
| | Medial mammillary nucleus lateral part | 12,01 | *0,3958* | 0,2100 | 0,0592 | 7 | 0,2352 | 0,0471 | 7 | 0,1704 | 0,155186 | 0,164314 | 0,252557 | 0,243429 | 0,173443 | 0,310371 | 0,273857 | 0,1917 | 0,261686 | 0,1704 | 0,237343 | 0,209957 | 0,301243 |
| | Medial mammillary nucleus medial part | -47,43 | *0,1234* | 0,1604 | 0,1041 | 7 | 0,0843 | 0,0572 | 7 | 0,097371 | 0,079114 | 0,103457 | 0,2343 | 0,252557 | 0,0426 | 0,313414 | 0,018257 | 0,112586 | 0,012171 | 0,066943 | 0,091286 | 0,173443 | 0,115629 |
| | Medial mammillary nucleus posterior part | 19,97 | *0,3573* | 0,0461 | 0,0102 | 7 | 0,0553 | 0,0209 | 6 | 0,039557 | 0,054771 | 0,036514 | 0,048686 | 0,039557 | 0,0639 | 0,039557 | 0,060857 | 0,015214 | 0,054771 | 0,076071 | 0,060857 | | 0,0639 |
| | Medial mammillary nucleus dorsal part | 9,17 | *0,5634* | 0,0948 | 0,0269 | 7 | 0,1035 | 0,0278 | 7 | 0,103457 | 0,112586 | 0,1065 | 0,088243 | 0,051729 | 0,130843 | 0,069986 | 0,121714 | 0,073029 | 0,109543 | 0,121714 | 0,1065 | 0,057814 | 0,133886 |
| | Supramammillary nucleus | 5,68 | *0,5663* | 0,2604 | 0,0557 | 7 | 0,2752 | 0,0353 | 7 | 0,3195 | 0,322543 | 0,304286 | 0,185614 | 0,206914 | 0,252557 | 0,231257 | 0,331672 | 0,267771 | 0,310371 | 0,258643 | 0,2343 | 0,279943 | 0,243429 |
| | Tuberomammillary nucleus | -15,53 | *0,6078* | 0,1400 | 0,0846 | 7 | 0,1182 | 0,0687 | 7 | 0,091286 | 0,082157 | 0,109543 | 0,1704 | 0,243429 | 0,030429 | 0,252557 | 0,036514 | 0,143014 | 0,030429 | 0,118671 | 0,112586 | 0,225171 | 0,161271 |
| | Tuberomammillary nucleus dorsal part | -13,85 | *0,7016* | 0,0494 | 0,0208 | 4 | 0,0426 | 0,0304 | 5 | 0,057814 | 0,060857 | 0,060857 | | | 0,018257 | | 0,015214 | 0,082157 | 0,012171 | 0,0639 | | 0,039557 | |
| | Tuberomammillary nucleus ventral part | -21,40 | *0,6268* | 0,1117 | 0,1071 | 7 | 0,0878 | 0,0667 | 7 | 0,033471 | 0,0213 | 0,048686 | 0,1704 | 0,243429 | 0,012171 | 0,252557 | 0,0213 | 0,060857 | 0,018257 | 0,054771 | 0,112586 | 0,185614 | 0,161271 |
| | Medial preoptic nucleus | 1,63 | *0,8383* | 0,3738 | 0,0388 | 7 | 0,3799 | 0,0663 | 7 | 0,380357 | 0,374272 | 0,328629 | 0,389486 | 0,4473 | 0,349929 | 0,346886 | 0,307329 | 0,486857 | 0,3621 | 0,343843 | 0,328628 | 0,453385 | 0,377314 |
| * | Dorsal premammillary nucleus | -20,97 | *0,0398* | 0,2708 | 0,0317 | 4 | 0,2140 | 0,0407 | 6 | 0,2556 | 0,289072 | 0,2343 | | | 0,304286 | | 0,228214 | 0,216043 | 0,237343 | 0,267772 | 0,155186 | | 0,179529 |
| | Ventral premammillary nucleus | -27,08 | *0,2182* | 0,2087 | 0,0640 | 7 | 0,1521 | 0,0946 | 7 | 0,2343 | 0,203872 | 0,216043 | 0,167357 | 0,216043 | 0,1065 | 0,316457 | 0,045643 | 0,219086 | 0,051729 | 0,237343 | 0,088243 | 0,279943 | 0,143014 |
| | Paraventricular hypothalamic nucleus descending division | -10,40 | *0,6488* | 0,2173 | 0,0635 | 7 | 0,1947 | 0,1013 | 6 | 0,2343 | 0,267772 | 0,219086 | 0,197786 | 0,0852 | 0,264729 | 0,252557 | 0,286029 | 0,024343 | 0,258643 | 0,1278 | 0,267771 | | 0,203871 |
| | Ventromedial hypothalamic nucleus | -4,40 | *0,5472* | 0,5634 | 0,0668 | 7 | 0,5386 | 0,0820 | 7 | 0,629871 | 0,587272 | 0,602486 | 0,480772 | 0,5112 | 0,492943 | 0,639 | 0,432086 | 0,6177 | 0,459471 | 0,593357 | 0,474685 | 0,632914 | 0,559886 |
| | Posterior hypothalamic nucleus | -18,70 | *0,2925* | 0,6577 | 0,2648 | 7 | 0,5347 | 0,1170 | 7 | 0,349928 | 0,4686 | 0,5112 | 0,824615 | 0,928071 | 0,4899 | 1,031529 | 0,486857 | 0,626829 | 0,456428 | 0,407743 | 0,550757 | 0,7455 | 0,4686 |
| | Hypothalamic lateral zone | -2,56 | *0,2302* | 6,04 | 0,24 | 7 | 5,89 | 0,21 | 7 | 6,228727 | 6,094846 | 5,811859 | 5,951832 | 5,659714 | 6,298717 | 6,2622 | 5,912275 | 6,030944 | 5,57147 | 5,964002 | 5,744911 | 5,781425 | 6,2196 |
| | Lateral hypothalamic area | -0,91 | *0,7635* | 2,35 | 0,15 | 7 | 2,32 | 0,11 | 7 | 2,501228 | 2,467758 | 2,327786 | 2,233458 | 2,078271 | 2,400815 | 2,409943 | 2,388644 | 2,355172 | 2,266928 | 2,373429 | 2,203027 | 2,193899 | 2,489057 |
| | Lateral preoptic area | -10,86 | *0,1925* | 0,6125 | 0,1057 | 7 | 0,5460 | 0,0690 | 7 | 0,499028 | 0,529457 | 0,514243 | 0,739415 | 0,730286 | 0,581186 | 0,693771 | 0,526415 | 0,517286 | 0,529457 | 0,462514 | 0,6816 | 0,581185 | 0,523371 |
| | Preparasubthalamic nucleus | -20,00 | *0,4718* | 0,0464 | 0,0152 | 4 | 0,0371 | 0,0213 | 5 | 0,057814 | 0,036514 | 0,030429 | | | 0,060857 | | 0,018257 | | 0,036514 | 0,033471 | 0,024343 | | 0,073029 |
| | Parasubthalamic nucleus | -2,34 | *0,7450* | 0,1856 | 0,0281 | 7 | 0,1813 | 0,0200 | 7 | 0,206914 | 0,222129 | 0,188657 | 0,158229 | 0,139971 | 0,188657 | 0,194743 | 0,222129 | 0,161271 | 0,173443 | 0,185614 | 0,185614 | 0,1704 | 0,1704 |
| | Perifornical nucleus | 2,25 | *0,7418* | 0,2126 | 0,0237 | 7 | 0,2173 | 0,0290 | 7 | 0,216043 | 0,222129 | 0,203871 | 0,194743 | 0,185614 | 0,206914 | 0,258643 | 0,200829 | 0,261686 | 0,1704 | 0,2343 | 0,203871 | 0,222128 | 0,228214 |
| | Retrochiasmatic area | 8,27 | *0,7243* | 0,1839 | 0,0679 | 7 | 0,1991 | 0,0882 | 7 | 0,203871 | 0,228214 | 0,176486 | 0,213 | 0,060857 | 0,267772 | 0,136929 | 0,264729 | 0,176486 | 0,261686 | 0,1704 | 0,164314 | 0,045643 | 0,310371 |
| | Subthalamic nucleus | 0,45 | *0,9552* | 0,1917 | 0,0261 | 7 | 0,1926 | 0,0304 | 7 | 0,228214 | 0,176486 | 0,155186 | 0,188657 | 0,173443 | 0,219086 | 0,200829 | 0,164314 | 0,200829 | 0,164314 | 0,252557 | 0,1917 | 0,197786 | 0,176486 |
| | Tuberal nucleus | -0,92 | *0,9110* | 0,5642 | 0,0933 | 7 | 0,5590 | 0,0767 | 7 | 0,663343 | 0,635957 | 0,626829 | 0,4047 | 0,480771 | 0,590315 | 0,547714 | 0,559886 | 0,635957 | 0,523371 | 0,629872 | 0,474685 | 0,632914 | 0,456429 |
| | Zona incerta | -4,55 | *0,2264* | 1,72 | 0,11 | 7 | 1,64 | 0,12 | 7 | 1,652271 | 1,576201 | 1,588372 | 1,81963 | 1,8105 | 1,783115 | 1,819629 | 1,567072 | 1,722257 | 1,445357 | 1,621843 | 1,615756 | 1,73747 | 1,792243 |
| | Fields of Forel | -12,95 | *0,2339* | 0,2317 | 0,0470 | 7 | 0,2017 | 0,0424 | 7 | 0,173443 | 0,1704 | 0,228214 | 0,261686 | 0,301243 | 0,237343 | 0,249514 | 0,161272 | 0,237343 | 0,173443 | 0,173443 | 0,249514 | 0,252557 | 0,164314 |
| | Median eminence | 4,21 | *0,7696* | 0,0826 | 0,0196 | 7 | 0,0861 | 0,0236 | 7 | 0,060857 | 0,079114 | 0,060857 | 0,082157 | 0,112586 | 0,103457 | 0,079114 | 0,109543 | 0,076071 | 0,100414 | 0,073029 | 0,097371 | 0,0426 | 0,103457 |
| | Midbrain | -2,09 | *0,3947* | 33,54 | 1,23 | 7 | 32,84 | 1,69 | 7 | 33,23408 | 32,96025 | 33,23714 | 33,21585 | 32,60726 | 33,26149 | 36,27695 | 33,85789 | 32,28168 | 32,01389 | 31,70962 | 31,81306 | 31,91651 | 36,29824 |
| | Midbrain sensory related | 0,05 | *0,9892* | 5,90 | 0,27 | 7 | 5,91 | 0,51 | 7 | 5,684055 | 5,629289 | 5,939659 | 5,857503 | 5,8788 | 5,884888 | 6,459986 | 6,103975 | 5,942701 | 5,534955 | 5,410202 | 5,51974 | 5,939654 | 6,904243 |
| | Superior colliculus sensory related | -2,21 | *0,5744* | 1,85 | 0,09 | 7 | 1,81 | 0,16 | 7 | 1,743557 | 1,780072 | 1,935258 | 1,859187 | 1,834843 | 1,795286 | 1,9809 | 1,853101 | 1,889615 | 1,691828 | 1,630972 | 1,679656 | 1,786156 | 2,111743 |
| | Superior colliculus optic layer | -6,98 | *0,1868* | 0,5790 | 0,0422 | 7 | 0,5386 | 0,0629 | 7 | 0,529457 | 0,514243 | 0,605529 | 0,593357 | 0,632914 | 0,587272 | 0,590314 | 0,584229 | 0,565972 | 0,477728 | 0,480772 | 0,526414 | 0,4899 | 0,645086 |
| | Superior colliculus superficial gray layer | 1,43 | *0,7194* | 0,9755 | 0,0756 | 7 | 0,9894 | 0,0655 | 7 | 0,9372 | 0,958501 | 1,0224 | 0,952415 | 0,921986 | 0,909815 | 1,125857 | 1,028486 | 0,9585 | 0,995014 | 0,925029 | 0,897642 | 1,043699 | 1,077171 |
| | Superior colliculus zonal layer | -4,90 | *0,6137* | 0,2925 | 0,0186 | 7 | 0,2782 | 0,0693 | 7 | 0,2769 | 0,307329 | 0,307329 | 0,313414 | 0,279943 | 0,2982 | 0,264729 | 0,240386 | 0,365143 | 0,219086 | 0,225172 | 0,2556 | 0,252557 | 0,389486 |
| | Inferior colliculus | 1,88 | *0,6512* | 3,79 | 0,19 | 7 | 3,86 | 0,36 | 7 | 3,703156 | 3,578402 | 3,727501 | 3,709245 | 3,764014 | 3,846173 | 4,177843 | 4,010488 | 3,837043 | 3,59057 | 3,54493 | 3,581441 | 3,882683 | 4,5582 |
| | Inferior colliculus central nucleus | 1,95 | *0,7394* | 0,9802 | 0,1184 | 7 | 0,9994 | 0,0894 | 7 | 0,985885 | 1,022401 | 0,803315 | 0,915901 | 1,010229 | 0,934158 | 1,189757 | 1,092386 | 0,815486 | 1,0224 | 1,065 | 1,013271 | 0,979799 | 1,007186 |
| | Inferior colliculus dorsal nucleus | -3,13 | *0,6716* | 1,08 | 0,09 | 7 | 1,05 | 0,19 | 7 | 1,004143 | 0,964586 | 1,201929 | 1,141072 | 1,1502 | 1,0437 | 1,083257 | 0,946329 | 1,180629 | 0,943285 | 0,903729 | 0,918942 | 1,049785 | 1,408843 |
| | Inferior colliculus external nucleus | 5,00 | *0,3651* | 1,72 | 0,12 | 7 | 1,81 | 0,21 | 7 | 1,713128 | 1,591415 | 1,722258 | 1,652272 | 1,603586 | 1,868315 | 1,904829 | 1,971773 | 1,840929 | 1,624885 | 1,576201 | 1,649228 | 1,853099 | 2,142171 |
| | Nucleus of the brachium of the inferior colliculus | 0,00 | *1,0000* | 0,0982 | 0,0198 | 7 | 0,0982 | 0,0208 | 7 | 0,112586 | 0,1065 | 0,097371 | 0,094329 | 0,057814 | 0,100414 | 0,118671 | 0,100414 | 0,051729 | 0,109543 | 0,103457 | 0,1065 | 0,1065 | 0,109543 |
| | Nucleus sagulum | -12,74 | *0,4143* | 0,0922 | 0,0285 | 7 | 0,0804 | 0,0231 | 7 | 0,0639 | 0,066943 | 0,091286 | 0,121714 | 0,139971 | 0,082157 | 0,079114 | 0,060857 | 0,121714 | 0,057814 | 0,076071 | 0,079114 | 0,100414 | 0,066943 |
| * | Parabigeminal nucleus | -26,55 | *0,0373* | 0,0491 | 0,0127 | 7 | 0,0361 | 0,0057 | 7 | 0,033471 | 0,051729 | 0,069986 | 0,054771 | 0,048686 | 0,033471 | 0,051729 | 0,039557 | 0,036514 | 0,045643 | 0,027386 | 0,036514 | 0,033471 | 0,033471 |
| * | Midbrain trigeminal nucleus | -53,13 | *0,0180* | 0,0162 | 0,0060 | 6 | 0,0076 | 0,0042 | 6 | 0,012171 | 0,012171 | 0,015214 | 0,012171 | 0,027386 | | 0,018257 | 0,012171 | 0,003043 | 0,009129 | | 0,006086 | 0,003043 | 0,012171 |
| | Subcommissural organ | 26,83 | *0,4756* | 0,0178 | 0,0134 | 7 | 0,0226 | 0,0107 | 7 | 0,015214 | 0,033471 | 0,003043 | 0,006086 | 0,006086 | 0,027386 | 0,033471 | 0,027386 | 0,003043 | 0,030429 | 0,027386 | 0,030429 | 0,027386 | 0,012171 |
| | Midbrain motor related | -4,31 | *0,0630* | 19,73 | 0,64 | 7 | 18,88 | 0,88 | 7 | 19,93679 | 19,53211 | 19,67208 | 19,59601 | 19,1487 | 19,16392 | 21,02614 | 19,453 | 18,14456 | 18,64358 | 18,51579 | 18,57663 | 18,1841 | 20,61231 |
| | Substantia nigra reticular part | -6,62 | *0,0760* | 1,59 | 0,12 | 7 | 1,48 | 0,07 | 7 | 1,539685 | 1,521429 | 1,445358 | 1,673572 | 1,786157 | 1,484915 | 1,664443 | 1,518387 | 1,442314 | 1,4697 | 1,533601 | 1,490999 | 1,363199 | 1,560986 |
| | Ventral tegmental area | 0,80 | *0,9296* | 0,4351 | 0,0755 | 7 | 0,4386 | 0,0684 | 7 | 0,483814 | 0,520329 | 0,477729 | 0,352972 | 0,325586 | 0,486857 | 0,398614 | 0,508157 | 0,365143 | 0,474686 | 0,441214 | 0,432086 | 0,334714 | 0,514243 |
| | Paranigral nucleus | 0,95 | *0,9644* | 0,0228 | 0,0039 | 4 | 0,0230 | 0,0114 | 7 | 0,027386 | 0,0213 | 0,018257 | | | 0,024343 | | 0,018257 | 0,039557 | 0,018257 | 0,039557 | 0,015214 | 0,015214 | 0,015214 |
| | Midbrain reticular nucleus retrorubral area | 9,92 | *0,6118* | 0,1052 | 0,0420 | 7 | 0,1156 | 0,0321 | 7 | 0,066943 | 0,054771 | 0,060857 | 0,143014 | 0,1278 | 0,136929 | 0,146057 | 0,088243 | 0,158229 | 0,082157 | 0,143014 | 0,100414 | 0,146057 | 0,091286 |

| | | | | | | | | | | | | | | | | | | | | | | | |
|---|---|---|---|---|---|---|---|---|---|---|---|---|---|---|---|---|---|---|---|---|---|---|---|
| | **Midbrain reticular nucleus** | -5,71 | *0,0771* | 4,98 | 0,23 | 7 | 4,69 | 0,31 | 7 | 5,026798 | 4,965946 | 5,03593 | 4,892917 | 4,634272 | 4,880745 | 5,413243 | 5,042017 | 4,372586 | 4,892913 | 4,54603 | 4,835098 | 4,24174 | 4,929429 |
| * | **Superior colliculus motor related** | -9,72 | *0,0218* | 4,96 | 0,32 | 7 | 4,48 | 0,36 | 7 | 5,191113 | 4,835103 | 4,829016 | 5,096789 | 5,1546 | 4,339116 | 5,255015 | 4,506474 | 4,473001 | 4,269127 | 4,150459 | 4,296512 | 4,387797 | 5,245886 |
| | **Superior colliculus motor related deep gray layer** | -2,77 | *0,5669* | 1,02 | 0,06 | 7 | 0,9907 | 0,1102 | 7 | 1,049785 | 1,055872 | 0,988929 | 0,970672 | 0,952414 | 0,988929 | 1,125857 | 1,061958 | 0,931114 | 0,940243 | 0,9798 | 0,964585 | 0,858085 | 1,198886 |
| | **Superior colliculus motor related deep white layer** | -0,68 | *0,9204* | 0,2543 | 0,0401 | 7 | 0,2526 | 0,0200 | 7 | 0,295157 | 0,267772 | 0,222129 | 0,203872 | 0,249514 | 0,228214 | 0,313414 | 0,249514 | 0,231257 | 0,243428 | 0,231257 | 0,2556 | 0,273857 | 0,282986 |
| * | **Superior colliculus motor related intermediate white layer** | -10,19 | *0,0176* | 1,86 | 0,13 | 7 | 1,67 | 0,13 | 7 | 1,834842 | 1,79833 | 1,877443 | 1,94743 | 1,999157 | 1,609672 | 1,953514 | 1,640101 | 1,713129 | 1,600542 | 1,542729 | 1,570114 | 1,710085 | 1,917 |
| * | **Superior colliculus motor related intermediate gray layer** | -14,37 | *0,0108* | 1,82 | 0,18 | 7 | 1,56 | 0,14 | 7 | 2,011328 | 1,71313 | 1,740515 | 1,974815 | 1,953514 | 1,512301 | 1,862229 | 1,554901 | 1,5975 | 1,484914 | 1,396672 | 1,506214 | 1,545771 | 1,847014 |
| | **Periaqueductal gray** | -0,70 | *0,8079* | 4,20 | 0,20 | 7 | 4,17 | 0,24 | 7 | 4,259999 | 4,156545 | 4,454744 | 4,092645 | 3,882686 | 4,14133 | 4,442572 | 4,153502 | 3,952672 | 4,074384 | 4,110901 | 4,159584 | 4,074383 | 4,698172 |
| * | **Precommissural nucleus** | 66,45 | *0,0075* | 0,1348 | 0,0585 | 7 | 0,2243 | 0,0429 | 7 | 0,109543 | 0,176486 | 0,139971 | 0,0639 | 0,0639 | 0,216043 | 0,173443 | 0,182572 | 0,2343 | 0,209957 | 0,209957 | 0,173443 | 0,282986 | 0,2769 |
| | **Interstitial nucleus of Cajal** | -14,69 | *0,2385* | 0,0769 | 0,0208 | 7 | 0,0656 | 0,0114 | 7 | 0,069986 | 0,0639 | 0,079114 | 0,1065 | 0,103457 | 0,051729 | 0,0639 | 0,069986 | 0,057814 | 0,045643 | 0,0639 | 0,066943 | 0,079114 | 0,076071 |
| | **Nucleus of Darkschewitsch** | -19,50 | *0,2341* | 0,0869 | 0,0318 | 7 | 0,0700 | 0,0146 | 7 | 0,051729 | 0,0639 | 0,124757 | 0,1065 | 0,1278 | 0,0639 | 0,069986 | 0,057814 | 0,082157 | 0,0639 | 0,066943 | 0,051729 | 0,073029 | 0,094329 |
| * | **Supraoculomotor periaqueductal gray** | -72,74 | *0,0141* | 0,0491 | 0,0279 | 7 | 0,0134 | 0,0067 | 5 | 0,048686 | 0,0213 | 0,0639 | 0,082157 | 0,069986 | 0,003043 | 0,054771 | 0,0213 | 0,012171 | 0,015214 | 0,003043 | 0,015214 | | |
| | **Pretectal region** | 6,73 | *0,2259* | 1,97 | 0,23 | 7 | 2,10 | 0,15 | 7 | 1,850057 | 2,078273 | 1,850058 | 1,792244 | 1,707043 | 2,333872 | 2,169557 | 2,312573 | 1,907872 | 2,005242 | 2,154344 | 1,965685 | 2,236499 | 2,126957 |
| | **Anterior pretectal nucleus** | 8,28 | *0,1037* | 1,19 | 0,12 | 7 | 1,29 | 0,08 | 7 | 1,144114 | 1,211058 | 1,071086 | 1,153244 | 1,138029 | 1,457529 | 1,168457 | 1,381458 | 1,208014 | 1,211057 | 1,3632 | 1,217142 | 1,329728 | 1,323643 |
| | **Medial pretectal area** | 19,73 | *0,3796* | 0,0639 | 0,0237 | 4 | 0,0765 | 0,0126 | 7 | | 0,045643 | 0,048686 | | | 0,0639 | 0,097371 | 0,088243 | 0,051729 | 0,0852 | 0,069986 | 0,0852 | 0,076071 | 0,079114 |
| | **Nucleus of the optic tract** | 29,35 | *0,1827* | 0,1926 | 0,0851 | 7 | 0,2491 | 0,0617 | 7 | 0,176486 | 0,258643 | 0,143014 | 0,091286 | 0,103457 | 0,279943 | 0,295157 | 0,316457 | 0,1491 | 0,279943 | 0,295157 | 0,2556 | 0,267771 | 0,179529 |
| | **Nucleus of the posterior commissure** | -15,99 | *0,5363* | 0,2556 | 0,1005 | 7 | 0,2147 | 0,1363 | 7 | 0,213 | 0,133886 | 0,401657 | 0,359057 | 0,301243 | 0,216043 | 0,164314 | 0,133886 | 0,352971 | 0,082157 | 0,088243 | 0,121714 | 0,352971 | 0,371229 |
| | **Olivary pretectal nucleus** | 10,09 | *0,8191* | 0,0553 | 0,0316 | 6 | 0,0609 | 0,0387 | 4 | 0,030429 | 0,015214 | 0,079114 | 0,091286 | 0,079114 | 0,036514 | | | 0,082157 | | | 0,009129 | 0,054771 | 0,097371 |
| | **Posterior pretectal nucleus** | 20,17 | *0,5701* | 0,1530 | 0,1062 | 7 | 0,1839 | 0,0909 | 7 | 0,143014 | 0,286029 | 0,069986 | 0,036514 | 0,060857 | 0,179529 | 0,295157 | 0,286029 | 0,0639 | 0,252557 | 0,270814 | 0,182571 | 0,155186 | 0,076071 |
| | **Retroparafascicular nucleus** | -1,30 | *0,9563* | 0,0917 | 0,0514 | 7 | 0,0905 | 0,0167 | 4 | 0,143014 | 0,1278 | 0,036514 | 0,060857 | 0,024343 | 0,100414 | 0,1491 | 0,1065 | | 0,094329 | 0,066943 | 0,094329 | | |
| * | **Cuneiform nucleus** | -19,32 | *0,0297* | 0,5851 | 0,0946 | 7 | 0,4721 | 0,0745 | 7 | 0,645086 | 0,556843 | 0,629872 | 0,6177 | 0,565971 | 0,395572 | 0,684643 | 0,517286 | 0,450343 | 0,562928 | 0,432086 | 0,508157 | 0,334714 | 0,499029 |
| | **Red nucleus** | 3,59 | *0,4422* | 0,6651 | 0,0415 | 7 | 0,6890 | 0,0673 | 7 | 0,657257 | 0,629872 | 0,635957 | 0,672472 | 0,690729 | 0,742457 | 0,626829 | 0,626829 | 0,730286 | 0,623786 | 0,763757 | 0,611614 | 0,763757 | 0,7029 |
| | **Oculomotor nucleus** | -13,24 | *0,1841* | 0,0416 | 0,0068 | 6 | 0,0361 | 0,0071 | 7 | 0,0426 | 0,051729 | 0,039557 | 0,030429 | | 0,0426 | 0,0426 | 0,039557 | 0,024343 | 0,039557 | 0,030429 | 0,033471 | 0,039557 | 0,045643 |
| | **Medial accesory oculomotor nucleus** | 28,57 | *0,2937* | 0,0101 | 0,0046 | 6 | 0,0130 | 0,0049 | 7 | 0,012171 | 0,012171 | | 0,003043 | 0,015214 | 0,012171 | 0,006086 | 0,006086 | 0,015214 | 0,012171 | 0,0213 | 0,012171 | 0,009129 | 0,015214 |
| | **Edinger-Westphal nucleus** | 6,82 | *0,6874* | 0,0191 | 0,0042 | 7 | 0,0204 | 0,0072 | 7 | 0,018257 | 0,015214 | 0,018257 | 0,018257 | 0,027386 | 0,0213 | 0,015214 | 0,012171 | 0,024343 | 0,009129 | 0,027386 | 0,0213 | 0,0213 | 0,027386 |
| | **Paratrochlear nucleus** | 44,76 | *0,3237* | 0,0114 | 0,0067 | 4 | 0,0165 | 0,0093 | 7 | 0,009129 | 0,009129 | | | 0,006086 | 0,0213 | | 0,009129 | 0,024343 | 0,009129 | 0,0213 | 0,003043 | 0,027386 | 0,0213 |
| | **Ventral tegmental nucleus** | -28,24 | *0,0976* | 0,0569 | 0,0113 | 7 | 0,0409 | 0,0202 | 7 | 0,060857 | 0,0639 | 0,073029 | 0,051729 | 0,036514 | 0,054771 | 0,057814 | 0,048686 | 0,012171 | 0,045643 | 0,054771 | 0,051729 | 0,012171 | 0,060857 |
| | **Anterior tegmental nucleus** | 29,17 | *0,7273* | 0,0365 | 0,0264 | 4 | 0,0472 | 0,0509 | 4 | | | 0,006086 | 0,039557 | 0,069986 | | 0,030429 | 0,003043 | 0,091286 | 0,003043 | | | 0,091286 | |
| | **Medial terminal nucleus of the accessory optic tract** | -15,89 | *0,1147* | 0,0465 | 0,0092 | 7 | 0,0391 | 0,0067 | 7 | 0,045643 | 0,039557 | 0,0426 | 0,057814 | 0,060857 | 0,036514 | 0,0426 | 0,0426 | 0,045643 | 0,036514 | 0,033471 | 0,030429 | 0,048686 | 0,036514 |
| * | **Midbrain behavioral state related** | -8,70 | *0,0102* | 1,78 | 0,11 | 7 | 1,63 | 0,06 | 7 | 1,722257 | 1,673572 | 1,801372 | 1,886573 | 1,834843 | 1,630972 | 1,932214 | 1,691829 | 1,606629 | 1,637057 | 1,597501 | 1,545771 | 1,585328 | 1,731386 |
| | **Substantia nigra compact part** | -4,28 | *0,6865* | 0,1930 | 0,0461 | 7 | 0,1847 | 0,0250 | 7 | 0,216043 | 0,188657 | 0,243429 | 0,173443 | 0,103457 | 0,197786 | 0,228214 | 0,179529 | 0,194743 | 0,1917 | 0,203871 | 0,173443 | 0,136928 | 0,213 |
| * | **Pedunculopontine nucleus** | -11,13 | *0,0041* | 0,8863 | 0,0539 | 7 | 0,7877 | 0,0503 | 7 | 0,8307 | 0,858086 | 0,928072 | 0,918943 | 0,943286 | 0,806357 | 0,918943 | 0,8307 | 0,760714 | 0,842871 | 0,769843 | 0,7455 | 0,721157 | 0,842871 |
| | **Midbrain raphe nuclei** | -6,86 | *0,1989* | 0,7038 | 0,0816 | 7 | 0,6555 | 0,0426 | 7 | 0,675514 | 0,626829 | 0,629872 | 0,794186 | 0,7881 | 0,626829 | 0,785057 | 0,6816 | 0,651172 | 0,602486 | 0,623786 | 0,626828 | 0,727242 | 0,675514 |
| | **Interfascicular nucleus raphe** | -21,78 | *0,2415* | 0,0878 | 0,0280 | 7 | 0,0687 | 0,0301 | 7 | 0,057814 | 0,0639 | 0,103457 | 0,115629 | 0,100414 | 0,054771 | 0,118671 | 0,109543 | 0,036514 | 0,069986 | 0,039557 | 0,100414 | 0,0426 | 0,082157 |
| | **Interpeduncular nucleus** | -9,88 | *0,1153* | 0,3564 | 0,0472 | 7 | 0,3212 | 0,0254 | 7 | 0,343843 | 0,325586 | 0,2982 | 0,4047 | 0,4047 | 0,313414 | 0,4047 | 0,301243 | 0,331671 | 0,304286 | 0,3408 | 0,289071 | 0,3621 | 0,3195 |
| | **Interpeduncular nucleus rostral** | -38,41 | *0,1477* | 0,0713 | 0,0242 | 7 | 0,0439 | 0,0393 | 7 | 0,069986 | 0,0426 | 0,045643 | 0,091286 | 0,097371 | 0,054771 | 0,097371 | 0,012171 | 0,082157 | 0,009129 | 0,066943 | 0,012171 | 0,103457 | 0,0213 |
| | **Interpeduncular nucleus caudal** | 0,57 | *0,9251* | 0,0761 | 0,0077 | 7 | 0,0765 | 0,0092 | 7 | 0,088243 | 0,082157 | 0,0639 | 0,076071 | 0,073029 | 0,076071 | 0,073029 | 0,066943 | 0,088243 | 0,073029 | 0,0852 | 0,082157 | 0,0639 | 0,076071 |
| | **Interpeduncular nucleus apical** | 15,09 | *0,6617* | 0,0323 | 0,0176 | 5 | 0,0371 | 0,0163 | 5 | | 0,012171 | 0,015214 | 0,051729 | 0,039557 | | 0,0426 | 0,048686 | | 0,048686 | | 0,039557 | 0,009129 | 0,039557 |
| | **Interpeduncular nucleus lateral** | 9,15 | *0,6157* | 0,0665 | 0,0259 | 7 | 0,0726 | 0,0173 | 7 | 0,091286 | 0,091286 | 0,066943 | 0,051729 | 0,045643 | 0,091286 | 0,027386 | 0,073029 | 0,082157 | 0,069986 | 0,094329 | 0,066943 | 0,039557 | 0,082157 |
| | **Interpeduncular nucleus intermediate** | 8,45 | *0,5116* | 0,0309 | 0,0057 | 7 | 0,0335 | 0,0084 | 7 | 0,033471 | 0,036514 | 0,027386 | 0,0213 | 0,027386 | 0,033471 | 0,036514 | 0,036514 | 0,024343 | 0,0426 | 0,030429 | 0,036514 | 0,0213 | 0,0426 |
| | **Interpeduncular nucleus dorsomedial** | 14,24 | *0,4740* | 0,0209 | 0,0079 | 7 | 0,0238 | 0,0065 | 6 | 0,012171 | 0,018257 | 0,030429 | 0,024343 | 0,027386 | 0,009129 | 0,024343 | 0,030429 | | 0,030429 | 0,015214 | 0,0213 | 0,018257 | 0,027386 |
| * | **Interpeduncular nucleus dorsolateral** | -18,00 | *0,0261* | 0,0435 | 0,0042 | 7 | 0,0356 | 0,0067 | 7 | 0,0426 | 0,039557 | 0,045643 | 0,039557 | 0,0426 | 0,0426 | 0,051729 | 0,033471 | 0,036514 | 0,030429 | 0,048686 | 0,030429 | 0,039557 | 0,030429 |
| | **Rostral linear nucleus raphe** | -27,12 | *0,0662* | 0,0513 | 0,0156 | 7 | 0,0374 | 0,0084 | 7 | 0,039557 | 0,033471 | 0,051729 | 0,0639 | 0,079114 | 0,045643 | 0,045643 | 0,033471 | 0,039557 | 0,024343 | 0,030429 | 0,0426 | 0,0426 | 0,048686 |
| | **Central linear nucleus raphe** | 23,02 | *0,1005* | 0,0604 | 0,0127 | 7 | 0,0743 | 0,0162 | 7 | 0,076071 | 0,069986 | 0,0426 | 0,051729 | 0,069986 | 0,0639 | 0,048686 | 0,0639 | 0,0852 | 0,057814 | 0,069986 | 0,060857 | 0,103457 | 0,079114 |
| | **Dorsal nucleus raphe** | 4,12 | *0,4656* | 0,1478 | 0,0140 | 7 | 0,1539 | 0,0161 | 7 | 0,158229 | 0,133886 | 0,133886 | 0,158229 | 0,133886 | 0,1491 | 0,167357 | 0,173443 | 0,158229 | 0,146057 | 0,143014 | 0,133886 | 0,176486 | 0,146057 |
| | **Hindbrain** | -3,29 | *0,0721* | 48,21 | 1,74 | 7 | 46,63 | 1,18 | 7 | 47,42596 | 47,80331 | 46,08713 | 47,35906 | 48,50923 | 48,65226 | 51,66163 | 47,26168 | 47,80938 | 45,18946 | 46,77482 | 45,63066 | 45,5698 | 48,14713 |
| | **Pons** | -3,39 | *0,1880* | 16,92 | 0,80 | 7 | 16,34 | 0,73 | 7 | 17,34732 | 17,3656 | 17,16476 | 15,9598 | 15,72549 | 17,96199 | 16,8909 | 16,49534 | 17,17085 | 15,89892 | 17,37472 | 15,30252 | 16,06628 | 16,09671 |
| | **Pons sensory related** | -6,00 | *0,2858* | 3,91 | 0,42 | 7 | 3,68 | 0,36 | 7 | 4,126113 | 4,308688 | 3,855301 | 3,593616 | 3,310629 | 4,50343 | 3,700115 | 3,937459 | 4,129158 | 3,471899 | 4,101773 | 3,404955 | 3,426255 | 3,283243 |
| | **Nucleus of the lateral lemniscus** | -5,79 | *0,7140* | 0,8037 | 0,2565 | 7 | 0,7572 | 0,2036 | 7 | 1,0437 | 1,007186 | 0,991972 | 0,471643 | 0,5112 | 0,9798 | 0,620743 | 0,824615 | 0,976757 | 0,587271 | 1,028486 | 0,587271 | 0,791142 | 0,505114 |
| | **Principal sensory nucleus of the trigeminal** | -9,49 | *0,2279* | 1,21 | 0,18 | 7 | 1,10 | 0,16 | 7 | 1,338857 | 1,344944 | 1,274958 | 0,921986 | 0,998057 | 1,384501 | 1,229314 | 1,138029 | 1,293214 | 0,949371 | 1,2993 | 0,949371 | 1,107599 | 0,949371 |
| | **Parabrachial nucleus** | -4,98 | *0,3716* | 1,09 | 0,13 | 7 | 1,04 | 0,08 | 7 | 0,998057 | 1,083258 | 0,943286 | 1,317558 | 1,004143 | 1,104558 | 1,1928 | 1,068043 | 1,0224 | 1,049785 | 0,943286 | 1,052828 | 0,940242 | 1,186714 |
| | **Koelliker-Fuse subnucleus** | 16,75 | *0,4428* | 0,1661 | 0,0534 | 7 | 0,1939 | 0,0754 | 7 | 0,143014 | 0,176486 | 0,133886 | 0,2769 | 0,112586 | 0,1491 | 0,1704 | 0,182572 | 0,139971 | 0,1917 | 0,130843 | 0,237343 | 0,133886 | 0,3408 |
| | **Superior olivary complex** | -2,32 | *0,7934* | 0,8051 | 0,1381 | 7 | 0,7864 | 0,1226 | 7 | 0,7455 | 0,8733 | 0,645086 | 0,882429 | 0,797229 | 1,034572 | 0,657257 | 0,906772 | 0,836786 | 0,885471 | 0,8307 | 0,815485 | 0,587271 | 0,642043 |
| | **Superior olivary complex periolivary region** | 2,49 | *0,9169* | 0,3138 | 0,1402 | 7 | 0,3217 | 0,1345 | 7 | 0,286028 | 0,422957 | 0,216043 | 0,310372 | 0,2343 | 0,569015 | 0,158229 | 0,462515 | 0,352971 | 0,4473 | 0,401657 | 0,304286 | 0,1278 | 0,155186 |
| | **Superior olivary complex medial part** | -5,97 | *0,6225* | 0,1821 | 0,0444 | 7 | 0,1713 | 0,0355 | 7 | 0,167357 | 0,136929 | 0,155186 | 0,216043 | 0,243429 | 0,133886 | 0,222129 | 0,133886 | 0,161271 | 0,152143 | 0,130843 | 0,203871 | 0,197786 | 0,219086 |
| | **Superior olivary complex lateral part** | -5,06 | *0,2938* | 0,3091 | 0,0300 | 7 | 0,2934 | 0,0226 | 7 | 0,292114 | 0,313414 | 0,273857 | 0,356014 | 0,3195 | 0,331672 | 0,2769 | 0,310372 | 0,322543 | 0,286028 | 0,2982 | 0,307328 | 0,261686 | 0,267771 |
| | **Pons motor related** | -3,75 | *0,3230* | 5,88 | 0,35 | 7 | 5,66 | 0,44 | 7 | 6,049198 | 5,881846 | 6,079631 | 5,361517 | 5,598857 | 6,426517 | 5,735786 | 5,559303 | 6,304801 | 5,279355 | 6,149616 | 5,090697 | 5,735782 | 5,471057 |
| | **Barrington's nucleus** | -64,29 | *0,3118* | 0,0284 | 0,0236 | 3 | 0,0101 | 0,0077 | 3 | | | | 0,009129 | 0,054771 | | 0,0213 | | 0,018257 | | | 0,003043 | 0,009129 | |
| | **Dorsal tegmental nucleus** | 4,35 | *0,8451* | 0,1565 | 0,0388 | 7 | 0,1633 | 0,0742 | 6 | 0,1917 | 0,1704 | 0,197786 | 0,146057 | 0,0852 | 0,133886 | 0,1704 | 0,209957 | | 0,179529 | 0,1704 | 0,197786 | 0,015214 | 0,206914 |
| | **Posterodorsal tegmental nucleus** | -18,09 | *0,5293* | 0,0771 | 0,0168 | 3 | 0,0631 | 0,0360 | 4 | | | | 0,057814 | 0,0852 | | 0,088243 | | 0,094329 | | | 0,033471 | 0,094329 | 0,030429 |
| | **Pontine central gray** | 4,14 | *0,7853* | 0,4621 | 0,0864 | 7 | 0,4812 | 0,1584 | 7 | 0,356014 | 0,356014 | 0,422957 | 0,505115 | 0,526414 | 0,581186 | 0,486857 | 0,374272 | 0,721157 | 0,4047 | 0,429043 | 0,3834 | 0,699857 | 0,356014 |
| | **Pontine gray** | 5,13 | *0,5844* | 1,00 | 0,13 | 7 | 1,05 | 0,20 | 7 | 0,961543 | 1,052829 | 1,010229 | 1,037615 | 0,912857 | 1,223229 | 0,803314 | 1,281044 | 0,879386 | 1,223228 | 0,970672 | 1,180628 | 0,7242 | 1,101514 |
| | **Pontine reticular nucleus caudal part** | -3,23 | *0,3701* | 2,37 | 0,11 | 7 | 2,29 | 0,18 | 7 | 2,464714 | 2,443416 | 2,458629 | 2,13913 | 2,361257 | 2,403858 | 2,327786 | 2,248673 | 2,495143 | 2,154342 | 2,510358 | 2,029585 | 2,416027 | 2,209114 |
| | **Supragenual nucleus** | 82,72 | *0,2273* | 0,0205 | 0,0046 | 4 | 0,0375 | 0,0173 | 3 | 0,018257 | 0,018257 | 0,027386 | | | 0,018257 | | | 0,051729 | | 0,018257 | | 0,0426 | |
| | **Supratrigeminal nucleus** | -3,21 | *0,8533* | 0,2573 | 0,0896 | 7 | 0,2491 | 0,0730 | 7 | 0,343843 | 0,249514 | 0,279943 | 0,188657 | 0,143014 | 0,395572 | 0,200829 | 0,231257 | 0,2982 | 0,173443 | 0,386443 | 0,200828 | 0,252557 | 0,200829 |
| | **Tegmental reticular nucleus** | -11,33 | *0,2154* | 0,7133 | 0,0813 | 7 | 0,6325 | 0,1394 | 7 | 0,803314 | 0,721158 | 0,751586 | 0,602486 | 0,6603 | 0,642043 | 0,812443 | 0,517286 | 0,797229 | 0,483814 | 0,7668 | 0,4686 | 0,718114 | 0,675514 |
| | **Motor nucleus of trigeminal** | -13,65 | *0,0628* | 0,3886 | 0,0516 | 7 | 0,3356 | 0,0448 | 7 | 0,398614 | 0,432086 | 0,398614 | 0,307329 | 0,371229 | 0,462514 | 0,349929 | 0,328629 | 0,371229 | 0,282986 | 0,4047 | 0,282986 | 0,352971 | 0,325586 |
| | **Peritrigeminal zone** | -10,20 | *0,4382* | 0,3282 | 0,0684 | 7 | 0,2947 | 0,0864 | 7 | 0,3834 | 0,313414 | 0,410786 | 0,249514 | 0,249514 | 0,395572 | 0,295157 | 0,249514 | 0,453386 | 0,252557 | 0,365143 | 0,200828 | 0,292114 | 0,249514 |
| | **Accessory trigeminal nucleus** | -36,72 | *0,3746* | 0,0325 | 0,0178 | 3 | 0,0205 | 0,0097 | 4 | | | | 0,012171 | 0,039557 | | 0,045643 | | 0,012171 | | | 0,012171 | 0,027386 | 0,030429 |
| | **Parvicellular motor 5 nucleus** | 6,06 | *0,8095* | 0,0717 | 0,0234 | 7 | 0,0761 | 0,0401 | 7 | 0,045643 | 0,088243 | 0,036514 | 0,100414 | 0,066943 | 0,079114 | 0,0852 | 0,118671 | 0,027386 | 0,124757 | 0,066943 | 0,094329 | 0,024343 | 0,076071 |

| | | | | | | | | | | | | | | | | | | | | | | | |
|---|---|---|---|---|---|---|---|---|---|---|---|---|---|---|---|---|---|---|---|---|---|---|---|
| | Intertrigeminal nucleus | -19,69 | *0,6007* | 0,0561 | 0,0313 | 7 | 0,0450 | 0,0367 | 5 | 0,082157 | 0,036514 | 0,0852 | 0,006086 | 0,0426 | 0,091286 | 0,048686 | | 0,0852 | | 0,060857 | 0,003043 | 0,066943 | 0,009129 |
| | Pons behavioral state related | -1,97 | *0,5262* | 3,56 | 0,19 | 7 | 3,49 | 0,21 | 7 | 3,374528 | 3,520588 | 3,441473 | 3,733588 | 3,508414 | 3,429301 | 3,916157 | 3,572316 | 3,243686 | 3,544927 | 3,447558 | 3,496241 | 3,27107 | 3,858343 |
| | Superior central nucleus raphe | -0,67 | *0,8846* | 0,6442 | 0,0602 | 7 | 0,6399 | 0,0488 | 7 | 0,690728 | 0,699858 | 0,696815 | 0,590315 | 0,547714 | 0,669429 | 0,614657 | 0,6816 | 0,666386 | 0,626828 | 0,690729 | 0,602485 | 0,5538 | 0,657257 |
| | Laterodorsal tegmental nucleus | 10,89 | *0,6459* | 0,2356 | 0,1041 | 7 | 0,2613 | 0,0995 | 7 | 0,304286 | 0,322543 | 0,267772 | 0,176486 | 0,045643 | 0,337757 | 0,194743 | 0,334714 | 0,069986 | 0,301243 | 0,325586 | 0,261686 | 0,1917 | 0,343843 |
| | Nucleus incertus | 8,32 | *0,6888* | 0,1652 | 0,0557 | 7 | 0,1789 | 0,0573 | 5 | 0,136929 | 0,176486 | 0,1704 | 0,222129 | 0,1917 | 0,054771 | 0,203871 | 0,188657 | | 0,194743 | 0,079114 | 0,209957 | | 0,222129 |
| | Pontine reticular nucleus | -0,97 | *0,6924* | 2,33 | 0,14 | 7 | 2,31 | 0,05 | 7 | 2,181728 | 2,24563 | 2,218244 | 2,422116 | 2,349086 | 2,306487 | 2,574257 | 2,251716 | 2,358215 | 2,285185 | 2,291272 | 2,236499 | 2,361256 | 2,355171 |
| | Nucleus raphe pontis | 4,91 | *0,7571* | 0,0974 | 0,0341 | 7 | 0,1022 | 0,0206 | 7 | 0,060857 | 0,076071 | 0,088243 | 0,143014 | 0,124757 | 0,060857 | 0,1278 | 0,103457 | 0,103457 | 0,121714 | 0,060857 | 0,094329 | 0,112586 | 0,118671 |
| * | Sublaterodorsal nucleus | -71,60 | *0,0019* | 0,1268 | 0,0183 | 3 | 0,0360 | 0,0378 | 6 | | | | 0,109543 | 0,146057 | | 0,124757 | 0,012171 | 0,006086 | 0,015214 | | 0,057814 | 0,0213 | 0,103457 |
| | Medulla | -3,24 | *0,2484* | 31,30 | 1,94 | 7 | 30,28 | 0,99 | 7 | 30,07863 | 30,43772 | 28,92237 | 31,39926 | 32,78374 | 30,69027 | 34,77073 | 30,76635 | 30,63853 | 29,29053 | 29,4001 | 30,32814 | 29,50353 | 32,05041 |
| | Medulla sensory related | -2,55 | *0,4989* | 7,89 | 0,63 | 7 | 7,69 | 0,43 | 7 | 7,378926 | 7,54629 | 7,308945 | 8,240062 | 8,337429 | 7,448917 | 8,946001 | 7,762333 | 7,531072 | 7,591926 | 7,135502 | 8,157896 | 7,305896 | 8,316129 |
| | Area postrema | -18,75 | *0,1144* | 0,0835 | 0,0187 | 7 | 0,0678 | 0,0155 | 7 | 0,097371 | 0,103457 | 0,094329 | 0,0639 | 0,054771 | 0,094329 | 0,076071 | 0,073029 | 0,073029 | 0,0639 | 0,091286 | 0,0639 | 0,069986 | 0,039557 |
| | Cochlear nuclei | -4,23 | *0,2921* | 1,66 | 0,13 | 7 | 1,59 | 0,10 | 7 | 1,789199 | 1,75573 | 1,710086 | 1,576201 | 1,430143 | 1,777029 | 1,603586 | 1,743558 | 1,710086 | 1,560985 | 1,570115 | 1,536642 | 1,591413 | 1,436229 |
| | Dorsal cochlear nucleus | -2,36 | *0,7969* | 0,6073 | 0,0931 | 7 | 0,5929 | 0,1101 | 7 | 0,675514 | 0,657258 | 0,669429 | 0,5112 | 0,474686 | 0,712029 | 0,550757 | 0,663343 | 0,687686 | 0,584228 | 0,645086 | 0,477728 | 0,684642 | 0,407743 |
| | Ventral cochlear nucleus | -5,31 | *0,1021* | 1,06 | 0,05 | 7 | 0,9998 | 0,0659 | 7 | 1,113685 | 1,098472 | 1,040657 | 1,065001 | 0,955457 | 1,065 | 1,052829 | 1,080215 | 1,0224 | 0,976757 | 0,925029 | 1,058914 | 0,906771 | 1,028486 |
| | Dorsal column nuclei | 14,32 | *0,1946* | 0,3430 | 0,0605 | 7 | 0,3921 | 0,0724 | 7 | 0,292114 | 0,295157 | 0,325586 | 0,322543 | 0,4047 | 0,310372 | 0,450343 | 0,3195 | 0,459471 | 0,3408 | 0,316457 | 0,371228 | 0,492943 | 0,444257 |
| | Cuneate nucleus | 3,09 | *0,6570* | 0,3095 | 0,0468 | 7 | 0,3191 | 0,0295 | 7 | 0,289071 | 0,289072 | 0,249514 | 0,289072 | 0,352971 | 0,307329 | 0,389486 | 0,286029 | 0,3195 | 0,301243 | 0,304286 | 0,313414 | 0,331671 | 0,377314 |
| | Gracile nucleus | 118,18 | *0,1346* | 0,0335 | 0,0303 | 7 | 0,0730 | 0,0562 | 7 | 0,003043 | 0,006086 | 0,076071 | 0,033471 | 0,051729 | 0,003043 | 0,060857 | 0,033471 | 0,139971 | 0,039557 | 0,012171 | 0,057814 | 0,161271 | 0,066943 |
| * | External cuneate nucleus | -9,47 | *0,0075* | 0,2386 | 0,0110 | 7 | 0,2160 | 0,0147 | 7 | 0,219086 | 0,240386 | 0,2343 | 0,237343 | 0,237343 | 0,252557 | 0,249514 | 0,237343 | 0,228214 | 0,200829 | 0,219086 | 0,194743 | 0,216043 | 0,216043 |
| | Nucleus of the trapezoid body | 9,88 | *0,5676* | 0,1408 | 0,0482 | 7 | 0,1548 | 0,0399 | 7 | 0,112586 | 0,097371 | 0,103457 | 0,209957 | 0,197786 | 0,103457 | 0,161271 | 0,130843 | 0,103457 | 0,173443 | 0,1065 | 0,1917 | 0,185614 | 0,1917 |
| | Nucleus of the solitary tract | -4,43 | *0,3664* | 0,9029 | 0,0930 | 7 | 0,8629 | 0,0625 | 7 | 0,821571 | 0,921986 | 0,815486 | 0,949372 | 0,955457 | 0,803315 | 1,052829 | 0,867215 | 0,833743 | 0,8946 | 0,852 | 0,858085 | 0,763757 | 0,970671 |
| | Spinal nucleus of the trigeminal caudal part | -0,74 | *0,9433* | 1,59 | 0,34 | 7 | 1,58 | 0,26 | 7 | 1,347985 | 1,363201 | 1,314515 | 1,704001 | 1,962643 | 1,332772 | 2,126957 | 1,536644 | 1,3632 | 1,524471 | 1,366243 | 1,920042 | 1,390585 | 1,968729 |
| | Spinal nucleus of the trigeminal interpolar part | -8,37 | *0,0600* | 1,95 | 0,10 | 7 | 1,79 | 0,18 | 7 | 2,014371 | 2,072187 | 1,917001 | 1,938301 | 1,840929 | 2,063058 | 1,816586 | 1,956558 | 1,834843 | 1,868314 | 1,907872 | 1,789199 | 1,427099 | 1,734429 |
| | Spinal nucleus of the trigeminal oral part | 8,23 | *0,6182* | 0,8607 | 0,2983 | 7 | 0,9315 | 0,2109 | 7 | 0,602486 | 0,584229 | 0,712029 | 1,101515 | 1,138029 | 0,611615 | 1,274957 | 0,760715 | 0,864172 | 0,861128 | 0,623786 | 1,101514 | 1,104557 | 1,204971 |
| | Paratrigeminal nucleus | -9,96 | *0,4615* | 0,1091 | 0,0223 | 7 | 0,0982 | 0,0304 | 7 | 0,082157 | 0,112586 | 0,082157 | 0,136929 | 0,115629 | 0,100414 | 0,133886 | 0,136929 | 0,060857 | 0,103457 | 0,082157 | 0,130843 | 0,0639 | 0,109543 |
| | Medulla motor related | -3,44 | *0,2370* | 17,69 | 1,13 | 7 | 17,08 | 0,57 | 7 | 16,98522 | 17,28648 | 16,17583 | 17,60598 | 18,44276 | 17,59685 | 19,72684 | 17,2165 | 17,38689 | 16,50749 | 16,94568 | 16,80569 | 16,544 | 18,15673 |
| | Facial motor nucleus | -9,66 | *0,2978* | 0,9359 | 0,1607 | 7 | 0,8455 | 0,1499 | 7 | 1,065 | 0,928072 | 1,037615 | 0,678558 | 0,772886 | 0,946329 | 1,122814 | 0,684643 | 1,061957 | 0,6816 | 0,852 | 0,736371 | 0,973714 | 0,928071 |
| | Nucleus ambiguus | -11,01 | *0,3292* | 0,0474 | 0,0065 | 7 | 0,0422 | 0,0117 | 7 | 0,045643 | 0,054771 | 0,036514 | 0,048686 | 0,0426 | 0,054771 | 0,048686 | 0,027386 | 0,060857 | 0,039557 | 0,051729 | 0,030429 | 0,039557 | 0,045643 |
| | Nucleus ambiguus dorsal division | -16,95 | *0,2096* | 0,0256 | 0,0030 | 7 | 0,0213 | 0,0079 | 7 | 0,027386 | 0,0213 | 0,024343 | 0,024343 | 0,024343 | 0,027386 | 0,030429 | 0,009129 | 0,033471 | 0,015214 | 0,024343 | 0,018257 | 0,024343 | 0,024343 |
| | Nucleus ambiguus ventral division | -4,00 | *0,8084* | 0,0217 | 0,0071 | 7 | 0,0209 | 0,0059 | 7 | 0,018257 | 0,033471 | 0,012171 | 0,024343 | 0,018257 | 0,027386 | 0,018257 | 0,018257 | 0,027386 | 0,024343 | 0,027386 | 0,012171 | 0,015214 | 0,0213 |
| | Dorsal motor nucleus of the vagus nerve | 0,48 | *0,9197* | 0,1826 | 0,0169 | 7 | 0,1834 | 0,0147 | 7 | 0,185614 | 0,188657 | 0,1491 | 0,176486 | 0,188657 | 0,185614 | 0,203871 | 0,206914 | 0,185614 | 0,167357 | 0,1704 | 0,182571 | 0,173443 | 0,197786 |
| | Gigantocellular reticular nucleus | -3,59 | *0,3006* | 2,60 | 0,12 | 7 | 2,51 | 0,19 | 7 | 2,635113 | 2,616859 | 2,443415 | 2,461673 | 2,674672 | 2,598601 | 2,7903 | 2,388644 | 2,723358 | 2,239542 | 2,543829 | 2,327785 | 2,647284 | 2,695971 |
| | Infracerebellar nucleus | -19,82 | *0,2203* | 0,0786 | 0,0234 | 6 | 0,0630 | 0,0187 | 7 | 0,100414 | 0,079114 | 0,097371 | 0,0639 | | 0,091286 | 0,039557 | 0,079114 | 0,054771 | 0,076071 | 0,088243 | 0,0426 | 0,039557 | 0,060857 |
| | Inferior olivary complex | -2,10 | *0,7103* | 0,5586 | 0,0620 | 7 | 0,5468 | 0,0530 | 7 | 0,559886 | 0,4899 | 0,474686 | 0,5964 | 0,654214 | 0,550757 | 0,584229 | 0,587272 | 0,486857 | 0,5538 | 0,550757 | 0,632914 | 0,483814 | 0,5325 |
| * | Intermediate reticular nucleus | -5,06 | *0,0498* | 2,86 | 0,13 | 7 | 2,72 | 0,12 | 7 | 2,790299 | 2,924187 | 2,677715 | 2,765959 | 2,887672 | 2,921144 | 3,0672 | 2,674673 | 2,8329 | 2,674671 | 2,86333 | 2,54687 | 2,616856 | 2,8116 |
| | Inferior salivatory nucleus | 8,24 | *0,6575* | 0,0129 | 0,0038 | 4 | 0,0140 | 0,0027 | 5 | | 0,012171 | | 0,018257 | 0,012171 | | 0,009129 | 0,015214 | | 0,018257 | 0,012171 | 0,012171 | | 0,012171 |
| | Linear nucleus of the medulla | 16,67 | *0,6780* | 0,0657 | 0,0457 | 5 | 0,0767 | 0,0336 | 5 | | 0,036514 | | 0,115629 | 0,115629 | 0,027386 | 0,033471 | 0,097371 | | 0,091286 | 0,045643 | 0,112586 | | 0,036514 |
| | Lateral reticular nucleus | -6,38 | *0,4833* | 0,5999 | 0,0842 | 7 | 0,5616 | 0,1113 | 7 | 0,623786 | 0,550757 | 0,642043 | 0,465557 | 0,593357 | 0,584229 | 0,739414 | 0,480772 | 0,584229 | 0,441214 | 0,505114 | 0,495985 | 0,708985 | 0,715071 |
| | Lateral reticular nucleus magnocellular part | -7,30 | *0,5047* | 0,5416 | 0,0920 | 7 | 0,5021 | 0,1208 | 7 | 0,572057 | 0,483815 | 0,5964 | 0,401657 | 0,526414 | 0,517286 | 0,693771 | 0,4047 | 0,514243 | 0,371228 | 0,432086 | 0,459471 | 0,6603 | 0,672471 |
| | Lateral reticular nucleus parvicellular part | 2,24 | *0,8615* | 0,0582 | 0,0102 | 7 | 0,0596 | 0,0164 | 7 | 0,051729 | 0,066943 | 0,045643 | 0,0639 | 0,066943 | 0,066943 | 0,045643 | 0,076071 | 0,069986 | 0,069986 | 0,073029 | 0,036514 | 0,048686 | 0,0426 |
| | Magnocellular reticular nucleus | -3,12 | *0,6484* | 0,5021 | 0,0548 | 7 | 0,4864 | 0,0694 | 7 | 0,5112 | 0,550757 | 0,483814 | 0,505115 | 0,544671 | 0,529457 | 0,389486 | 0,547715 | 0,502071 | 0,5112 | 0,562929 | 0,495985 | 0,410785 | 0,374271 |
| | Medullary reticular nucleus | 4,10 | *0,7190* | 1,97 | 0,45 | 7 | 2,05 | 0,37 | 7 | 1,646185 | 1,676615 | 1,518386 | 2,279101 | 2,443414 | 1,652272 | 2,586429 | 2,130001 | 1,685743 | 1,959599 | 1,761815 | 2,513399 | 1,73747 | 2,580343 |
| | Medullary reticular nucleus dorsal part | 4,46 | *0,7114* | 1,02 | 0,22 | 7 | 1,07 | 0,23 | 7 | 0,842871 | 0,864172 | 0,8307 | 1,083258 | 1,250614 | 0,903729 | 1,3845 | 0,958501 | 0,8946 | 0,9585 | 0,876343 | 1,250614 | 1,037614 | 1,503171 |
| | Medullary reticular nucleus ventral part | 3,71 | *0,7701* | 0,9489 | 0,2355 | 7 | 0,9841 | 0,2039 | 7 | 0,803314 | 0,812443 | 0,687686 | 1,195844 | 1,1928 | 0,748543 | 1,201929 | 1,171501 | 0,791143 | 1,0011 | 0,885472 | 1,262785 | 0,699857 | 1,077171 |
| | Parvicellular reticular nucleus | -5,93 | *0,0734* | 2,33 | 0,14 | 7 | 2,19 | 0,12 | 7 | 2,300399 | 2,452544 | 2,172601 | 2,251716 | 2,169557 | 2,513401 | 2,452543 | 2,276058 | 2,376472 | 2,203028 | 2,203029 | 2,160428 | 1,990027 | 2,136086 |
| | Parasolitary nucleus | 45,83 | *0,4649* | 0,0183 | 0,0157 | 5 | 0,0266 | 0,0164 | 4 | | | 0,006086 | 0,012171 | 0,036514 | 0,003043 | 0,033471 | | 0,006086 | | | 0,030429 | 0,024343 | 0,045643 |
| | Paragigantocellular reticular nucleus | -2,87 | *0,6670* | 0,9394 | 0,1074 | 7 | 0,9124 | 0,1207 | 7 | 0,812443 | 0,921986 | 0,785057 | 1,071086 | 1,034571 | 0,970672 | 0,9798 | 1,071086 | 0,7242 | 1,031528 | 0,891557 | 0,891557 | 0,815485 | 0,961543 |
| | Paragigantocellular reticular nucleus dorsal part | -12,36 | *0,1808* | 0,2708 | 0,0230 | 7 | 0,2373 | 0,0558 | 7 | 0,267771 | 0,295157 | 0,231257 | 0,258643 | 0,264729 | 0,279943 | 0,2982 | 0,2556 | 0,130843 | 0,270814 | 0,273857 | 0,240386 | 0,197786 | 0,292114 |
| | Paragigantocellular reticular nucleus lateral part | 0,98 | *0,8974* | 0,6686 | 0,1016 | 7 | 0,6751 | 0,0826 | 7 | 0,544671 | 0,626829 | 0,5538 | 0,812443 | 0,769843 | 0,690729 | 0,6816 | 0,815486 | 0,593357 | 0,760714 | 0,6177 | 0,651171 | 0,6177 | 0,669429 |
| | Perihypoglossal nuclei | 14,85 | *0,2970* | 0,2517 | 0,0746 | 7 | 0,2891 | 0,0507 | 7 | 0,143014 | 0,194743 | 0,252557 | 0,334714 | 0,313414 | 0,200829 | 0,322543 | 0,334714 | 0,289071 | 0,328628 | 0,182571 | 0,282986 | 0,301243 | 0,304286 |
| | Nucleus of Roller | -17,98 | *0,2982* | 0,0387 | 0,0109 | 7 | 0,0317 | 0,0129 | 7 | 0,027386 | 0,0426 | 0,048686 | 0,048686 | 0,024343 | 0,048686 | 0,030429 | 0,0426 | 0,036514 | 0,0426 | 0,0426 | 0,024343 | 0,009129 | 0,024343 |
| | Nucleus prepositus | 20,82 | *0,2336* | 0,2130 | 0,0757 | 7 | 0,2573 | 0,0541 | 7 | 0,115629 | 0,152143 | 0,203871 | 0,286029 | 0,289071 | 0,152143 | 0,292114 | 0,292114 | 0,252557 | 0,286028 | 0,139971 | 0,258643 | 0,292114 | 0,279943 |
| | Parapyramidal nucleus | -7,52 | *0,6875* | 0,0982 | 0,0325 | 7 | 0,0909 | 0,0346 | 7 | 0,1278 | 0,112586 | 0,115629 | 0,057814 | 0,0639 | 0,073029 | 0,136929 | 0,054771 | 0,143014 | 0,0639 | 0,091286 | 0,054771 | 0,109543 | 0,118671 |
| | Vestibular nuclei | -4,25 | *0,2109* | 3,22 | 0,22 | 7 | 3,09 | 0,15 | 7 | 3,027642 | 3,05503 | 2,966787 | 3,264988 | 3,347143 | 3,301501 | 3,605786 | 3,121973 | 3,310629 | 2,966785 | 3,09763 | 2,82377 | 3,128055 | 3,161529 |
| | Lateral vestibular nucleus | 17,98 | *0,2335* | 0,2708 | 0,0584 | 7 | 0,3195 | 0,0837 | 7 | 0,279943 | 0,346886 | 0,264729 | 0,322543 | 0,167357 | 0,2769 | 0,237343 | 0,453386 | 0,279943 | 0,422957 | 0,2982 | 0,286028 | 0,249514 | 0,246471 |
| | Medial vestibular nucleus | -6,20 | *0,2187* | 1,85 | 0,19 | 7 | 1,74 | 0,13 | 7 | 1,673571 | 1,673572 | 1,700958 | 1,837887 | 2,081314 | 1,840929 | 2,148257 | 1,704001 | 1,941343 | 1,597499 | 1,777029 | 1,573156 | 1,798328 | 1,761814 |
| | Spinal vestibular nucleus | -5,26 | *0,2897* | 0,7433 | 0,0588 | 7 | 0,7042 | 0,0724 | 7 | 0,712028 | 0,693772 | 0,669429 | 0,748543 | 0,757671 | 0,775929 | 0,845914 | 0,651172 | 0,800272 | 0,6177 | 0,708986 | 0,629871 | 0,763757 | 0,757671 |
| | Superior vestibular nucleus | -8,84 | *0,0718* | 0,3591 | 0,0259 | 7 | 0,3273 | 0,0334 | 7 | 0,3621 | 0,3408 | 0,331672 | 0,356014 | 0,3408 | 0,407743 | 0,374271 | 0,313414 | 0,289071 | 0,328628 | 0,313414 | 0,334714 | 0,316457 | 0,395571 |
| | Nucleus x | -17,14 | *0,1440* | 0,0609 | 0,0118 | 7 | 0,0504 | 0,0131 | 7 | 0,054771 | 0,054771 | 0,045643 | 0,0639 | 0,073029 | 0,054771 | 0,079114 | 0,048686 | 0,030429 | 0,045643 | 0,060857 | 0,060857 | 0,039557 | 0,066943 |
| | Hypoglossal nucleus | -3,03 | *0,7687* | 0,3443 | 0,0452 | 7 | 0,3338 | 0,0792 | 7 | 0,337757 | 0,374272 | 0,2556 | 0,349929 | 0,380357 | 0,325586 | 0,386443 | 0,389486 | 0,246471 | 0,410786 | 0,395572 | 0,3408 | 0,203871 | 0,349929 |
| | Nucleus y | -51,67 | *0,0931* | 0,0365 | 0,0122 | 3 | 0,0176 | 0,0090 | 5 | | | | 0,024343 | 0,036514 | | 0,048686 | | 0,018257 | 0,003043 | | 0,027386 | 0,018257 | 0,0213 |
| | Medulla behavioral state related | -8,27 | *0,1174* | 0,2208 | 0,0153 | 7 | 0,2026 | 0,0238 | 7 | 0,2343 | 0,225172 | 0,219086 | 0,188657 | 0,219086 | 0,228214 | 0,231257 | 0,203872 | 0,246471 | 0,194743 | 0,216043 | 0,176486 | 0,200828 | 0,179529 |
| | Nucleus raphe magnus | -6,50 | *0,5420* | 0,1069 | 0,0185 | 7 | 0,1000 | 0,0227 | 7 | 0,115629 | 0,1278 | 0,112586 | 0,091286 | 0,0852 | 0,1278 | 0,088243 | 0,103457 | 0,1278 | 0,109543 | 0,124757 | 0,076071 | 0,088243 | 0,069986 |
| | Nucleus raphe pallidus | -3,73 | *0,6624* | 0,0700 | 0,0138 | 7 | 0,0674 | 0,0064 | 7 | 0,066943 | 0,060857 | 0,057814 | 0,0639 | 0,091286 | 0,060857 | 0,088243 | 0,073029 | 0,066943 | 0,0639 | 0,054771 | 0,073029 | 0,069986 | 0,069986 |
| | Nucleus raphe obscurus | -19,80 | *0,1094* | 0,0439 | 0,0080 | 7 | 0,0352 | 0,0105 | 7 | 0,051729 | 0,036514 | 0,048686 | 0,033471 | 0,0426 | 0,039557 | 0,054771 | 0,027386 | 0,051729 | 0,0213 | 0,036514 | 0,027386 | 0,0426 | 0,039557 |
| | Cerebellum | -3,98 | *0,1004* | 49,93 | 2,22 | 7 | 47,94 | 1,94 | 7 | 47,80023 | 48,49708 | 47,72723 | 51,0896 | 50,73356 | 49,69901 | 53,93769 | 49,33387 | 47,62681 | 48,12277 | 46,98477 | 44,68129 | 47,88542 | 50,94352 |
| | Cerebellar cortex | -3,60 | *0,1195* | 47,35 | 1,95 | 7 | 45,64 | 1,86 | 7 | 45,51504 | 46,09018 | 45,50899 | 48,27496 | 48,02846 | 47,04868 | 50,95265 | 46,99695 | 45,32641 | 45,83758 | 44,7361 | 42,45088 | 45,70065 | 48,43012 |
| * | Vermal regions | -5,08 | *0,0353* | 19,72 | 0,42 | 7 | 18,71 | 0,97 | 7 | 19,50471 | 19,43778 | 19,58688 | 19,58688 | 19,4895 | 19,75424 | 20,64883 | 19,15175 | 18,84137 | 18,87484 | 18,75922 | 16,64442 | 19,0148 | 19,70554 |
| | Lingula (I) | -11,25 | *0,5351* | 0,1430 | 0,0272 | 7 | 0,1269 | 0,0599 | 7 | 0,143014 | 0,115629 | 0,167357 | 0,112586 | 0,136929 | 0,188657 | 0,136929 | 0,197786 | 0,051729 | 0,158229 | 0,179529 | 0,130843 | 0,0426 | 0,1278 |
| | Central lobule | -3,42 | *0,3061* | 3,88 | 0,18 | 7 | 3,75 | 0,27 | 7 | 3,773142 | 3,739673 | 3,703158 | 3,943545 | 4,1322 | 3,748802 | 4,113943 | 3,903988 | 3,462772 | 3,767056 | 3,709244 | 3,636212 | 3,496241 | 4,250871 |

| | | | | | | | | | | | | | | | | | | | | | | | |
|---|---|---|---|---|---|---|---|---|---|---|---|---|---|---|---|---|---|---|---|---|---|---|---|
| | Lobule II | 4,27 | *0,3351* | 1,24 | 0,10 | 7 | 1,30 | 0,09 | 7 | 1,2141 | 1,256701 | 1,089343 | 1,299301 | 1,259743 | 1,414929 | 1,168457 | 1,302344 | 1,229314 | 1,2993 | 1,281043 | 1,162371 | 1,329728 | 1,4697 |
| | Lobule III | -7,04 | *0,1257* | 2,64 | 0,21 | 7 | 2,45 | 0,21 | 7 | 2,559042 | 2,482973 | 2,613815 | 2,644244 | 2,872457 | 2,333872 | 2,945486 | 2,601644 | 2,233457 | 2,467756 | 2,428201 | 2,473842 | 2,166513 | 2,781171 |
| * | Culmen | -7,06 | *0,0424* | 6,12 | 0,42 | 7 | 5,69 | 0,25 | 7 | 6,332184 | 5,991389 | 5,897059 | 6,110061 | 6,651686 | 5,388902 | 6,499543 | 5,671889 | 5,431501 | 6,009641 | 5,875759 | 5,367597 | 5,589725 | 5,9001 |
| * | Lobules IV-V | -7,06 | *0,0424* | 6,12 | 0,42 | 7 | 5,69 | 0,25 | 7 | 6,332184 | 5,991389 | 5,897059 | 6,110061 | 6,651686 | 5,388902 | 6,499543 | 5,671889 | 5,431501 | 6,009641 | 5,875759 | 5,367597 | 5,589725 | 5,9001 |
| | Declive (VI) | 4,42 | *0,5410* | 2,85 | 0,30 | 7 | 2,98 | 0,43 | 7 | 2,379514 | 2,933316 | 2,702058 | 2,71423 | 2,784214 | 3,286287 | 3,149357 | 2,985044 | 3,490158 | 2,519485 | 2,647287 | 2,492099 | 3,468855 | 3,228471 |
| | Folium-tuber vermis (VII) | -9,40 | *0,1803* | 1,19 | 0,10 | 7 | 1,08 | 0,18 | 7 | 1,186714 | 1,089343 | 1,189758 | 1,247572 | 1,162371 | 1,068043 | 1,378414 | 1,001101 | 0,9159 | 1,204971 | 1,208015 | 0,943285 | 0,912857 | 1,354071 |
| | Pyramus (VIII) | -14,28 | *0,1894* | 1,54 | 0,27 | 7 | 1,32 | 0,32 | 7 | 1,691828 | 1,509258 | 1,728343 | 1,679658 | 0,961543 | 1,707044 | 1,506214 | 1,688787 | 1,204972 | 1,560985 | 1,573158 | 0,794185 | 1,338856 | 1,083257 |
| | Uvula (IX) | -2,21 | *0,8265* | 2,21 | 0,30 | 7 | 2,16 | 0,49 | 7 | 2,239542 | 2,339958 | 2,479929 | 1,913958 | 1,886571 | 2,647287 | 1,932214 | 1,871358 | 2,884629 | 1,859185 | 2,011329 | 1,746599 | 2,832898 | 1,892657 |
| | Nodulus (X) | -9,38 | *0,0934* | 1,78 | 0,08 | 7 | 1,62 | 0,22 | 7 | 1,758771 | 1,719215 | 1,719215 | 1,865273 | 1,773986 | 1,719215 | 1,932214 | 1,831801 | 1,399714 | 1,795285 | 1,554901 | 1,533599 | 1,332771 | 1,868314 |
| | Hemispheric regions | -2,55 | *0,3555* | 27,63 | 1,62 | 7 | 26,93 | 1,04 | 7 | 26,01033 | 26,6524 | 25,92211 | 28,68807 | 28,53896 | 27,29444 | 30,30382 | 27,8452 | 26,48503 | 26,96275 | 25,97688 | 25,80646 | 26,68584 | 28,72457 |
| | Simple lobule | -1,30 | *0,6701* | 5,10 | 0,29 | 7 | 5,04 | 0,28 | 7 | 5,057227 | 4,950731 | 4,755987 | 5,181989 | 5,057229 | 5,032888 | 5,693186 | 5,282403 | 4,807715 | 5,008541 | 5,060273 | 4,612969 | 5,038969 | 5,4528 |
| | Ansiform lobule | -2,22 | *0,4794* | 9,44 | 0,67 | 7 | 9,23 | 0,35 | 7 | 8,976426 | 9,076848 | 8,60216 | 9,767577 | 10,05664 | 9,168132 | 10,46439 | 9,505891 | 8,979473 | 9,210725 | 9,274632 | 8,76951 | 9,067709 | 9,834514 |
| | Crus 1 | -1,35 | *0,6840* | 5,12 | 0,35 | 7 | 5,05 | 0,26 | 7 | 4,929427 | 4,984203 | 4,564287 | 5,25806 | 5,270229 | 5,151559 | 5,693186 | 5,297617 | 4,731644 | 5,111998 | 5,154602 | 4,689041 | 5,00854 | 5,373686 |
| | Crus 2 | -3,26 | *0,3580* | 4,32 | 0,35 | 7 | 4,18 | 0,14 | 7 | 4,046999 | 4,092645 | 4,037873 | 4,509517 | 4,786414 | 4,016573 | 4,7712 | 4,208274 | 4,247829 | 4,098727 | 4,12003 | 4,080469 | 4,059169 | 4,460829 |
| | Paramedian lobule | -2,45 | *0,5061* | 4,28 | 0,34 | 7 | 4,18 | 0,22 | 7 | 3,806613 | 4,040916 | 4,016573 | 4,591674 | 4,530814 | 4,29043 | 4,704257 | 4,253917 | 4,326943 | 4,183927 | 3,77923 | 4,025698 | 4,244783 | 4,433443 |
| | Copula pyramidis | -1,18 | *0,7308* | 2,29 | 0,14 | 7 | 2,26 | 0,14 | 7 | 2,081314 | 2,218244 | 2,227372 | 2,263887 | 2,288229 | 2,498187 | 2,443414 | 2,248673 | 2,403857 | 2,154342 | 2,126958 | 2,081313 | 2,403856 | 2,412986 |
| | Paraflocculus | -4,25 | *0,2247* | 5,14 | 0,33 | 7 | 4,93 | 0,30 | 7 | 4,704256 | 5,032888 | 4,938559 | 5,574518 | 5,2611 | 4,929431 | 5,565386 | 5,191117 | 4,634272 | 5,157641 | 4,561244 | 5,023755 | 4,637312 | 5,270229 |
| * | Flocculus | -5,60 | *0,0211* | 1,37 | 0,04 | 7 | 1,29 | 0,06 | 7 | 1,3845 | 1,332772 | 1,381458 | 1,308429 | 1,344943 | 1,375372 | 1,433186 | 1,363201 | 1,332772 | 1,247571 | 1,174543 | 1,293214 | 1,293214 | 1,3206 |
| * | Cerebellar nuclei | -9,90 | *0,0230* | 1,76 | 0,15 | 7 | 1,59 | 0,09 | 7 | 1,637057 | 1,643144 | 1,551858 | 1,856144 | 1,862229 | 1,850058 | 1,929172 | 1,643144 | 1,491 | 1,643142 | 1,609672 | 1,545771 | 1,463613 | 1,713129 |
| | Fastigial nucleus | -8,12 | *0,3178* | 0,5299 | 0,0740 | 7 | 0,4869 | 0,0803 | 7 | 0,584228 | 0,535543 | 0,562929 | 0,520329 | 0,395571 | 0,623786 | 0,486857 | 0,569015 | 0,4473 | 0,5538 | 0,569014 | 0,356014 | 0,465557 | 0,4473 |
| | Interposed nucleus | -11,02 | *0,2075* | 0,8563 | 0,1414 | 7 | 0,7620 | 0,1226 | 7 | 0,763757 | 0,775929 | 0,705943 | 0,967629 | 1,010229 | 0,736372 | 1,034571 | 0,754629 | 0,648129 | 0,803314 | 0,712029 | 0,879385 | 0,5964 | 0,940243 |
| | Dentate nucleus | -8,53 | *0,1991* | 0,3160 | 0,0471 | 7 | 0,2891 | 0,0194 | 7 | 0,270814 | 0,279943 | 0,252557 | 0,331672 | 0,349929 | 0,3621 | 0,365143 | 0,2769 | 0,2982 | 0,286028 | 0,264729 | 0,279943 | 0,292114 | 0,325586 |
| | Vestibulocerebellar nucleus | 16,32 | *0,6686* | 0,0591 | 0,0414 | 7 | 0,0688 | 0,0341 | 5 | 0,018257 | 0,051729 | 0,030429 | 0,036514 | 0,1065 | 0,1278 | 0,0426 | 0,0426 | 0,097371 | | 0,0639 | 0,030429 | 0,109543 | |
| | fiber tracts | -3,45 | *0,1460* | 45,46 | 2,23 | 7 | 43,89 | 1,40 | 7 | 44,27356 | 44,32228 | 43,11121 | 45,32034 | 44,45006 | 47,03042 | 49,7142 | 44,18535 | 44,18838 | 42,31091 | 43,43984 | 42,67909 | 43,8232 | 46,61962 |
| | cranial nerves | -3,42 | *0,2502* | 12,21 | 0,73 | 7 | 11,80 | 0,54 | 7 | 11,76064 | 11,29813 | 11,57808 | 12,44529 | 12,42703 | 12,50919 | 13,47377 | 11,69675 | 12,09536 | 11,2251 | 11,24336 | 11,78194 | 11,72108 | 12,80434 |
| | vomeronasal nerve | 23,53 | *0,5085* | 0,0172 | 0,0115 | 6 | 0,0213 | 0,0070 | 4 | 0,0213 | 0,024343 | 0,003043 | 0,006086 | 0,033471 | 0,015214 | | 0,015214 | | | 0,027386 | 0,027386 | | 0,015214 |
| | olfactory nerve | -1,95 | *0,5235* | 5,68 | 0,39 | 7 | 5,57 | 0,20 | 7 | 5,553213 | 5,127217 | 5,385859 | 5,766218 | 5,614072 | 6,037031 | 6,295672 | 5,617117 | 5,738829 | 5,291527 | 5,35543 | 5,614069 | 5,522783 | 5,863586 |
| | olfactory nerve layer of main olfactory bulb | -1,26 | *0,7575* | 3,89 | 0,35 | 7 | 3,84 | 0,21 | 7 | 3,706199 | 3,33193 | 3,654473 | 4,092645 | 3,9831 | 4,101773 | 4,375629 | 3,885731 | 4,050043 | 3,627084 | 3,617958 | 3,757927 | 3,773141 | 4,190014 |
| | lateral olfactory tract general | -4,60 | *0,1024* | 1,04 | 0,05 | 7 | 0,9911 | 0,0550 | 7 | 1,031528 | 1,016315 | 0,995015 | 1,068043 | 0,9798 | 1,098472 | 1,083257 | 0,964586 | 0,988929 | 0,931114 | 0,940243 | 1,083257 | 1,043699 | 0,985886 |
| | lateral olfactory tract body | -5,51 | *0,0579* | 0,9224 | 0,0291 | 7 | 0,8716 | 0,0548 | 7 | 0,912857 | 0,928072 | 0,882429 | 0,937201 | 0,888514 | 0,952415 | 0,955457 | 0,839829 | 0,842872 | 0,806357 | 0,836786 | 0,961542 | 0,915899 | 0,897643 |
| | dorsal limb | 2,61 | *0,7793* | 0,1165 | 0,0211 | 7 | 0,1195 | 0,0186 | 7 | 0,118671 | 0,088243 | 0,112586 | 0,130843 | 0,091286 | 0,146057 | 0,1278 | 0,124757 | 0,146057 | 0,124757 | 0,103457 | 0,121714 | 0,1278 | 0,088243 |
| | anterior commissure olfactory limb | -1,85 | *0,7254* | 0,7516 | 0,0922 | 7 | 0,7377 | 0,0420 | 7 | 0,815485 | 0,778972 | 0,736372 | 0,605529 | 0,651171 | 0,836786 | 0,836786 | 0,7668 | 0,699857 | 0,733328 | 0,797229 | 0,772885 | 0,705942 | 0,687686 |
| | optic nerve | -6,08 | *0,0546* | 1,49 | 0,08 | 7 | 1,40 | 0,08 | 7 | 1,503171 | 1,481872 | 1,369286 | 1,457529 | 1,481871 | 1,527515 | 1,637057 | 1,436229 | 1,372329 | 1,4484 | 1,417972 | 1,375371 | 1,259742 | 1,5123 |
| | brachium of the superior colliculus | 6,46 | *0,1392* | 0,1817 | 0,0163 | 7 | 0,1934 | 0,0105 | 7 | 0,173443 | 0,206914 | 0,179529 | 0,161272 | 0,173443 | 0,176486 | 0,200829 | 0,185614 | 0,206914 | 0,185614 | 0,200829 | 0,179528 | 0,1917 | 0,203871 |
| | optic chiasm | -6,84 | *0,3422* | 0,3747 | 0,0478 | 7 | 0,3491 | 0,0492 | 7 | 0,401657 | 0,3621 | 0,389486 | 0,337757 | 0,2982 | 0,386443 | 0,4473 | 0,337757 | 0,4047 | 0,352971 | 0,3621 | 0,322543 | 0,261686 | 0,401657 |
| | optic tract | -2,91 | *0,3157* | 0,8064 | 0,0546 | 7 | 0,7829 | 0,0186 | 7 | 0,7668 | 0,7881 | 0,721157 | 0,791143 | 0,867214 | 0,848958 | 0,861129 | 0,800272 | 0,757672 | 0,797228 | 0,797229 | 0,778971 | 0,757671 | 0,791143 |
| | oculomotor nerve | -7,69 | *0,4089* | 0,1978 | 0,0308 | 7 | 0,1826 | 0,0355 | 7 | 0,213 | 0,1704 | 0,219086 | 0,197786 | 0,231257 | 0,209957 | 0,143014 | 0,1491 | 0,231257 | 0,143014 | 0,182571 | 0,152143 | 0,216043 | 0,203871 |
| | medial longitudinal fascicle | -0,60 | *0,8831* | 0,1448 | 0,0127 | 7 | 0,1439 | 0,0086 | 7 | 0,146057 | 0,152143 | 0,143014 | 0,130843 | 0,167357 | 0,143014 | 0,130843 | 0,139972 | 0,161271 | 0,139971 | 0,146057 | 0,143014 | 0,143014 | 0,133886 |
| | posterior commissure | -32,38 | *0,3521* | 0,0456 | 0,0251 | 7 | 0,0309 | 0,0315 | 7 | 0,045643 | 0,009129 | 0,066943 | 0,066943 | 0,0639 | 0,054771 | 0,012171 | 0,009129 | 0,051729 | 0,003043 | 0,006086 | 0,006086 | 0,069986 | 0,069986 |
| | trochlear nerve | -36,36 | *0,1798* | 0,0096 | 0,0054 | 7 | 0,0061 | 0,0025 | 4 | 0,015214 | 0,006086 | 0,006086 | 0,009129 | 0,009129 | 0,003043 | 0,018257 | | 0,006086 | 0,009129 | | 0,006086 | 0,003043 | |
| | trigeminal nerve | -4,22 | *0,1575* | 2,42 | 0,10 | 7 | 2,32 | 0,15 | 7 | 2,349085 | 2,339958 | 2,315615 | 2,473844 | 2,476886 | 2,379515 | 2,601643 | 2,291273 | 2,443415 | 2,084356 | 2,166515 | 2,406899 | 2,358213 | 2,4708 |
| | motor root of the trigeminal nerve | 14,29 | *0,4425* | 0,0609 | 0,0163 | 7 | 0,0696 | 0,0238 | 7 | 0,0639 | 0,079114 | 0,060857 | 0,0639 | 0,039557 | 0,079114 | 0,039557 | 0,103457 | 0,054771 | 0,100414 | 0,057814 | 0,066943 | 0,039557 | 0,0639 |
| | sensory root of the trigeminal nerve | -4,70 | *0,1629* | 2,36 | 0,12 | 7 | 2,25 | 0,16 | 7 | 2,285185 | 2,260844 | 2,254758 | 2,409944 | 2,437329 | 2,300401 | 2,562086 | 2,187815 | 2,388643 | 1,983942 | 2,108701 | 2,339956 | 2,318656 | 2,4069 |
| | spinal tract of the trigeminal nerve | -2,63 | *0,4544* | 1,62 | 0,12 | 7 | 1,57 | 0,09 | 7 | 1,494042 | 1,731387 | 1,421015 | 1,640101 | 1,6401 | 1,710086 | 1,685743 | 1,716172 | 1,5762 | 1,506214 | 1,509258 | 1,630971 | 1,466656 | 1,6188 |
| | facial nerve | -27,88 | *0,0545* | 0,1356 | 0,0310 | 7 | 0,0978 | 0,0352 | 7 | 0,124757 | 0,115629 | 0,118671 | 0,094329 | 0,1704 | 0,146057 | 0,179529 | 0,0426 | 0,088243 | 0,060857 | 0,133886 | 0,115629 | 0,112586 | 0,130843 |
| | genu of the facial nerve | 11,86 | *0,8280* | 0,0299 | 0,0216 | 6 | 0,0335 | 0,0292 | 5 | 0,0426 | | 0,057814 | 0,003043 | 0,015214 | 0,045643 | 0,015214 | | 0,060857 | | 0,027386 | 0,006086 | 0,066943 | 0,006086 |
| | vestibulocochlear nerve | -6,02 | *0,3807* | 1,61 | 0,23 | 7 | 1,51 | 0,16 | 7 | 1,335814 | 1,424058 | 1,500129 | 1,746601 | 1,7679 | 1,512301 | 1,993072 | 1,448401 | 1,637057 | 1,433185 | 1,317558 | 1,402756 | 1,588371 | 1,773986 |
| | vestibular nerve | -7,03 | *0,2090* | 0,2534 | 0,0181 | 7 | 0,2356 | 0,0301 | 7 | 0,258643 | 0,222129 | 0,279943 | 0,249514 | 0,243429 | 0,2556 | 0,264729 | 0,231257 | 0,2769 | 0,203871 | 0,213 | 0,206914 | 0,270814 | 0,246471 |
| | cochlear nerve | -5,83 | *0,4528* | 1,36 | 0,23 | 7 | 1,28 | 0,14 | 7 | 1,077171 | 1,201929 | 1,220186 | 1,497087 | 1,524471 | 1,256701 | 1,728343 | 1,217144 | 1,360157 | 1,229314 | 1,104558 | 1,195842 | 1,317556 | 1,527514 |
| | trapezoid body | 11,65 | *0,5277* | 0,2426 | 0,0888 | 7 | 0,2708 | 0,0727 | 7 | 0,185614 | 0,346886 | 0,1917 | 0,179529 | 0,219086 | 0,392529 | 0,182571 | 0,365143 | 0,279943 | 0,343843 | 0,310372 | 0,185614 | 0,209957 | 0,200829 |
| | dorsal acoustic stria | 61,34 | *0,2469* | 0,0086 | 0,0059 | 6 | 0,0139 | 0,0094 | 7 | | 0,003043 | 0,003043 | 0,009129 | 0,012171 | 0,018257 | 0,006086 | 0,012171 | 0,018257 | 0,009129 | 0,009129 | 0,009129 | 0,033471 | 0,006086 |
| | lateral lemniscus | -12,59 | *0,4033* | 0,8563 | 0,2762 | 7 | 0,7485 | 0,1752 | 7 | 0,639 | 0,599443 | 0,763757 | 1,080215 | 1,068043 | 0,581186 | 1,262786 | 0,593357 | 0,845914 | 0,620743 | 0,529457 | 0,785057 | 0,839828 | 1,025443 |
| | brachium of the inferior colliculus | 1,63 | *0,8081* | 0,2400 | 0,0290 | 7 | 0,2439 | 0,0299 | 7 | 0,252557 | 0,252557 | 0,222129 | 0,209957 | 0,200829 | 0,264729 | 0,2769 | 0,246472 | 0,203871 | 0,2556 | 0,2556 | 0,216043 | 0,2343 | 0,295157 |
| | vagus nerve | -12,50 | *0,5742* | 0,0070 | 0,0015 | 7 | 0,0061 | 0,0033 | 6 | 0,006086 | 0,006086 | 0,006086 | 0,006086 | 0,009129 | 0,006086 | 0,009129 | 0,003043 | 0,006086 | | 0,006086 | 0,003043 | 0,006086 | 0,012171 |
| | solitary tract | -12,50 | *0,5742* | 0,0070 | 0,0015 | 7 | 0,0061 | 0,0033 | 6 | 0,006086 | 0,006086 | 0,006086 | 0,006086 | 0,009129 | 0,006086 | 0,009129 | 0,003043 | 0,006086 | | 0,006086 | 0,003043 | 0,006086 | 0,012171 |
| | dorsal roots | 7,26 | *0,1999* | 0,6407 | 0,0339 | 7 | 0,6873 | 0,0813 | 7 | 0,639 | 0,602486 | 0,654215 | 0,687686 | 0,632914 | 0,672472 | 0,5964 | 0,693772 | 0,572057 | 0,754628 | 0,635957 | 0,678557 | 0,654214 | 0,821571 |
| | cervicothalamic tract | 7,26 | *0,1999* | 0,6407 | 0,0339 | 7 | 0,6873 | 0,0813 | 7 | 0,639 | 0,602486 | 0,654215 | 0,687686 | 0,632914 | 0,672472 | 0,5964 | 0,693772 | 0,572057 | 0,754628 | 0,635957 | 0,678557 | 0,654214 | 0,821571 |
| | medial lemniscus | 4,75 | *0,4702* | 0,6403 | 0,0337 | 7 | 0,6707 | 0,1002 | 7 | 0,639 | 0,602486 | 0,651172 | 0,687686 | 0,632914 | 0,672472 | 0,5964 | 0,693772 | 0,523372 | 0,754628 | 0,635957 | 0,678557 | 0,587271 | 0,821571 |
| | cerebellum related fiber tracts | -2,95 | *0,2795* | 9,36 | 0,52 | 7 | 9,08 | 0,38 | 7 | 8,745169 | 9,146834 | 8,924703 | 9,262463 | 9,302015 | 10,1175 | 10,011 | 9,274634 | 9,438944 | 8,854711 | 9,015989 | 8,358724 | 9,274623 | 9,356786 |
| | cerebellar commissure | -17,26 | *0,2917* | 0,0856 | 0,0210 | 7 | 0,0709 | 0,0284 | 7 | 0,094329 | 0,079114 | 0,103457 | 0,079114 | 0,045643 | 0,109543 | 0,088243 | 0,057814 | 0,0213 | 0,088243 | 0,1065 | 0,088243 | 0,054771 | 0,079114 |
| * | cerebellar peduncles | -6,11 | *0,0224* | 2,95 | 0,15 | 7 | 2,77 | 0,11 | 7 | 2,814642 | 2,802473 | 2,893758 | 2,975916 | 2,890714 | 3,09763 | 3,188914 | 2,94853 | 2,793343 | 2,808556 | 2,708144 | 2,726399 | 2,607727 | 2,808557 |
| | superior cerebelar peduncles | -6,03 | *0,2013* | 0,8507 | 0,0648 | 7 | 0,7994 | 0,0764 | 7 | 0,769843 | 0,772886 | 0,812443 | 0,906772 | 0,885471 | 0,879386 | 0,928071 | 0,858086 | 0,7242 | 0,821571 | 0,712029 | 0,845914 | 0,730285 | 0,903729 |
| * | superior cerebellar peduncle decussation | -33,81 | *0,0440* | 0,0604 | 0,0182 | 7 | 0,0400 | 0,0156 | 7 | 0,045643 | 0,051729 | 0,076071 | 0,076071 | 0,057814 | 0,033471 | 0,082157 | 0,066943 | 0,027386 | 0,054771 | 0,030429 | 0,039557 | 0,024343 | 0,036514 |
| | uncinate fascicle | -6,09 | *0,5076* | 0,0856 | 0,0147 | 7 | 0,0804 | 0,0138 | 7 | 0,069986 | 0,066943 | 0,091286 | 0,082157 | 0,1065 | 0,082157 | 0,100414 | 0,073029 | 0,076071 | 0,057814 | 0,076071 | 0,094329 | 0,088243 | 0,097371 |
| | ventral spinocerebellar tract | 3,60 | *0,6063* | 0,2773 | 0,0358 | 7 | 0,2873 | 0,0349 | 7 | 0,237343 | 0,237343 | 0,246472 | 0,3195 | 0,307329 | 0,304286 | 0,289071 | 0,316457 | 0,246471 | 0,307328 | 0,246472 | 0,316457 | 0,258643 | 0,3195 |
| | middle cerebellar peduncle | -6,55 | *0,1035* | 1,23 | 0,06 | 7 | 1,15 | 0,10 | 7 | 1,220185 | 1,211058 | 1,2141 | 1,244529 | 1,138029 | 1,293215 | 1,3206 | 1,287129 | 1,1502 | 1,278 | 1,1715 | 1,092385 | 1,004142 | 1,192386 |
| | inferior cerebellar peduncle | -5,57 | *0,1505* | 0,8668 | 0,0495 | 7 | 0,8185 | 0,0662 | 7 | 0,824614 | 0,818529 | 0,867215 | 0,824615 | 0,867214 | 0,925029 | 0,940243 | 0,803315 | 0,918943 | 0,708985 | 0,824615 | 0,7881 | 0,873299 | 0,812443 |
| | dorsal spinocerebellar tract | 0,59 | *0,8997* | 0,1465 | 0,0137 | 7 | 0,1474 | 0,0115 | 7 | 0,133886 | 0,1278 | 0,136929 | 0,155186 | 0,161271 | 0,1491 | 0,161271 | 0,161272 | 0,133886 | 0,158229 | 0,143014 | 0,139971 | 0,136928 | 0,158229 |
| | arbor vitae | -1,29 | *0,7038* | 6,32 | 0,39 | 7 | 6,24 | 0,39 | 7 | 5,836198 | 6,265246 | 5,927488 | 6,207432 | 6,365657 | 6,910332 | 6,733843 | 6,268289 | 6,624301 | 5,957912 | 6,201345 | 5,544083 | 6,612125 | 6,469114 |

| | | | | | | | | | | | | | | | | | | | | | | | |
|---|---|---|---|---|---|---|---|---|---|---|---|---|---|---|---|---|---|---|---|---|---|---|---|
| | supra-callosal cerebral white matter | -8,46 | *0,0509* | 0,9611 | 0,0770 | 7 | 0,8798 | 0,0616 | 7 | 1,037614 | 1,052829 | 0,931115 | 0,934158 | 0,852 | 0,897643 | 1,0224 | 0,885472 | 0,778972 | 0,855043 | 0,988929 | 0,888514 | 0,879385 | 0,882429 |
| | lateral forebrain bundle system | -2,67 | *0,3063* | 13,16 | 0,79 | 7 | 12,81 | 0,29 | 7 | 13,206 | 12,98388 | 12,21403 | 13,00822 | 12,38747 | 13,75981 | 14,53269 | 12,92302 | 12,70697 | 12,54874 | 12,95953 | 12,59742 | 12,56091 | 13,33989 |
| | corpus callosum | -2,03 | *0,4882* | 7,25 | 0,51 | 7 | 7,10 | 0,15 | 7 | 7,235912 | 7,117247 | 6,688202 | 7,135504 | 6,727757 | 7,70756 | 8,109215 | 7,22679 | 7,065515 | 6,892069 | 7,308945 | 6,955968 | 7,102025 | 7,141586 |
| | corpus callosum anterior forceps | 4,73 | *0,5197* | 1,59 | 0,28 | 7 | 1,67 | 0,09 | 7 | 1,536642 | 1,341901 | 1,481872 | 1,481872 | 1,357114 | 1,813544 | 2,117829 | 1,652272 | 1,7466 | 1,755728 | 1,637058 | 1,664442 | 1,710085 | 1,491 |
| | external capsule | -6,69 | *0,2155* | 0,8385 | 0,0908 | 7 | 0,7824 | 0,0671 | 7 | 0,888514 | 0,778972 | 0,812443 | 0,821572 | 0,730286 | 0,824615 | 1,013271 | 0,806358 | 0,721157 | 0,797228 | 0,848957 | 0,855042 | 0,669428 | 0,778971 |
| | corpus callosum extreme capsule | 10,11 | *0,2704* | 0,0774 | 0,0134 | 7 | 0,0852 | 0,0119 | 7 | 0,0639 | 0,088243 | 0,060857 | 0,079114 | 0,066943 | 0,091286 | 0,091286 | 0,0639 | 0,0852 | 0,088243 | 0,088243 | 0,079114 | 0,103457 | 0,088243 |
| * | genu of corpus callosum | -11,36 | *0,0090* | 0,8724 | 0,0536 | 7 | 0,7733 | 0,0643 | 7 | 0,909814 | 0,900686 | 0,906772 | 0,864172 | 0,827657 | 0,921986 | 0,775929 | 0,675515 | 0,775929 | 0,705943 | 0,858086 | 0,797228 | 0,772885 | 0,827657 |
| | corpus callosum posterior forceps | -2,46 | *0,6531* | 1,20 | 0,12 | 7 | 1,17 | 0,11 | 7 | 1,080214 | 1,189758 | 1,244529 | 1,159329 | 1,034571 | 1,390586 | 1,308429 | 1,214101 | 1,390586 | 1,1076 | 1,025443 | 1,138028 | 1,125856 | 1,198886 |
| | corpus callosum body | -3,28 | *0,4218* | 2,85 | 0,23 | 7 | 2,75 | 0,18 | 7 | 3,021556 | 2,936359 | 2,437329 | 2,689887 | 2,8116 | 2,881587 | 3,149357 | 2,99113 | 2,522529 | 2,656413 | 3,018515 | 2,656413 | 2,741613 | 2,686843 |
| | corpus callosum splenium | -1,19 | *0,8832* | 0,6581 | 0,0966 | 7 | 0,6503 | 0,0984 | 7 | 0,623786 | 0,6603 | 0,556843 | 0,861129 | 0,629871 | 0,608572 | 0,666386 | 0,629872 | 0,544672 | 0,578143 | 0,6816 | 0,620743 | 0,648128 | 0,848957 |
| | corticospinal tract | -4,08 | *0,3468* | 3,83 | 0,33 | 7 | 3,68 | 0,26 | 7 | 4,004399 | 3,903988 | 3,39583 | 3,687945 | 3,496243 | 3,983102 | 4,351286 | 3,785316 | 3,368443 | 3,815742 | 3,867473 | 3,745755 | 3,234555 | 3,910071 |
| | internal capsule | -3,35 | *0,5950* | 2,11 | 0,28 | 7 | 2,04 | 0,20 | 7 | 2,342999 | 2,199987 | 1,950472 | 1,810501 | 1,728343 | 2,355172 | 2,403857 | 2,099573 | 1,862229 | 2,108699 | 2,315615 | 1,996113 | 1,731385 | 2,181729 |
| | cerebal peduncle | -3,89 | *0,6192* | 0,9607 | 0,1533 | 7 | 0,9233 | 0,1182 | 7 | 0,848957 | 0,918943 | 0,824615 | 1,128901 | 0,943286 | 0,842872 | 1,217143 | 0,982843 | 0,8307 | 1,004143 | 0,769843 | 1,025442 | 0,800271 | 1,049786 |
| | pyramid | -2,72 | *0,6967* | 0,5595 | 0,0828 | 7 | 0,5442 | 0,0570 | 7 | 0,5325 | 0,441215 | 0,459472 | 0,614658 | 0,642043 | 0,632915 | 0,593357 | 0,587272 | 0,456429 | 0,578143 | 0,499029 | 0,614657 | 0,508157 | 0,565971 |
| | pyramidal decussation | -22,36 | *0,4377* | 0,1069 | 0,0663 | 7 | 0,0830 | 0,0417 | 7 | 0,188657 | 0,197786 | 0,066943 | 0,076071 | 0,079114 | 0,121714 | 0,018257 | 0,076071 | 0,112586 | 0,097371 | 0,133886 | 0,066943 | 0,091286 | 0,003043 |
| | thalamus related | -2,28 | *0,6128* | 2,08 | 0,09 | 7 | 2,03 | 0,22 | 7 | 1,965685 | 1,962644 | 2,130001 | 2,184773 | 2,163471 | 2,069144 | 2,072186 | 1,910915 | 2,273015 | 1,840928 | 1,783115 | 1,895699 | 2,224327 | 2,288229 |
| | external medullary lamina of the thalamus | 1,24 | *0,8880* | 0,1048 | 0,0136 | 7 | 0,1061 | 0,0197 | 7 | 0,097371 | 0,103457 | 0,097371 | 0,0852 | 0,112586 | 0,1278 | 0,109543 | 0,112586 | 0,091286 | 0,121714 | 0,094329 | 0,136929 | 0,079114 | 0,1065 |
| | optic radiation | -1,14 | *0,7758* | 1,65 | 0,05 | 7 | 1,63 | 0,16 | 7 | 1,630971 | 1,609672 | 1,606629 | 1,676615 | 1,5975 | 1,713129 | 1,685743 | 1,603587 | 1,822672 | 1,484914 | 1,460572 | 1,478828 | 1,71617 | 1,822671 |
| | auditory radiation | -9,15 | *0,5615* | 0,3278 | 0,1011 | 7 | 0,2978 | 0,0862 | 7 | 0,237343 | 0,249514 | 0,426 | 0,422957 | 0,453386 | 0,228214 | 0,2769 | 0,194743 | 0,359057 | 0,2343 | 0,228214 | 0,279943 | 0,429043 | 0,359057 |
| | extrapyramidal fiber systems | -0,62 | *0,9113* | 1,18 | 0,13 | 7 | 1,18 | 0,11 | 7 | 1,119771 | 1,068043 | 1,174543 | 1,095429 | 1,268871 | 1,119772 | 1,436229 | 1,122815 | 1,296257 | 1,061957 | 1,101515 | 1,071085 | 1,335814 | 1,241486 |
| | cerebral nuclei related | -2,82 | *0,6018* | 0,1078 | 0,0122 | 7 | 0,1048 | 0,0088 | 7 | 0,109543 | 0,100414 | 0,091286 | 0,103457 | 0,118671 | 0,103457 | 0,1278 | 0,115629 | 0,109543 | 0,103457 | 0,1065 | 0,088243 | 0,100414 | 0,109543 |
| | nigrostriatal tract | -2,82 | *0,6018* | 0,1078 | 0,0122 | 7 | 0,1048 | 0,0088 | 7 | 0,109543 | 0,100414 | 0,091286 | 0,103457 | 0,118671 | 0,103457 | 0,1278 | 0,115629 | 0,109543 | 0,103457 | 0,1065 | 0,088243 | 0,100414 | 0,109543 |
| | tectospinal pathway | -3,93 | *0,3019* | 0,4873 | 0,0370 | 7 | 0,4682 | 0,0286 | 7 | 0,450343 | 0,4473 | 0,5538 | 0,4899 | 0,492943 | 0,4686 | 0,508157 | 0,4686 | 0,495986 | 0,450343 | 0,456429 | 0,429043 | 0,514243 | 0,462514 |
| | doral tegmental decussation | 150,00 | *0,1780* | 0,0030 | 0,0000 | 4 | 0,0076 | 0,0071 | 6 | 0,003043 | 0,003043 | 0,003043 | | | | 0,003043 | 0,003043 | 0,018257 | | 0,003043 | 0,003043 | 0,015214 | 0,003043 |
| | crossed tectospinal pathway | -4,92 | *0,1757* | 0,4856 | 0,0369 | 7 | 0,4616 | 0,0229 | 7 | 0,4473 | 0,444257 | 0,550757 | 0,4899 | 0,492943 | 0,4686 | 0,505114 | 0,465557 | 0,477729 | 0,450343 | 0,453386 | 0,426 | 0,499028 | 0,459471 |
| | rubrospinal tract | 2,51 | *0,7811* | 0,5881 | 0,1062 | 7 | 0,6029 | 0,0874 | 7 | 0,559886 | 0,520329 | 0,529457 | 0,502072 | 0,657257 | 0,547715 | 0,800271 | 0,538586 | 0,690729 | 0,508157 | 0,538586 | 0,5538 | 0,721157 | 0,669429 |
| | ventral tegmental decussation | 14,98 | *0,3260* | 0,0416 | 0,0131 | 6 | 0,0478 | 0,0065 | 7 | 0,051729 | 0,054771 | 0,0426 | 0,018257 | | 0,045643 | 0,036514 | 0,039557 | 0,048686 | 0,054771 | 0,057814 | 0,045643 | 0,0426 | 0,045643 |
| * | medial forebrain bundle system | -6,50 | *0,0319* | 7,07 | 0,38 | 7 | 6,61 | 0,32 | 7 | 6,885984 | 7,302861 | 6,855559 | 7,059433 | 6,648643 | 6,943803 | 7,8171 | 6,931632 | 6,210472 | 6,356526 | 6,578659 | 6,636468 | 6,463025 | 7,117243 |
| * | cerebrum related | -6,92 | *0,0184* | 6,20 | 0,29 | 7 | 5,78 | 0,30 | 7 | 5,982255 | 6,301761 | 6,131359 | 6,307847 | 5,8788 | 6,06746 | 6,761229 | 5,964003 | 5,501486 | 5,458884 | 5,747959 | 5,729697 | 5,693182 | 6,332186 |
| | amygdalar capsule | 9,85 | *0,1828* | 0,1413 | 0,0197 | 7 | 0,1552 | 0,0169 | 7 | 0,124757 | 0,146057 | 0,136929 | 0,121714 | 0,124757 | 0,167357 | 0,167357 | 0,1491 | 0,164314 | 0,146057 | 0,182571 | 0,158228 | 0,1278 | 0,158229 |
| | anterior commissure temporal limb | -23,82 | *0,2706* | 0,3138 | 0,1009 | 7 | 0,2391 | 0,1375 | 7 | 0,380357 | 0,331672 | 0,295157 | 0,270814 | 0,130843 | 0,331672 | 0,456429 | 0,395572 | 0,057814 | 0,337757 | 0,313414 | 0,3195 | 0,060857 | 0,188657 |
| | cingulum bundle | -3,73 | *0,2220* | 1,09 | 0,04 | 7 | 1,05 | 0,07 | 7 | 1,0863 | 1,110643 | 1,083258 | 1,083258 | 1,074129 | 1,055872 | 1,165414 | 1,022401 | 1,007186 | 0,9585 | 1,0437 | 1,034571 | 1,168456 | 1,138029 |
| * | fornix system | -7,41 | *0,0280* | 4,34 | 0,22 | 7 | 4,02 | 0,26 | 7 | 4,043956 | 4,390845 | 4,305644 | 4,558203 | 4,241743 | 4,193059 | 4,670786 | 4,110902 | 3,955715 | 3,724456 | 3,876601 | 3,885727 | 4,059169 | 4,539943 |
| | alveus | -9,61 | *0,1170* | 1,32 | 0,16 | 7 | 1,19 | 0,11 | 7 | 1,119771 | 1,162372 | 1,463615 | 1,539687 | 1,351029 | 1,195843 | 1,3845 | 1,119772 | 1,223229 | 1,125857 | 1,083258 | 1,211057 | 1,147156 | 1,421014 |
| | dorsal fornix | 4,94 | *0,8082* | 0,0352 | 0,0137 | 7 | 0,0369 | 0,0125 | 7 | 0,018257 | 0,039557 | 0,054771 | 0,018257 | 0,039557 | 0,045643 | 0,030429 | 0,045643 | 0,057814 | 0,030429 | 0,0426 | 0,027386 | 0,033471 | 0,0213 |
| | fimbria | -9,86 | *0,1132* | 1,76 | 0,24 | 7 | 1,59 | 0,11 | 7 | 1,804414 | 2,090444 | 1,612715 | 1,731387 | 1,433186 | 1,618801 | 2,026543 | 1,731387 | 1,414929 | 1,615757 | 1,634015 | 1,542728 | 1,484913 | 1,679657 |
| | postcommissural fornix | -20,91 | *0,2831* | 0,2765 | 0,1149 | 7 | 0,2187 | 0,0706 | 7 | 0,179529 | 0,194743 | 0,331672 | 0,407743 | 0,441214 | 0,1704 | 0,209957 | 0,164314 | 0,304286 | 0,146057 | 0,197786 | 0,176486 | 0,328628 | 0,213 |
| | columns of the fornix | -23,90 | *0,2073* | 0,2765 | 0,1149 | 7 | 0,2104 | 0,0575 | 7 | 0,179529 | 0,194743 | 0,331672 | 0,407743 | 0,441214 | 0,1704 | 0,209957 | 0,164314 | 0,273857 | 0,146057 | 0,197786 | 0,176486 | 0,301243 | 0,213 |
| | hippocampal commissures | 3,59 | *0,6015* | 0,9555 | 0,1101 | 7 | 0,9898 | 0,1286 | 7 | 0,921985 | 0,903729 | 0,842872 | 0,861129 | 0,976757 | 1,162372 | 1,019357 | 1,049786 | 0,955457 | 0,806357 | 0,918943 | 0,928071 | 1,064999 | 1,204971 |
| | dorsal hippocampal commissure | 6,09 | *0,4592* | 0,8563 | 0,1047 | 7 | 0,9085 | 0,1462 | 7 | 0,797228 | 0,797229 | 0,791143 | 0,839829 | 0,782014 | 1,068043 | 0,918943 | 0,955458 | 0,897643 | 0,708985 | 0,812443 | 0,821571 | 1,013271 | 1,1502 |
| | ventral hippocampal commissure | -17,98 | *0,4576* | 0,0991 | 0,0550 | 7 | 0,0813 | 0,0253 | 7 | 0,124757 | 0,1065 | 0,051729 | 0,0213 | 0,194743 | 0,094329 | 0,100414 | 0,094329 | 0,057814 | 0,097371 | 0,1065 | 0,1065 | 0,051729 | 0,054771 |
| | stria terminalis | -1,81 | *0,6406* | 0,3117 | 0,0223 | 7 | 0,3060 | 0,0219 | 7 | 0,346886 | 0,322543 | 0,310372 | 0,273857 | 0,307329 | 0,3195 | 0,301243 | 0,286029 | 0,316457 | 0,292114 | 0,331672 | 0,331671 | 0,2769 | 0,307329 |
| | commissural branch of stria terminalis | 4,81 | *0,8287* | 0,0452 | 0,0207 | 7 | 0,0474 | 0,0157 | 7 | 0,073029 | 0,054771 | 0,060857 | 0,018257 | 0,027386 | 0,054771 | 0,027386 | 0,0426 | 0,048686 | 0,0639 | 0,060857 | 0,024343 | 0,030429 | 0,060857 |
| | hypothalamus related | -3,55 | *0,6130* | 0,8690 | 0,1279 | 7 | 0,8381 | 0,0906 | 7 | 0,903728 | 1,001101 | 0,7242 | 0,751586 | 0,769843 | 0,876343 | 1,055871 | 0,967629 | 0,708986 | 0,897643 | 0,8307 | 0,906771 | 0,769842 | 0,785057 |
| | medial forebrain bundle | -27,78 | *0,5251* | 0,0365 | 0,0317 | 7 | 0,0264 | 0,0239 | 6 | 0,027386 | 0,006086 | 0,009129 | 0,057814 | 0,076071 | 0,006086 | 0,073029 | 0,006086 | 0,060857 | 0,003043 | 0,027386 | 0,012171 | 0,048686 | |
| | supraoptic commissures | -20,48 | *0,1395* | 0,0361 | 0,0113 | 7 | 0,0287 | 0,0030 | 7 | 0,024343 | 0,051729 | 0,024343 | 0,045643 | 0,045643 | 0,030429 | 0,030429 | 0,027386 | 0,033471 | 0,027386 | 0,027386 | 0,030429 | 0,030429 | 0,024343 |
| | mammillary related | -4,06 | *0,3760* | 0,2465 | 0,0174 | 7 | 0,2365 | 0,0228 | 7 | 0,219086 | 0,267772 | 0,252557 | 0,243429 | 0,258643 | 0,228214 | 0,2556 | 0,252557 | 0,216043 | 0,252557 | 0,209957 | 0,2343 | 0,219086 | 0,270814 |
| | principal mammillary tract | -18,45 | *0,4178* | 0,0448 | 0,0104 | 7 | 0,0365 | 0,0234 | 7 | 0,024343 | 0,051729 | 0,054771 | 0,045643 | 0,045643 | 0,039557 | 0,051729 | 0,039557 | 0,006086 | 0,051729 | 0,018257 | 0,060857 | 0,015214 | 0,0639 |
| | mammillothalamic tract | 1,18 | *0,8664* | 0,1104 | 0,0130 | 7 | 0,1117 | 0,0153 | 7 | 0,121714 | 0,124757 | 0,115629 | 0,097371 | 0,118671 | 0,091286 | 0,103457 | 0,115629 | 0,124757 | 0,094329 | 0,1278 | 0,088243 | 0,109543 | 0,121714 |
| | mammillotegmental tract | -6,57 | *0,3362* | 0,0596 | 0,0076 | 7 | 0,0556 | 0,0070 | 7 | 0,054771 | 0,057814 | 0,060857 | 0,073029 | 0,057814 | 0,048686 | 0,0639 | 0,051729 | 0,0639 | 0,060857 | 0,0426 | 0,057814 | 0,054771 | 0,057814 |
| | mammillary peduncle | 2,74 | *0,8804* | 0,0317 | 0,0104 | 7 | 0,0326 | 0,0108 | 7 | 0,018257 | 0,033471 | 0,0213 | 0,027386 | 0,036514 | 0,048686 | 0,036514 | 0,045643 | 0,0213 | 0,045643 | 0,0213 | 0,027386 | 0,039557 | 0,027386 |
| | epithalamus related | 0,08 | *0,9946* | 0,5499 | 0,1338 | 7 | 0,5503 | 0,1007 | 7 | 0,632914 | 0,675515 | 0,438172 | 0,4047 | 0,389486 | 0,611615 | 0,696814 | 0,6816 | 0,398614 | 0,614657 | 0,565972 | 0,629871 | 0,471643 | 0,4899 |
| | stria medullaris | -11,18 | *0,4958* | 0,3617 | 0,1211 | 7 | 0,3212 | 0,0918 | 7 | 0,444257 | 0,429043 | 0,249514 | 0,225172 | 0,237343 | 0,422957 | 0,523371 | 0,444257 | 0,228214 | 0,380357 | 0,349929 | 0,389486 | 0,213 | 0,243429 |
| | fasciculus retroflexus | 11,99 | *0,3269* | 0,1378 | 0,0304 | 7 | 0,1543 | 0,0301 | 7 | 0,1491 | 0,182572 | 0,155186 | 0,1491 | 0,1278 | 0,1065 | 0,094329 | 0,133886 | 0,1278 | 0,143014 | 0,143014 | 0,143014 | 0,176486 | 0,213 |
| | habenular commissure | 48,28 | *0,1016* | 0,0504 | 0,0241 | 7 | 0,0748 | 0,0271 | 7 | 0,039557 | 0,0639 | 0,033471 | 0,030429 | 0,024343 | 0,082157 | 0,079114 | 0,103457 | 0,0426 | 0,091286 | 0,073029 | 0,097371 | 0,082157 | 0,033471 |
| | ventricular systems | -6,81 | *0,0811* | 6,66 | 0,33 | 7 | 6,21 | 0,53 | 7 | 6,828169 | 6,995532 | 6,459988 | 6,517804 | 6,107014 | 6,68516 | 7,019872 | 6,612132 | 5,659715 | 6,286541 | 6,246988 | 5,98834 | 5,577554 | 7,068557 |
| | lateral ventricle | -3,74 | *0,3231* | 4,03 | 0,31 | 7 | 3,88 | 0,22 | 7 | 4,095684 | 4,272174 | 3,71533 | 3,727502 | 3,694029 | 4,41823 | 4,299557 | 3,989188 | 3,742715 | 3,764013 | 4,07743 | 3,602741 | 3,754884 | 4,235657 |
| | subependymal zone | -0,34 | *0,9709* | 0,1282 | 0,0121 | 7 | 0,1278 | 0,0281 | 7 | 0,103457 | 0,124757 | 0,139971 | 0,136929 | 0,133886 | 0,130843 | 0,1278 | 0,118671 | 0,164314 | 0,1065 | 0,109543 | 0,094329 | 0,164314 | 0,136929 |
| | choroid plexus | 2,02 | *0,7801* | 1,55 | 0,25 | 7 | 1,58 | 0,15 | 7 | 1,710085 | 1,618801 | 1,271915 | 1,302344 | 1,396671 | 1,965687 | 1,567072 | 1,591415 | 1,527515 | 1,585328 | 1,567072 | 1,338856 | 1,591413 | 1,850057 |
| | third ventricle | -8,42 | *0,1384* | 1,01 | 0,07 | 7 | 0,9224 | 0,1217 | 7 | 1,010228 | 1,138029 | 0,998057 | 0,951529 | 1,028486 | 0,9372 | 0,955457 | 0,876343 | 1,034572 | 0,852 | 0,8307 | 0,867214 | 0,845914 | 1,1502 |
| | cerebral aqueduct | -14,47 | *0,2604* | 0,4656 | 0,1260 | 7 | 0,3982 | 0,0807 | 7 | 0,5112 | 0,438172 | 0,645086 | 0,456429 | 0,282986 | 0,346886 | 0,578143 | 0,453386 | 0,261686 | 0,453386 | 0,377314 | 0,441214 | 0,322543 | 0,477729 |
| | fourth ventricle | -13,03 | *0,2148* | 1,15 | 0,11 | 7 | 1,00 | 0,27 | 7 | 1,211057 | 1,147158 | 1,101515 | 1,351029 | 1,101514 | 0,982843 | 1,186714 | 1,293215 | 0,620743 | 1,217142 | 0,961543 | 1,077171 | 0,654214 | 1,204971 |
| | lateral recess | -7,99 | *0,1742* | 0,5003 | 0,0495 | 7 | 0,4603 | 0,0540 | 7 | 0,535543 | 0,486857 | 0,495986 | 0,572772 | 0,432086 | 0,4686 | 0,495986 | 0,517286 | 0,429043 | 0,462514 | 0,432086 | 0,426 | 0,4047 | 0,550757 |
| | #brainvol | -3,04 | *0,1126* | 483,65 | 17,29 | 7 | 468,94 | 14,68 | 7 | 477,0042 | 474,8807 | 464,3036 | 484,1219 | 477,7164 | 488,6861 | 518,8376 | 471,847 | 471,6581 | 455,2234 | 461,6168 | 457,9711 | 465,3195 | 498,9677 |
| | #maskvol | -3,04 | *0,1126* | 483,65 | 17,29 | 7 | 468,94 | 14,68 | 7 | 477,0042 | 474,8807 | 464,3036 | 484,1219 | 477,7164 | 488,6861 | 518,8376 | 471,847 | 471,6581 | 455,2234 | 461,6168 | 457,9711 | 465,3195 | 498,9677 |
