## Supplementary Table 2 for "Disruption of autism-associated *Pcdh9* gene leads to transcriptional alterations, synapses overgrowth and aberrant excitatory transmission in the CA1"

| Gene | p_val | avg_log2FC | pct.1 | pct.2 | p_val_adj | Cluster |
|---|---|---|---|---|---|---|
| Gm47283 | 2,3197E-54 | -0,869744 | 0,41 | 0,716 | 7,4892E-50 | 0 |
| Bc1 | 3,7534E-48 | 0,91991678 | 0,657 | 0,404 | 1,2118E-43 | 0 |
| Gm42439 | 6,1674E-10 | -0,3150512 | 0,832 | 0,897 | 1,9911E-05 | 0 |
| Epha5 | 7,5219E-10 | 0,3115842 | 0,938 | 0,905 | 2,4284E-05 | 0 |
| AC149090.1 | 1,0007E-09 | 0,32031353 | 0,983 | 0,978 | 3,2308E-05 | 0 |
| 530059O14Ri | 1,036E-09 | 0,31117188 | 0,651 | 0,55 | 3,3446E-05 | 0 |
| Add2 | 1,3111E-09 | 0,29894449 | 0,825 | 0,776 | 4,2327E-05 | 0 |
| Apoe | 1,4507E-09 | 0,26709076 | 0,931 | 0,908 | 4,6835E-05 | 0 |
| App | 6,7424E-09 | 0,2724201 | 0,746 | 0,656 | 0,00021768 | 0 |
| Egfem1 | 1,6853E-08 | 0,27918132 | 0,986 | 0,982 | 0,00054411 | 0 |
| Ncam2 | 8,0041E-07 | 0,30381234 | 0,865 | 0,845 | 0,0258413 | 0 |
| Gm47283 | 1,1764E-40 | -0,750008 | 0,386 | 0,667 | 3,798E-36 | 1 |
| Bc1 | 1,768E-39 | 0,9596951 | 0,641 | 0,381 | 5,708E-35 | 1 |
| Cox8a | 7,9739E-09 | 0,29869369 | 0,734 | 0,645 | 0,00025744 | 1 |
| Ddx5 | 3,6038E-08 | -0,2826844 | 0,88 | 0,916 | 0,00116348 | 1 |
| Olfm1 | 5,9698E-08 | -0,2559951 | 0,696 | 0,807 | 0,00192736 | 1 |
| Baz2b | 1,7001E-07 | -0,2512051 | 0,421 | 0,578 | 0,0054888 | 1 |
| Bc1 | 4,2717E-53 | 1,12244761 | 0,667 | 0,363 | 1,3791E-48 | 2 |
| Gm47283 | 1,9998E-13 | -0,329529 | 0,204 | 0,392 | 6,4564E-09 | 2 |
| Adipor2 | 1,155E-10 | 0,38929467 | 0,61 | 0,468 | 3,729E-06 | 2 |
| Mbp | 1,3328E-10 | 0,36251083 | 0,989 | 0,979 | 4,3028E-06 | 2 |
| Edil3 | 3,5402E-10 | 0,38931255 | 0,934 | 0,88 | 1,1429E-05 | 2 |
| Gm42418 | 5,3659E-09 | 0,37435237 | 0,974 | 0,958 | 0,00017324 | 2 |
| Mobp | 1,2441E-08 | 0,35769914 | 0,779 | 0,694 | 0,00040164 | 2 |
| Meg3 | 2,0552E-08 | 0,41103135 | 0,733 | 0,688 | 0,00066353 | 2 |
| St18 | 2,5075E-08 | 0,3749767 | 0,924 | 0,93 | 0,00080953 | 2 |
| Vmp1 | 3,4045E-08 | 0,29466482 | 0,831 | 0,74 | 0,00109914 | 2 |
| Phlpp1 | 5,3265E-08 | 0,32228737 | 0,908 | 0,875 | 0,00171966 | 2 |
| Apoe | 8,0428E-08 | 0,25200013 | 0,945 | 0,918 | 0,00259662 | 2 |
| Plp1 | 9,3861E-08 | 0,33877445 | 0,95 | 0,939 | 0,00303029 | 2 |
| Thra | 1,0388E-07 | 0,25954656 | 0,551 | 0,411 | 0,00335363 | 2 |
| Rab31 | 2,1476E-07 | 0,29301014 | 0,661 | 0,534 | 0,0069335 | 2 |
| Sept4 | 6,1672E-07 | 0,26601712 | 0,687 | 0,589 | 0,01991066 | 2 |
| Map7 | 8,6786E-07 | 0,298065 | 0,744 | 0,683 | 0,02801895 | 2 |
| Ncam1 | 9,2785E-07 | 0,27923227 | 0,807 | 0,743 | 0,02995553 | 2 |
| AC149090.1 | 1,2425E-06 | 0,3203292 | 0,652 | 0,546 | 0,04011484 | 2 |
| Snhg11 | 1,4732E-06 | 0,31919053 | 0,619 | 0,548 | 0,04756357 | 2 |
| Kif1b | 1,6608E-06 | 0,27010342 | 0,781 | 0,721 | 0,05361752 | 2 |
| Rps21 | 1,7677E-06 | 0,25988792 | 0,656 | 0,568 | 0,05706943 | 2 |
| Ppp1r16b | 1,9349E-06 | 0,27155827 | 0,628 | 0,567 | 0,06246795 | 2 |
| Tmeff2 | 2,071E-06 | 0,34016341 | 0,969 | 0,968 | 0,06686099 | 2 |
| Clasp2 | 2,0948E-06 | 0,25942773 | 0,715 | 0,665 | 0,06763032 | 2 |
| Ano4 | 3,0437E-06 | 0,29794115 | 0,829 | 0,821 | 0,09826651 | 2 |
| Bc1 | 3,7487E-51 | 1,45610944 | 0,759 | 0,413 | 1,2103E-46 | 3 |
| Pcdh9 | 5,7998E-23 | 0,71434923 | 0,984 | 0,965 | 1,8725E-18 | 3 |

| | | | | | | |
|---|---|---|---|---|---|---|
| Gm47283 | 9,2309E-17 | -0,3887437 | 0,179 | 0,408 | 2,9802E-12 | 3 |
| Ttr | 5,2093E-13 | -0,2757227 | 0,049 | 0,202 | 1,6818E-08 | 3 |
| Msmo1 | 2,6349E-10 | 0,37044186 | 0,501 | 0,383 | 8,5067E-06 | 3 |
| Sik3 | 2,9535E-10 | 0,39838616 | 0,633 | 0,523 | 9,5354E-06 | 3 |
| Kcnn2 | 2,837E-09 | 0,37597153 | 0,748 | 0,592 | 9,1591E-05 | 3 |
| AC149090.1 | 3,4853E-09 | 0,41138454 | 0,788 | 0,655 | 0,00011252 | 3 |
| Pcyt2 | 5,2959E-09 | 0,32884404 | 0,539 | 0,401 | 0,00017098 | 3 |
| Mir99ahg | 1,083E-08 | 0,36794921 | 0,94 | 0,927 | 0,00034964 | 3 |
| Plpp3 | 1,2565E-08 | 0,3338547 | 0,932 | 0,901 | 0,00040566 | 3 |
| Fam171b | 2,1758E-08 | 0,31596383 | 0,629 | 0,502 | 0,00070245 | 3 |
| Cpe | 3,9995E-07 | 0,31788008 | 0,947 | 0,937 | 0,01291252 | 3 |
| Nedd4 | 4,3264E-07 | 0,27030716 | 0,519 | 0,385 | 0,0139678 | 3 |
| Gnao1 | 6,4492E-07 | 0,30642265 | 0,81 | 0,721 | 0,02082114 | 3 |
| Sparcl1 | 6,8403E-07 | 0,31840084 | 0,863 | 0,815 | 0,02208396 | 3 |
| Rpl41 | 8,2379E-07 | 0,35949674 | 0,653 | 0,554 | 0,026596 | 3 |
| Nrcam | 8,7296E-07 | 0,30266695 | 0,777 | 0,746 | 0,02818344 | 3 |
| Vcl | 1,0077E-06 | 0,28502913 | 0,525 | 0,39 | 0,032535 | 3 |
| Nfia | 1,0377E-06 | 0,31559475 | 0,881 | 0,869 | 0,03350086 | 3 |
| Uba52 | 1,0509E-06 | 0,27429584 | 0,324 | 0,228 | 0,03392784 | 3 |
| Glul | 1,1121E-06 | 0,30804346 | 0,766 | 0,695 | 0,03590508 | 3 |
| mt-Atp6 | 1,127E-06 | 0,41318066 | 0,982 | 0,974 | 0,03638409 | 3 |
| Kif1b | 1,2119E-06 | 0,29569416 | 0,808 | 0,718 | 0,03912572 | 3 |
| Gpm6b | 1,3482E-06 | 0,30626847 | 0,92 | 0,894 | 0,04352608 | 3 |
| Slc1a3 | 1,3856E-06 | 0,30014023 | 0,931 | 0,892 | 0,04473321 | 3 |
| Kif5b | 1,4037E-06 | 0,25684296 | 0,413 | 0,275 | 0,04531697 | 3 |
| Mt3 | 1,9051E-06 | 0,32897248 | 0,709 | 0,655 | 0,06150737 | 3 |
| Tspan7 | 2,4931E-06 | 0,25566724 | 0,971 | 0,955 | 0,08048897 | 3 |
| Prnp | 2,5771E-06 | 0,25846875 | 0,516 | 0,42 | 0,08320072 | 3 |
| Gabra2 | 2,903E-06 | 0,31809246 | 0,547 | 0,439 | 0,09372435 | 3 |
| Gm47283 | 8,3071E-38 | -0,8391577 | 0,355 | 0,675 | 2,682E-33 | 4 |
| Bc1 | 7,1108E-36 | 0,97408721 | 0,66 | 0,405 | 2,2957E-31 | 4 |
| Ttr | 8,3553E-11 | -0,2795218 | 0,065 | 0,219 | 2,6975E-06 | 4 |
| Rtn1 | 1,6872E-08 | 0,28577577 | 0,96 | 0,93 | 0,00054472 | 4 |
| Pcdh9 | 2,8034E-08 | 0,37675927 | 0,995 | 0,994 | 0,00090509 | 4 |
| Bsn | 7,5983E-07 | 0,28144623 | 0,736 | 0,669 | 0,02453118 | 4 |
| Gabrb3 | 1,0406E-06 | 0,30929435 | 0,97 | 0,951 | 0,03359534 | 4 |
| Hnrnpr | 1,2138E-06 | 0,25644731 | 0,501 | 0,346 | 0,03918787 | 4 |
| Ddx5 | 1,3244E-06 | -0,2785644 | 0,927 | 0,958 | 0,04275773 | 4 |
| Srgap3 | 1,918E-06 | 0,27767024 | 0,826 | 0,775 | 0,06192382 | 4 |
| Ptprd | 2,3574E-06 | 0,32125954 | 0,997 | 0,994 | 0,07610912 | 4 |
| Gm47283 | 4,4484E-56 | -1,0842702 | 0,449 | 0,791 | 1,4362E-51 | 5 |
| Bc1 | 5,0356E-37 | 1,03085025 | 0,816 | 0,642 | 1,6258E-32 | 5 |
| 530059O14Ri | 7,2291E-10 | 0,39396484 | 0,908 | 0,856 | 2,3339E-05 | 5 |
| Camk2a | 4,5992E-09 | 0,31432603 | 0,993 | 0,982 | 0,00014848 | 5 |
| Ksr1 | 8,2551E-09 | 0,37873756 | 0,665 | 0,572 | 0,00026652 | 5 |
| AC149090.1 | 5,0238E-08 | 0,32585034 | 0,993 | 0,973 | 0,00162193 | 5 |
| Gm10848 | 7,5583E-08 | -0,3184924 | 0,76 | 0,809 | 0,0024402 | 5 |
| Pcdh9 | 1,3618E-07 | 0,43041593 | 0,971 | 0,971 | 0,00439667 | 5 |

| | | | | | | |
|---|---|---|---|---|---|---|
| 930415C11Ri | 4,3943E-07 | -0,2833536 | 0,319 | 0,435 | 0,01418705 | 5 |
| Gm21798 | 1,1111E-06 | -0,2683425 | 0,391 | 0,545 | 0,03587291 | 5 |
| Dnaja2 | 1,2639E-06 | -0,2542399 | 0,485 | 0,574 | 0,04080359 | 5 |
| Specc1 | 1,5431E-06 | 0,29664397 | 0,778 | 0,676 | 0,04981758 | 5 |
| Csmd2 | 1,6423E-06 | 0,30925742 | 0,739 | 0,662 | 0,05302247 | 5 |
| Rps8 | 1,9294E-06 | 0,25912253 | 0,804 | 0,721 | 0,06228976 | 5 |
| Frmd4a | 2,3182E-06 | 0,30890232 | 0,926 | 0,926 | 0,07484465 | 5 |
| Gm47283 | 2,6579E-46 | -0,9844643 | 0,494 | 0,736 | 8,581E-42 | 6 |
| Bc1 | 2,3526E-45 | 1,1716248 | 0,86 | 0,674 | 7,5954E-41 | 6 |
| Gm36264 | 1,0116E-10 | 0,27740076 | 0,286 | 0,124 | 3,266E-06 | 6 |
| Apoe | 5,3901E-10 | 0,33129479 | 0,956 | 0,908 | 1,7402E-05 | 6 |
| 530059O14Ri | 6,2724E-10 | 0,37509644 | 0,947 | 0,901 | 2,025E-05 | 6 |
| Pcdh9 | 1,3365E-09 | 0,55744204 | 0,971 | 0,968 | 4,315E-05 | 6 |
| Ksr1 | 1,8269E-09 | 0,3774154 | 0,787 | 0,713 | 5,8981E-05 | 6 |
| Shank2 | 5,946E-09 | 0,35876134 | 0,809 | 0,711 | 0,00019197 | 6 |
| Dapk1 | 7,0015E-09 | 0,37356366 | 0,86 | 0,78 | 0,00022604 | 6 |
| Cttn | 1,2517E-08 | 0,31785677 | 0,528 | 0,374 | 0,0004041 | 6 |
| Snhg11 | 1,4362E-08 | 0,34390797 | 1 | 1 | 0,00046366 | 6 |
| Larp1 | 1,5057E-08 | -0,2880223 | 0,385 | 0,534 | 0,0004861 | 6 |
| Ddx5 | 3,7175E-08 | -0,3166938 | 0,954 | 0,943 | 0,00120019 | 6 |
| Sv2b | 6,6266E-08 | 0,38458113 | 0,821 | 0,759 | 0,00213939 | 6 |
| Dnmt3a | 8,663E-08 | 0,32979018 | 0,683 | 0,589 | 0,00279686 | 6 |
| Camk2a | 3,3206E-07 | 0,26887639 | 0,995 | 0,989 | 0,01072065 | 6 |
| Pabpc1 | 4,895E-07 | 0,2622183 | 0,429 | 0,261 | 0,0158034 | 6 |
| Meg3 | 5,6022E-07 | 0,30220819 | 1 | 1 | 0,01808685 | 6 |
| Olfm1 | 7,2332E-07 | -0,2656863 | 0,77 | 0,83 | 0,02335249 | 6 |
| Pkp2 | 1,1302E-06 | 0,28220025 | 0,799 | 0,695 | 0,0364879 | 6 |
| Galnt9 | 1,3446E-06 | 0,29999148 | 0,763 | 0,693 | 0,04341056 | 6 |
| Bc1 | 1,1539E-44 | 1,09012876 | 0,773 | 0,444 | 3,7253E-40 | 7 |
| Gm19951 | 1,9593E-11 | 0,43372649 | 0,454 | 0,256 | 6,3257E-07 | 7 |
| Uba52 | 3,1719E-10 | 0,30979597 | 0,321 | 0,158 | 1,024E-05 | 7 |
| Apoe | 5,7609E-08 | 0,32049958 | 0,966 | 0,889 | 0,00185992 | 7 |
| Hbb-bs | 1,8385E-07 | 2,30678506 | 0,202 | 0,16 | 0,00593552 | 7 |
| Gm42418 | 1,8468E-07 | 0,36339276 | 0,989 | 0,984 | 0,00596233 | 7 |
| mt-Co1 | 3,643E-07 | -0,3685258 | 0,819 | 0,897 | 0,01176135 | 7 |
| Gm47283 | 5,5681E-07 | -0,2618618 | 0,154 | 0,323 | 0,01797654 | 7 |
| Tshz2 | 2,8207E-06 | 0,25271074 | 0,124 | 0,039 | 0,09106666 | 7 |
| Gm47283 | 1,3548E-48 | -1,1712278 | 0,587 | 0,835 | 4,3739E-44 | 9 |
| Bc1 | 2,1564E-25 | 0,9967819 | 0,79 | 0,635 | 6,9619E-21 | 9 |
| Gm12296 | 8,2627E-12 | -0,5147032 | 0,852 | 0,883 | 2,6676E-07 | 9 |
| Gm2000 | 1,4408E-10 | 0,30117936 | 0,258 | 0,1 | 4,6515E-06 | 9 |
| Larp1 | 5,5496E-10 | -0,3670748 | 0,419 | 0,628 | 1,7917E-05 | 9 |
| Gm13269 | 2,6337E-09 | -0,4484968 | 0,681 | 0,796 | 8,503E-05 | 9 |
| Acp1 | 5,552E-09 | -0,3222455 | 0,777 | 0,856 | 0,00017925 | 9 |
| Gm10848 | 3,3782E-08 | -0,4135804 | 0,568 | 0,715 | 0,00109065 | 9 |
| Rps8 | 2,6401E-07 | 0,32411686 | 0,855 | 0,798 | 0,00852364 | 9 |
| Gm36264 | 2,7048E-07 | 0,25911656 | 0,2 | 0,085 | 0,00873236 | 9 |
| Kpna4 | 4,6873E-07 | -0,3180854 | 0,539 | 0,686 | 0,01513287 | 9 |

| | | | | | | |
|---|---|---|---|---|---|---|
| Cacnb4 | 5,1053E-07 | 0,4017678 | 0,732 | 0,706 | 0,01648242 | 9 |
| Ptp4a2 | 1,4148E-06 | -0,2972035 | 0,377 | 0,574 | 0,04567725 | 9 |
| Rph3a | 1,7918E-06 | 0,39468077 | 0,632 | 0,489 | 0,05784988 | 9 |
| mt-Atp6 | 2,8647E-06 | 0,38947937 | 0,984 | 0,988 | 0,09248671 | 9 |
| Bc1 | 1,5983E-28 | 0,99442845 | 0,641 | 0,38 | 5,16E-24 | 10 |
| Gm47283 | 6,0809E-19 | -0,6190418 | 0,455 | 0,687 | 1,9632E-14 | 10 |
| Pcdh9 | 3,7182E-12 | 0,45048017 | 0,997 | 0,992 | 1,2004E-07 | 10 |
| Camk2a | 4,1976E-12 | 0,36778534 | 0,994 | 0,992 | 1,3552E-07 | 10 |
| Cst3 | 3,2452E-07 | 0,32680891 | 0,881 | 0,763 | 0,01047707 | 10 |
| Ncam2 | 5,4163E-07 | 0,36728653 | 0,939 | 0,961 | 0,01748646 | 10 |
| Larp1 | 5,94E-07 | -0,2532745 | 0,243 | 0,434 | 0,0191772 | 10 |
| Lrrtm4 | 1,119E-06 | 0,28119825 | 1 | 1 | 0,03612731 | 10 |
| Anp32a | 2,5477E-06 | 0,26005726 | 0,394 | 0,239 | 0,08225388 | 10 |
| Bc1 | 1,5719E-30 | 1,08400147 | 0,7 | 0,458 | 5,0748E-26 | 11 |
| Gm47283 | 9,8573E-24 | -0,772695 | 0,58 | 0,751 | 3,1824E-19 | 11 |
| Pcdh9 | 5,7334E-11 | 0,42432307 | 1 | 1 | 1,851E-06 | 11 |
| Egfem1 | 5,5904E-09 | 0,35756026 | 1 | 0,993 | 0,00018049 | 11 |
| Cntn5 | 7,4112E-09 | 0,42619014 | 0,98 | 0,94 | 0,00023927 | 11 |
| Ank2 | 1,8297E-08 | 0,32563714 | 1 | 0,993 | 0,00059071 | 11 |
| Ncam2 | 2,6367E-08 | 0,4384807 | 0,977 | 0,973 | 0,00085126 | 11 |
| Camk2a | 4,8889E-08 | 0,28411641 | 1 | 1 | 0,00157838 | 11 |
| Gm2000 | 1,9255E-07 | 0,27677817 | 0,268 | 0,106 | 0,00621648 | 11 |
| Tmeff2 | 1,9542E-07 | 0,41264401 | 0,939 | 0,934 | 0,00630915 | 11 |
| Cacna2d1 | 2,2922E-07 | 0,33178362 | 0,983 | 0,973 | 0,00740049 | 11 |
| Kctd16 | 2,56E-07 | 0,398958 | 0,983 | 0,997 | 0,00826501 | 11 |
| Ptprd | 5,9979E-07 | 0,31826683 | 1 | 1 | 0,01936412 | 11 |
| Kcnd2 | 6,8943E-07 | 0,30783666 | 1 | 1 | 0,02225829 | 11 |
| AC149090.1 | 7,8978E-07 | 0,32922895 | 1 | 0,993 | 0,02549795 | 11 |
| Prr16 | 8,1473E-07 | 0,49198004 | 0,481 | 0,425 | 0,02630362 | 11 |
| Rbbp6 | 9,1622E-07 | 0,3410042 | 0,589 | 0,475 | 0,02958013 | 11 |
| Ppfia2 | 1,4708E-06 | 0,32031567 | 1 | 1 | 0,04748486 | 11 |
| Kbtbd11 | 1,5491E-06 | 0,32235139 | 0,708 | 0,625 | 0,05001387 | 11 |
| Larp1 | 1,5783E-06 | -0,2842041 | 0,385 | 0,515 | 0,05095385 | 11 |
| Mctp1 | 1,9785E-06 | 0,33230966 | 0,983 | 0,987 | 0,06387682 | 11 |
| Stau2 | 2,0281E-06 | 0,33726533 | 0,764 | 0,654 | 0,06547834 | 11 |
| Add2 | 2,3556E-06 | 0,30085128 | 0,913 | 0,894 | 0,0760496 | 11 |
| App | 2,5063E-06 | 0,30289578 | 0,895 | 0,844 | 0,08091513 | 11 |
| Kcnb2 | 2,5359E-06 | 0,36381787 | 0,936 | 0,934 | 0,08187206 | 11 |
| Faah | 2,6797E-06 | 0,30258853 | 0,691 | 0,588 | 0,08651364 | 11 |
| Ppm1h | 2,8862E-06 | 0,31814635 | 0,79 | 0,691 | 0,09318007 | 11 |
| Bc1 | 3,9333E-38 | 1,29655358 | 0,728 | 0,45 | 1,2699E-33 | 12 |
| Gm47283 | 1,015E-21 | -0,7006503 | 0,616 | 0,795 | 3,2768E-17 | 12 |
| Lrrtm4 | 9,9693E-17 | 0,52529709 | 1 | 1 | 3,2186E-12 | 12 |
| Pcdh9 | 2,552E-15 | 0,54857978 | 1 | 1 | 8,239E-11 | 12 |
| Tenm2 | 3,5896E-15 | 0,50329152 | 1 | 1 | 1,1589E-10 | 12 |
| Camk2a | 4,5158E-15 | 0,44501844 | 1 | 0,997 | 1,4579E-10 | 12 |
| Negr1 | 1,294E-14 | 0,50882962 | 1 | 0,997 | 4,1776E-10 | 12 |
| Kctd16 | 1,4944E-14 | 0,56374388 | 1 | 0,997 | 4,8247E-10 | 12 |

| | | | | | | |
|---|---|---|---|---|---|---|
| Cacna2d1 | 3,5501E-13 | 0,50898676 | 0,989 | 0,968 | 1,1462E-08 | 12 |
| Tmeff2 | 6,055E-13 | 0,5630064 | 0,975 | 0,983 | 1,9548E-08 | 12 |
| Ptprd | 2,2804E-12 | 0,46588912 | 1 | 1 | 7,3624E-08 | 12 |
| Shisa9 | 2,6486E-12 | 0,50354987 | 0,978 | 0,951 | 8,5511E-08 | 12 |
| Egfem1 | 2,7297E-12 | 0,5040579 | 0,993 | 0,994 | 8,8129E-08 | 12 |
| Ncam2 | 2,8439E-12 | 0,59472303 | 0,968 | 0,977 | 9,1814E-08 | 12 |
| Meg3 | 7,3662E-12 | 0,45106897 | 1 | 1 | 2,3782E-07 | 12 |
| Grik2 | 7,7599E-12 | 0,50101931 | 0,993 | 1 | 2,5053E-07 | 12 |
| Anks1b | 4,3003E-11 | 0,41826785 | 1 | 0,997 | 1,3884E-06 | 12 |
| Ralyl | 4,9497E-11 | 0,54964015 | 0,975 | 0,948 | 1,598E-06 | 12 |
| Arhgap39 | 5,5705E-11 | 0,42580621 | 0,903 | 0,787 | 1,7984E-06 | 12 |
| Prickle2 | 5,6008E-11 | 0,45227661 | 0,928 | 0,856 | 1,8082E-06 | 12 |
| Sobp | 5,6566E-11 | 0,44966175 | 0,943 | 0,873 | 1,8262E-06 | 12 |
| Mctp1 | 6,9595E-11 | 0,46368049 | 0,989 | 0,974 | 2,2469E-06 | 12 |
| Dcc | 9,145E-11 | 0,49526355 | 0,996 | 0,994 | 2,9525E-06 | 12 |
| Kcnip4 | 3,608E-10 | 0,3953686 | 1 | 1 | 1,1648E-05 | 12 |
| Tcf20 | 3,6712E-10 | 0,45538362 | 0,828 | 0,709 | 1,1853E-05 | 12 |
| Gabra2 | 4,2915E-10 | 0,47140586 | 0,975 | 0,963 | 1,3855E-05 | 12 |
| Nkain2 | 4,6142E-10 | 0,41226085 | 1 | 1 | 1,4897E-05 | 12 |
| Dapk1 | 4,8654E-10 | 0,41911866 | 0,95 | 0,928 | 1,5708E-05 | 12 |
| Csmd1 | 5,0181E-10 | 0,39441442 | 1 | 1 | 1,6201E-05 | 12 |
| Lsamp | 6,6781E-10 | 0,40185605 | 1 | 1 | 2,156E-05 | 12 |
| Dock4 | 6,8661E-10 | 0,41815954 | 0,989 | 0,971 | 2,2167E-05 | 12 |
| Pip5k1b | 1,1373E-09 | 0,42881705 | 0,978 | 0,931 | 3,6718E-05 | 12 |
| Dpyd | 1,1506E-09 | 0,46693055 | 0,86 | 0,787 | 3,7148E-05 | 12 |
| Dlg2 | 1,1989E-09 | 0,38835006 | 1 | 1 | 3,8707E-05 | 12 |
| Slc8a1 | 1,7358E-09 | 0,54593202 | 0,728 | 0,608 | 5,6041E-05 | 12 |
| 530059O14Ri | 1,9065E-09 | 0,42920354 | 0,781 | 0,706 | 6,1552E-05 | 12 |
| Tnik | 2,4598E-09 | 0,3824719 | 0,978 | 0,974 | 7,9416E-05 | 12 |
| Setbp1 | 2,6231E-09 | 0,40280854 | 0,939 | 0,931 | 8,4687E-05 | 12 |
| Robo2 | 3,6994E-09 | 0,45985797 | 0,925 | 0,882 | 0,00011944 | 12 |
| Slit3 | 3,8038E-09 | 0,34348175 | 0,996 | 0,994 | 0,00012281 | 12 |
| Csmd3 | 4,0972E-09 | 0,41613644 | 1 | 0,994 | 0,00013228 | 12 |
| Arpp21 | 5,2494E-09 | 0,41141227 | 0,964 | 0,963 | 0,00016948 | 12 |
| Phlpp1 | 6,6216E-09 | 0,42143915 | 0,688 | 0,591 | 0,00021378 | 12 |
| Frmd4a | 7,8567E-09 | 0,36819106 | 0,996 | 0,991 | 0,00025365 | 12 |
| Ank2 | 7,8949E-09 | 0,36288341 | 1 | 0,991 | 0,00025489 | 12 |
| Atp8a1 | 8,0317E-09 | 0,41469353 | 0,885 | 0,847 | 0,0002593 | 12 |
| Gripap1 | 1,107E-08 | 0,32583265 | 0,588 | 0,383 | 0,00035739 | 12 |
| Epb41l1 | 1,214E-08 | 0,39211463 | 0,796 | 0,683 | 0,00039195 | 12 |
| Nbea | 1,2538E-08 | 0,38508057 | 1 | 0,991 | 0,0004048 | 12 |
| Dock3 | 1,2566E-08 | 0,40607517 | 0,907 | 0,896 | 0,0004057 | 12 |
| Fgf12 | 1,3109E-08 | 0,40139094 | 0,925 | 0,893 | 0,00042324 | 12 |
| Prkce | 1,5497E-08 | 0,34502593 | 1 | 0,991 | 0,00050031 | 12 |
| Pitpnc1 | 1,6081E-08 | 0,44175991 | 0,742 | 0,605 | 0,00051918 | 12 |
| Ksr2 | 1,9469E-08 | 0,37612922 | 0,864 | 0,83 | 0,00062854 | 12 |
| Kcnh1 | 3,2505E-08 | 0,39185855 | 0,746 | 0,64 | 0,00104942 | 12 |
| Slc1a2 | 3,3157E-08 | 0,42500445 | 0,789 | 0,695 | 0,00107048 | 12 |

| | | | | | | |
|---|---|---|---|---|---|---|
| Nlgn1 | 3,6483E-08 | 0,36882924 | 1 | 1 | 0,00117786 | 12 |
| Ppfia2 | 3,6583E-08 | 0,38365817 | 0,993 | 1 | 0,00118108 | 12 |
| 110051M20R | 3,8306E-08 | 0,37482541 | 0,792 | 0,729 | 0,00123671 | 12 |
| Adgrb3 | 4,028E-08 | 0,37053314 | 1 | 1 | 0,00130043 | 12 |
| Castor2 | 4,109E-08 | 0,32716148 | 0,437 | 0,271 | 0,00132658 | 12 |
| Safb | 4,3026E-08 | 0,32477703 | 0,642 | 0,481 | 0,0013891 | 12 |
| Gabrb3 | 4,9147E-08 | 0,38173125 | 0,993 | 0,997 | 0,00158671 | 12 |
| Ak5 | 5,0314E-08 | 0,34960818 | 0,968 | 0,948 | 0,00162439 | 12 |
| Nav2 | 5,0992E-08 | 0,33293737 | 0,986 | 0,965 | 0,00164629 | 12 |
| Cdh13 | 5,2961E-08 | 0,42417843 | 0,903 | 0,905 | 0,00170985 | 12 |
| Tspan5 | 5,4991E-08 | 0,41420621 | 0,882 | 0,804 | 0,00177537 | 12 |
| Tmcc1 | 5,5115E-08 | 0,38614993 | 0,914 | 0,853 | 0,00177938 | 12 |
| Pcbp3 | 6,0621E-08 | 0,35474798 | 0,477 | 0,334 | 0,00195716 | 12 |
| Add2 | 6,5767E-08 | 0,3697119 | 0,892 | 0,827 | 0,00212329 | 12 |
| Crim1 | 6,8517E-08 | 0,32383054 | 0,448 | 0,291 | 0,00221208 | 12 |
| Cttnbp2 | 7,1698E-08 | 0,32631608 | 0,996 | 0,994 | 0,00231477 | 12 |
| Lrfn5 | 7,7395E-08 | 0,38678597 | 0,993 | 0,994 | 0,00249871 | 12 |
| Abr | 7,8711E-08 | 0,34748711 | 0,896 | 0,89 | 0,00254119 | 12 |
| Gm26871 | 8,1018E-08 | 0,41212542 | 0,964 | 0,931 | 0,00261565 | 12 |
| Dlgap2 | 8,447E-08 | 0,33273792 | 1 | 1 | 0,00272712 | 12 |
| Ust | 8,6406E-08 | 0,41355 | 0,667 | 0,611 | 0,00278961 | 12 |
| Unc5c | 8,9291E-08 | 0,4973951 | 0,753 | 0,683 | 0,00288275 | 12 |
| Dscaml1 | 9,7095E-08 | 0,34586061 | 0,975 | 0,951 | 0,00313472 | 12 |
| Sez6l | 9,7184E-08 | 0,37737847 | 0,878 | 0,839 | 0,00313759 | 12 |
| Lrrc7 | 9,8354E-08 | 0,36262261 | 0,996 | 0,997 | 0,00317535 | 12 |
| Snx30 | 9,9049E-08 | 0,33205262 | 0,487 | 0,337 | 0,0031978 | 12 |
| Grm5 | 9,9515E-08 | 0,37134435 | 1 | 1 | 0,00321283 | 12 |
| Il1rapl1 | 1,003E-07 | 0,38193108 | 1 | 1 | 0,00323809 | 12 |
| Macrod2 | 1,0486E-07 | 0,38006202 | 1 | 1 | 0,0033854 | 12 |
| Kansl1 | 1,1693E-07 | 0,36892782 | 0,896 | 0,821 | 0,0037752 | 12 |
| Zfp804b | 1,2667E-07 | 0,55246212 | 0,792 | 0,718 | 0,00408949 | 12 |
| Cpne4 | 1,2876E-07 | 0,42575101 | 0,835 | 0,801 | 0,00415712 | 12 |
| Snph | 1,3468E-07 | 0,36473754 | 0,645 | 0,496 | 0,00434802 | 12 |
| Lcorl | 1,3531E-07 | 0,46897485 | 0,692 | 0,669 | 0,00436856 | 12 |
| Zfp638 | 1,4053E-07 | 0,37093876 | 0,71 | 0,588 | 0,00453693 | 12 |
| Nrg2 | 1,4539E-07 | 0,36906568 | 0,882 | 0,761 | 0,00469404 | 12 |
| Map1b | 1,4784E-07 | 0,32143766 | 0,986 | 0,991 | 0,00477302 | 12 |
| Cntnap5a | 1,5064E-07 | 0,39763546 | 0,993 | 0,991 | 0,00486348 | 12 |
| Cntn5 | 1,6223E-07 | 0,41185195 | 0,957 | 0,934 | 0,00523763 | 12 |
| Snhg14 | 1,8104E-07 | 0,39243137 | 0,925 | 0,916 | 0,00584495 | 12 |
| Ryr2 | 1,813E-07 | 0,34158304 | 1 | 1 | 0,00585314 | 12 |
| Apba2 | 1,8461E-07 | 0,33566144 | 0,828 | 0,758 | 0,00596018 | 12 |
| Klhl29 | 2,0773E-07 | 0,34620611 | 0,896 | 0,89 | 0,00670652 | 12 |
| Pygb | 2,207E-07 | 0,34715839 | 0,452 | 0,303 | 0,00712535 | 12 |
| Pacsin1 | 2,208E-07 | 0,32436394 | 0,824 | 0,784 | 0,00712862 | 12 |
| Dnajc6 | 2,2847E-07 | 0,37500107 | 0,774 | 0,68 | 0,00737622 | 12 |
| Gria3 | 2,8523E-07 | 0,34228616 | 0,968 | 0,968 | 0,00920859 | 12 |
| Dbn1 | 2,8593E-07 | 0,31467054 | 0,749 | 0,654 | 0,00923124 | 12 |

| | | | | | | |
|---|---|---|---|---|---|---|
| Gpatch8 | 2,9056E-07 | 0,33413384 | 0,9 | 0,89 | 0,0093808 | 12 |
| Nrg3 | 3,0027E-07 | 0,34906604 | 1 | 1 | 0,00969426 | 12 |
| Tmem108 | 3,2434E-07 | 0,34428869 | 0,989 | 0,974 | 0,01047144 | 12 |
| Ube2h | 3,2721E-07 | 0,33753465 | 0,663 | 0,527 | 0,01056398 | 12 |
| Auts2 | 3,5806E-07 | 0,28761257 | 1 | 1 | 0,01155987 | 12 |
| Ccnd2 | 3,626E-07 | 0,35047126 | 0,624 | 0,45 | 0,01170663 | 12 |
| Kalrn | 3,64E-07 | 0,31413312 | 1 | 0,997 | 0,01175172 | 12 |
| Cntn1 | 3,6987E-07 | 0,36777119 | 0,943 | 0,925 | 0,01194134 | 12 |
| Nos1ap | 3,7836E-07 | 0,35967053 | 0,86 | 0,795 | 0,01221546 | 12 |
| Adgrl3 | 3,7999E-07 | 0,36620084 | 0,946 | 0,911 | 0,0122681 | 12 |
| Ncam1 | 3,8291E-07 | 0,34456292 | 0,946 | 0,928 | 0,01236235 | 12 |
| Plekhg5 | 3,9666E-07 | 0,34702878 | 0,871 | 0,79 | 0,01280608 | 12 |
| Plekha5 | 4,0592E-07 | 0,36413922 | 0,91 | 0,844 | 0,01310513 | 12 |
| Xkr4 | 4,2812E-07 | 0,37475576 | 0,957 | 0,98 | 0,01382182 | 12 |
| Mdga2 | 4,3576E-07 | 0,36637809 | 0,982 | 0,983 | 0,01406846 | 12 |
| Pde10a | 4,7447E-07 | 0,4050732 | 0,839 | 0,807 | 0,01531833 | 12 |
| Rapgef4 | 4,8607E-07 | 0,34175264 | 0,892 | 0,847 | 0,01569265 | 12 |
| Dab1 | 4,8735E-07 | 0,31188676 | 1 | 1 | 0,01573407 | 12 |
| Osbpl6 | 4,9116E-07 | 0,35654129 | 0,778 | 0,663 | 0,01585711 | 12 |
| Ahcyl2 | 5,6811E-07 | 0,34458359 | 0,993 | 0,988 | 0,01834152 | 12 |
| Sltm | 5,7139E-07 | 0,36778188 | 0,756 | 0,663 | 0,0184474 | 12 |
| Dlgap1 | 5,9065E-07 | 0,30241307 | 1 | 1 | 0,01906915 | 12 |
| Phactr1 | 5,9952E-07 | 0,35797239 | 0,971 | 0,954 | 0,01935541 | 12 |
| Lingo2 | 6,1207E-07 | 0,34603287 | 1 | 1 | 0,0197606 | 12 |
| Gabrb2 | 6,3754E-07 | 0,36758843 | 0,885 | 0,859 | 0,02058304 | 12 |
| Dis3l2 | 6,4793E-07 | 0,32700134 | 0,717 | 0,614 | 0,02091857 | 12 |
| Apoe | 6,6426E-07 | 0,34466083 | 0,957 | 0,899 | 0,02144549 | 12 |
| Gpr158 | 6,6971E-07 | 0,3796678 | 0,9 | 0,856 | 0,02162144 | 12 |
| Otud7a | 6,9554E-07 | 0,34832609 | 0,961 | 0,919 | 0,02245537 | 12 |
| Nrcam | 7,0377E-07 | 0,33450929 | 0,964 | 0,957 | 0,02272116 | 12 |
| Chrna7 | 7,191E-07 | 0,3437679 | 0,606 | 0,467 | 0,02321614 | 12 |
| Tbc1d5 | 7,2242E-07 | 0,36382705 | 0,778 | 0,605 | 0,02332333 | 12 |
| Atl1 | 7,377E-07 | 0,29956365 | 0,455 | 0,32 | 0,02381679 | 12 |
| Rplp1 | 7,4187E-07 | 0,3582504 | 0,81 | 0,784 | 0,02395119 | 12 |
| Shank1 | 7,4246E-07 | 0,34342953 | 0,613 | 0,47 | 0,02397042 | 12 |
| R3hdm1 | 7,4526E-07 | 0,31974153 | 0,943 | 0,905 | 0,02406075 | 12 |
| Lrrn2 | 7,6076E-07 | 0,3251782 | 0,706 | 0,588 | 0,02456127 | 12 |
| Iqsec2 | 8,3652E-07 | 0,33860205 | 0,789 | 0,692 | 0,0270072 | 12 |
| Prkg1 | 8,7958E-07 | 0,33830029 | 0,989 | 0,988 | 0,02839722 | 12 |
| Pitpnm2 | 8,9367E-07 | 0,32982504 | 0,842 | 0,79 | 0,02885219 | 12 |
| Gsk3b | 9,1203E-07 | 0,34241553 | 0,703 | 0,611 | 0,02944478 | 12 |
| Prdm5 | 9,5808E-07 | 0,34807742 | 0,699 | 0,562 | 0,03093152 | 12 |
| Rgs7 | 1,0628E-06 | 0,33365364 | 0,986 | 0,968 | 0,03431376 | 12 |
| Rfx7 | 1,0966E-06 | 0,36552943 | 0,62 | 0,513 | 0,03540385 | 12 |
| Ccnjl | 1,1131E-06 | 0,33673428 | 0,606 | 0,429 | 0,03593526 | 12 |
| Sergef | 1,1451E-06 | 0,30649661 | 0,53 | 0,403 | 0,03696897 | 12 |
| Kirrel3 | 1,1551E-06 | 0,31623416 | 1 | 0,997 | 0,03729307 | 12 |
| Cadm2 | 1,1648E-06 | 0,35850824 | 1 | 1 | 0,03760453 | 12 |

| | | | | | | |
|---|---|---|---|---|---|---|
| Agap2 | 1,2298E-06 | 0,30277196 | 0,466 | 0,311 | 0,03970265 | 12 |
| Celf4 | 1,2832E-06 | 0,31753872 | 0,946 | 0,957 | 0,04142663 | 12 |
| Trerf1 | 1,2871E-06 | 0,32742655 | 0,903 | 0,821 | 0,04155291 | 12 |
| Atrx | 1,35E-06 | 0,32422485 | 0,806 | 0,755 | 0,04358478 | 12 |
| Sez6l2 | 1,3554E-06 | 0,31881622 | 0,706 | 0,594 | 0,04375872 | 12 |
| Cacna1c | 1,3561E-06 | 0,29567039 | 0,993 | 0,997 | 0,04378042 | 12 |
| Rbfox1 | 1,3618E-06 | 0,31380743 | 1 | 1 | 0,04396473 | 12 |
| Ppm1h | 1,389E-06 | 0,33304907 | 0,749 | 0,651 | 0,04484429 | 12 |
| Prr16 | 1,4071E-06 | 0,48421832 | 0,821 | 0,738 | 0,04542981 | 12 |
| Ralgps1 | 1,4158E-06 | 0,32984553 | 0,677 | 0,556 | 0,04570932 | 12 |
| Schip1 | 1,5078E-06 | 0,3569891 | 0,971 | 0,945 | 0,04867848 | 12 |
| Pde2a | 1,5085E-06 | 0,30604024 | 0,47 | 0,354 | 0,04870197 | 12 |
| Mmp16 | 1,6015E-06 | 0,35645101 | 0,953 | 0,968 | 0,05170362 | 12 |
| Sgsm2 | 1,6031E-06 | 0,30693489 | 0,738 | 0,646 | 0,05175493 | 12 |
| Pclo | 1,6634E-06 | 0,36110611 | 0,892 | 0,844 | 0,05370325 | 12 |
| Kcnd2 | 1,7645E-06 | 0,32749766 | 1 | 1 | 0,05696815 | 12 |
| Cers6 | 1,7877E-06 | 0,32944423 | 0,9 | 0,862 | 0,05771707 | 12 |
| Grid1 | 1,8799E-06 | 0,35199769 | 0,95 | 0,939 | 0,06069261 | 12 |
| Nedd4l | 1,9059E-06 | 0,33312745 | 0,925 | 0,928 | 0,06153307 | 12 |
| Atxn1 | 1,9205E-06 | 0,33846811 | 0,95 | 0,934 | 0,06200485 | 12 |
| Sorbs1 | 1,9444E-06 | 0,30717113 | 0,964 | 0,916 | 0,06277595 | 12 |
| Gabrg3 | 1,9501E-06 | 0,418712 | 0,602 | 0,484 | 0,06295826 | 12 |
| Rere | 1,9515E-06 | 0,30364913 | 0,968 | 0,96 | 0,06300438 | 12 |
| Ptprn2 | 1,9525E-06 | 0,32500425 | 0,907 | 0,847 | 0,06303493 | 12 |
| Enox1 | 2,0308E-06 | 0,32734437 | 0,964 | 0,928 | 0,06556354 | 12 |
| Stag1 | 2,1126E-06 | 0,33410834 | 0,932 | 0,89 | 0,0682038 | 12 |
| Mical2 | 2,135E-06 | 0,30987753 | 0,896 | 0,847 | 0,06892742 | 12 |
| Med12l | 2,1485E-06 | 0,33713788 | 0,792 | 0,741 | 0,06936411 | 12 |
| Nr3c2 | 2,1726E-06 | 0,36775501 | 0,982 | 0,971 | 0,07014277 | 12 |
| Pde4dip | 2,1873E-06 | 0,32283291 | 0,932 | 0,882 | 0,07061539 | 12 |
| Dmd | 2,2409E-06 | 0,37484378 | 0,839 | 0,807 | 0,07234846 | 12 |
| Spock3 | 2,2459E-06 | 0,39306561 | 0,778 | 0,726 | 0,07250735 | 12 |
| Fbxw11 | 2,2893E-06 | 0,30752257 | 0,631 | 0,501 | 0,07391014 | 12 |
| Trim2 | 2,2927E-06 | 0,32130363 | 0,918 | 0,888 | 0,0740184 | 12 |
| Ablim3 | 2,3177E-06 | 0,31467054 | 0,71 | 0,582 | 0,07482634 | 12 |
| Edil3 | 2,3207E-06 | 0,39138551 | 0,889 | 0,888 | 0,074924 | 12 |
| Mrtfa | 2,4221E-06 | 0,33281789 | 0,81 | 0,715 | 0,07819644 | 12 |
| Eif4g3 | 2,4265E-06 | 0,32833253 | 0,849 | 0,827 | 0,07834082 | 12 |
| Kcnd3 | 2,4302E-06 | 0,33391431 | 0,907 | 0,862 | 0,07845755 | 12 |
| Dnmt3a | 2,4884E-06 | 0,31855921 | 0,731 | 0,625 | 0,08033683 | 12 |
| Nrxn1 | 2,5131E-06 | 0,32251662 | 1 | 1 | 0,08113689 | 12 |
| Slc8a3 | 2,5272E-06 | 0,3115713 | 0,405 | 0,259 | 0,08159044 | 12 |
| Znrf3 | 2,5292E-06 | 0,32792664 | 0,67 | 0,513 | 0,08165671 | 12 |
| Miat | 2,5344E-06 | 0,31467054 | 0,846 | 0,818 | 0,08182169 | 12 |
| Oxr1 | 2,6024E-06 | 0,35048568 | 0,903 | 0,882 | 0,08401957 | 12 |
| Rnf150 | 2,6054E-06 | 0,35050265 | 0,659 | 0,539 | 0,08411393 | 12 |
| Hivep2 | 2,6281E-06 | 0,31888074 | 0,968 | 0,963 | 0,08484961 | 12 |
| Fgf14 | 2,6371E-06 | 0,32328882 | 1 | 1 | 0,08513921 | 12 |

| | | | | | | |
|---|---|---|---|---|---|---|
| Rnf165 | 2,6698E-06 | 0,32201252 | 0,588 | 0,45 | 0,08619389 | 12 |
| Adcy8 | 2,7026E-06 | 0,32342301 | 0,466 | 0,294 | 0,08725372 | 12 |
| [illegible]730522E02Ri | 2,8538E-06 | 0,35541456 | 0,928 | 0,928 | 0,09213407 | 12 |
| Phf21a | 2,861E-06 | 0,30157488 | 0,86 | 0,798 | 0,09236641 | 12 |
| Kifap3 | 2,8844E-06 | 0,29498817 | 0,771 | 0,657 | 0,09312366 | 12 |
| Gbf1 | 3,0047E-06 | 0,31688837 | 0,652 | 0,527 | 0,09700641 | 12 |
| Trim9 | 3,0355E-06 | 0,33473927 | 0,896 | 0,859 | 0,09800034 | 12 |
| Kcnb1 | 3,0712E-06 | 0,30271774 | 0,642 | 0,504 | 0,09915495 | 12 |
| Gm47283 | 4,8415E-37 | -1,0544224 | 0,38 | 0,752 | 1,5631E-32 | 13 |
| Bc1 | 2,1424E-20 | 0,98063655 | 0,806 | 0,632 | 6,9166E-16 | 13 |
| Gm13269 | 2,7463E-07 | -0,4048819 | 0,654 | 0,808 | 0,00886656 | 13 |
| Ttr | 1,1172E-06 | -0,2631733 | 0,053 | 0,176 | 0,03606851 | 13 |
| Bc1 | 1,0329E-29 | 1,21521984 | 0,693 | 0,376 | 3,3348E-25 | 14 |
| Gm47283 | 9,2265E-14 | -0,5644219 | 0,343 | 0,57 | 2,9788E-09 | 14 |
| AC149090.1 | 7,5671E-11 | 0,64927563 | 0,896 | 0,855 | 2,443E-06 | 14 |
| Gm19951 | 2,8048E-07 | 0,35847375 | 0,291 | 0,128 | 0,00905515 | 14 |
| Kif1b | 3,3959E-07 | 0,41938813 | 0,649 | 0,496 | 0,01096364 | 14 |
| mt-Atp6 | 9,5628E-07 | 0,51479476 | 0,98 | 0,963 | 0,03087355 | 14 |
| Ttr | 1,2075E-06 | -0,2634474 | 0,056 | 0,211 | 0,03898431 | 14 |
| Usp31 | 2,2535E-06 | 0,2526966 | 0,239 | 0,079 | 0,07275343 | 14 |
| Bc1 | 6,0476E-24 | 1,17871417 | 0,68 | 0,31 | 1,9525E-19 | 16 |
| Gm47283 | 2,4455E-08 | -0,4830688 | 0,258 | 0,44 | 0,00078954 | 16 |
| Ikzf1 | 2,8333E-07 | 0,41911286 | 0,674 | 0,547 | 0,00914747 | 16 |
| Ttr | 7,2082E-07 | -0,3063376 | 0,051 | 0,254 | 0,02327172 | 16 |
| Tmem135 | 1,1816E-06 | 0,34217012 | 0,421 | 0,224 | 0,03814716 | 16 |
| Bc1 | 5,444E-22 | 1,43651723 | 0,873 | 0,551 | 1,7576E-17 | 21 |
| Msi2 | 4,8432E-19 | -1,6953299 | 0,508 | 0,83 | 1,5636E-14 | 21 |
| Dclk1 | 4,6572E-18 | 1,31483867 | 0,968 | 0,714 | 1,5036E-13 | 21 |
| Hivep2 | 5,0448E-17 | 1,18194038 | 1 | 0,707 | 1,6287E-12 | 21 |
| Lrrtm3 | 3,4146E-16 | 1,4846003 | 0,889 | 0,429 | 1,1024E-11 | 21 |
| Pdzrn3 | 7,3627E-16 | 1,61928257 | 0,635 | 0,095 | 2,3771E-11 | 21 |
| Kcnb2 | 1,8181E-15 | 1,53243096 | 0,952 | 0,68 | 5,8698E-11 | 21 |
| Nwd2 | 4,3707E-15 | -2,068032 | 0,444 | 0,694 | 1,4111E-10 | 21 |
| Rbms3 | 1,2364E-14 | -1,681799 | 0,222 | 0,789 | 3,9918E-10 | 21 |
| Camk2a | 5,1059E-14 | 1,17416414 | 0,889 | 0,66 | 1,6484E-09 | 21 |
| Sipa1l1 | 1,101E-13 | 1,12765193 | 0,81 | 0,34 | 3,5547E-09 | 21 |
| Gm49678 | 1,4703E-13 | 1,11896283 | 0,492 | 0,061 | 4,7469E-09 | 21 |
| Ext1 | 1,6363E-13 | 1,30064387 | 0,905 | 0,49 | 5,2828E-09 | 21 |
| Dscaml1 | 2,2248E-13 | 1,13027422 | 0,667 | 0,19 | 7,1827E-09 | 21 |
| Tcerg1l | 2,3249E-13 | 0,85725983 | 0,587 | 0,061 | 7,5058E-09 | 21 |
| Ptprd | 3,1536E-13 | 1,40320236 | 0,984 | 0,966 | 1,0181E-08 | 21 |
| Vav3 | 3,4837E-13 | -1,5552152 | 0,159 | 0,66 | 1,1247E-08 | 21 |
| Gpm6b | 4,3007E-13 | 1,03052487 | 0,825 | 0,449 | 1,3885E-08 | 21 |
| Csmd1 | 9,5286E-13 | 1,83509991 | 1 | 0,748 | 3,0763E-08 | 21 |
| Phactr1 | 1,0916E-12 | 1,75461346 | 0,984 | 0,558 | 3,5241E-08 | 21 |
| Sorbs2 | 2,5736E-12 | 1,11518282 | 0,937 | 0,612 | 8,3089E-08 | 21 |
| Dlg2 | 2,7317E-12 | 1,15809874 | 1 | 0,973 | 8,8192E-08 | 21 |
| [illegible]010300C02Ri | 2,9881E-12 | 0,84857758 | 0,603 | 0,204 | 9,6472E-08 | 21 |

| | | | | | | |
|---|---|---|---|---|---|---|
| Celf2 | 4,2931E-12 | 1,56232847 | 1 | 0,898 | 1,386E-07 | 21 |
| Gm42418 | 6,5623E-12 | 1,01227137 | 0,984 | 0,966 | 2,1186E-07 | 21 |
| Gm47283 | 7,0415E-12 | -1,0939221 | 0,476 | 0,81 | 2,2733E-07 | 21 |
| Chsy3 | 7,1795E-12 | 1,39449179 | 0,794 | 0,184 | 2,3179E-07 | 21 |
| Pdzd2 | 8,7064E-12 | 0,89174711 | 0,492 | 0,116 | 2,8109E-07 | 21 |
| Ctnna3 | 8,9271E-12 | 1,48222946 | 0,937 | 0,633 | 2,8821E-07 | 21 |
| Camk4 | 9,8261E-12 | 1,07122835 | 0,635 | 0,177 | 3,1724E-07 | 21 |
| Meis2 | 2,1727E-11 | 1,39497188 | 0,587 | 0,082 | 7,0147E-07 | 21 |
| Gria2 | 2,4342E-11 | 1,0446053 | 1 | 0,912 | 7,8589E-07 | 21 |
| Galntl6 | 2,5315E-11 | -1,6287104 | 0,302 | 0,728 | 8,1729E-07 | 21 |
| Ntrk3 | 3,5554E-11 | 1,03327615 | 0,952 | 0,578 | 1,1479E-06 | 21 |
| Nav2 | 4,2417E-11 | 1,01703181 | 0,952 | 0,646 | 1,3694E-06 | 21 |
| Trpm3 | 4,7132E-11 | -1,6786573 | 0,603 | 0,83 | 1,5217E-06 | 21 |
| Scube1 | 6,8813E-11 | -1,469983 | 0,063 | 0,531 | 2,2216E-06 | 21 |
| Kcnq5 | 7,3719E-11 | 2,16568532 | 0,857 | 0,354 | 2,38E-06 | 21 |
| Cdh20 | 7,8441E-11 | 1,45322991 | 0,73 | 0,177 | 2,5325E-06 | 21 |
| Adgrl3 | 8,1211E-11 | 1,18997094 | 0,968 | 0,741 | 2,6219E-06 | 21 |
| R3hdm1 | 9,4825E-11 | 1,13057171 | 0,921 | 0,612 | 3,0614E-06 | 21 |
| Nrxn3 | 1,0643E-10 | 1,63096128 | 0,968 | 0,741 | 3,436E-06 | 21 |
| Nlgn1 | 1,3775E-10 | 1,26413696 | 0,952 | 0,952 | 4,4473E-06 | 21 |
| Ppp2r2b | 1,3911E-10 | 1,02419969 | 0,952 | 0,524 | 4,4911E-06 | 21 |
| Dkk3 | 1,5432E-10 | 0,87970577 | 0,508 | 0,129 | 4,9823E-06 | 21 |
| Prkca | 1,5454E-10 | 1,34831486 | 0,825 | 0,408 | 4,9892E-06 | 21 |
| Gda | 1,6586E-10 | 0,69564453 | 0,54 | 0,122 | 5,3548E-06 | 21 |
| Plcb1 | 1,7239E-10 | 1,52221633 | 0,905 | 0,381 | 5,5656E-06 | 21 |
| Dlgap2 | 1,7836E-10 | 1,37100692 | 0,937 | 0,401 | 5,7583E-06 | 21 |
| Caln1 | 2,4163E-10 | 0,9987664 | 0,683 | 0,156 | 7,801E-06 | 21 |
| Tmtc1 | 2,5081E-10 | 0,84262025 | 0,698 | 0,374 | 8,0972E-06 | 21 |
| Gramd1b | 2,6891E-10 | 0,75704984 | 0,778 | 0,306 | 8,6816E-06 | 21 |
| Syt9 | 2,7297E-10 | -1,3814291 | 0,111 | 0,51 | 8,8129E-06 | 21 |
| Atp8a1 | 2,9183E-10 | 0,86079418 | 0,667 | 0,293 | 9,4219E-06 | 21 |
| Sntb2 | 2,9472E-10 | 1,11769504 | 0,571 | 0,224 | 9,5149E-06 | 21 |
| Tshz2 | 2,9547E-10 | 2,29342667 | 0,857 | 0,272 | 9,5392E-06 | 21 |
| Dpp10 | 3,6055E-10 | 1,45118508 | 0,857 | 0,844 | 1,164E-05 | 21 |
| Cdh12 | 5,0084E-10 | 1,39875472 | 0,857 | 0,395 | 1,617E-05 | 21 |
| Mapt | 5,3029E-10 | 0,79612767 | 0,889 | 0,639 | 1,7121E-05 | 21 |
| Nav3 | 7,4164E-10 | 1,10959182 | 0,952 | 0,85 | 2,3944E-05 | 21 |
| Ralyl | 8,1936E-10 | 1,25700338 | 0,937 | 0,932 | 2,6453E-05 | 21 |
| Kalrn | 8,207E-10 | 1,48917896 | 0,968 | 0,51 | 2,6496E-05 | 21 |
| Ptk2b | 8,4122E-10 | 0,70436093 | 0,619 | 0,15 | 2,7159E-05 | 21 |
| Pde4b | 9,2744E-10 | 1,18563387 | 0,857 | 0,456 | 2,9942E-05 | 21 |
| Tiam1 | 1,1553E-09 | 0,90923454 | 0,635 | 0,265 | 3,73E-05 | 21 |
| Chn1 | 1,2268E-09 | 0,83615895 | 0,778 | 0,517 | 3,9606E-05 | 21 |
| Ankrd33b | 1,2934E-09 | 0,73534368 | 0,571 | 0,136 | 4,1756E-05 | 21 |
| Prkce | 1,4847E-09 | 0,9422845 | 0,905 | 0,558 | 4,7933E-05 | 21 |
| Gm43507 | 1,6521E-09 | 0,60090404 | 0,444 | 0,075 | 5,3337E-05 | 21 |
| Mdga2 | 1,7319E-09 | 0,95224836 | 0,937 | 0,884 | 5,5915E-05 | 21 |
| Nrgn | 1,7743E-09 | 0,77069645 | 0,857 | 0,524 | 5,7282E-05 | 21 |

| | | | | | | |
|---|---|---|---|---|---|---|
| Ppm1l | 1,915E-09 | 0,89205209 | 0,73 | 0,259 | 6,1827E-05 | 21 |
| Tacc1 | 1,9546E-09 | 0,77059637 | 0,667 | 0,299 | 6,3105E-05 | 21 |
| Egfem1 | 2,4012E-09 | 1,28940604 | 0,921 | 0,707 | 7,7523E-05 | 21 |
| Tshz3 | 2,6891E-09 | 1,03086096 | 0,603 | 0,109 | 8,6818E-05 | 21 |
| Kctd16 | 2,843E-09 | 1,59405037 | 0,937 | 0,374 | 9,1786E-05 | 21 |
| Ppp3ca | 2,9219E-09 | 1,06369468 | 0,937 | 0,68 | 9,4333E-05 | 21 |
| Ntrk2 | 3,2828E-09 | 0,87725694 | 0,857 | 0,463 | 0,00010599 | 21 |
| Cdc14b | 3,3601E-09 | 0,84236137 | 0,571 | 0,204 | 0,00010848 | 21 |
| Ssbp2 | 3,3743E-09 | 0,93008309 | 0,667 | 0,347 | 0,00010894 | 21 |
| Dab1 | 3,8409E-09 | 2,04584303 | 0,952 | 0,429 | 0,000124 | 21 |
| Lmo4 | 3,8557E-09 | 0,76203093 | 0,667 | 0,211 | 0,00012448 | 21 |
| Ephb2 | 3,8965E-09 | 0,62004784 | 0,492 | 0,102 | 0,0001258 | 21 |
| Pde4a | 5,2077E-09 | 0,66335839 | 0,619 | 0,259 | 0,00016813 | 21 |
| Rapgef2 | 5,5392E-09 | 0,81839319 | 0,667 | 0,313 | 0,00017883 | 21 |
| Vsnl1 | 5,8642E-09 | 0,87611447 | 0,825 | 0,313 | 0,00018933 | 21 |
| Ldb2 | 5,9239E-09 | 1,52061626 | 0,857 | 0,197 | 0,00019125 | 21 |
| Rgs7 | 5,9829E-09 | 1,20391976 | 0,952 | 0,51 | 0,00019316 | 21 |
| Zfyve28 | 5,9927E-09 | 0,6291623 | 0,54 | 0,15 | 0,00019347 | 21 |
| Snap25 | 6,3636E-09 | 0,86612233 | 0,905 | 0,769 | 0,00020545 | 21 |
| Pcdh15 | 7,1753E-09 | 1,57911536 | 0,857 | 0,469 | 0,00023165 | 21 |
| Flrt1 | 7,3401E-09 | 0,58193481 | 0,429 | 0,061 | 0,00023697 | 21 |
| Crtac1 | 7,897E-09 | 0,66599907 | 0,54 | 0,163 | 0,00025495 | 21 |
| Cacna2d1 | 8,8005E-09 | 1,2425703 | 0,762 | 0,354 | 0,00028412 | 21 |
| Tcf4 | 1,0424E-08 | 1,29532394 | 0,841 | 0,537 | 0,00033654 | 21 |
| Camk1d | 1,0507E-08 | 0,83967264 | 0,857 | 0,524 | 0,00033922 | 21 |
| Gm19410 | 1,1078E-08 | 0,64670773 | 0,365 | 0,041 | 0,00035765 | 21 |
| Homer1 | 1,1234E-08 | 0,5849625 | 0,571 | 0,184 | 0,0003627 | 21 |
| Cacnb4 | 1,2337E-08 | 1,00491549 | 0,841 | 0,435 | 0,00039831 | 21 |
| Rfx3 | 1,3536E-08 | 0,99051895 | 0,714 | 0,293 | 0,00043702 | 21 |
| 810034E14Ri | 1,4824E-08 | 0,73400318 | 0,651 | 0,197 | 0,00047859 | 21 |
| Rapgef5 | 1,5075E-08 | 0,83144054 | 0,651 | 0,204 | 0,00048669 | 21 |
| Gabrb3 | 1,8629E-08 | 0,8593938 | 0,952 | 0,776 | 0,00060143 | 21 |
| Kcnh1 | 1,9138E-08 | 0,6209418 | 0,54 | 0,136 | 0,00061788 | 21 |
| Ttc3 | 1,983E-08 | 0,7457004 | 0,984 | 0,803 | 0,0006402 | 21 |
| Dlg1 | 2,037E-08 | 0,81623307 | 0,683 | 0,272 | 0,00065764 | 21 |
| Ank | 2,0732E-08 | 0,6412215 | 0,651 | 0,19 | 0,00066933 | 21 |
| Robo2 | 2,1058E-08 | 1,22764815 | 0,635 | 0,395 | 0,00067986 | 21 |
| Lncpint | 2,9179E-08 | 0,74909552 | 0,889 | 0,687 | 0,00094206 | 21 |
| Ank2 | 3,0299E-08 | 0,81785533 | 0,968 | 0,905 | 0,0009782 | 21 |
| Nrg1 | 3,1083E-08 | 1,23817699 | 0,905 | 0,803 | 0,00100352 | 21 |
| Rap1gds1 | 3,2758E-08 | 0,78226394 | 0,762 | 0,347 | 0,00105758 | 21 |
| Npas2 | 3,3692E-08 | 0,7419605 | 0,476 | 0,116 | 0,00108774 | 21 |
| Rasgef1a | 3,4345E-08 | 0,67377177 | 0,587 | 0,252 | 0,00110884 | 21 |
| Kcnma1 | 3,5112E-08 | -1,2343818 | 0,937 | 0,918 | 0,0011336 | 21 |
| Schip1 | 3,5767E-08 | 1,4647334 | 0,81 | 0,735 | 0,00115473 | 21 |
| Grm5 | 3,7641E-08 | 1,17224631 | 0,952 | 0,687 | 0,00121524 | 21 |
| Satb2 | 3,854E-08 | 0,70781925 | 0,381 | 0,054 | 0,00124427 | 21 |
| 130073E24Ri | 4,0429E-08 | -1,2511488 | 0,095 | 0,429 | 0,00130525 | 21 |

| | | | | | | |
|---|---|---|---|---|---|---|
| Gm3294 | 4,1794E-08 | 0,50085687 | 0,381 | 0,048 | 0,00134933 | 21 |
| Spon1 | 4,8126E-08 | 0,51189904 | 0,444 | 0,088 | 0,00155376 | 21 |
| Gm2164 | 4,8575E-08 | 0,91795175 | 0,651 | 0,197 | 0,00156824 | 21 |
| Gm49906 | 5,2212E-08 | 1,0730148 | 0,587 | 0,075 | 0,00168566 | 21 |
| Cap2 | 5,5072E-08 | 0,78181983 | 0,667 | 0,34 | 0,001778 | 21 |
| Sobp | 6,1546E-08 | 0,89675401 | 0,794 | 0,469 | 0,00198702 | 21 |
| Nbea | 6,7125E-08 | 0,77640633 | 0,984 | 0,83 | 0,00216711 | 21 |
| Slco3a1 | 7,3103E-08 | 0,6918777 | 0,492 | 0,286 | 0,00236014 | 21 |
| Dapk1 | 7,3355E-08 | 0,74052341 | 0,762 | 0,367 | 0,00236827 | 21 |
| Tspan5 | 7,7561E-08 | 0,91457009 | 0,635 | 0,32 | 0,00250407 | 21 |
| Raver2 | 8,1624E-08 | 0,57527344 | 0,429 | 0,109 | 0,00263522 | 21 |
| :130071C03Ri | 8,8308E-08 | 0,57853623 | 0,651 | 0,272 | 0,00285103 | 21 |
| Unc5d | 8,8818E-08 | 1,63576687 | 0,841 | 0,347 | 0,00286749 | 21 |
| Arhgap39 | 9,0258E-08 | 0,6409444 | 0,603 | 0,32 | 0,00291397 | 21 |
| Cdc42bpa | 9,569E-08 | 0,75032398 | 0,714 | 0,456 | 0,00308935 | 21 |
| 921534H16Ri | 1,0979E-07 | 0,55401391 | 0,556 | 0,15 | 0,00354472 | 21 |
| Sorbs2os | 1,1219E-07 | 0,78276928 | 0,651 | 0,279 | 0,00362218 | 21 |
| Bcl11a | 1,1498E-07 | 0,85982234 | 0,762 | 0,293 | 0,00371202 | 21 |
| Msra | 1,1586E-07 | 1,09541957 | 0,762 | 0,327 | 0,00374048 | 21 |
| Grin2b | 1,2841E-07 | 1,13842138 | 0,968 | 0,667 | 0,00414572 | 21 |
| Fam110b | 1,2923E-07 | 0,53187575 | 0,397 | 0,088 | 0,00417213 | 21 |
| Lhfpl3 | 1,5011E-07 | -1,1769402 | 0,27 | 0,755 | 0,00484643 | 21 |
| Adcy9 | 1,5913E-07 | 0,6338721 | 0,571 | 0,279 | 0,00513737 | 21 |
| Calm1 | 1,6254E-07 | 0,59162623 | 0,968 | 0,844 | 0,0052477 | 21 |
| Dach1 | 1,7489E-07 | -1,0013553 | 0,063 | 0,476 | 0,00564622 | 21 |
| Slc4a4 | 1,8198E-07 | 0,78024969 | 0,54 | 0,218 | 0,00587533 | 21 |
| Sh3rf3 | 1,8838E-07 | 0,64134497 | 0,524 | 0,177 | 0,00608201 | 21 |
| Kcnip3 | 1,8902E-07 | 0,60151035 | 0,476 | 0,204 | 0,00610249 | 21 |
| Sorcs1 | 1,9462E-07 | 1,45456586 | 0,365 | 0,15 | 0,00628338 | 21 |
| Cdh13 | 1,9481E-07 | 0,9851672 | 0,778 | 0,361 | 0,00628951 | 21 |
| Csrnp3 | 1,9944E-07 | 0,81274818 | 0,81 | 0,381 | 0,00643879 | 21 |
| March1 | 2,3098E-07 | 1,34642346 | 0,889 | 0,367 | 0,00745734 | 21 |
| Trim9 | 2,3408E-07 | 0,71384304 | 0,73 | 0,388 | 0,00755737 | 21 |
| Psd3 | 2,3912E-07 | 0,70282434 | 0,73 | 0,422 | 0,00772001 | 21 |
| Cntn5 | 2,6395E-07 | 1,42037752 | 0,889 | 0,66 | 0,00852154 | 21 |
| Mical2 | 2,6396E-07 | 0,80519681 | 0,683 | 0,279 | 0,00852187 | 21 |
| Gabrb1 | 2,9388E-07 | 1,07617168 | 0,905 | 0,517 | 0,00948797 | 21 |
| Grip1 | 3,05E-07 | -0,8977995 | 0,492 | 0,823 | 0,00984707 | 21 |
| Cacna1a | 3,0993E-07 | 1,07555103 | 0,937 | 0,449 | 0,01000606 | 21 |
| St7 | 3,1535E-07 | 0,54432052 | 0,635 | 0,265 | 0,01018115 | 21 |
| Gm48742 | 3,1741E-07 | 0,55942741 | 0,46 | 0,122 | 0,01024774 | 21 |
| Tspan7 | 3,1784E-07 | 0,61233894 | 0,905 | 0,701 | 0,01026153 | 21 |
| Arhgef3 | 3,2156E-07 | 0,62449086 | 0,429 | 0,095 | 0,01038163 | 21 |
| Ly6h | 3,3245E-07 | 0,6520767 | 0,81 | 0,578 | 0,01073311 | 21 |
| Cnksr2 | 3,5073E-07 | 0,92494042 | 0,762 | 0,354 | 0,01132316 | 21 |
| Lzts1 | 3,569E-07 | 0,68038207 | 0,524 | 0,15 | 0,01152244 | 21 |
| Atp2b4 | 3,934E-07 | 0,57288967 | 0,349 | 0,068 | 0,01270104 | 21 |
| Ndst3 | 3,9495E-07 | 0,79672337 | 0,714 | 0,279 | 0,01275088 | 21 |

| | | | | | | |
|---|---|---|---|---|---|---|
| Clstn2 | 3,9889E-07 | 1,15180045 | 0,841 | 0,435 | 0,01287807 | 21 |
| Gphn | 4,1015E-07 | 0,70730775 | 0,857 | 0,721 | 0,01324177 | 21 |
| Hs6st3 | 4,2343E-07 | 1,62574812 | 0,81 | 0,422 | 0,01367058 | 21 |
| Mrtfb | 4,6365E-07 | 0,67974073 | 0,73 | 0,449 | 0,01496897 | 21 |
| Atrx | 4,7024E-07 | 0,59911299 | 0,746 | 0,469 | 0,01518182 | 21 |
| Kctd8 | 4,8155E-07 | -1,2110049 | 0,254 | 0,653 | 0,01554683 | 21 |
| Dleu2 | 5,1631E-07 | 0,62535709 | 0,46 | 0,17 | 0,01666922 | 21 |
| Kcnq3 | 5,9361E-07 | 1,00231758 | 0,825 | 0,395 | 0,01916478 | 21 |
| [illegible]730522E02Ri | 6,0558E-07 | 1,24739643 | 0,81 | 0,252 | 0,01955118 | 21 |
| Dip2b | 6,1048E-07 | 0,65344224 | 0,587 | 0,388 | 0,01970946 | 21 |
| Arntl | 6,1158E-07 | 0,5126483 | 0,476 | 0,156 | 0,01974495 | 21 |
| Gas7 | 6,2746E-07 | 0,65328387 | 0,476 | 0,163 | 0,02025767 | 21 |
| Homer2 | 6,3742E-07 | 0,64620013 | 0,746 | 0,415 | 0,02057921 | 21 |
| Mir99ahg | 6,5012E-07 | 0,89196715 | 0,778 | 0,442 | 0,02098904 | 21 |
| Arhgef28 | 6,5404E-07 | 0,53726576 | 0,365 | 0,095 | 0,02111569 | 21 |
| Slc16a2 | 6,8345E-07 | 0,51428169 | 0,413 | 0,095 | 0,02206503 | 21 |
| Lingo2 | 6,8908E-07 | 1,44726583 | 0,937 | 0,592 | 0,0222471 | 21 |
| Gria3 | 6,9656E-07 | 1,09813432 | 0,857 | 0,503 | 0,02248849 | 21 |
| Ppfia2 | 8,1371E-07 | 0,68549658 | 0,905 | 0,755 | 0,02627051 | 21 |
| Otud7a | 8,6916E-07 | 0,67949398 | 0,905 | 0,592 | 0,02806087 | 21 |
| Acap2 | 8,7834E-07 | 0,69494238 | 0,603 | 0,265 | 0,02835737 | 21 |
| Tcf20 | 8,79E-07 | 0,57213846 | 0,698 | 0,388 | 0,02837854 | 21 |
| Fam171b | 8,8793E-07 | 0,53865177 | 0,651 | 0,279 | 0,02866669 | 21 |
| Osbpl6 | 9,2877E-07 | 0,71424552 | 0,778 | 0,395 | 0,02998523 | 21 |
| Grm7 | 9,7609E-07 | 1,42845802 | 0,984 | 0,51 | 0,03151297 | 21 |
| Syn2 | 9,9596E-07 | 0,7728165 | 0,825 | 0,517 | 0,03215447 | 21 |
| Pde10a | 1,0952E-06 | 1,09548031 | 0,905 | 0,442 | 0,0353575 | 21 |
| Fntb | 1,1115E-06 | 0,50618539 | 0,444 | 0,245 | 0,03588526 | 21 |
| Ahi1 | 1,1273E-06 | 0,67147149 | 0,937 | 0,918 | 0,03639355 | 21 |
| Srgap3 | 1,2028E-06 | 0,71628534 | 0,73 | 0,449 | 0,03883322 | 21 |
| Khdrbs2 | 1,2105E-06 | 1,16405911 | 0,825 | 0,531 | 0,03908223 | 21 |
| Sgcz | 1,2392E-06 | 1,29460046 | 0,698 | 0,66 | 0,0400081 | 21 |
| Limch1 | 1,2639E-06 | 0,6576078 | 0,603 | 0,293 | 0,04080398 | 21 |
| Rbfox3 | 1,2891E-06 | 0,70974413 | 0,905 | 0,531 | 0,04161762 | 21 |
| Tmem117 | 1,3442E-06 | -0,659956 | 0,238 | 0,599 | 0,04339613 | 21 |
| Wscd2 | 1,359E-06 | 0,53114513 | 0,476 | 0,15 | 0,04387683 | 21 |
| Nrp1 | 1,4369E-06 | 0,6727054 | 0,524 | 0,272 | 0,04639157 | 21 |
| L3mbtl3 | 1,4445E-06 | 0,46133115 | 0,365 | 0,082 | 0,04663429 | 21 |
| Rora | 1,5661E-06 | 1,11413953 | 1 | 0,728 | 0,05056172 | 21 |
| Mbnl1 | 1,8383E-06 | 0,5987683 | 0,571 | 0,313 | 0,05934819 | 21 |
| Trerf1 | 1,8484E-06 | 0,67349918 | 0,73 | 0,354 | 0,05967539 | 21 |
| Astn2 | 2,0413E-06 | 0,8442545 | 0,825 | 0,619 | 0,06590212 | 21 |
| Gm10115 | 2,1646E-06 | 0,47643804 | 0,333 | 0,082 | 0,06988474 | 21 |
| Cntnap2 | 2,2389E-06 | -0,8312887 | 0,952 | 0,993 | 0,07228133 | 21 |
| Camta1 | 2,3371E-06 | 0,82043475 | 0,889 | 0,578 | 0,0754539 | 21 |
| Smad3 | 2,3903E-06 | 0,47279922 | 0,254 | 0,041 | 0,07717203 | 21 |
| Auts2 | 2,4116E-06 | 0,81833756 | 0,984 | 0,952 | 0,07785814 | 21 |
| Plxna2 | 2,4562E-06 | 0,5930358 | 0,54 | 0,259 | 0,07929924 | 21 |

| | | | | | | |
|---|---|---|---|---|---|---|
| Man1a2 | 2,4744E-06 | 0,54568735 | 0,587 | 0,333 | 0,07988702 | 21 |
| Galnt18 | 2,5314E-06 | 0,74385198 | 0,603 | 0,245 | 0,08172687 | 21 |
| Wipf3 | 2,5813E-06 | 0,58902529 | 0,54 | 0,245 | 0,08333842 | 21 |
| Edil3 | 2,6003E-06 | 0,80316713 | 0,619 | 0,286 | 0,08394959 | 21 |
| Dync1i2 | 2,6242E-06 | 0,56342934 | 0,651 | 0,476 | 0,08472091 | 21 |
| Arpp21 | 2,6387E-06 | 1,25626006 | 0,905 | 0,381 | 0,08518904 | 21 |
| Usp9x | 2,7907E-06 | 0,54559105 | 0,603 | 0,429 | 0,0900978 | 21 |
| Dip2c | 2,8332E-06 | 0,59571967 | 0,937 | 0,735 | 0,09147093 | 21 |
| Tmem163 | 2,8948E-06 | -0,8008399 | 0,111 | 0,469 | 0,0934589 | 21 |
| Plcl2 | 2,9884E-06 | 0,55496776 | 0,587 | 0,361 | 0,09648031 | 21 |
| Lyst | 3,0466E-06 | 0,45606477 | 0,508 | 0,197 | 0,09836011 | 21 |
