## Supplementary Tables 3-6 for "Disruption of autism-associated *Pcdh9* gene leads to transcriptional alterations, synapses overgrowth and aberrant excitatory transmission in the CA1"

**Supplementary Table 3**

| |
|---|
| Apoe |
| Bc1 |
| Camk2a |
| Csmd2 |
| Cttn |
| Dapk1 |
| Ddx5 |
| Dnaja2 |
| Dnmt3a |
| Frmd4a |
| Galnt9 |
| Gm10848 |
| Gm13269 |
| Gm21798 |
| Gm36264 |
| Gm47283 |
| Ksr1 |
| Larp1 |
| Meg3 |
| Olfm1 |
| Pabpc1 |
| Pcdh9 |
| Pisd (AC149090.1) |
| Pkp2 |
| Rps8 |
| Shank2 |
| Snhg11 |
| Specc1 |
| Sv2b |
| Ttr |
| 4930415C11Rik |
| 9530059O14Rik |

#### Supplementary Table 4

| #node1 | node2 | ode1_string_i | ode2_string_i | hood_on_chro | gene_fusion | enetic_coocc | homology | coexpression | ly_determine | abase_annota | nated_textm | mbined_score |
|---|---|---|---|---|---|---|---|---|---|---|---|---|
| APOE | TTR | ENSP0000025 | ENSP0000023 | 0 | 0 | 0 | 0 | 0 | 0 | 0.500 | 0.903 | 0.949 |
| CAMK2A | SHANK2 | ENSP000005( | ENSP0000034 | 0 | 0 | 0 | 0 | 0 | 0.086 | 0 | 0.399 | 0.427 |
| CAMK2A | SV2B | ENSP000005( | ENSP0000037 | 0 | 0 | 0 | 0 | 0 | 0 | 0 | 0.480 | 0.480 |
| CAMK2A | KSR1 | ENSP000005( | ENSP0000038 | 0 | 0 | 0 | 0.560 | 0 | 0.045 | 0.400 | 0.043 | 0.403 |
| CTTN | SHANK2 | ENSP000003( | ENSP0000034 | 0 | 0 | 0 | 0 | 0 | 0.311 | 0 | 0.957 | 0.969 |
| DAPK1 | DNMT3A | ENSP0000038 | ENSP0000026 | 0 | 0 | 0 | 0 | 0 | 0.069 | 0 | 0.415 | 0.432 |
| DDX5 | PABPC1 | ENSP0000022 | ENSP0000031 | 0 | 0 | 0 | 0 | 0 | 0.127 | 0 | 0.623 | 0.656 |
| LARP1 | PABPC1 | ENSP0000033 | ENSP0000031 | 0 | 0 | 0 | 0 | 0 | 0.409 | 0 | 0.845 | 0.904 |
| SHANK2 | SV2B | ENSP0000034 | ENSP0000037 | 0 | 0 | 0 | 0 | 0 | 0 | 0 | 0.460 | 0.459 |

#### Supplementary Table 5

| source | term_name | term_id | highlighted | adjusted_p_value | _log10_of_adjusted_ | term_size | query_size | tersection_si | tive_domain | intersections |
|---|---|---|---|---|---|---|---|---|---|---|
| GO:BP | cellular component maintenance | GO:0043954 | true | 0,002925863 | 2,533746082 | 95 | 25 | 4 | 26856 | APOE,CTTN,PKP2,SHANK2 |
| GO:CC | glutamatergic synapse | GO:0098978 | true | 0,000122154 | 3,913091811 | 612 | 24 | 7 | 26959 | APOE,CAMK2A,CTTN,DAPK1,OLFM1,SHANK2,SV2B |
| GO:CC | cell junction | GO:0030054 | false | 0,001916504 | 2,717490232 | 2275 | 24 | 10 | 26959 | APOE,CAMK2A,CTTN,DAPK1,FRMD4A,OLFM1,PABPC1,PKP2,SHANK2,SV2B |
| GO:CC | synapse | GO:0045202 | false | 0,010089288 | 1,996139485 | 1660 | 24 | 8 | 26959 | APOE,CAMK2A,CTTN,DAPK1,OLFM1,PABPC1,SHANK2,SV2B |
| GO:CC | supramolecular complex | GO:0099080 | true | 0,026620379 | 1,574785765 | 1404 | 24 | 7 | 26959 | APOE,CTTN,LARP1,PABPC1,PKP2,SHANK2,SPECC1 |
| TF | Factor: KROX | | | | | | | | | |
| CORUM | Ctnn-Glua2 complex | CORUM:7414 | false | 0,049957536 | 1,301398989 | 1 | 6 | 1 | 1082 | CTTN |

### Supplementary Table 6

| your genelist input | gene hgnc_id | gene symbol | gene name | gene synonyms | GO term ID | GO term name | GO domain | SynGO annotation ID |
|---|---|---|---|---|---|---|---|---|
| CTTN | HGNC:3338 | CTTN | cortactin | EMS1 | GO:0098871 | postsynaptic actin cytoskeleton (GO:0098871) | CC | 223 |
| APOE | HGNC:613 | APOE | apolipoprotein E | AD2 | GO:0043083 | synaptic cleft (GO:0043083) | CC | 405 |
| SHANK2 | HGNC:14295 | SHANK2 | SH3 and multiple ankyrin repeat domains 2 | CTTNBP1,PROSAP1,SHANK,SPANK-3,CORTBP1 | GO:0014069 | postsynaptic density (GO:0014069) | CC | 522 |
| PABPC1 | HGNC:8554 | PABPC1 | poly(A) binding protein cytoplasmic 1 | PAB1,PABPC2 | GO:0045202 | synapse (GO:0045202) | CC | 565 |
| CTTN | HGNC:3338 | CTTN | cortactin | EMS1 | GO:0098885 | modification of postsynaptic actin cytoskeleton (GO:0098885) | BP | 766 |
| CAMK2A | HGNC:1460 | CAMK2A | calcium/calmodulin dependent protein kinase II alpha | KIAA0968,CAMKIIA,CAMKA | GO:0099523 | presynaptic cytosol (GO:0099523) | CC | 886 |
| CAMK2A | HGNC:1460 | CAMK2A | calcium/calmodulin dependent protein kinase II alpha | KIAA0968,CAMKIIA,CAMKA | GO:0099524 | postsynaptic cytosol (GO:0099524) | CC | 887 |
| CAMK2A | HGNC:1460 | CAMK2A | calcium/calmodulin dependent protein kinase II alpha | KIAA0968,CAMKIIA,CAMKA | GO:0014069 | postsynaptic density (GO:0014069) | CC | 889 |
| CAMK2A | HGNC:1460 | CAMK2A | calcium/calmodulin dependent protein kinase II alpha | KIAA0968,CAMKIIA,CAMKA | GO:0099148 | regulation of synaptic vesicle docking (GO:0099148) | BP | 981 |
| CTTN | HGNC:3338 | CTTN | cortactin | EMS1 | GO:0098871 | postsynaptic actin cytoskeleton (GO:0098871) | CC | 995 |
| CTTN | HGNC:3338 | CTTN | cortactin | EMS1 | GO:0098885 | modification of postsynaptic actin cytoskeleton (GO:0098885) | BP | 1005 |
| SV2B | HGNC:16874 | SV2B | synaptic vesicle glycoprotein 2B | KIAA0735,HST19680,SLC22B2 | GO:0030285 | integral component of synaptic vesicle membrane (GO:0030285) | CC | 1234 |
| APOE | HGNC:613 | APOE | apolipoprotein E | AD2 | GO:0050807 | regulation of synapse organization (GO:0050807) | BP | 1698 |
| PABPC1 | HGNC:8554 | PABPC1 | poly(A) binding protein cytoplasmic 1 | PAB1,PABPC2 | GO:0045202 | synapse (GO:0045202) | CC | 1900 |
| SHANK2 | HGNC:14295 | SHANK2 | SH3 and multiple ankyrin repeat domains 2 | CTTNBP1,PROSAP1,SHANK,SPANK-3,CORTBP1 | GO:0098919 | structural constituent of postsynaptic density (GO:0098919) | BP | 1958 |
| SHANK2 | HGNC:14295 | SHANK2 | SH3 and multiple ankyrin repeat domains 2 | CTTNBP1,PROSAP1,SHANK,SPANK-3,CORTBP1 | GO:0099175 | regulation of postsynapse organization (GO:0099175) | BP | 1962 |
| CAMK2A | HGNC:1460 | CAMK2A | calcium/calmodulin dependent protein kinase II alpha | KIAA0968,CAMKIIA,CAMKA | GO:0098696 | regulation of neurotransmitter receptor localization to postsynaptic specialization membrane (GO:0098696) | BP | 2075 |
| OLFM1 | HGNC:17187 | OLFM1 | olfactomedin 1 | NOE1,OLFA,AMY,NOELIN | GO:0099243 | extrinsic component of synaptic membrane (GO:0099243) | CC | 2185 |
| DAPK1 | HGNC:2674 | DAPK1 | death associated protein kinase 1 | DAPK | GO:0014069 | postsynaptic density (GO:0014069) | CC | 3089 |
| RPS8 | HGNC:10441 | RPS8 | ribosomal protein S8 | ES8 | YNGO:postsyn_ribosom | postsynaptic ribosome | CC | 3243 |
| SV2B | HGNC:16874 | SV2B | synaptic vesicle glycoprotein 2B | KIAA0735,HST19680,SLC22B2 | GO:0099509 | regulation of presynaptic cytosolic calcium levels (GO:0099509) | BP | 3651 |
| CTTN | HGNC:3338 | CTTN | cortactin | EMS1 | GO:0098794 | postsynapse (GO:0098794) | CC | 3710 |
| CTTN | HGNC:3338 | CTTN | cortactin | EMS1 | GO:0099010 | modification of postsynaptic structure (GO:0099010) | BP | 3712 |
| SV2B | HGNC:16874 | SV2B | synaptic vesicle glycoprotein 2B | KIAA0735,HST19680,SLC22B2 | GO:2000300 | regulation of synaptic vesicle exocytosis (GO:2000300) | BP | 3795 |
| SV2B | HGNC:16874 | SV2B | synaptic vesicle glycoprotein 2B | KIAA0735,HST19680,SLC22B2 | GO:0030285 | integral component of synaptic vesicle membrane (GO:0030285) | CC | 3947 |
| OLFM1 | HGNC:17187 | OLFM1 | olfactomedin 1 | NOE1,OLFA,AMY,NOELIN | GO:0099147 | extrinsic component of postsynaptic density membrane (GO:0099147) | CC | 4819 |
| OLFM1 | HGNC:17187 | OLFM1 | olfactomedin 1 | NOE1,OLFA,AMY,NOELIN | GO:0098696 | regulation of neurotransmitter receptor localization to postsynaptic specialization membrane (GO:0098696) | BP | 4820 |
| CSMD2 | HGNC:19290 | CSMD2 | CUB and Sushi multiple domains 2 | KIAA1884 | GO:0099061 | integral component of postsynaptic density membrane (GO:0099061) | CC | 5152 |
| CSMD2 | HGNC:19290 | CSMD2 | CUB and Sushi multiple domains 2 | KIAA1884 | GO:0099562 | maintenance of postsynaptic density structure (GO:0099562) | BP | 5153 |
| CSMD2 | HGNC:19290 | CSMD2 | CUB and Sushi multiple domains 2 | KIAA1884 | GO:0097107 | postsynaptic density assembly (GO:0097107) | BP | 5154 |
